# Mechanisms Generating Cancer Genome Complexity From A Single Cell Division Error

**DOI:** 10.1101/835058

**Authors:** Neil T. Umbreit, Cheng-Zhong Zhang, Luke D. Lynch, Logan J. Blaine, Anna M. Cheng, Richard Tourdot, Lili Sun, Hannah F. Almubarak, Kim Judge, Thomas J. Mitchell, Alexander Spektor, David Pellman

**Affiliations:** Howard Hughes Medical Institute, Chevy Chase, MD, USA; Department of Cell Biology, Harvard Medical School, Boston, MA, USA; Department of Pediatric Oncology, Dana-Farber Cancer Institute, Boston, MA, USA; Department of Biomedical Informatics, Harvard Medical School, Boston, MA, USA; Department of Data Sciences, Dana-Farber Cancer Institute, Boston, MA, USA; Wellcome Sanger Institute, Hinxton, Cambridgeshire, CB10 1SA, UK; Cambridge University Hospitals NHS Foundation Trust, Cambridge, CB2 0QQ, UK; Single-Cell Sequencing Program, Dana-Farber Cancer Institute, Boston, MA, USA; Department of Radiation Oncology, Dana-Farber Cancer Institute, Boston, MA, USA

## Abstract

The chromosome breakage-fusion-bridge (BFB) cycle is a mutational process that produces gene amplification and genome instability. Signatures of BFB cycles can be observed in cancer genomes with chromothripsis, another catastrophic mutational process. Here, we explain this association by identifying a mutational cascade downstream of chromosome bridge formation that generates increasing amounts of chromothripsis. We uncover a new role for actomyosin forces in bridge breakage and mutagenesis. Chromothripsis then accumulates starting with aberrant interphase replication of bridge DNA, followed by an unexpected burst of mitotic DNA replication, generating extensive DNA damage. Bridge formation also disrupts the centromeric epigenetic mark, leading to micronucleus formation that itself promotes chromothripsis. We show that this mutational cascade generates the continuing evolution and sub-clonal heterogeneity characteristic of many human cancers.

## INTRODUCTION

Cancer genomes can contain thousands of chromosomal rearrangements (1). Traditionally, it was assumed that these genomes evolve gradually by accruing small-scale changes successively over many generations. However, the extent of genomic rearrangement in many cancers suggests a non-exclusive, alternative view: these genomes may evolve rapidly via discrete episodes that generate bursts of genomic alterations (2–6). This latter model provides a parsimonious explanation for the origin of extreme genomic complexity.

Three classes of catastrophic events have been described that may account for a substantial fraction of chromosome alterations in cancer: whole-genome duplication, chromothripsis, and chromosome breakage-fusion-bridge cycles. The first class, whole-genome duplication (WGD), can promote tumorigenesis and is now appreciated to occur during the development of ∼40% of human solid tumors (2, 7). Whole-genome duplication causes genome instability by several mechanisms, including doubling the number of centrosomes, distorting spindle architecture, generating chromosome segregation errors, and producing micronuclei, abnormal nuclear structures common in cancer (8–11).

The second class of catastrophic event, chromothripsis, is a massive rearrangement of only one or a few chromosomes resulting in an unusual DNA copy number pattern (3, 5, 12). Chromothripsis occurs with reported frequencies of 20–65% in many common tumor types (1, 13). We previously found that chromothripsis can originate from micronuclei (14–17), which arise from mitotic segregation errors or unrepaired DNA breaks that generate acentric chromosome fragments. Due to aberrant nuclear envelope assembly around these chromosomes, micronuclei undergo defective DNA replication and spontaneous loss of nuclear envelope integrity, which results in extensive DNA damage by unknown mechanisms (18, 19).

The third class of catastrophic event, the chromosome breakage-fusion-bridge (BFB) cycle (20, 21), starts with cell division errors that trigger the formation of another abnormal nuclear structure, a chromosome bridge. Bridges arise from telomere crisis and end-to-end chromosome fusions but can also occur from end fusions at DNA breaks, incomplete DNA replication, or a failure to resolve chromosome catenation (22). Bridge breakage then initiates a process that can generate gene amplification after multiple cell generations. Evidence of BFB cycles has been reported in a broad range of cancer types, with estimated frequencies ranging up to ∼80% in pancreatic cancer (13).

Although BFB cycles are major sources of genome instability, the perfect palindromic sequence pattern expected from the originally proposed BFB model is not commonly observed in cancer genomes (1, 13, 23). Whether this is due to subsequent chromosomal rearrangement obscuring the simple BFB pattern, or whether the BFB process itself is inherently more complex than originally envisioned, has been unclear. Recently, comprehensive cancer genome sequencing has uncovered examples where signatures of the BFB cycle are intermingled with chromothripsis, raising the possibility that coupling of these two processes could add complexity to BFB cycles (23). However, to determine the relationship between BFB cycles and chromothripsis, it is first necessary to better understand the BFB cycle, for which many key mechanistic steps, particularly how chromosome bridges are broken, remain unclear.

Proposed models for chromosome bridge breakage have included breakage by spindle forces during the mitosis in which they are formed or DNA cleavage by the cytokinesis/abscission apparatus (21, 24–26). Yet recent work indicates that chromosome bridges are rarely, if ever, broken during mitosis or cytokinesis, but rather persist for many hours into interphase (26, 27). It was then proposed that interphase bridges are severed by the cytoplasmic, endoplasmic reticulum-associated exonuclease, TREX1 (26). Transient nuclear envelope (NE) disruption was suggested to allow TREX1 to enter the nucleus and gain access to the bridge DNA, simultaneously breaking the bridge and fragmenting the bridge DNA to generate chromothripsis (26). Although the TREX1-model could explain the association between BFB cycles and chromothripsis in cancer genomes (23), loss of TREX1 was reported to delay, but not block, bridge breakage (26).

Below, we present data supporting a new model that explains the linkage between BFB cycles and chromothripsis. Rather than being generated simultaneously by a single mechanism, we demonstrate that chromothripsis accumulates through a cascade of new mutational events initiated by the formation of a chromosome bridge. The first such event appears to involve defective DNA replication of bridge DNA, which, in a minority of cells, is associated with a newly-identified signature of DNA rearrangement in broken bridges during the interphase after bridge formation. Next, we observed extensive DNA damage and frequent chromothripsis associated with an unexpected burst of aberrant DNA replication on broken bridge “stubs” during the next mitosis. Then, because of compromised maintenance of the centromere epigenetic mark, CENP-A, the chromosomes with the broken bridge stubs mis-segregate with high frequency into micronuclei, which will generate further rounds of chromothripsis. Analysis of clonal populations after bridge breakage established that these events initiate iterative cycles of genome instability, causing extensive subclonal heterogeneity downstream of the formation of a single chromosome bridge. An analogous series of events was observed after the formation of micronuclei, indicating that similar mechanisms generate chromothripsis irrespective of the nuclear structure initiating the mutational cascade. Together, these findings reveal how a single cell division error rapidly generates extreme genomic complexity.

## RESULTS

We used four methods to generate chromosome bridges to study their breakage and genomic impact: transient expression of a dominant negative variant of telomeric repeat-binding factor 2 (TRF2-DN) (28), partial knockdown of condensin (siSMC2) (29), low-dose topoisomerase II inhibition (ICRF-193) (30), and CRISPR/Cas9-mediated telomere loss on chromosome 4 (Chr4g1, Fig. S1A-C). TRF2-DN was employed to generate chromosome bridges in most experiments in this study, unless otherwise specified. Chromosome bridges were visualized in live cells with GFP-BAF (barrier-to-autointegration factor). GFP-BAF is a sensitive reporter for these structures because, unlike histones (26), DNA binding by BAF is not compromised by the stretching of chromosome bridges during bridge extension (Fig. 1). For TRF2-DN, we developed transient expression and live-cell imaging conditions that avert the previously reported strong inhibition of cell cycle progression (26). In our conditions, cells with bridges entered S phase with similar timing after mitotic exit as unperturbed parental cells lacking bridges (8.3 versus 7.3 hr, respectively; Fig. S1D and accompanying legend). Importantly, bridges generated by each of the above four approaches all had similar lifetimes (t_1/2_): ∼10 hours from the completion of mitosis (Fig. 1A).

**Figure 1.**
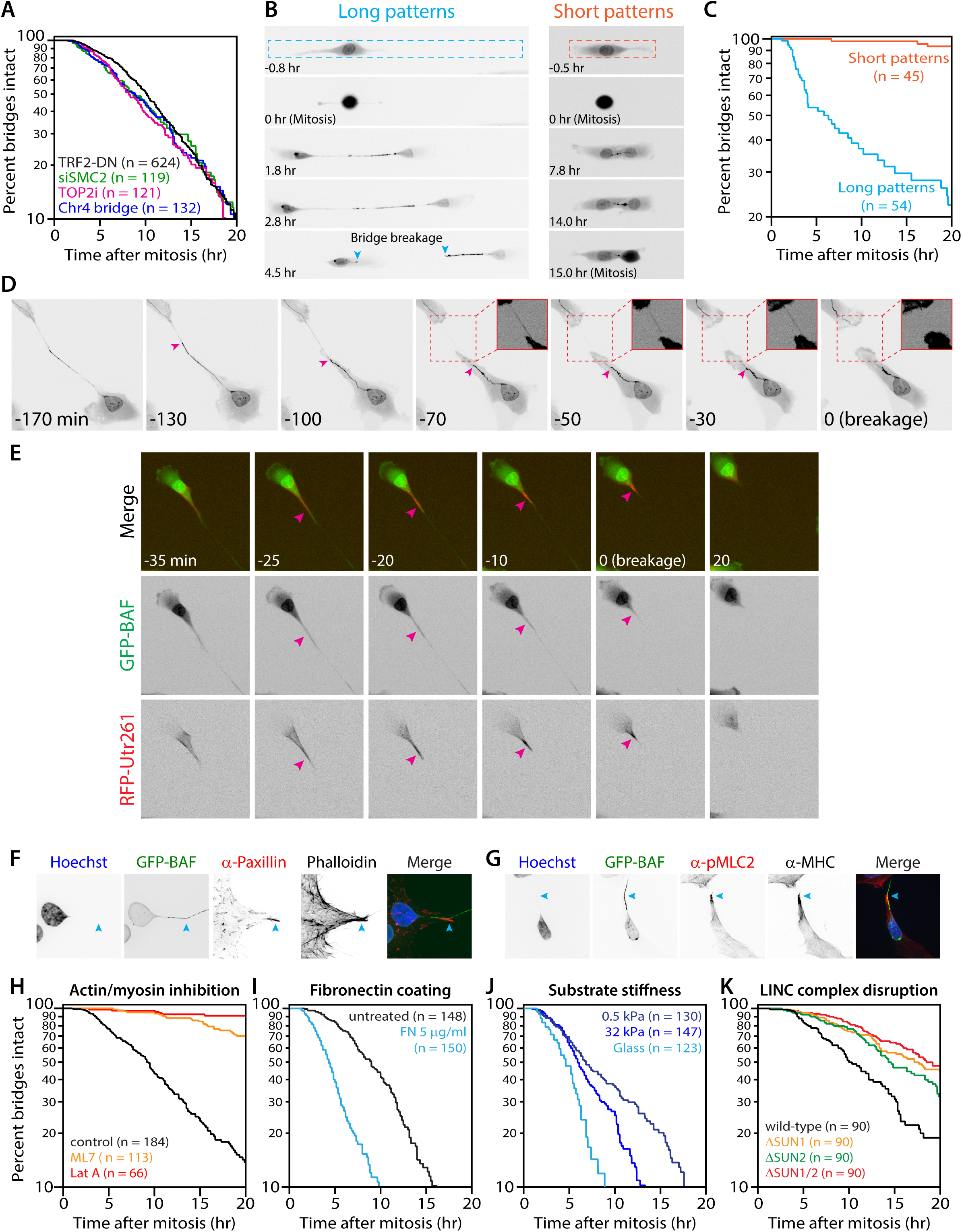
Chromosome bridge breakage requires actomyosin contractility. (A) Indistinguishable chromosome bridge lifetimes observed with different experimental methods for bridge induction. Survival plot shows bridge lifetimes (time interval from bridge formation after mitosis until bridge breakage or the next mitosis; visualized with GFP-BAF). Methods of bridge induction were: inducible dominant-negative TRF2 (black, n = 624 bridges analyzed), partial knockdown of condensin (siSMC2, green, n = 119), low-dose topoisomerase II inhibition (100 nM ICRF-193, magenta, n = 121), and inducible CRISPR/Cas9-targeted telomere loss on chromosome 4 (Chr4g1, blue, n = 132). No significant difference in mean lifetime is found among the four methods (*p* = 0.14, one-way ANOVA). (B) Extension of chromosome bridges is required for their breakage. Representative time-lapse images (GFP-BAF) of cells with bridges on “long” (20×300 μm, left) or “short” (20×100 μm, right) fibronectin micropatterns. Note: due to the area occupied by the main cell body of each daughter, bridge length does not exceed ∼50 µm on short patterns. Dashed lines: micropattern borders; teal arrowheads: broken bridge ends. Timestamp is relative to completion of the previous mitosis (0 hr). (C) Quantification of data from (B). Orange and teal traces show data for cells on short (n = 45) and long (n = 54) micropatterns; *p* < 0.0001 (Mann-Whitney). (D) A representative chromosome bridge breakage event. Prior to breakage, there is apparent non-uniform stretching of the bridge (GFP-BAF). Magenta arrowhead indicates a transition between “taut” and “slack” regions of the bridge. The taut region appears to progressively stretch, whereas the slack region progressively retracts; breakage occurs in the taut region. Inset images: high contrast of the regions marked by dashed red boxes to visualize the taut region before and after breakage. Timestamp is relative to bridge breakage (min). (E) Actin dynamics during chromosome bridge breakage. Actin is detected with RFP-Utr261; bridges with GFP-BAF. Contraction of the actin-rich structure (magenta arrowheads) occurs immediately preceding and up to the time of bridge breakage (−25 to 0 min). After bridge breakage, the actin structure rapidly disassembles (20 min). Timestamp is relative to bridge breakage (min). (F) Representative images show large focal adhesions (α-Paxillin) and actin fibers (phalloidin) at the bent region of a chromosome bridge (GFP-BAF), indicated by cyan arrowheads. (G) As in (B), representative accumulation of contractile myosin II (α-myosin heavy chain, MHC; α-phospho-myosin light chain 2, pMLC2) at the transition between taut and slack segments of a chromosome bridge. (H) Actomyosin contractility is required for bridge breakage. Bridge lifetime plots show the effect of actin disruption (0.5 μM Latrunculin A; red trace, n = 66) or myosin II inhibition (20 μM ML7; orange trace, n = 113) relative to controls (DMSO; black trace, n = 184). Cells were allowed to divide, form and extend chromosome bridges and were then exchanged into drug medium (see Fig. S5B). (I) Increasing cellular contractility decreases bridge lifetime. Bridge lifetimes for cells plated on untreated glass (black, n = 148) or fibronectin (FN)-coated glass (light blue, n = 150). (J) Bridge breakage depends on substrate stiffness. Bridge lifetimes were measured for cells plated on substrates of varying stiffness, each coated with 5 μg/ml fibronectin: glass (>10^6^ kPa; light blue, n = 123), stiff gel (32 kPa; medium blue, n = 147), and soft gel (0.5 kPa; dark blue, n = 130). (K) Bridge breakage in part requires the LINC complex. Bridge lifetimes for wild-type (black, n = 90), SUN1 knockout (orange, n = 90), SUN2 knockout (green, n = 90), and SUN1/SUN2 double knockout (red, n = 90) RPE-1 cells.

### Mechanical force triggers chromosome bridge breakage

The cytoplasmic, endoplasmic reticulum-associated exonuclease, TREX1, has been suggested to mediate chromosome bridge breakage after rupture of the primary nucleus in bridged cells, as genetic ablation of TREX1 had a partial effect on bridge resolution (26). However, using the same cell lines and bridge induction method (and an additional method) but imaging conditions with less light exposure, we were unable to detect an effect of TREX1 knockout on bridge lifetime (six independent clones, two different knockout strategies, Fig. S2A-C). Additionally, a notable fraction of bridge breakage events occurred in the absence of any detectable rupture of the primary nucleus (36%, n = 58, Movie S1), and bridge lifetime showed no correlation with the occurrence or duration of nuclear envelope disruption (Fig. S2D). Therefore, fundamental aspects of the mechanism for bridge breakage remain to be identified.

A clue for alternative mechanisms comes from the fact that as interphase cells migrate in culture, bridges can reach hundreds of microns in length before breaking, raising the possibility that bridge breakage might have a mechanical component. Consistent with this idea, BJ foreskin fibroblasts, which exhibited similar motility to RPE-1 cells (cell velocity 0.48 and 0.51 µm/min, respectively), extended and broke chromosome bridges during interphase with similar timing as bridges in RPE-1 cells (t_1/2_ = 7.4 hr; Fig. S3A and Movie S2). By contrast, two cell lines that exhibited low motility (HeLa and U2OS) almost never extended bridges beyond 100 µm and rarely underwent breakage before the next mitosis (10% and 20% interphase breakage, respectively; Fig. S3B-C and Movies S3-4).

We hypothesized that the extension of chromosome bridges is required for their breakage. To test this idea, we controlled bridge extension using rectangular fibronectin “micropatterns” that constrain cell migration to the fibronectin-containing pattern (31). When RPE-1 cells were plated on long (300 µm) patterns, newly formed chromosome bridges extended to ∼160 μm on average, and ∼85% of bridges broke during interphase with similar kinetics as unconfined cells on glass coverslips (Fig. 1B-C, Movie S5). By contrast, restricting bridge extension with short (100 µm) micropatterns limited bridge extension to <50 μm and almost completely blocked bridge breakage (<10% bridge cleavage prior to entry into the next mitosis; Fig. 1B-C, Movie S6). Although there was less spontaneous NE rupture on short patterns, increasing NE ruptures >8-fold with Lamin B1 knockdown failed to accelerate bridge breakage on short patterns (Fig. S4). Therefore, the extension of chromosome bridges, but not NE rupture, is required for their breakage.

Mechanical forces could stretch a bridge across its length or act locally within a section of a bridge. Live-cell imaging supported the latter model: bridges often formed acute angle bends and/or exhibited non-uniform stretching prior to breakage, with one segment appearing taut and adjacent segments appearing slack, followed by breakage of the bridge within the taut segment (23 of 25 cases examined, Fig. 1D, Movie S7). Supporting the idea that local actomyosin contractile forces contribute to bridge breakage, live cell imaging with the actin reporter RFP-Utr261 (32) revealed large concentrations of actin filaments immediately adjacent to the taut segments of the bridge just prior to breakage in all cases examined (n = 30; Fig. 1E, Fig. S5A, Movies S8-9 and see (27) for similar results). Actin accumulation was transient and dissolved after bridge breakage. Immunostaining to detect paxillin revealed large focal adhesions at sites of F-actin accumulation where bridges appeared taut, indicating strong cell-extracellular matrix attachments (Fig. 1F). These sites also had extensive accumulations of non-muscle myosin II, which co-stained for the active, phosphorylated form of myosin regulatory light chain, indicating high contractile forces (Fig. 1G). Local myosin accumulation and contractility is known to be induced by increased membrane tension (33), which is expected to occur at the base of extending chromosome bridges.

We next asked if actomyosin contractility is required for chromosome bridge breakage. During live imaging, chromosome bridges were generated and allowed to extend, and then small-molecule inhibitors of myosin activation (ML7) or actin assembly (Latrunculin A) were added. By comparison with controls, ML7 addition substantially delayed, and Latrunculin A addition abolished, bridge breakage (Fig. 1H, Fig. S5B, and Movie S10) demonstrating that a functional actomyosin network is required for bridge breakage. Moreover, when cells were plated on fibronectin, which increases focal adhesions and intracellular actomyosin contractile forces (34), bridge breakage was accelerated two-fold (*p* < 0.0001; Fig 1I). Because fibronectin also affects cell signaling (35, 36), we plated cells on hydrogels of varying stiffness, all coated with the same concentration of fibronectin. Consistent with the known effect of reduced substrate stiffness diminishing actomyosin contractility (37), bridge lifetime increased with decreasing substrate stiffness (Fig. 1J). Finally, we asked whether bridge breakage depends on the LINC complex, the best characterized pathway by which actomyosin forces can be transmitted across the nuclear envelope (38–40). Knockout of the major inner nuclear membrane LINC components, SUN1 and SUN2, had a partial effect, delaying bridge breakage (t_1/2_ = 18 hr; Fig 1K and Fig. S6). Together, these data establish a critical role for cytoplasmic actomyosin contractile forces in chromosome bridge breakage.

### Single-cell sequencing reveals the immediate impact of chromosome bridge breakage

#### Copy number alterations immediately after bridge breakage

We investigated both the immediate and long-term consequences of chromosome bridge breakage on genome structure. To define the immediate outcome(s) of bridge breakage, we employed our previously developed approach, which combines live-cell imaging with single-cell whole-genome sequencing [Look-Seq (17)]. Chromosome bridges were induced, their breakage was monitored by live-cell imaging, and the two daughter cells were isolated ∼8 hr after bridge breakage for single-cell whole genome sequencing. Sequencing was performed to ∼25× mean depth, which allowed us to interrogate ∼90% of the unique sequence of each homologous chromosome with one or more reads (Fig. 2A, Methods). This approach enabled us to observe the immediate consequences of bridge breakage without confounding genomic alterations during subsequent cell divisions.

**Figure 2.**
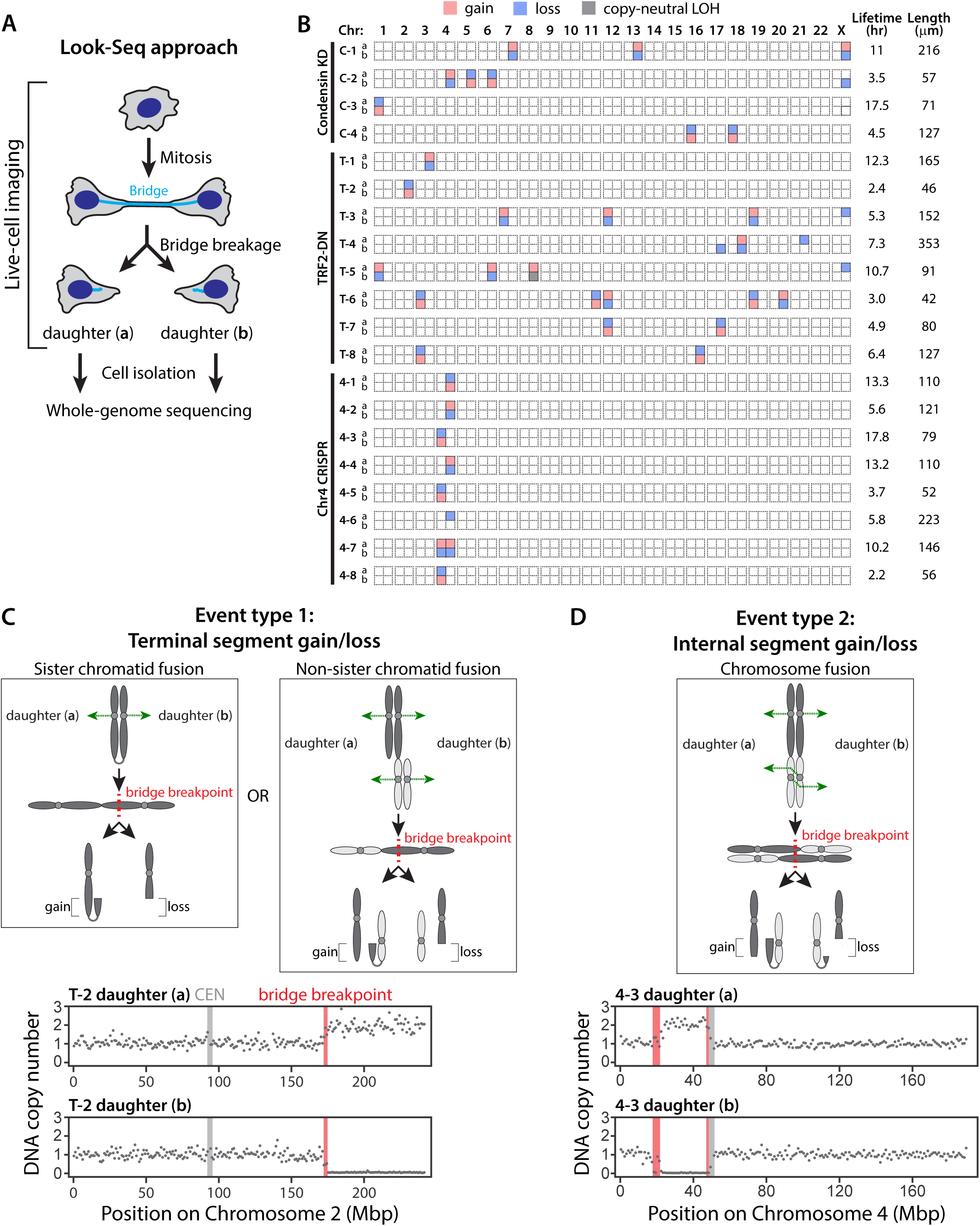
Immediate effect of chromosome bridge breakage on DNA copy number. (A) Cartoon illustrating the Look-Seq experiment. Bridge formation and breakage was monitored during live imaging. After bridge breakage, individual daughter cells were isolated for whole-genome sequencing. (B) Schematic summary of large-scale (≥2.5 Mb) DNA copy number alterations after bridge breakage. Dashed boxes show the p- and q-arms of each chromosome. Chromosome arms that contain a subregion with copy number alterations are colored as follows: white, diploid; red, gain; blue, loss; gray, copy-neutral loss of heterozygosity. Right: bridge lifetime and bridge length at the time of breakage for each sample. (C) Type 1 events are daughter cells with reciprocal gain and loss of a terminal chromosome segment. Cartoon depicts chromatid fusion events initiated by DNA breaks or telomere uncapping. Left: sister chromatid fusion; Right: fusion of single chromatids from different chromosomes (G2 cell). The resulting dicentric fusions are segregated in mitosis (green dashed arrows) to form a bridge. Breakage of the bridge (dashed red line) generates the depicted reciprocal copy number alterations. Bottom: representative plot of DNA copy number (gray dots show mean copy number of 1-Mb bins) for the affected haplotype resulting from a Type 1 bridge breakage event involving the q-arm of chromosome 2. Red bar: inferred bridge breakpoint. Light gray bar: centromere. (D) Type 2 events are reciprocal gain and loss of an internal chromosome segment between the daughter cells. Top: cartoon depicts a chromosome fusion (28) (e.g. from replication of an interchromosomal fusion occurring in G1). If the kinetochores of the dicentric chromatids attach to microtubule bundles from opposite poles (dashed green arrows), one dicentric will assume an inverted orientation relative to the other (middle panel). In contrast to the scenarios depicted in (C), cleavage of both chromatids in the resulting bridge at the indicated position yields reciprocal copy number alterations of an internal chromosome segment. Bottom, plot of DNA copy number as in (C).

In all 20 cell pairs after bridge breakage, we observed copy number alterations affecting a segment (>2.5 Mb) of one or more chromosome arms, distributed in a reciprocal pattern between the daughter cells (Fig. 2B, Fig. S7, and Movies S11-13). Using previously developed haplotype copy number analysis, we could unambiguously identify the homologous chromosome that underwent breakage (17). Most commonly, we found terminal segment reciprocal gain and loss patterns, which, as in the original BFB model (21), are expected from breakage of dicentric fusions between sister chromatids or single chromatids from different chromosomes (“chromatid fusions,” Fig. 2C). Interestingly, in four daughter cell pairs, we observed the reciprocal gain and loss of internal chromosome segments. This pattern can be explained by breakage of replicated dicentric chromosomes derived from two different chromosomes (“chromosome fusions,” (41), Fig. 2D), coupled to an inverted orientation of the dicentric chromatids along the mitotic spindle. Although bridge breakage sometimes affected only one chromosome, in nine cases, two or more different chromosomes were involved, as expected from the methods employed to induce bridges (41); the exception was the CRISPR-based method, which, as expected, exclusively produced chromosome 4 bridges (Fig. 2B).

Closer inspection of the bridge breakpoints revealed a spectrum of genomic outcomes, where some bridges underwent simple breakage and others experienced fragmentation, which was specifically localized to the region of the main copy number transition (Fig. 3). In cases where bridge breakage occurred with local fragmentation, fragments as small as ∼100 kb could be readily detected (see Methods for details) if these fragments were retained within a larger region of complete haplotype loss. The retention of larger (∼1Mb) fragments in one daughter could also be validated by reciprocal loss of this fragment in the sister cell. Rearrangements involving fragment ends often provided additional support for the identification of fragmentation.

**Figure 3.**
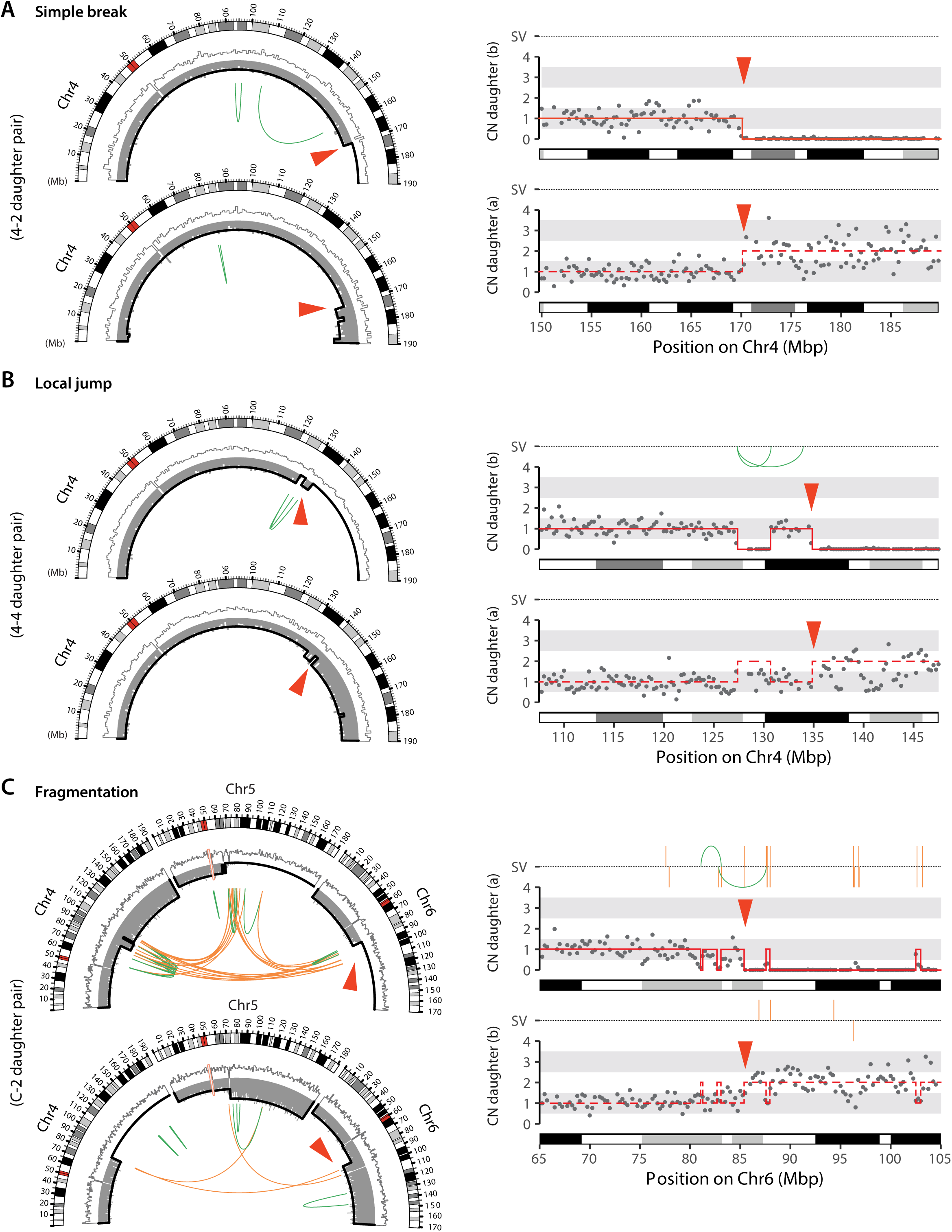
Small-scale, highly localized DNA breakage and rearrangement with bridge breakage. (A) Simple breakage of a bridge chromosome. Left: CIRCOS plots showing the bridge chromosome (Chr4) in the 4-2 daughter pair (Fig. 3B). DNA copy number is shown for the bridge haplotype (gray bars/black outline) and the non-bridge haplotype (white bars/grey outline); intrachromosomal rearrangements are shown as green arcs, chromosome band pattern is shown in the outer arc. Red arrowhead indicates the bridge breakpoint. Right: Expanded view of DNA copy number at the breakpoint transition (gray dots show 250-kb bins). Copy-number segments (red solid lines) are determined using SNP-level coverage in the top daughter (see Methods). The reciprocal pattern is shown for the bottom daughter (red dashed lines). This line represents the expected copy number in the bottom daughter based on fragmentation detected in the top daughter, assuming that two copies of the shown homolog were distributed between both daughters. Structural variants (SV) are shown above the copy-number plots as in the CIRCOS plots. (B) Generation of the “local jump” pattern from bridge breakage. As in (A), shown are CIRCOS plots (Left) and DNA copy number plots with rearrangements (Right) for the bridge chromosome (Chr4) in the 4-4 daughter pair. (C) Local fragmentation with complex rearrangement associated with bridge breakage. As in (A), CIRCOS plots (Left) and DNA copy number plots with rearrangement (Right) in the C-2 daughter pair, whose bridge contained three different chromosomes (Chrs 4, 5, and 6). Each bridge chromosome contains multiple breaks resulting in local fragmentation. The pattern of rearrangements in daughter (b) indicates end-joining of these fragments, including the formation of inter-chromosomal rearrangements (orange arcs or lines). In addition to fragmentation, daughter (a) evidences the TST jump rearrangement pattern (see Fig. 5).

Importantly, both simple breaks and local fragmentation could be generated by mechanical force-dependent bridge breakage, because direct breakage of bridges with a glass capillary yielded a very similar spectrum of outcomes (Fig. 4A and Fig. S8A). Moreover, we observed similar local fragmentation patterns for spontaneous bridge breakage in TREX1-null cells, reinforcing the conclusion that TREX1 is not required to break or fragment chromosome bridges (Fig. 4B and Fig. S8B).

**Figure 4.**
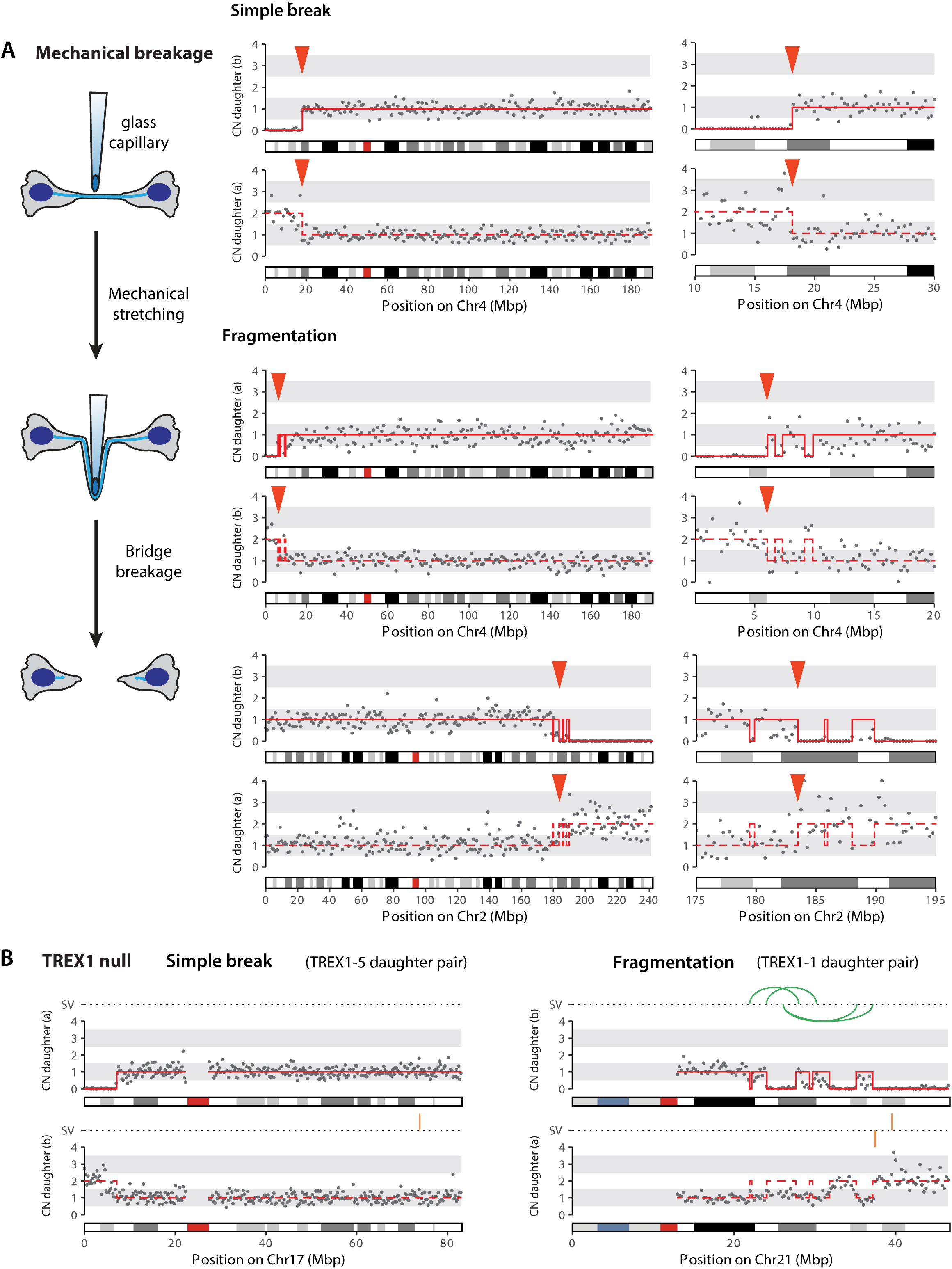
Local fragmentation accompanies mechanical breakage of bridges and does not require TREX1. (A) Mechanical breakage of chromosome bridges produces a spectrum of outcomes from simple breakage to local fragmentation. Left: schematic of the experiment using a glass capillary to mechanically stretch and break chromosome bridges. The daughter cells were collected for sequencing immediately after mechanical bridge breakage, to determine its direct consequences. Therefore, cells did not have time for DNA repair to generate chromosomal rearrangements. Top right: an example of simple bridge breakage. Plots as in Fig. 3, with the exception that grey dots in the whole-chromosome plot indicate 1 Mb bins due to lower (5×) sequence depth. Bottom right: two examples of local fragmentation. (B) A similar spectrum from simple breakage to local fragmentation after spontaneous bridge breakage in cells lacking TREX1. Left: DNA copy number plots, as in (A), showing simple bridge breakage in TREX1-null cells. Right: local fragmentation of the bridge chromosome, with chromosome rearrangements (green arcs and orange lines) due to end-joining of the resulting fragments.

In sum, these findings demonstrate that the immediate consequences of bridge breakage are relatively simple patterns of copy number alterations localized to the bridge. This localized pattern contrasts with that reported from bulk sequencing of populations of cells isolated many generations after telomere crisis. These populations often contained complex copy number oscillations and rearrangements that encompassed most of a chromosome arm and/or spanned the centromere (26). We also observed these complex patterns in long-term population evolution experiments and will present evidence defining a cascade of events downstream of initial bridge breakage that can explain them (see Figs. 7-9 below).

#### Chromosome rearrangements associated with bridge breakage

We next analyzed chromosome rearrangements associated with the above described DNA copy number alterations. Many cell pairs exhibiting simple breakage or small-scale fragmentation contained the approximate number of rearrangements expected from ligation of the fragments (Fig. 3B). This pattern of rearrangements closely resembles what has been termed “local jump footprints,” a rearrangement signature in cancer genomes of unknown origin (42). We also identified several examples of local fragmentation involving two or more chromosomes, where subsequent end-joining produced a pattern of intra- and inter-chromosomal rearrangements (Fig. 3C, bottom cell). Overall, these findings suggest that “local jumps” can be generated by DNA ligation after local fragmentation, and thus may share a common underlying mechanism with many cases of chromothripsis, consistent with our previous proposal (17).

Four daughter cell pairs showed a distinct and particularly extreme pattern of complex rearrangement (n = 4 of 20 cell pairs; Fig. 5A). In two of these cases we additionally observed kataegis near the rearrangements. Kataegis is a phenomenon in which local clusters of point mutations are generated in a strand-coordinated manner and in trinucleotide contexts implicating the action of APOBEC family cytosine deaminases on single-stranded DNA (Fig. S9) (43, 44). Thus, complex rearrangements and kataegis can occur at or around the time of chromosome bridge breakage, albeit in a minority of cases.

**Figure 5.**
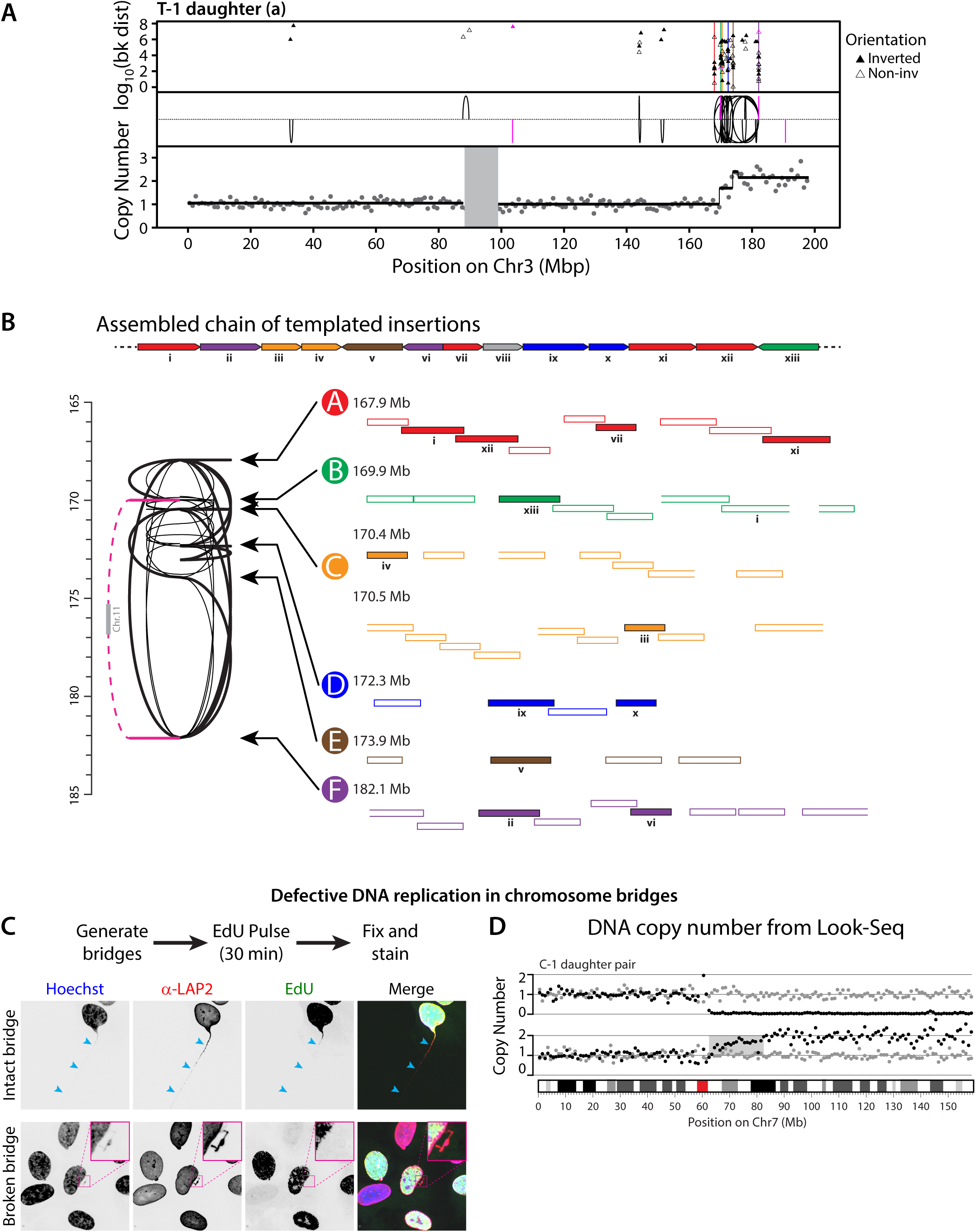
The Tandem Short Template (TST) jump rearrangement signature and aberrant DNA replication within broken chromosome bridges. (A) Extreme breakpoint clustering near the site of chromosome bridge breakage. Copy number (1-Mb bins, gray dots) and rearrangements (black curves and magenta lines) are shown as in Fig. 3 for daughter cell (a) of the T-1 sample. Copy-number segmentation based on the 250-kb bin-level analysis (see Methods) is shown (bottom, black line). Uppermost (“rainfall”) plot shows the distance between adjacent breakpoints (log_10_ scale); clustering of breakpoints is indicated from marked drops in inter-breakpoint distance (colored lines correspond to rearrangement hotspots in panel B). (B) Features of the TST jump signature. Top: One chain of short insertions, colored according to their respective hotspot origin. The origin of each insertion in this chain is labeled in the hotspot plot below, showing its order in the template chain (i, ii, iii, etc.). Bottom left: The chain of insertions depicted above is shown as thick black lines, and other chains are thin black lines. Magenta line shows a short insertion derived from Chr11. Bottom right: Each hotspot (A-F) spans 1-2 kb and contains 4-8 short insertions (8-16 break ends); numbers indicate the position of each hotspot on the reference genome. The TST insertions (not drawn true-to-scale) within each hotspot often display partial overlap and form multiple chains. Filled shapes: TST insertions in the single, long chain shown above. Unfilled shapes: other TST insertions involved in different chains. Shapes open on one side represent the start or end of chains, where only one segment boundary could be determined via SV analysis. (C) Cytological observation of under-replication of chromatin in bridges. Cells with intact (top) or broken (bottom) chromosome bridges were pulse-labeled in S phase with EdU. Bridges, marked by cyan arrowheads, were visualized by staining for LAP2. Insets (magenta boxes) show the broken bridge stub. (D) Example of interphase under-replication of DNA in bridges detected by single-cell sequencing. Cells were isolated after a Look-Seq experiment (Fig. 3A). Shown are copy number plots for the bridge haplotype (black dots) and the control, non-bridge haplotype (gray dots). Gray shading: region of under-replication of bridge haplotype. The mean copy number in this 20-Mb region (grey rectangle) for the bridge haplotype, 1.56, is lower than the expected gain (CN = 2) for this region. Partial retention of that haplotype in the sister cell (median CN of bridge haplotype = 0.05) does not explain the extent of “missing” DNA.

Analysis of rearrangement junctions from these four samples revealed surprising features that are inconsistent with an origin from simple fragmentation followed by ligation in random order and orientation. Instead, the rearrangement pattern suggests it originates from errors during DNA replication. First, rather than being randomly distributed, in these samples, breakpoints were tightly clustered into local hotspots, as has been previously noted in chromothripsis samples but not understood mechanistically (45). Second, tracking the connections between rearrangements revealed chains of short insertions (median 183 bp) that were arrayed in tandem, hereafter referred to as “Tandem Short Template” (TST) jumps (Fig. 5B). The TST insertions were typically derived from multiple hotspots on chromosomes within the bridge, but occasionally, insertions originated from other chromosomes not involved in the bridge. One plausible explanation is that these insertions were generated by template-switching DNA replication errors, as in the microhomology-mediated break-induced replication (MMBIR) model (12, 46).

Accordingly, we analyzed microhomology at the junctions between TST insertions. Although a minority of junctions showed blunt-end joining (≤ 1bp microhomology), junctions with microhomology were also infrequent. For example, of the 13 junctions in the TST chain shown in Fig. 5B, five contained microhomology or insertion of ≤1 bp, and two showed ≥2 bp microhomology. The remaining six junctions contained 2–20 bp of sequence with ambiguous origin. It is possible that these sequences reflect junctional microhomology, but that detection of microhomology is obscured because the sequences are derived from repeats and/or partial mismatches that are difficult to map (47).

In light of the above findings implicating aberrant DNA replication in the generation of the TST jump signature, we characterized the efficiency of DNA replication in chromosome bridges. Pulse-labeling with the nucleoside analog, 5-ethynyl-2′-deoxyuridine (EdU) was used to assess replication in chromosome bridges. Cells in S phase labeled strongly for EdU in the primary nucleus, but EdU intensity dropped off where chromatin extruded from the main nucleus and was absent from most of the bridge (Fig. 5C). Control experiments demonstrated that the absence of EdU signal was not due to limited detection sensitivity for the small amount of DNA in bridges (Fig. S10). The same defect was observed in intact bridges and broken bridge stubs (Fig. 5C), indicating that both structures have replication defects. Further support for the conclusion that bridge DNA is poorly replicated came from our single-cell sequencing experiments. In approximately half of the daughter cell pairs, we could identify a region of the bridge chromosome that was present at lower copy number than the intact homologue that was not in the bridge (Fig. 5D). Therefore, chromosome bridges exhibit severe DNA replication defects similar to those previously identified in micronuclei (14, 16, 19).

We observed the TST jump signature in two additional contexts by bulk DNA sequencing. First, we identified the TST jump signature by bulk sequencing of a population of cells derived from a single cell with a broken bridge. We use CRISPR to induce breakage of the termini of Chr4, observed the formation and breakage of chromosome bridges and then isolated individual cells that were then grown into large populations (>10^6^ cells each). One out of 12 such populations evidenced clustering of rearrangements into breakpoint hotspots, with chains of short insertions with a similar size range as the single cell experiments (Fig. 6A). Second, we identified the TST jump signature by long-read sequencing of a primary tumor sample obtained from a patient with renal cell carcinoma. In this patient sample, the TST jumps are associated with a chromothripsis event that generated the unbalanced translocation between Chr3p and Chr5q (Fig. 6B), which is the canonical driver event in this cancer type (48). The median size of the insertions (199 bp) was similar to what we observed by single-cell sequencing of broken bridges (Fig. 6C). These findings indicate that the TST jump signature reflects a specific mutational process that can be stably inherited over many generations.

**Figure 6.**
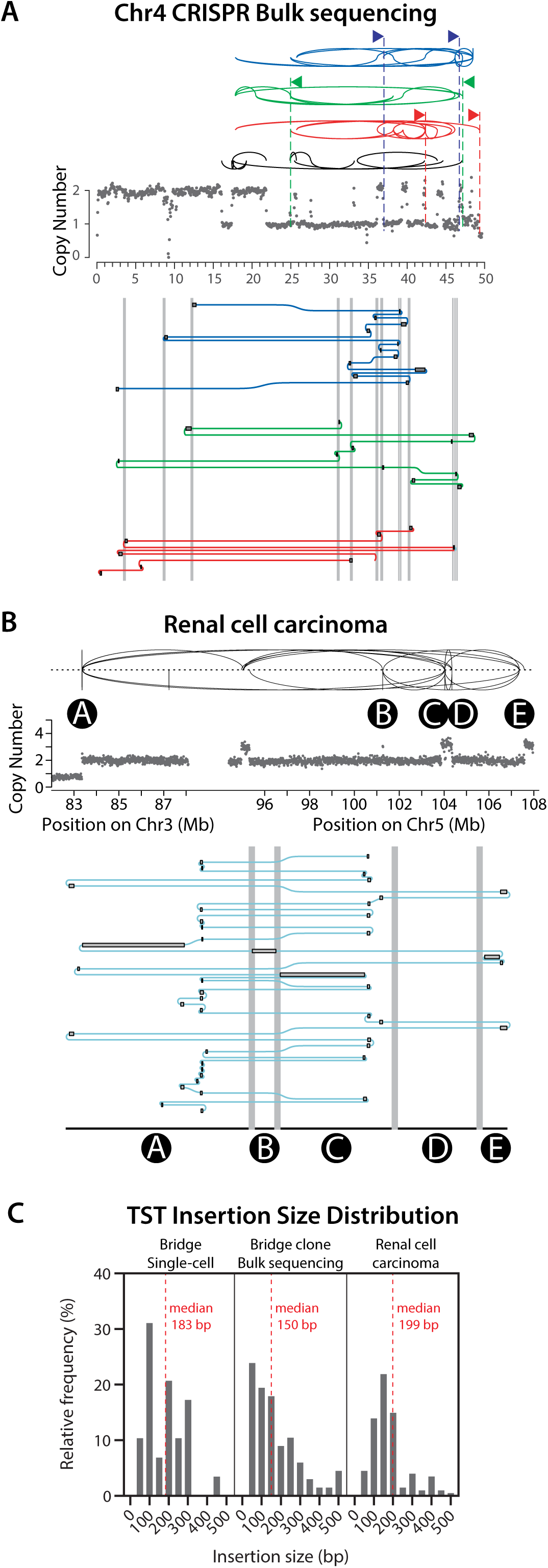
The Tandem Short Template (TST) jump rearrangement signature in primary clones from bridged cells and in a primary tumor sample. (A) The TST jump signature is observed in bulk sequencing of the progeny of a single cell after bridge breakage. Top: DNA copy number of the bridge chromosome (Chr4) is shown as gray dots (250-kb bins). Long-range rearrangements (distance between breakpoints > 1Mb) on Chr4 are shown as black or colored curves. Three chains of complex rearrangements (similar to the chain illustrated in Fig. 5B), each consisting of 8-14 short templates originating from 11 breakpoint hotspots, are shown with blue, green, and red curves. Black curves indicate rearrangements not obviously linked in chains. Bottom: the chains of insertions are shown schematically, where templated insertions (gray boxes with black outline) are connected as shown by blue, green, or red lines in an expanded view for each breakpoint cluster (≤10-kb window in each region). Grey vertical lines are axis breaks indicating distances larger than 10 kb. (B) TST jump signature in a renal cell carcinoma sample. As in (A), upper plot shows copy number (gray dots: 10-kb bins) and rearrangements (black lines) for the region of unbalanced translocation between Chr3 and Chr5. Schematic below depicts one chain of templated insertions from long-read sequencing data, in an expanded view for each breakpoint cluster (3- to 10-kb windows in the hotspot regions labeled A-E). (C) Chains of short insertions identified in bridged cells and in a primary tumor sample exhibit a similar fragment size distribution. Histograms show the size distribution for chained short insertions from single-cell sequencing of a daughter cell after bridge breakage (left; data from Fig. 5B), from bulk sequencing of progeny derived from a single cell after bridge breakage (center; data from panel A), and from long-read sequencing data from the renal cell carcinoma sample (right; data from panel B).

In summary, sequencing cells after the breakage of chromosome bridges demonstrates that most rearrangements result from ligation after localized fragmentation but that highly complex rearrangements do occur in a minority of cases. The unique sequence features of these rearrangements (TST jumps) suggest an origin from template-switching errors in DNA replication (12, 47, 49, 50).

### Mechanisms generating DNA damage downstream of chromosome bridge breakage

#### Damage from aberrant mitotic DNA replication

Although there is a low frequency of complex rearrangement initially associated with chromosome bridges in the first interphase after the bridge has formed, we hypothesized that additional DNA damage might arise downstream of bridge breakage. First, chromosome bridges contain segments of incomplete DNA replication and probably stalled replication forks that could undergo replication fork breakage upon entry into mitosis (51, 52). Second, we found that complex rearrangements were frequent in the second generation (i.e. grand-daughter cells) derived from cells with the stubs of broken chromosome bridges. In all three of the second-generation lineages that we examined by single-cell sequencing, we detected complex rearrangements localized near the bridge breakpoints (Fig. S11). These considerations motivated experiments to determine if the broken stubs of chromosome bridges acquire additional damage upon entry into the next mitosis.

We assayed DNA damage in mitosis using a protocol of live-cell imaging followed by fixation and staining for γ-H2AX in these same cells. Relative to primary nuclei, most broken bridges exhibited little to no detectable damage during interphase, even when cells were held in extended G2-arrest with CDK1 inhibition. However, if cells with broken bridge stubs were released into mitosis, γ-H2AX labeling intensity increased ∼5-fold (Fig 7A-B). This damage was localized to one or a few mitotic chromosomes and was observed only in cells that had a bridge in the prior interphase (Fig 7A). Heavy mitotic γ-H2AX labeling was consistently associated with extensive replication protein A (RPA) accumulation, indicating the generation of single-stranded DNA (ssDNA) (Fig. 7A-B). Surprisingly, pulse-labeling with EdU revealed that RPA and γ-H2AX accumulation coincided with extensive DNA synthesis that occurred specifically on the bridge DNA during mitosis (Fig. 7C). Similar findings were obtained in BJ cells with bridges induced by topoisomerase inhibition (Fig. S12). Live-cell imaging of GFP-RPA2 established that the mitotic replication specifically occurred on the stub of the broken chromosome bridge (Fig. 7D and Movie S14). Therefore, the stubs of broken chromosome bridges undergo a second wave of DNA damage during a burst of aberrant, mitosis-specific DNA replication.

**Figure 7.**
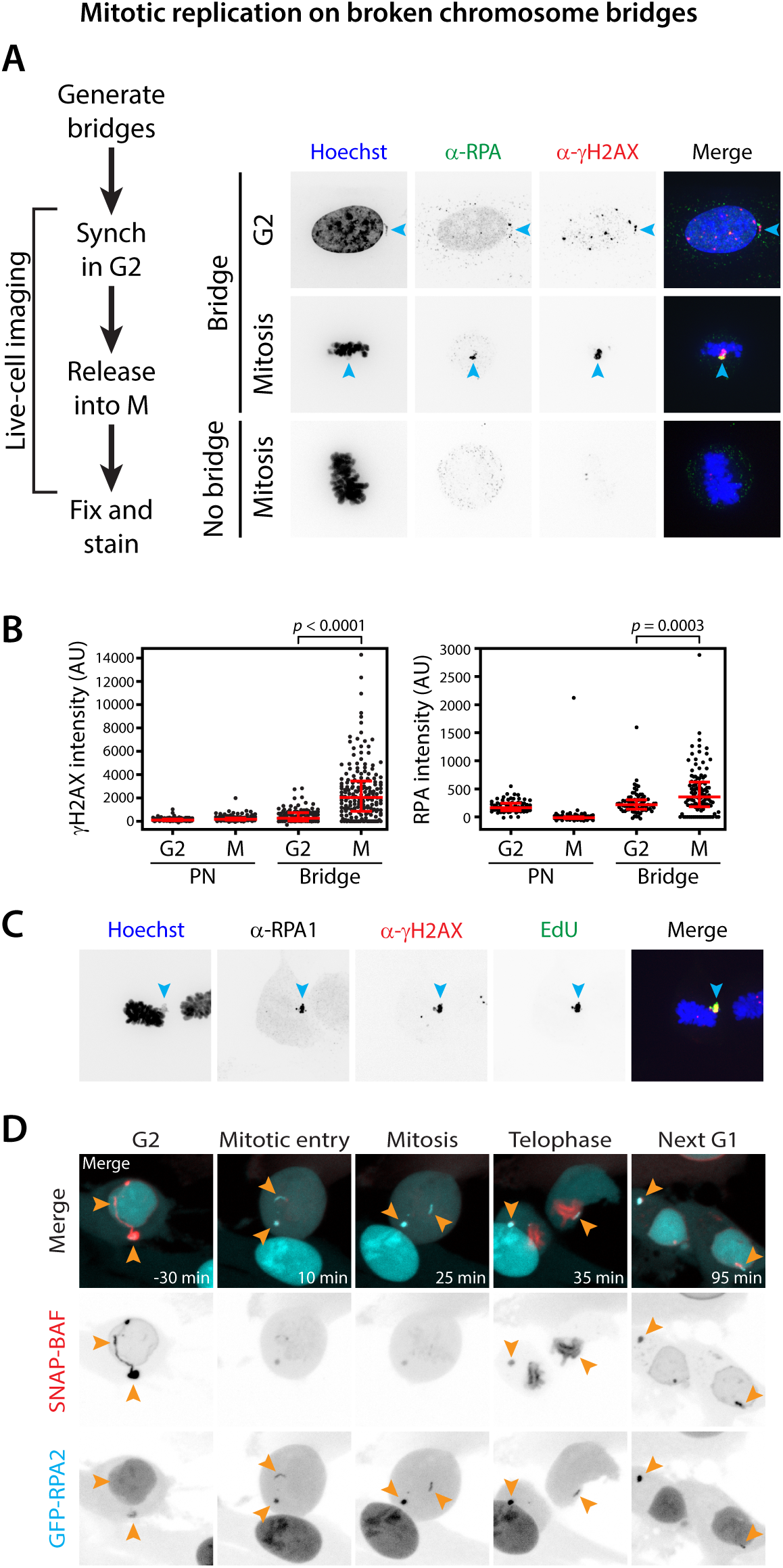
Aberrant DNA replication and extensive DNA damage on bridge DNA after mitotic entry. (A) Correlative live-cell/fixed-cell imaging was used to monitor broken bridge chromosomes entering the next mitosis. Left: schematic of the experiment. Right: example images show cells with broken bridges in G2 arrest and after release into mitosis, as well as a control mitotic cell that did not have a bridge in the prior interphase. Cyan arrowheads: bridge chromosome. (B) Quantification from (A). (C) Images showing correlation of DNA damage (γ-H2AX) with RPA accumulation and active DNA replication (EdU). Cyan arrowheads: bridge chromosome. (D) Representative time-lapse images showing a burst of mitotic DNA replication specifically on a chromosome from a broken bridge. Mitotic replication was visualized by GFP-RPA2; bridge with SNAP-BAF. During mitosis, high activity of vaccinia related kinase (VRK) inactivates DNA binding by BAF (10 to 25 min). Orange arrowheads indicate the broken bridge chromosome. Note that an unrelated interphase cell migrates through the bottom of the field of view in several frames (10 to 35 min). Confocal imaging was performed with a 40× objective, 7 z-slices at 1-µm spacing, acquired every 5 min.

#### Chromosome bridges generate micronuclei

If chromosome bridge formation generated micronuclei, the frequency of chromothripsis and the size of the rearrangement footprint would be further increased (14, 17). This could contribute to the extensive pattern of rearrangements previously reported by bulk sequencing of cell clones derived after telomere crisis (26).

Although it has been previously reported that there is no increase in the frequency of micronuclei immediately after chromosome bridge breakage (26), whether the resulting broken chromosomes segregate normally in subsequent cell divisions has not been examined. To address this question, we used live-cell imaging to track these chromosomes over two generations (Fig. 8A). We confirmed that micronucleation is not an immediate consequence of chromosome bridge breakage in the first cell cycle when the bridge forms and breaks. However, a different result was obtained when we examined daughter cells with broken bridges that went through the next mitosis: 52% of divisions resulted in grand-daughter cells with micronuclei (n = 82 daughter cell divisions examined; Fig. 8A and Fig. S13A). This frequency was higher still when the bridge did not break during the first cell division (65%, n = 20). By comparison, cells without a bridge divided normally and did not produce micronuclei (n = 82 divisions), even though they were present in the same imaging dish and were treated identically. BJ cells induced to form bridges by topoisomerase inhibition also showed an increase in micronucleation rate (>5 fold) in the second cell cycle (Fig. S13A). Therefore, micronucleation is a major downstream consequence of chromosome bridge formation.

**Figure 8.**
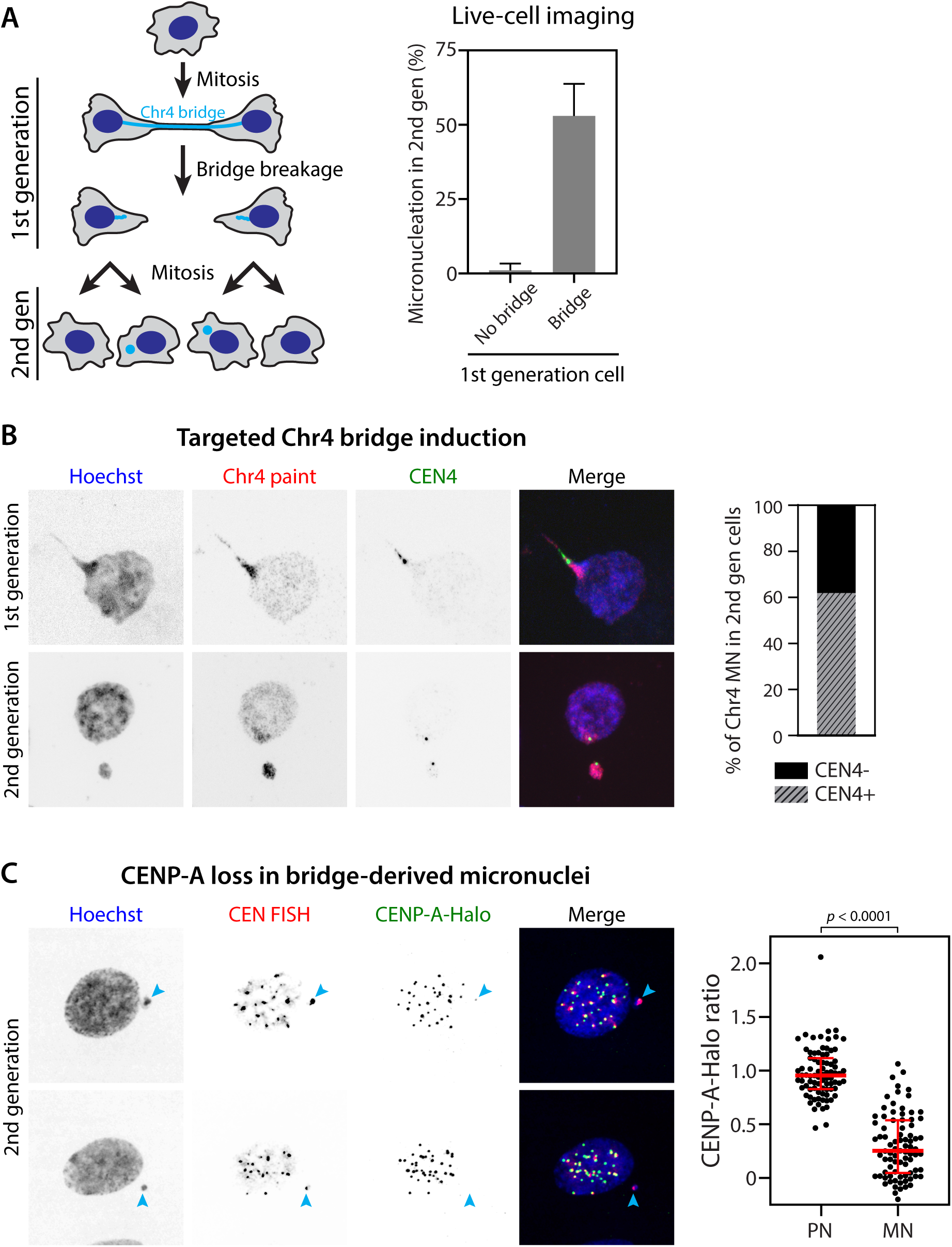
Centromere inactivation on chromosomes within bridges leads to frequent micronucleation. (A) Frequent micronucleation in the second generation after bridge formation. Left: schematic of the live-cell imaging experiment. A cell divides, forming a bridge between two daughter cells in the first generation. After the bridge breaks, the daughter cells with broken bridge stubs divide, generating four “grand-daughter” cells in the second generation. The frequency of micronucleation in second generation cells was measured in control cells that did not have a bridge in the first generation (No bridge) as compared to cells that did (Bridge). Right: quantification of data from the experiment. (B) Micronuclei derived from a bridge chromosome usually retain their centromeres. Left: fluorescence in situ hybridization (FISH) to detect whole-chromosome 4 (Chr4 paint, red) and Chr4 centromere (CEN4, green) in first- and second-generation cells after CRISPR-mediated Chr4 bridge formation. Right: quantification of CEN4 status of Chr4-containing MN in second-generation cells. (C) Centromere defects explain the high rate of micronucleation after chromosome bridge formation. Left: representative images showing CENP-A-Halo labeling (green) and anti-centromere FISH (red) to assess centromere integrity of chromosomes in micronuclei (second-generation cells, as in (A)). Cyan arrowheads indicate the location of the centromere (FISH signal) in micronuclei. Right: quantification of CENP-A-Halo levels at centromeres in micronuclei (MN), relative to those in primary nuclei (PN), *p* < 0.0001 (paired *t*-test). Bridges were induced in the first generation by transient TRF2-DN expression. Only centromere DNA-containing micronuclei were analyzed.

To determine whether the above described micronuclei contain chromosomes from bridges, we generated CRISPR-mediated Chr4 bridges and used fluorescence in situ hybridization (FISH) to detect DNA from Chr4 (Fig. 8B). In the first cell cycle after induction, almost all bridges contained Chr4 sequence (Fig. S1C). Chr4 centromeres (CEN4) sometimes localized to the “base” of the bridge, with some CEN4 spots appearing stretched, suggesting that they were under mechanical tension (Fig. 8B). In the second cell cycle, most micronuclei contained DNA from Chr4 (80%, n = 105), indicating that the majority of chromosomes from bridges mis-segregate in the next mitosis. Surprisingly, most of these micronuclei contained CEN4 DNA (62%, n = 84), indicating that the high rate of mis-segregation cannot be explained by loss of centromeric DNA. As most bridge-derived micronuclei exhibited little or no staining for CENP-A (Fig. S13B), it appeared that the functionality of centromeres trapped within bridges might be compromised.

Defective loading of CENP-A, presumably due to nuclear import defects, could contribute to centromere defects in chromosome bridges, as was recently reported for micronuclei (53). However, failure to load new CENP-A would only cause a 2-fold dilution each cell cycle, which on its own (54) would be unlikely to explain the timing and extent of chromosome mis-segregation we observed. This frequent mis-segregation suggested a more severe defect, perhaps due to active stripping of previously loaded CENP-A from the centromeres of bridge chromosomes. To address this possibility, we pulse-labeled cells expressing Halo-tagged CENP-A from its endogenous locus (55) prior to the induction of chromosome bridges, enabling preferential visualization of the preexisting population of CENP-A that was loaded prior to bridge formation. After labeling and bridge induction (TRF2-DN), cells were given sufficient time to divide twice—first to form bridges, and again to allow bridges to be converted to micronuclei (Fig. 8C). We then measured CENP-A levels at centromeres in micronuclei and determined that they were reduced ∼4-fold relative to the primary nucleus, with ∼25% of micronuclear centromeres lacking any detectable CENP-A (Fig. 8C). By contrast, in control micronuclei induced by nocodazole washout, CENP-A levels were only ∼1.5-fold reduced using the same labeling strategy (Fig. S13C), as expected from defective CENP-A loading alone. Therefore, chromosomes in bridges are prone to CENP-A depletion, which likely compromises centromere identity, leading to frequent micronucleation and additional chromothripsis.

### Common mechanisms for DNA damage in micronuclei and chromosome bridges

We hypothesized that chromosome bridges and micronuclei, although morphologically distinct, might nevertheless have a similarly defective nucleoplasm leading to similar defects in DNA replication—both during interphase and then later in mitosis. This idea was motivated by our previous work showing that the replication defect in micronuclei stems from aberrant nuclear envelope assembly and defective nucleo-cytoplasmic transport (19). Chromosome bridges share this same defect in nuclear envelope assembly (19).

We determined if micronuclei acquire DNA damage during interphase like chromosome bridges. Because nuclear envelope disruption itself causes DNA damage (18), we characterized micronuclei with intact nuclear envelopes. Micronuclei were generated by a nocodazole washout procedure (17) and EdU labeling was used to assess the extent of DNA replication in micronuclei relative to the primary nucleus (Fig. 9A). Intact micronuclei were identified by their accumulation of a nuclear import reporter (RFP fused with a nuclear localization signal, RFP-NLS) (17). As expected, many micronuclei in G2 cells had detectable, but strongly reduced DNA replication relative to the primary nucleus (median EdU ratio = 27%). In these G2 cells, 23% of intact micronuclei displayed DNA damage (Fig. 9A-B). Interestingly, almost all DNA damage occurred in micronuclei with the strongest replication defect, as 90% of damaged micronuclei were at or below the median EdU level (27% of the primary nucleus signal, Fig. 9A-B). Furthermore, nearly all DNA damage in intact micronuclei could be eliminated by blocking the initiation of DNA replication with small molecule inhibitors of either cyclin-dependent kinase (CDK) or Dbf4-dependent kinase (DDK) (Fig. 9A-B). We note that although γ-H2AX intensity measurements were reliable for assessing DNA damage in micronuclei, similar measurements are not feasible for chromosome bridges because of the tension-induced nucleosome loss that occurs on bridge chromosomes (26). Single-cell sequencing showed extensive chromothripsis-like rearrangements in one of ten G2 cells with intact micronuclei (Fig. 9C). Thus, like chromosome bridges, intact micronuclei undergo defective DNA replication in interphase during the first cell cycle after their formation, leading to a low frequency of DNA damage and chromothripsis.

**Figure 9.**
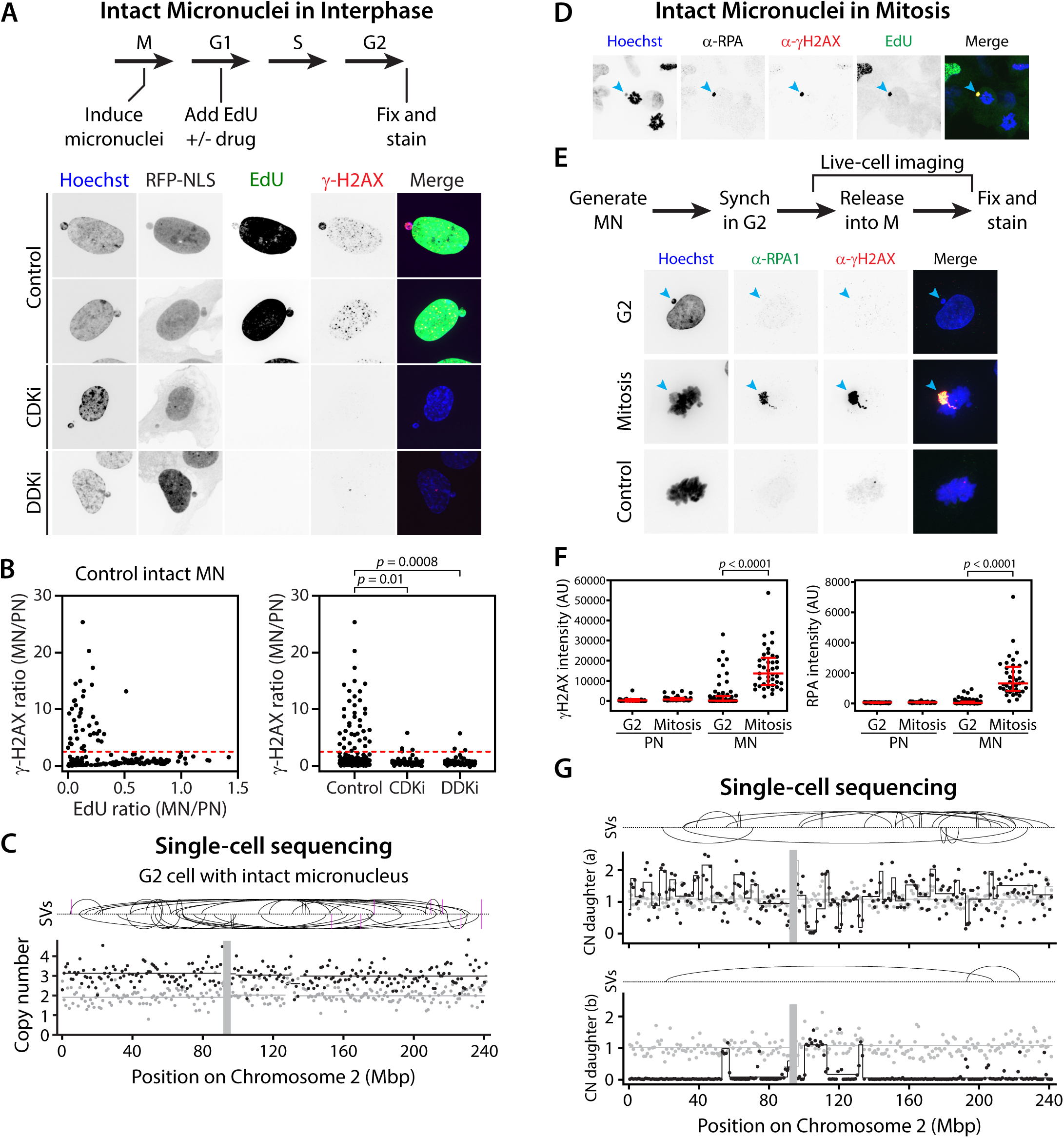
Micronuclei develop extensive DNA damage associated with a burst of mitotic DNA synthesis, which promotes chromothripsis. (A) Modest DNA damage is associated with defective replication in intact micronuclei of G2 cells. Top: schematic of the experiment. Micronuclei were induced by a nocodazole washout procedure (17), and EdU was added in G1 to visualize all DNA replication during the following S phase. Cells were then fixed in G2 (22 hours after mitosis). Where indicated, small-molecule inhibitors of Dbf4-dependent kinase (PHA-767491) or cyclin-dependent kinase (flavopiridol) were also added in G1 to block the initiation of DNA replication. Bottom: example images show intact micronuclei (assessed by RFP-NLS), with counter-staining to relate the extent of DNA replication (EdU) to the amount of DNA damage (γ-H2AX). Robust γ-H2AX signal was correlated with diminished EdU signal and was blocked by the DDK or CDK inhibitors. (B) Quantification of (A) showing DNA damage in micronuclei with poor DNA replication. Left: DNA damage in intact micronuclei (ratio of γ-H2AX intensity in the micronucleus relative to the primary nucleus) relative to replication proficiency (EdU ratio). Dashed red line indicates the threshold (three standard deviations above the mean intensity for primary nuclei) above which micronuclei were scored as positive for DNA damage. Right: compared to control, CDKi (*p* = 0.01) or DDKi (*p* = 0.0008) prevents DNA damage in intact micronuclei; *p*-values determined by Mann-Whitney test. (C) Complex rearrangement of a chromosome from an intact micronucleus in a G2 cell. The chromosome from the micronucleus (Chr2) is under-replicated and was identified by its odd-numbered copy number state. The mis-segregation generating this micronucleus resulted in a diploid cell with an extra copy of Chr2 from the micronucleus (2N+1) (17). Black dots: 1-Mb bins for the haplotype of the micronuclear chromosome, which together with the fully replicated copy of Chr2 haplotype in the primary nucleus, leads to black copy number of ∼3. Gray dots: the other Chr2 haplotype, which is also in the primary nucleus and present at a copy number of 2. n = 1 of 10 cells examined; the remaining 9 G2 micronucleated cells did not exhibit rearrangement of the micronucleated chromosome. (D) Mitotic DNA replication and DNA damage (synchronized fixed cells), similar to Fig. 7C, for cells induced to form micronuclei by nocodazole washout. Cyan arrowheads: micronucleated chromosome. (E) Parallel experiments as in Fig. 7A-B, for cells with intact micronuclei that were released into mitosis. To avoid confounding DNA damage from interphase nuclear envelope rupture, only cells with intact micronuclei (RFP-NLS) were analyzed. Cyan arrowheads: micronucleated chromosome. (F) Quantification from (E). Levels of DNA damage on the micronucleated chromosome increased ∼10-fold in mitotic cells compared to G2 cells (*p* < 0.0001, left plot), concomitant with abrupt initiation of mitotic DNA synthesis as indicated by RPA1 accumulation (*p* < 0.0001, right plot); *p*-values calculated by Mann-Whitney test. (G) Complex rearrangement of a chromosome from an intact micronucleus after passing through mitosis. Copy number and rearrangements are shown for the micronucleated chromosome (one haplotype of Chr2), identified by its odd copy number as described in (C). 8 of 9 daughter pairs examined evidenced complex rearrangements on the micronucleated chromosome.

We next asked if micronuclear chromosomes, like broken chromosome bridges, undergo mitotic replication and secondary DNA damage. Although most intact micronuclei in G2 cells lacked DNA damage, after entering mitosis, there was a ∼10-fold increase in damage levels on micronuclear chromosomes, accompanied by mitotic DNA synthesis and the accumulation of ssDNA (Fig. 9D-F, Fig. S14A, and Movie S15). These findings were corroborated in BJ and HeLa cells (Fig. S14B).

We used single-cell sequencing to determine if transit through mitosis promotes complex rearrangement of micronuclear chromosomes. By live-cell imaging, we identified cells with intact micronuclei that subsequently went through mitosis, generating daughter cells. Unlike the parental G2 cells where only one of ten cells exhibited chromothripsis (Fig. 9C), we observed chromothripsis in eight of the nine daughter pairs after passing through mitosis (Fig. 9G and Fig. S15). Thus, incompletely replicated chromosomes from either micronuclei or bridges undergo aberrant replication upon entry into mitosis linked to a high frequency of complex rearrangement in the next generation.

Therefore, at a low frequency, DNA from chromosome bridges or micronuclei undergo fragmentation and rearrangement during defective DNA replication in interphase. Subsequently, a second wave of abnormal replication and heavy DNA damage occurs when cells enter mitosis. For chromosomes bridges, DNA damage and chromothripsis is further amplified by the induction of micronuclei.

### Complex genome evolution from the formation of a chromosome bridge

Our results identify a cascade of events that should amplify both the frequency and extent of chromosomal aberrations downstream of chromosome bridge formation. These findings predict that the formation of a chromosome bridge should initiate ongoing genome instability where episodes of chromothripsis would necessarily accompany breakage-fusion-bridge cycles (3, 56).

To more directly test our model, we induced the formation of CRISPR-generated Chr4 bridges as discussed above, enabling us to track the evolution of the bridge chromosome in a long-term population growth assay. Bulk genome sequencing analysis of 12 such populations (hereafter “primary clones”) revealed copy number alterations that affected one or both copies of Chr4 in every primary clone (Fig. 10A, Fig. S16). Only one primary clone showed simple breakage of Chr4, whereas the remainder showed complex copy number patterns and rearrangements: seven primary clones had complex copy number alterations confined to one arm of Chr4, and four exhibited alterations of both Chr4 arms (Fig. S16).

**Figure 10.**
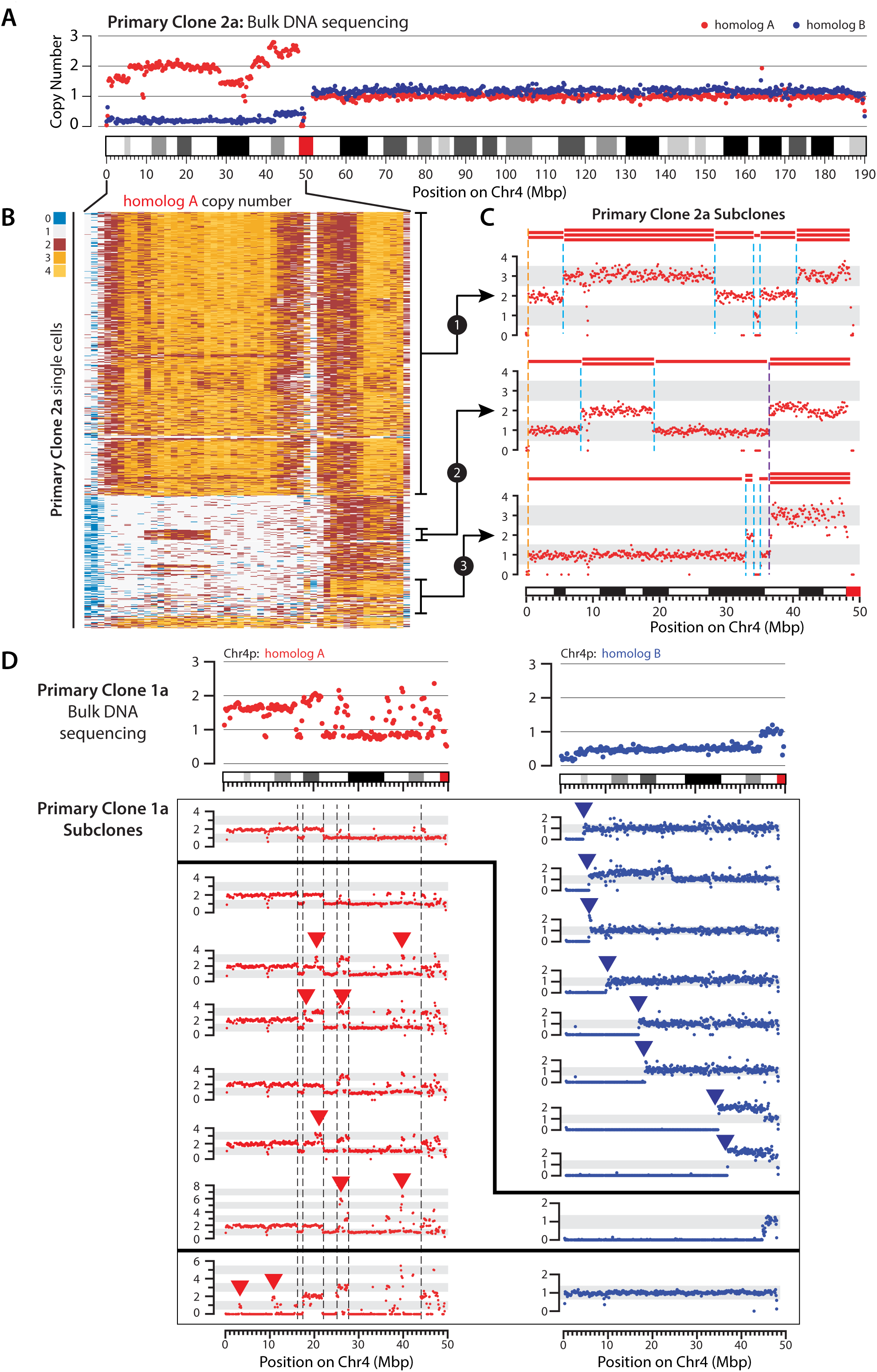
Extensive genetic heterogeneity after chromosome bridge formation. (A) Multiple co-existing subclones in a population derived from a cell with a broken chromosome bridge (Chr4, see Fig. S1). Top: DNA copy number of the two Chr4 homologs (red and blue dots, 25-kb bins) from bulk DNA sequencing of Primary Clone 2a. Non-integer copy number indicates the presence of multiple subclones with different copy number states. (B) Heatmap of DNA copy number for Homolog A on the p-arm of Chr4 (0-50 Mb) in ∼800 single cells, where each row represents one cell. Different subclonal populations can be identified with copy number profiles consistent with those seen in single cell-derived subclones shown in (C). (C) DNA copy number of Chr4p Homolog A (red dots, 25-kb bins) in three subclones grown from single cells isolated from Primary Clone 2a. The copy number state is shown schematically above each plot. The copy number change point shared across all subclones (dashed orange line) is inferred to have resulted from breakage of the Chr4 bridge chromosome in the first generation. Other copy number changes that were shared only among a subset of subclones (dashed purple line), or were private to individual subclones (dashed cyan lines), suggest that these more complex patterns arose during later generations downstream of the initial bridge breakage event. (D) Evidence of ongoing chromosomal instability in Chr4 bridge primary clones indicated by copy number variations across subclones. Top: Bulk DNA copy number of Homolog A (red) and Homolog B (blue) on Chr4p (0-50Mb) in Primary Clone 1a. Bottom: Copy number profiles of each homolog in single-cell derived subclones. The first profile of Homolog A was observed together with each of the first eight profiles of Homolog B, as shown by the thick black lines. Similarly, the next five profiles of Homolog A were each observed together with the ninth profile of Homolog B. The last profiles of Homologs A and B were observed together (two independent subclones). Copy number breakpoints on Homolog A that are shared among all subclones are indicated by vertical dashed black lines. Variations in Homolog A range from focal copy number changes (red arrows), including focal amplifications, to near-complete arm loss (last profile). Profiles of Homolog B show different degrees of terminal loss, a pattern that suggests ongoing instability affecting this chromosome. This is supported by the observation that many of these Homolog B profiles were present at a low clonal fraction (i.e. observed in only one subclone). Therefore, this progressive terminal loss pattern might represent a snapshot of ongoing evolution that eventually results in complete loss of the chromosome arm.

Across all 12 primary clones, there were 26 additional karyotype abnormalities (copy number alterations and/or chromosome fusions) affecting a total of 8 different non-targeted chromosomes (Table S1). Interestingly, these fusions primarily involved acrocentric chromosomes (85% of cases), which were typically seen fused at their p-arms to the aberrant Chr4 (Fig. S17). A similar enrichment of acrocentric fusions has been reported previously in samples after TRF2-DN expression (57), so this effect is unlikely to be an off-target artifact of the CRISPR-generated Chr4 bridge system. These results suggest that broken chromosome ends might be more likely to fuse with acrocentric chromosomes whose terminal rDNA repeat sequences are fragile and prone to spontaneous breakage (58). The high frequency of acrocentric fusions may also be explained by selection, as they provide an efficient path to stabilize broken bridge chromosomes by supplying a single functional centromere and telomere. Importantly, the non-Chr4 aberrations were typically subclonal within each primary clone (Table S1), suggesting their occurrence during downstream evolution of the initial bridge involving Chr4.

Multiple additional lines of evidence indicated a high degree of ongoing genome instability within most of the primary clones, including (i) high frequencies of micronuclei and chromosome bridges (not shown); (ii) non-clonal aberrations of Chr4 observed by cytogenetic analyses (Table S1 and Fig. S17); and (iii) non-integer copy number states in the bulk sequencing data, indicating subclonal copy number heterogeneity (Fig. 10A, Fig. S16, and Fig. S18). Subclonal heterogeneity was directly validated with low-pass sequencing followed by DNA copy number analysis on ∼500–800 single cells from each of nine primary clones (Fig. 10B and Fig. S19). This heterogeneity primarily occurred on Chr4, but was also evident on acrocentric Chrs 13, 14, 15, and 22 at lower penetrance (Fig. 10B and Fig. S19).

To better understand the evolution of copy number variation, we derived subclones from the primary clones, and performed bulk whole-genome sequencing (Fig. 10C-D). These data provided clear evidence for complex copy number alterations and rearrangements occurring downstream of the initial bridge breakage event. Three representative subclone copy number profiles, shown in Fig. 10C, illustrate a shared ancestral breakpoint near the Chr4 p-arm terminus (dotted black line), as well as additional breakpoints specific to each lineage (dotted blue lines). The breakpoints private to each lineage cannot be explained by an early rearrangement event with subsequent loss in some subclones. Instead, these breakpoints can only have been acquired after the ancestral break.

Other examples provided evidence for least three modes of ongoing genome evolution in the primary clone. First, we observed kataegis in 22 of 23 subclones derived from primary clone 1a; however, only a few of these kataegis events were shared across the entire set of subclones (Fig. S20). Most kataegis events were identified only in particular lineages or were even private to just one subclone, which is best explained by some kataegis events occurring late in the evolution of the population (Fig. S20). Second, as shown by the analysis of subclones from one primary clone in Fig. 10D, one homolog (homolog A) exhibited shared breaks as well as private focal changes, including amplifications, which varied in both magnitude and location across the different subclones. Third, in other subclones from this experiment that all share a common homolog A profile, we observed variable loss of the p-arm terminus for homolog B in a pattern suggestive of progressive shortening. This finding, together with our observation that one subclonal lineage exhibits an intact homolog B profile (Fig. 10D bottom), indicates that the evolution of homolog B occurred late during growth of the primary clone and postdated the alterations that gave rise to their shared homolog A profile.

The apparent progressive shortening of homolog B in Fig. 10D likely reflects ongoing BFB cycles. The absence of cells with gain of this region, as predicted by the BFB model, may be explained by gene amplification compromising the fitness of these cells, and/or a bias towards segmental loss due to under-replication of bridge DNA (Fig. 5C-D). This progressive pattern of terminal segment loss generates a sloping average copy number level in the bulk sequencing data (Fig. 10D, homolog B). This pattern is present in most of our primary clones (Fig. S18), and is also commonly observed in cancer genomes (C.Z. Zhang, unpublished). The pattern may therefore represent a useful sequence-based biomarker for ongoing genome instability.

In summary, our results demonstrate that the formation and breakage of a chromosome bridge initiates a cascade of events that rapidly generate a high degree of genomic complexity and cellular heterogeneity. Because later genome evolution obscures the initial genomic alterations, elucidating early events requires a combination of live-cell imaging and single cell genomic analysis (Look-Seq), enabling a direct correspondence to be established between the phenotype of a cell division error and its genomic consequences.

## DISCUSSION

Our results identify a cascade of events that generate increasing amounts of chromothripsis after the formation of a chromosome bridge. These findings substantially revise the chromosome breakage-fusion-bridge model (21, 59, 60) and establish that episodes of chromothripsis will be inherently interwoven with BFB cycles, explaining the implied association between these processes in cancer genomes.

We propose the following model (Fig. S21). Like micronuclei, nuclear envelope assembly around chromosome bridges is aberrant, leading to a depletion of nuclear pores (19) and a defective nucleoplasm. This results in poor DNA replication in the bridge, including stalled replication forks and replication origins that have not fired. The bridge is then broken by a mechanism that requires stretching force from the actin cytoskeleton. Bridge breakage produces simple breaks and local fragmentation, generating free DNA ends that can engage in end-joining and/or in error-prone replicative repair, potentially MMBIR (12, 46). In some cells, this produces the rearrangement signature that we term Tandem Short Template (TST) jumps. These events lead to a low frequency of chromothripsis during the interphase when the bridge forms and breaks. Subsequently, after cells enter mitosis, the stubs of broken chromosome bridges undergo a burst of aberrant mitotic DNA replication, similar to what occurs for micronuclear chromosmes. This leads to significantly more DNA damage and increases the frequency of chromothripsis. Finally, bridge formation disrupts the centromere histone epigenetic mark, compromises centromere function and thereby increases the rate of micronucleation during the next cell division after bridge formation. These micronuclei generate further cycles of chromothripsis, as previously described (15, 17). Combined, these mutational events rapidly generate hallmark features of cancer genome complexity, producing continuing cycles of genome evolution and subclonal heterogeneity from a single cell division error.

### Mutagenesis and DNA fragmentation from actomyosin-based force

It was previously proposed that bridge breakage might occur by mechanical forces generated during chromosome segregation in mitosis (21), cytokinetic furrow ingression, or abscission (24, 25). However, recent findings (26) and our own observations indicate that most bridges remain intact throughout mitosis, cytokinesis, and abscission, arguing against a major role for these processes in bridge breakage. This work suggested that bridges are cleaved enzymatically via a mechanism partially dependent upon the cytoplasmic, endomembrane-associated exonuclease TREX1 (26). Our data disfavor a role for TREX1 and, instead, demonstrate that bridge breakage requires mechanical forces from interphase actomyosin-based contractility (Fig. 1). These forces appear to be exerted locally on DNA near the base of the bridge and are associated with transient actin accumulation and large focal adhesions. This actin accumulation may be triggered by plasma membrane tension, consistent with well-described force-response properties of cytoskeletal contractility (37, 61, 62). Actomyosin forces are transmitted across the nuclear envelope to the bridge chromatin in part by the LINC complex (38, 40).

A simple interpretation of our results is that actomyosin-dependent forces are capable of rupturing the phosphodiester bonds in bridge DNA. The force required to break DNA is estimated to be in the range of 0.5–2 nN (63, 64), which can be achieved or exceeded by traction forces generated from individual focal adhesions, which range from 2∼20 nN (37, 65–67). Although non-covalent interactions connecting actin to chromatin (LINC-dependent and -independent) are individually weak, large numbers of attachments acting in parallel could support the high mechanical load needed to break DNA.

It is also possible that bridge breakage involves DNA processing enzyme(s) whose activity or access to DNA requires mechanical tension. Force-mediated ejection of nucleosomes from DNA (68), which explains the rapid loss of histone signal we and others have observed in bridges (26), might increase nuclease accessibility to bridge DNA. In principle, loss of nuclear envelope integrity could enable access of cytoplasmic nucleases such as TREX1 (26); yet we observed that knockout of TREX1 had no effect on bridge lifetime. Moreover, we did not detect an impact of nuclear envelope rupture on bridge breakage, which disfavors a mechanism based on NE-restricted access of cytoplasmic nucleases to bridge DNA. We therefore propose that mechanical force is either sufficient for DNA breakage or facilitates the action of one or more nuclear-localized factors, such as a nuclease or topoisomerase.

Whole-genome sequencing of single cells after chromosome bridge breakage during the interphase after they were formed revealed that bridge breakage resulted in either simple breaks or local DNA fragmentation, consistent with a breakage mechanism involving mechanical force. Indeed, we also observed both simple breakage and fragmentation when we mechanically broke intact chromosome bridges with a glass capillary. Mechanical bridge breakage could, in principle, cause localized chromosome fragmentation if forces were applied to multiple sites on chromatin, as might occur if the chromatin were in a looped conformation.

We highlight that our ability to draw mechanistic conclusions about bridge breakage benefitted from specific features of our analysis and experimental design: haplotype-specific DNA copy number measurement and the comparison of sister cells (Fig. 2). For example, we noted puzzling examples of internal chromosome segment copy number alterations after bridge breakage (Fig. 2C). Duplication of an internal chromosome segment, as we observed in those daughter cells with copy number gain, would commonly be attributed to a DNA replication error involving template switching or to unequal sister chromatid exchange between repeat sequences (69). However, the comparison between sister cells instead showed this class of copy number alteration in our experiments is best explained by DNA breakage after chromosome-type fusions.

### Chromosomal rearrangements from abnormal nuclear architecture

When bridge breakage was accompanied by fragmentation, we often detected chromosome rearrangements. In most cases, rearrangements resulted from ligation of fragments generated during bridge breakage. This gave rise to a range of outcomes (Fig. 3), from simpler patterns similar to the “local jump” footprint described in cancer genomes (42), to more complex events meeting the criteria for chromothripsis (45). In cases where multiple chromosomes appeared to be present in the bridge, we often also detected interchromosomal rearrangements between breakage sites.

A subset of bridge breakage events (4 of 20) showed a distinct pattern of extreme localized rearrangements, where small (∼2–3 kb) regions contained focal clusters of ∼10 breakpoints each. These “hotspots” were extensively inter-connected by rearrangements, despite being situated megabases apart in the reference genome or, occasionally, on different chromosomes. This generates a signature of multiple short (median ∼150–200 bp) insertions present in tandem within rearrangement junctions (TST jumps; Fig. 5). We think TST jumps are likely generated by aberrant DNA replication involving replication template switching (12) for the following reasons. First, local breakpoint clusters are not expected from a random fragmentation process but could be generated by localized cycles of replication fork collapse, breakage, and aberrant replicative repair. Second, the size distribution of inserted segments (Fig. 6C) is inconsistent with random fragmentation and re-ligation. In agreement with generally similar functional defects in micronuclei and chromosome bridges, we previously identified an example of multiple short tandem insertions in single cell analysis of chromothripsis derived from a micronucleus (17).

The TST jump signature does not result from artifacts during single-cell whole-genome amplification because a similar pattern was observed in bulk sequencing analysis of clonal populations of cells after bridge breakage. Furthermore, we also observed a similar signature by long-read sequencing of a renal cell carcinoma genome. Features of the TST jump signature have been described in large cancer data sets (1, 70) and tandem arrays of short insertions have also been noted in lung cancer genomes, where they may be common ((71) and J. Lee, personal communication). Although the cause of the TST jump signature is unknown, an origin for the insertions from Okazaki fragments might explain the size distribution of the insertions, which is strikingly similar in all of the contexts in which the pattern has been observed.

We demonstrate that similar DNA replication abnormalities occur in bridges and in micronuclei whose nuclear envelopes are intact. This finding demonstrates that nuclear envelope rupture (18) is not absolutely necessary for complex rearrangements on micronuclear chromosomes. Common functional defects of the nucleoplasm in chromosome bridges and micronuclei make sense, given these structures share a common defect in nuclear envelope assembly (19). Generally, DNA replication errors are thought to be major sources of structural variation in cancer genomes. However, what triggers these replication errors in the first place remains poorly understood. We propose that nuclear architecture defects, a hallmark feature of human cancer termed nuclear atypia (72) are a major trigger for cancer-associated DNA replication errors.

### A wave of DNA damage from aberrant mitotic DNA replication

Based on the high frequency of chromothripsis after cells pass through mitosis, we performed a series of experiments that uncovered an unexpected burst of DNA replication that occurs during mitosis, specifically on the stubs of broken chromosome bridges or on micronuclear chromosomes. This mitotic DNA replication is highly aberrant as it produces heavy DNA damage and ssDNA formation. Although some ssDNA forms on bridges in interphase (26), the amount of ssDNA is far greater in mitosis (Fig. 7). Mitotic replication may therefore make a significant contribution to the kataegis pattern that has been linked to chromosome bridge breakage (Figs. S9 and S19; also see (26)).

The mechanism triggering mitotic DNA replication on bridge stubs or micronuclear chromosomes is not known. However, because bridge and micronuclear DNA is incompletely replicated during interphase, these structures likely contain stalled DNA replication forks and licensed replication origins that have not fired. Our prior experiments demonstrate that for micronuclei, incomplete DNA replication occurs because of defective nucleocytoplasmic transport, leading to a failure to accumulate key proteins required for DNA replication and repair (14, 19). However, when cells enter mitosis, the nuclear envelope surrounding the primary nucleus, micronucleus or chromosome bridge will be broken down near-simultaneously. When this occurs, under-replicated bridge or micronuclear DNA will suddenly gain access to the pool of replication factors that had been sequestered in the primary nucleus throughout interphase. Access to replication factors, coupled with high mitotic cyclin-dependent kinase activity (51, 73), likely then activates replication on this incompletely replicated DNA. The DNA damage resulting from mitotic DNA replication may have a number of causes including the well-described activation of structure-specific endonucleases in mitosis (74) and/or the recently discovered cleavage of stalled DNA replication forks that occurs because of removal of the MCM2-7 replicative helicase from mitotic chromosomes (51, 75).

### Loss of the centromeric epigenetic mark in chromosome bridges

In addition to the above described mutational events, we found that chromosome bridge formation predisposes to micronucleation, which could then initiate another round of chromothripsis downstream of bridge breakage (14, 17, 76). We attribute the high rate at which bridge chromosomes mis-segregate in the second mitosis to depletion of CENP-A nucleosomes, which provide the epigenetic specification of centromere identity (77, 78). Two mechanisms likely contribute to CENP-A loss. First, because bridges largely lack nuclear pore complexes (19, 26) and have dimensions that will impede diffusion (79), they should fail in the normal replenishment of CENP-A nucleosomes that occurs each cell division. However, in the timeframe of our experiments, this dilution cannot account for the observed magnitude of CENP-A loss. Instead, our data suggests active CENP-A loss, which we propose may originate from stripping of CENP-A containing nucleosomes by actomyosin forces when centromeric chromatin is trapped within the bridge (Movie S16). The forces required to strip nucleosomes from DNA (∼20 pN) are ∼50-fold lower than those required to break covalent bonds in the backbone (63, 64, 68). Thus, in addition to promoting mutagenesis, actomyosin contractility may disrupt epigenetic marks on chromatin.

### Rapid genome evolution from a single cell division error

The above described cascade of events is predicted to generate ongoing cycles of complex genome evolution. We tested this hypothesis with a CRISPR-based system to track the fate of a defined chromosome (Chr4) bridge. In populations derived from a single cell after bridge breakage, we detected extensive genetic heterogeneity, with evidence that chromothripsis recurs downstream of initial bridge breakage.

Together, these findings identify mechanisms that explain the remarkable potential of a single unrepaired DNA break to compromise the integrity of the genome. In human cells, a single DNA break only weakly, if at all, activates the DNA damage response to block cell cycle progression (80, 81). This means that unrepaired breaks can be amplified into many additional breaks when the cell divides due to the generation of micronuclei or additional chromosome bridges. Because de novo telomere addition is inefficient (82), stable end-capping of chromosomes is primarily achieved through chromosome translocation or break-induced DNA replication (83). For a stable chromosome to result, the DNA segment with the capped ends must additionally contain only one functional centromere. The end result is that downstream of chromosome bridge formation, it is easy for the accumulating burden of DNA breakage to exceed the capacity to stabilize broken DNA ends. Complex genome evolution with subclonal heterogeneity is therefore a virtually inevitable consequence of chromosome bridge formation, a common cell division error during tumor development.

## ACKNOWLEDGEMENTS

We would like to thank M. Bao, P. Campbell, M. Kwon, J. Lee, S. Liu, M. Meyerson, J. Walter, and K. Xie for comments on the manuscript; I. Cheeseman, Maciejowski and T. de Lange for reagents; P. Campbell, J. Maciejowski and T. de Lange for sharing unpublished results. N.T.U. was an HHMI Fellow of the Damon Runyon Cancer Institute and is supported by the Claudia Adams Barr Program for Innovative Cancer Research. T.M. is supported by Cancer Research UK and the Royal College of Surgeons (C63474/A27176). C.-Z.Z. is supported by an NCI career transition award (K22CA216319). C.-Z.Z and L.S. are supported by an NCI Cancer Moonshot award (1R33CA225344.). R.T. and H.F.A. are supported by the Harvard University Milton Fund. A.S. is supported by an NCI Mentored Clinical Scientist Research Career Development Award (K08CA208008) and a Burroughs-Wellcome Career Award for Medical Scientists (CAMS). D.P. is a HHMI investigator and is supported by NIH grant GM083299, a Research Investigator Award from the Lustgarten Foundation, and an award from the G. Harold and Leila Y. Mathers Charitable Foundation.

## AUTHOR CONTRIBUTIONS

D.P., N.T.U., and C.-Z.Z. conceived the project. D.P. and N.T.U. designed the biological experiments, which were performed by A.M.C., L.D.L. and N.T.U. D.P., N.T.U., and C.-Z.Z. designed the sequencing experiments, which were performed by L.S. and N.T.U. C.-Z.Z. designed and performed the analysis of single-cell and bulk sequencing data of RPE-1 samples, with help from L.J.B. The low-pass single-cell CNV analysis was performed by R.T. and H.F.A. T.J.M., and K.J. contributed the data and analysis in Fig. 5D. A.S. contributed the data in Figs. 7C and 7G, and to early experiments on the project. D.P. and N.T.U. wrote the manuscript with edits from other authors.

## SUPPLEMENTAL MATERIALS

Materials and Methods

Table S1.

Figures S1-S20

Movies S1-S16

References 84-95

