## Supplemental Materials for "Mechanisms Generating Cancer Genome Complexity From A Single Cell Division Error"

#### MATERIALS AND METHODS

##### Cell culture and generation of cell lines

All cells were cultured at 37ºC in 5% $CO_2$ atmosphere with 100% humidity. U2OS, Hela, and telomerase-immortalized RPE-1 cells were grown in DMEM/F12 (1:1) media (Gibco) supplemented with 10% FBS, 100 IU/ml penicillin, and 100 µg/ml streptomycin. Telomerase-immortalized BJ cells were grown in DMEM/Medium 199 (4:1) supplemented with 10% FBS, 100 IU/ml penicillin, and 100 µg/ml streptomycin. For cell lines with doxycycline-inducible constructs, tetracycline-free FBS (X&Y Cell Culture) was used in all culture media.

Protein and shRNA expression constructs were cloned into pLenti-CMV-Neo-DEST or pLenti-CMV-Blast-DEST (Addgene) and co-transfected with lentiviral packaging plasmids, pMD2.G and psPAX2, into 293FT cells using Lipofectamine 2000 (Life Technologies) according to the manufacturer's instructions. RPE-1, U2OS, Hela, or BJ cells were virally transduced in the presence of 10 µg/ml polybrene for 16 hours. Transduced cells were obtained either by antibiotic selection (500 µg/ml geneticin or 10 µg/ml blasticidin) started 24 hours after transduction, or by fluorescence activated cell sorting 5-7 days after transduction.

| Expression construct | Reference |
|---|---|
| GFP-BAF | This study |
| SNAP-BAF | This study |
| rtTA3 | Addgene #26429 |
| TRF2-DN (Tet-ON) | (*28*) |
| Cas9 (Tet-ON) | (*84*) |
| Chr4 subtelomere sgRNA | This study |
| TagRFP-T-Utrophin 261 | (*85*) |
| RFP-NLS | (*17*) |
| CENP-A-Halo | (*55*) |
| shRNA p21 / shRNA Rb | (*86*) |

##### Cell cycle synchronization and methods to generate chromosome bridges or micronuclei

To prevent cell cycle arrest (e.g. from telomere dysfunction), all experiments were performed in cells depleted of p53 using 40 nM ON-TARGETplus siRNA SMARTpool L-003329-00-0050 (Dharmacon), or in cells constitutively expressing shRNAs against p21 and Retinoblastoma (*86*). siRNA was transfected with Lipofectamine 3000 (Life Technologies) according to the manufacturer's instructions 24 hours before the start of the experiment.

Chromosome bridges were generated by the following procedures: (1) TRF2-DN induction with 0.1 µg/ml doxycycline for 14-16 hours; (2) Cas9 induction with 1 µg/ml doxycycline for 14-16 hours in cells constitutively expressing sgRNA targeting the Chr4 subtelomere (see details below); (3) topoisomerase II inhibition with 100 nM ICRF-193; (4) partial depletion of condensin by transfection of 1 nM SMC2 ON-TARGETplus siRNA SMARTpool L-006836-01-0005 (Dharmacon) using Lipofectamine 3000 (Life Technologies). Each method was optimized to generate chromosome bridges at a moderate frequency, ~20-30% per cell division (frequency in untreated RPE-1 and BJ hTert cells is <<1%). For TRF2-DN and CRISPR-based methods, bridges begin forming during cell divisions occurring at least ~8 hours after washout of doxycycline, and ~1-2 days after transfection for SMC2 knockdown, whereas ICRF-193 effects are essentially immediate. Hela and U2OS cells generated bridges at a basal frequency of ~5-10% per cell division and did not require experimental induction.

Cell synchronization and induction of micronuclei by nocodazole block and washout was performed as described (*17*); cells were treated with 100 ng/ml nocodazole for 6 hours, followed by mitotic-shakeoff and three washes with warm medium. To detect interphase DNA replication, cells were incubated continuously with 10 µM EdU, added three hours after mitotic shakeoff. EdU incorporation was detected using the Click-iT Plus EdU Alex Fluor 647 Imaging Kit (Life Technologies). To block initiation of DNA replication, 10 µM PHA-767491 or 250 nM flavopiridol was added three hours after shakeoff, together with 50 uM Z-VAD to prevent apoptosis during prolonged arrest in cells depleted of p21 and Rb (*87*). For G2 synchronization, cells were treated with 9 µM RO-3306 for 18 hours. G2-arrested cells were released into mitosis by washing seven times with warm medium. To label mitotic DNA synthesis, cells were released from G2 block into 10 µM EdU. For generation of micronuclei by MPS1 inhibition, G2-arrested cells were released into 0.8 µM NMS-P715.

| Drug | Product details | Concentration |
|---|---|---|
| Doxycycline | Clontech Labs #631311 | 0.1-1 µg/ml |
| ICRF-193 | Sigma #I4659 | 100 nM |
| Latrunculin A | Life Technologies #L12370 | 0.5 µM |
| ML7 | Abcam #ab120848 | 20 µM |
| Nocodazole | Sigma #M1404 | 100 ng/ml |
| NMS-P715 | MilliporeSigma #4759495MG | 0.8 µM |
| 5-ethynyl-2'-deoxyuridine | Life Technologies #C10640 | 10 µM |
| RO-3306 | MilliporeSigma #2176995MG | 9 µM |
| PHA-767491 | Selleck #S2742 | 10 µM |
| Flavopiridol | Selleck #S1230 | 250 nM |
| Z-VAD-FMK | Selleck # S7023 | 50 µM |
| Etoposide | Selleck #S1225 | 10 µM |

**CRISPR-mediated gene knockouts and Chr4 telomere loss**

For RPE-1 cells with doxycycline-inducible Cas9 expression (*84*), sgRNA constructs were cloned into plasmid pLenti-Guide-Puro (Addgene) and delivered by lentiviral transducton. Guide RNA targeting sequences: TREX1, 5′-GAGAGCTTGTCTACCACACG-3′; Chr4 subtelomere, 5′-TTTAGTGCCCGGCCGCAAGG-3′. Transduced cells were obtained by selection with 12 µg/ml puromycin.

To generate CRISPR knockouts by Cas9/sgRNA transfection, TrueCut Cas9 v2 (Invitrogen) was incubated with TrueGuide modified synthetic sgRNA (Thermo Fisher) and transfected using Lipofectamine CRISPRMAX (Invitrogen) according to the manufacturer's instructions. Knockout clones were generated by single-cell flow sorting as above. Guide RNA targeting sequences: TREX1, 5′-GCAGGTACGTACCCAACCAT-3′; SUN1, 5′-CAAGACTCGTCCAATACAGG-3′; SUN2, 5′-ACTGCATGGTGACGCCAACT-3′.

#### DNA isolation and Western blotting

Bulk DNA isolation was performed using PureLink Genomic DNA kit (Invitrogen) according to the manufacturer's instructions. For Western blots, cells were collected by trypsinization, pelleted, washed in PBS, and lysed by addition of an equal volume of 2X SDS lysis buffer (0.1 M Tris pH 6.8, 4% SDS, and 12% β-mercaptoethanol). Samples were denatured at 100ºC for 10 minutes, passed through a 28 ½ gauge needle to shear genomic DNA, run on NUPAGE 4-12% Bis-Tris polyacrylamide gel (Life Technologies), and transferred to PVDF membrane using iBlot (Life Technologies). Odyssey Blocking Buffer (LI-COR) was used for membrane blocking and antibody dilutions. Blots were incubated with primary antibodies for 1 hour at room temperature or overnight at 4ºC. Three washes were performed with PBS-T, followed by incubation with fluorescent secondary antibodies (LI-COR) for 1 hour at room temperature, and three more PBS-T washes. Membranes were visualized using a ChemiDoc MP Imaging System (BioRad).

| Antibody | Manufacturer | Dilution |
|---|---|---|
| Myc | Life Technologies #MA121316 | 1:1000 |
| Cas9 | Cell Signaling Technology # 14697S | 1:1000 |
| GAPDH | Abcam #ab8245 | 1:5000 |
| γ-H2AX | MilliporeSigma #05-636-I | 1:1000 |
| SMC2 | Abcam #ab10412 | 1:5000 |
| α-Tubulin | Sigma #T9026 | 1:5000 |
| TREX1 | Abcam #ab185228 | 1:1000 |
| SUN1 | Abcam #ab124770 | 1:1000 |
| SUN2 | Abcam #ab124916 | 1:1000 |
| IR680 anti-mouse | LI-COR #926-68072 | 1:5000 |
| IR680 anti-rabbit | LI-COR #926-68073 | 1:5000 |
| IR800 anti-mouse | LI-COR #926-32212 | 1:5000 |
| IR800 anti-rabbit | LI-COR #926-32213 | 1:5000 |

#### Live-cell imaging

Cells were plated on #1.5 glass-bottom plates (MatTek), or on ibiTreat 24-well μ-Plates or 35 mm dishes (ibidi). Micropatterns (CYTOO) were custom-made on #1.5 coverslips and mounted in a CYTOOchamber for imaging. Substrate stiffness experiments were performed using CytoSoft Imaging plates (Advanced BioMatrix) coated with 5 μg/ml fibronectin (Sigma). Prior to the start of imaging, SNAP-tagged proteins were labeled using SNAP-Cell 647-SiR dye (New England Biolabs) according to the manufacturer's instructions.

Widefield fluorescence microscopy was performed on a Nikon inverted microscope (Ti-E or Ti2) with Perfect Focus, outfitted with an environmental enclosure to maintain cell culture conditions (37ºC and humidified 5% $CO_2$). Imaging was performed using a 20×/0.75 NA Plan Apochromat Lambda objective (Nikon); Z-stacks of four images at 2-μm spacing were acquired with a Zyla 4.2 sCMOS camera (Andor), at intervals of 5-10 minutes for up to 48 hours.

Actin/myosin contractility was inhibited during live-imaging experiments as follows. Cells were induced to generate chromosome bridges during the first 12-14 hours of imaging in normal growth medium. During a gap between image acquisition intervals, the medium was then replaced to introduce 0.5 µM Latrunculin A (Life Technologies) to disrupt actin dynamics, or 20 µM ML7 (Abcam) to inhibit myosin activation by myosin light chain kinase. Live-imaging continued in drug medium for the remainder of the experiment. Bridges that were intact at the start of drug treatment were analyzed.

Live-cell confocal microscopy was performed on a Nikon Ti-E inverted microscope fitted with a Yokogawa CSU-22 spinning disk head with the Borealis modification, as described above. Z-stacks (seven images at 1-µm spacing) were collected every 10 minutes for 24 hours, using a Prime BSI back-thinned sCMOS camera (Photometrics), and a 40×/0.95 NA Plan Apochromat Lambda objective (Nikon) with the correction collar set to 0.17.

**Indirect immunofluorescence microscopy**

Cells were seeded on #1.5 coverslips and were washed once with PBS prior to fixation. For staining of RPA1, γ-H2AX, and LAP2, and for labeling of EdU incorporation, cells were fixed with 4% paraformaldehyde for 15 minutes. For staining of actin, myosin heavy chain, phospho-myosin light chain 2, paxillin, SUN1, and SUN2, cells were fixed with PTEMF buffer (20 mM PIPES pH 6.8, 1 mM $MgCl_2$, 10 mM EGTA, 0.2% Triton X-100, 4% paraformaldehyde) for 15 minutes.

Fixed samples were washed three times with PBS, permeabilized for 5 minutes at room temperature with PBS+0.5% Triton X-100, and washed three more times with PBS. Samples were then blocked in PBS+3% BSA for one hour, incubated with primary antibodies diluted in blocking buffer, washed three times with PBS+0.05% Triton X-100, incubated with secondary antibodies diluted in blocking buffer, and washed again three times in PBS+0.05% Triton X-100. After DNA staining in PBS+2.5 µg/ml Hoechst 33342 (Life Technologies) for 20 minutes, samples were washed twice in PBS and mounted in ProLong Gold antifade (Life Technologies) on glass slides.

Imaging was performed on a spinning disk confocal microscope (described above). Z-stacks of nine images at 0.5-µm spacing were collected using 60×/1.40 NA or 100×/1.45 NA Plan Apochromat oil immersion objectives.

| Antibody | Manufacturer | Dilution |
|---|---|---|
| γ-H2AX | MilliporeSigma #05-636-I | 1:500 |
| RPA1 | Cell Signaling Technology #2267S | 1:100 |
| LAP2 | BD Biosciences #611000 | 1:750 |
| SUN1 | Abcam #ab124770 | 1:200 |
| SUN2 | Abcam #ab124916 | 1:200 |
| pSer19-Myosin light chain | Cell Signaling Technology #3675S | 1:200 |
| Myosin heavy chain | Biomedical Technologies Inc. #BT-567 | 1:100 |
| Paxillin | Abcam #ab32084 | 1:250 |
| CENP-A | Abcam #ab13939 | 1:500 |

**Same cell correlative live- and fixed-imaging**

Cells were seeded in 35-mm ibiTreat Grid-500 dishes (ibidi) with a gridded imaging surface. For live/fixed imaging of chromosome bridges, asynchronous cells were treated with 100 ng/ml doxycycline for 14-16 hours, followed by washout for 16-20 hours to induce TRF2-DN and generate chromosome bridges. Cells were treated with 9 μM RO-3306, and immediately mounted on the microscope for live-cell imaging throughout the 18-hour arrest in G2. During a gap between image acquisition intervals, samples were released into mitosis by washing seven times. After 45 minutes, cells were fixed with 4% paraformaldehyde and indirect immunofluorescence was performed as described above. Based on live-cell imaging videos and using the gridded dish surface for cell location, cells with intact chromosome bridges at the start of the experiment were analyzed by spinning disk confocal microscopy.

For live/fixed imaging of intact micronuclei, asynchronous cells were treated with 0.8 μM NMS-P715 (MPS1i) for 16-20 hours to generate micronuclei and synchronized in G2 with 9 μM RO-3306 for 18 hours. Live-cell imaging was started in the last 1-2 hours of G2 arrest. During a gap between image acquisition intervals, samples were released into mitosis by washing seven times. After 45 minutes, extraction was performed for 1 minute in CSK buffer (10 mM PIPES pH 6.8, 100 mM NaCl, 1 mM $MgCl_2$, 1 mM EGTA, 0.5% Triton X-100) prior to fixation with 4% paraformaldehyde. Extraction was used before fixation to remove RFP-NLS signal, enabling staining in the near-red fluorescence channel; indirect immunofluorescence was performed as described above. Based on live-cell imaging videos and using the gridded dish surface for cell location, cells with micronuclei that remained intact until mitotic entry (judged by RFP-NLS signal) were analyzed by spinning disk confocal microscopy.

**Fluorescence in situ hybridization**

Cells were seeded on #1.5 glass coverslips and synchronized by serum starvation: samples were exchanged to serum-free DMEM/F12 for 24 hours, followed by release into medium with 10% FBS and 1 μg/ml doxycycline to induce CRISPR-mediated Chr4 bridge formation at the next mitosis. Doxycycline was washed out after 14-16 hours, and samples were collected 24 and 48 hours after washout to analyze 1$^{st}$ generation and 2$^{nd}$ generation cells as follows. Cells were swelled in hypo-osmotic solution (75 mM KCl) for 15-30 minutes before fixation by dropwise addition of a one-half volume of -20ºC Carnoy's solution (3:1 mixture of methanol:acetic acid). After 5 minutes, samples were exchanged to fresh -20ºC Carnoy's solution twice more, incubating at room temperature for 20 minutes each time. Coverslips were removed from the fixative and left to air-dry overnight at room temperature. Chr4 centromere and whole-chromosome 4 "paint" probes (Leica Biosystems) were prepared in hybridization buffer according to the manufacturer's instructions and applied to samples, sealed onto glass slides with rubber cement, and denatured at 75ºC for 2-5 minutes before hybridization at 37ºC in a humidified chamber for 1-2 days. Samples were removed from slides, washed twice in 50% formamide/2X SSC at 42ºC for 10 minutes each, washed twice in 2X SSC at room temperature for 5 minutes each, and mounted on glass slides in ProLong Gold antifade with DAPI (Life Technologies).

To visualize CENP-A in bridge-derived micronuclei, cells expressing endogenously tagged CENP-A-Halo were synchronized by serum starvation and released into doxycycline for TRF2-DN induction of bridges. After 14-16 hours, doxycycline was washed out and cells were labeled with Janelia Fluor 646 HaloTag ligand (Promega) according to the manufacturer's instructions. After 48 hours ($2^{nd}$ generation timepoint), cells were swelled in 75 mM KCl, fixed in Carnoy's solution, hybridized with pan-centromere (α-satellite) FISH probe (Leica Biosystems), and mounted on glass slides using the procedure described above. To visualize CENP-A in micronuclei induced by nocodazole washout, cells were pulse-labeled with Janelia Fluor 646 (Promega) 24 hours prior to the nocodazole block and release procedure described above, and then processed identically for FISH with pan-centromere probe.

**Image analysis**

Live-cell imaging videos were analyzed using NIS-Elements (Nikon) or Metamorph (Molecular Devices). In all experiments, bridge lifetime was measured as the time from cytokinetic furrow ingression to bridge breakage, scored visually based on GFP-BAF. Cell cycle progression was scored visually in cells expressing an mCherry-tagged fragment of Geminin (*88*). Timing of S phase onset was measured as the time interval from completion of mitosis to the first appearance of mCherry-Geminin signal. Total cell cycle duration was measured as the time interval from completion of the first mitosis until nuclear envelope breakdown in the second mitosis. Nuclear envelope (NE) rupture was scored visually in cells expressing RFP-NLS, imaged at 5-minute intervals, and was defined as the time from loss of nuclear accumulation until the first evidence of re-accumulation. For cell motility measurements, cell tracking was performed with the ImageJ/FIJI plugin, TrackMate, applied to detect primary nuclei based on GFP-BAF signal. Cell velocity was measured as the average root-mean-square distance traveled over 10-minute intervals, as previously described (*34*). Only cells that were tracked over at least five imaging intervals were analyzed.

Quantitative analysis of fixed-cell images was performed using ImageJ/FIJI. Image segmentation was performed using maximum intensity projections from Z-stacks. For interphase cells, images were segmented based on Hoechst staining (primary nuclei and micronuclei) or GFP-BAF signal (bridges). For mitotic cells, the main chromosome mass was segmented with Hoechst and the micronuclear/bridge chromosome was segmented based on RPA1 staining. Live-cell imaging showed that nearly all broken chromosome bridges or intact micronuclei become strongly positive for RPA1 upon mitotic entry, validating RPA as a reliable marker for bridge or micronuclear chromosomes in mitosis (Fig. 7E and Fig. S13A). The resulting image masks were used to measure γ-H2AX and/or EdU levels in the primary nucleus and micronucleus/bridge from a sum intensity projection. Centromere FISH signal was used for segmentation to measure CENP-A-Halo intensity at the centromere in the micronucleus relative to the mean intensity from ten well-resolved centromeres in the primary nucleus. In all cases, a random area of the cytoplasm was sampled for background subtraction.

**Single-cell isolation, genome amplification, and DNA sequencing**

Long-term live-imaging and correlative single-cell whole-genome sequencing ("Look-Seq") was performed as described (*17*) or with modifications detailed below. Flow sorting was used to deposit single cells into the wells of a 384-well μClear plate (Greiner). Widefield fluorescence imaging was performed at 15-20 minute intervals, and cells were treated to form bridges in the next mitosis. Following bridge breakage, the two daughter cells were trypsinized and isolated by limiting dilution into a new 384-well μClear plate, where they were allowed to attach for 3 hours before lysis.

During the course of this study, we developed a new method to isolate cells directly from imaging dishes, and this was employed for some Look-Seq experiments. Specifically, cells were plated in a 35-mm gridded ibiTreat dish (ibidi), and imaged at 10-minute intervals. After sufficient time for bridge formation and breakage, the plate was transferred to another Nikon inverted microscope equipped with a CellEctor single-cell isolation system (Molecular Machines & Industries). The sample was exchanged into PBS-based, non-enzymatic dissociation reagent (Sigma) to loosen cell attachment for isolation as follows. Cells of interest were identified based on video recordings, using the gridded dish surface for cell location. Using a glass capillary with an inner diameter of 40 μm, single cells were directly picked from the imaging dish in a volume of 80 nL, and transferred into a ~5-μL droplet of PBS in a PCR tube cap. Cell picking from a single imaging dish was performed within 30 minutes of applying cell dissociation reagent, and cells were lysed within 5-20 minutes after isolation.

In our Look-Seq experiments, we collected cells on average 8.8 hr after bridge breakage. This likely provides ample time for DNA repair by non-homologous end joining (NHEJ), which occurs on a timescale of <2 hours (*89*).

Cell lysis and whole-genome amplification was then performed using the REPLI-g Single Cell kit (Qiagen), and the amplified DNA was purified, sheared to ~500 bp size, and processed with a Library Preparation Kit (KAPA) for multiplexed next-generation sequencing, as previously described (*17*).

**Chromium single-cell CNV library construction**

Hundreds of single cells were individually barcoded for whole-genome sequencing using the Chromium Single Cell DNA kit (10X Genomics). Libraries were successfully generated for nine primary clones (Fig. S18), while libraries for the remaining three primary clones failed to capture sufficient numbers of single cells (<100 unique cells identified).

**Quality assessment of sequencing libraries by low-pass whole-genome sequencing**

Sequencing libraries of single cells and bulk populations were subjected to low-pass whole-genome sequencing (~0.1× mean coverage) on the MiSeq (Illumina) platform to assess library quality based on library complexity and chimeric frequency. Low-pass whole-genome sequencing further enabled us to (i) assess the uniformity of single-cell whole-genome amplification, (ii) estimate haplotype DNA copy number to identify bridge chromosomes or micronucleated chromosomes, and (iii) estimate the number of cells captured in the Chromium single-cell CNV libraries. Sequencing libraries passing quality control were sequenced to a greater depth on the HiSeq 2500 (Illumina) or the NovaSeq (Illumina) platforms. Single-cell samples isolated after bridge resolution were sequenced to ~25× mean coverage (range 20-34×), except for those for which the bridge was broken mechanically (~5× mean coverage). Single-cell samples with intact micronuclei (either before or after cell division/reincorporation) were sequenced to 5-10× mean coverage. Single-cell derived bulk populations were each sequenced to ~10× mean coverage. Single-cell derived subclones were each sequenced to ~1× mean coverage. Each of the Chromium single-cell CNV libraries contained ~ 800 cells and were sequenced in aggregate to ~60× mean coverage, yielding ~0.08×  per cell.

**Alignment and preprocessing of reads from whole-genome sequencing**

All sequencing reads were aligned to the human genome reference (GRCh38 primary assembly) with additional "sponge reference" compiled from repeat-rich sequences (*90*) using **bwa** (http://bio-bwa.sourceforge.net/) v.0.7.12 in the paired-end mode by "bwa mem". For read pairs with multiple alignments, the choice of primary alignment is based on the following criteria in order of priority: proper alignment of both mates (i.e., following the forward-reverse orientation and having a fragment size within the inferred insert size distribution), accurate mapping (highest combined mapping qualities of alignments for one or both mates), most aligned bases (largest combined blocks aligned to the reference); the other alignments are marked as supplementary alignments. Read pairs with identical alignment positions are marked as PCR duplicates by **MarkDuplicates** from the Picard software suite (http://picard.sourceforge.net/). For read pairs with multiple alignments, PCR duplicate status was determined based on primary alignment positions but marked for both primary and supplementary alignments.

**Single-cell variant calling and detection of kataegis**

We performed short variant detection using GATK HaplotypeCaller v4.1.2.0 running jointly on all single-cell bridge samples sequenced to ~ 25× mean depth. In addition to the default read filters, we further excluded reads of low mapping quality (--minimum-mapping-quality 10), excessive clipping (--filter-too-short 50), or having discordant alignment positions (NonChimericOriginalAlignmentReadFilter, MateOnSameContigOrNoMappedMateReadFilter). These filters served to eliminate potential errors due to alignment inaccuracy, including both misalignment of the entire read and incorrect placement of mismatched bases in split or discordant reads. We note that the exclusion of split reads and discordant reads may reduce evidential support for variants near rearrangement breakpoints. The rationale for imposing a more stringent filter was that the majority of split or discordant reads are not related to true structural variants but result from single-cell amplification artifacts. To further exclude variants caused by amplification artifacts that do not generate chimeric DNA, we removed all variants with fewer than four supporting reads.

To identify de novo single-nucleotide variants (SNVs) in each single-cell sample, we excluded variants detected in the bulk RPE-1 data and those with supporting reads in more than one single-cell sample. We further imposed the requirement that the reference base at a de novo variant site be observed in at least 50% of all samples in order to exclude poorly mappable regions of the genome, which are particularly prone to alignment artifacts. Finally, we also excluded variants present in the dbSNP database (build 146) to exclude both common variants not detected in the bulk data and frequent false variants due to recurrent mapping artifacts.

Even with this stringent filtering, the list of de novo variants was dominated by a high frequency of C>T SNVs (~1 per 100 kb). We believe most of these variants reflect cytosine deamination due to heating during library preparation, as previously reported (*91*). However, in a few samples we observed clusters of mutations ("kataegis") where the tight localization of strand-coordinated C>T variants could not be explained by the background rate of deamination. To identify bona fide kataegis events due to AID/APOBEC activity, we searched for strand-coordinated clusters of C>T, C>G, and C>A mutations in the TpC context as described previously (*43*). We further required that kataegis clusters contain at least five strand-coordinated mutations in the TpC context with a minimum inter-mutation distance of 2 kb (*26*). Based on these criteria, kataegis was either only observed on the bridge chromosome near the site of breakage (T-2 and 4-8), or in multiple loci across the genome where they were not associated with detectable copy number alterations (daughter pairs 4-4, 4-5, and 4-6).

**Calculation of total sequence coverage**

*Raw sequence coverage.* We first counted the number of reads in each 10-kb bins using the GATK (v.4.0.12.0-6) CollectReadCounts module with the following read filters to remove non-properly aligned read fragments:

--read-filter FragmentLengthReadFilter --max-fragment-length 1000
--read-filter MateOnSameContigOrNoMappedMateReadFilter
--read-filter MateDifferentStrandReadFilter

*Normalization of GC-dependent bias.* We then normalized read counts in 10-kb bins based on the local average GC content. For bulk libraries, the local average GC content was calculated directly from the reference sequence in each bin. For single-cell libraries, the local average GC content was calculated from the sequence in a 50-kb flanking region of each bin as follows. The average GC content of the $i$th bin was calculated as the arithmetic mean of GC content in bins $i$-2 to $i$+2, excluding any bin with 5,000 or more non-N bases. For bins at chromosome ends, the GC-averaging interval was truncated at the ends. Each 10-kb bin was first grouped into a GC stratum based on the local average GC content. Normalization of sequencing coverage was then performed independently for each (single-cell or bulk) sequencing library. First, the raw read counts were converted to log-read counts and centered by the genome-wide median of log-counts estimated from all bins having ≥ 9,000 non-N bases and average GC percentage in the range of [0.32,0.61] (roughly corresponding to 0.1 to 99.9 percentile of average GC content of all bins). Second, for bins within each GC stratum, we calculated the median of normalized log-read counts, again excluding those with ≥ 9,000 non-N bases or those having zero coverage (these bins reflect incompleteness in the human genome reference or sequence variation between the RPE-1 genome and the human genome reference). For GC strata with ≤100 bins, the estimates of median coverage were inaccurate, and we excluded all the bins in these strata from downstream calculations. Third, we determined the multiplicative GC correction factor for each GC stratum by normalizing the median of log-scale coverage of that stratum against the mean of medians across all strata. This normalization ensured uniform median log-scale coverage in all GC strata. The multiplicative GC correction factors were then applied to each bin by their GC strata to generate GC-normalized log-scale coverage. Finally, we centered the GC-normalized log-scale coverage by the median in all 10-kb bins (excluding those with zero coverage) and then converted the centered, normalized log-scale coverage to linear-scale normalized coverage. For bulk libraries, we further calculated bin-level normalized sequence coverage (30 kb, 90 kb, 250 kb) as the mean of 10-kb normalized sequence coverage in each bin. For single-cell libraries, we performed additional normalization of recurrent amplification bias.

*Estimation and normalization of recurrent bias.* After correcting for GC-dependent bias, we still observed significant variation that was often recurrent among independently amplified libraries. The source of this recurrent bias is unclear but may be related to priming during multiple-displacement amplification, which depends on local sequence composition and chromatin accessibility. To correct for this locus-specific bias, we took advantage of our large number of independently amplified single RPE-1 libraries and determined the median of normalized sequencing coverage in each 10-kb bin across all samples after GC-correction. Variation in the median coverage across bins (assuming no copy-number changes) reflects recurrent amplification bias; we normalized this variation by dividing GC-normalized sequencing coverage in each bin in each sample by the median across all samples, with the exception of de novo or recurrent copy number variants which were identified as follows.

To exclude regions with copy-number variation in the calculation of median coverage, we first calculated arm-level coverage in each sample using the GC-normalized coverage and excluded any arm with average coverage deviating from the median (across all samples) by 25% or more. The median values determined in this way were robust to sporadic segmental or arm-level copy-number changes in one or a few samples. Special treatment was given to regions with altered DNA copy number in a large number of cells. For RPE-1 cells these regions were Chr10q (clonal single-copy gain from 61.8 Mb to q-ter as determined from bulk DNA-Seq) and Chr12 (frequent 2-copy gain of 12p from iso-Chr12p or loss-of-heterozygosity in 12p). For Chr12p, we excluded any sample with an average sequencing depth above 1.45 in the calculation of expected median coverage; this excludes samples with iso-Chr12p. For Chr10q, we first calculated the median coverage for every bin in the gained region and then multiplied the median by the ratio between the average median coverage in 10p relative to the average median coverage in the 10q gain; this ensures a constant median coverage across Chr10.

Putting all of this information together, we determined the expected sequencing coverage for every 10-kb bin in disomic regions reflecting recurrent amplification bias. For each single-cell sample, we divided the GC-normalized coverage by the mean coverage to normalize away recurrent amplification bias. While the correction for GC content resulted in a modest improvement in coverage uniformity, the correction for recurrent bias at the bin level resulted in a far greater improvement. With these two corrections, the normalized sequencing coverage better reflected the relative abundance of DNA sequence in each bin. We further calculated the bin-level normalized sequence coverage (50kb, 250kb, 1Mb) as the mean of 10-kb normalized sequence coverage in each bin.

**Calculation of the haplotype fraction**

*Determination of complete RPE-1 haplotypes.* Whole-chromosome haplotypes of the RPE-1 genome were determined from a combination of linked-reads and Hi-C bulk sequencing of RPE-1 cells using a recently developed computational method (*92*) and validated using our own sequencing data of monosomic cells. Details of variant calling, filtering, haplotype determination and validation can be found in our bioRxiv preprint (*92*).

*Calculation of haplotype copy ratio from allelic coverage.* We first counted the number of reads showing either the reference or the alternate genotype at each heterozygous site (only single-nucleotide variants were included) using an in-house version of the GATK (v.4.0.12.0-6)

ASEReadCounter module that was modified to output counts at every site (including those with zero coverage). We then converted allelic coverage to haplotype coverage using the haplotype phase at each variant site, generating read counts supporting either parental haplotype (**A** or **B**) at all phased variant sites. Finally, we calculated the haplotype fraction at each variant site, defined as the fraction of reads originating from either haplotype. We further calculated the haplotype fraction in 50-kb, 250-kb, and 1-Mb bins as the average haplotype fraction over all variants in each bin. For bulk libraries, the haplotype fraction was calculated for 30-kb, 90-kb, and 250-kb bins.

**Calculation of haplotype DNA copy number**

The haplotype coverage in each bin (50 kb, 250 kb, or 1 Mb for single-cell libraries, 30 kb, 90 kb, and 250 kb for bulk libraries) was calculated as

$$C_{A,B}^{(i)} = D^{(i)} \cdot \overline{R_{A,B}^{(i)}},$$

where $D^{(i)}$ is the normalized total sequence coverage and $\overline{R_{A,B}^{(i)}}$ is the mean haplotype fraction in the *i*th bin. The haplotype copy number was determined by normalizing $C_{A,B}^{(i)}$ by the median of $C_{A,B}^{(i)}$ across the genome (of both haplotypes). In a diploid genome, this median value corresponds to a single copy and haplotype copy number should take integer values (0, 1, 2, …); in tetraploid or close-to-tetraploid genomes, this corresponds to two copies and haplotype copy number can take half-integer values (0, ½, 1, 1½, 2,…). Determining the mean coverage for a single homolog can be challenging if half of the chromosomes have a different copy number (e.g., in a perfect triploid genome, half of all chromosomes have one copy and the other half have two); such cases are rare (not encountered in the current study) but can be easily identified and corrected by manual review.

**Segmentation of haplotype DNA copy number**

Haplotype DNA copy number segmentation was done using a similar iterative strategy as described before (*17*) and performed on the 250-kb bin-level haplotype copy number data. During each round of iteration, the copy number in each bin was first calculated as the local average of 5 consecutive bins (1.25 Mb window) and then rounded to the nearest integer. This procedure was repeated until the copy number of all bins converged to a constant value. The choice of window size (1.25 Mb) determines the minimum length of copy-number alteration (CNA) that is retained after iterative local averaging. For single-copy changes (i.e., one copy loss or gain), the minimum CNA length is 1.25 Mb; for a half-copy change (e.g., in a G2 cell), the minimum CNA would be 2.5 Mb; for 2-copy changes (rare), the minimum CNA length is ~0.6 Mb. We further coalesced segmental breakpoints within 1.5 Mb to eliminate short CNA segments. The segmental haplotype DNA copy number was then calculated as the mean haplotype copy number across all 250-kb bins within each segment. The threshold of minimum CNA length was conservative by choice to accommodate varying amplification variability in a large number of samples. For samples with better amplification uniformity, or for events with better detection power (in particular, the detection of loss or retention), the threshold could be significantly smaller (~ 100 kb).

**Fine-scale segmentation of haplotype retention and loss**

Our haplotype phasing information enabled us to detect smaller (> 100 kb) single-copy deletions in our samples, evident from a near-complete loss of SNPs supporting one of the two haplotypes. For each single-cell sample sequenced to ~30× mean coverage, we divided the genome into 10-kb bins and defined haplotype retention or loss within each bin as the majority coverage status of all SNPs in that bin (0: majority absent, 1: majority covered by at least one read). Bins not containing a phased SNP site were excluded from subsequent analysis.

To identify haplotype copy number change-points, we modeled allelic retention or loss using a Hidden Markov Model (HMM) with two "hidden" states ("RET" and "DEL"), and two observed states (0 and 1) denoting bin-level haplotype coverage as defined above. Using this framework, the emission probabilities reflected the frequency of two types of errors: erroneous coverage of a haplotype that was lost due to sequencing or phasing errors ($\beta_1 = p(1 \mid DEL)$, and coverage dropout in a region of haplotype retention ($\beta_2 = p(0 \mid RET)$).

We determined the emission and transition probabilities empirically from a sample with average amplification uniformity (T-6, daughter a). This sample exhibited an arm-level loss of Chr19p that allowed for estimation of $\beta_1 = 0.03$. Calculating the rate of haplotype coverage dropout on disomic chromosomes yielded $\beta_2 = 0.15$. We assumed equal transition probability ($\alpha$) between states (RET→DEL and DEL→RET) and set its value such that the detection limit of CNAs would be 100 kb (or 10 bins) by the following argument. The probability that the HMM is in the DEL state and emits ten "1" observations is equal to ${\beta_2}^{10}$. Alternatively, the probability that the HMM switches from DEL to RET, emits ten "1" observations, and then switches back is $\alpha^2(1-\beta_1)^{10}$. Setting these two expressions equal to each other and solving for $\alpha$ yielded $\alpha = 5.5 \times 10^{-8}$. With these choices of emission and transition probabilities, the minimum detectable length of loss flanked by large segmental retention was ~200 kb by a similar mathematical argument. The better detection of segmental retention than segmental loss by our model is a consequence of the rate of sequencing errors resulting in erroneous haplotype retention calls ($\beta_1$) being lower than the rate of bin-level coverage dropouts ($\beta_2$). We used the Viterbi algorithm to solve for the most likely sequence of hidden (RET or DEL) states given the observed data.

For samples sequenced to lower depth (~5×), we modified our approach to account for the higher rate of haplotype coverage dropouts. To accomplish this, we increased the bin size from 10 kb to 20 kb. Additionally, we determined the haplotype coverage status in each bin based on the presence of any read support for the observed haplotype (0: no supporting reads in bin, 1: at least one supporting read). We then calculated $\beta_1 = 0.06$ and $\beta_2 = 0.15$ using the same procedure as described above. We set the transition probability $\alpha = 1.75 \times 10^{-6}$ to set the minimum size of CNAs to 10 bins (> 200 kb genomic distance).

Our choices of parameters ensured that when fine-scale segmentation was performed on the whole chromosome, copy number breakpoint calls were restricted to within a few Mb of the main copy number transition. This indicates that the segments identified here are of biological origin and not due to dropout from uneven coverage or sequencing artifacts. Validating this general approach, we observed a striking concordance between breakpoints detected from this segmentation analysis and, independently, structural variants detected in one or both kindred cells (Fig. 4).

**Detection of chromosomal rearrangements**

Chromosomal rearrangements were detected from discordantly mapped read pairs (including split reads) using a previously described algorithm (*17*). For 25× sequencing data, we required at least 3 discordant reads within the range of the average insert size (300 bp) on both breakpoints to be considered as a rearrangement event. We estimate that this minimal threshold allows detection of 78% of genetic variants on each homolog (median across all samples, 53-87%). To exclude amplification artifacts that can appear as false variants with low allele fractions, we used more stringent thresholds for structural variant detection. For genome-wide structural variant detection, we set a conservative threshold by requiring at least 10 variant-supporting reads (including one split read) for each structural variant; with this stringent threshold, the median detection sensitivity is estimated to be 45% (29-58%). For detection of structural variants on the bridge chromosome, we relaxed the threshold to 5 or more variant-supporting reads to increase the median sensitivity to 66% (43-79%).

For 5-10× sequencing data, we required at least 2 discordant reads within the range of the average insert size (300 bp) on both breakpoints to be considered as a rearrangement event. We removed intra-chromosomal rearrangements with distance between breakpoints <150 kb based on the analysis of false positive rearrangement detection due to MDA chimeras (*17*). The only exception to the exclusion of short-range rearrangements was when such events were supported by additional evidence including copy-number alterations (fine-scale retention and loss) or chained templates (see below).

**Assembly of rearrangement chains (TST jumps)**

We assembled complex chains of short templates in three steps. We first performed read-based assembly around each breakpoint detected by discordant reads using SvABA (https://github.com/walaj/svaba). SvABA can both identify the exact breakpoint location at base-pair level resolution and assemble longer contigs from multiple reads spanning the same breakpoint. Second, we classified regions containing more than 3 breakpoints within 10 kb as rearrangement hotspots and determined all the rearrangement junctions in these regions. Some of these junctions were already assembled by SvABA. Those missed by SvABA were usually located at or near regions of interspersed repeats (Alu or L1) and/or had very few supporting split reads (1-2); these events were manually assembled using both split reads and discordant mates. For most breakpoints in hotspots, we were able to assemble short contigs consisting of more than one rearrangement junction. In the last step, we merged these short contigs into rearrangement chains using three types of long-range linkage from discordant read pairs. The first type (most frequent) is junction-junction linkage provided by discordant pairs with each pairmate is mapped to a different rearrangement junction (i.e., a split read spanning both breakpoints of a rearrangement junction) in different contigs. The second type is junction-segment linkage provided by discordant pairs with one pairmate mapped to a rearrangement junction but the other mapped to a segment in a different contig. The third type (less frequent) is segment-segment linkage provided by discordant pairs with pairmates mapped to segments in different contigs. Across each contig, every rearrangement junction was supported by at least one read providing junction-junction type linkage or junction-segment linkage to a non-adjacent segment; segment-segment linkage was used for additional support.

**Haplotype copy number variation from massively parallel single-cell sequencing**

*Cell barcode extraction and filtration.* We first extracted the cell barcodes from the fastq files using the Cell Ranger DNA software (10X Genomics). After barcode extraction, all the sequencing reads were processed using the same workflow as other data. To identify cell barcodes associated with real DNA sequence from single cells, we generated a histogram of read counts for each cell barcode. The distribution of read counts was bimodal for all libraries and we excluded barcodes with lower read counts.

*Haplotype coverage calculation.* We calculated the haplotype coverage at each phased variant site from sequence reads associated with each eligible cell barcode using the same workflow as described above. As the sequence yield is very low for each cell, most variant sites only show coverage of one haplotype, resulting in haplotype fractions at most sites to be either 0 or 1. We therefore used a different approach to estimate haplotype copy number. First, we calculated the fraction of haplotype coverage as the percentage of phased variants showing coverage for either haplotype in each 1 Mb bin. Second, we estimated haplotype copy number from haplotype coverage $f$ using

$$c = \ln(1-f).$$

The rationale behind this calculation is as follows. Assume that the fraction of coverage of a single chromatid to be $f_0$. As $f_0$ is a constant in each library, it can be estimated from the haplotype coverage across the genome when most chromatids have only one copy. When a homolog is present in only one copy, the expected fraction of non-coverage is $1-f_0$. When a homolog has $n$ independently amplified copies, the expected fraction of this homolog that is not covered by any of the $n$ copies is given by given by $(1-f_0)^n$. Therefore, the DNA copy number can be estimated from

$$n = \ln(1-f)/\ln(1-f_0).$$

In practice, it is convenient to use $\ln(1-f)$ as the signal of haplotype coverage after normalization by its median across the genome for both haplotypes. This normalization will produce haplotype copy number that is centered at 1 copy. We finally performed average-linkage hierarchical clustering (UPGMA) to group single cells based on the Euclidean distance between haplotype copy number estimates of all bins on chromosome 4.

**Sequencing of sample from patient with renal cell carcinoma**

We identified a previously published clear cell renal cell carcinoma (*93*) where chromothripsis was associated with a chromosome fusion event that joined Chrs 3p and 5q. Structural variants were called from the short read sequencing data using the BRASS (breakpoint via assembly) algorithm (*94*). Additional DNA from the same tumor from fresh frozen tissue (DIAMOND study; Evaluation of biomarkers in urological disease - NHS National Research Ethics Service reference 03/018) was extracted and eluted into nuclease-free water. DNA (4.6 mg) was prepared using library construction kit SQK-LSK109 (Oxford Nanopore Technologies) in accordance with the manufacturer's protocol. The library was sequenced using the PromethION device (flow cell version FLO-PRO002 using MinKNOW software version 1.14.2, Oxford Nanopore Technologies) with standard run time. Sequencing reads were then aligned to GRCh37d5 using Minimap2 (*95*), allowing the reconstruction of split reads traversing the known rearrangement hotspots. The average sequencing depth for covered regions was 15×. Data are shown for one long read (~40 kb) spanning a complex structural variant exhibiting the TST jump signature. Other reads across this region (not shown) independently supported the presence of the TST jump signature in this sample.

#### SUPPLEMENTAL FIGURE LEGENDS

**Figure S1. Four methods to induce the formation of chromosome bridges.**

(A) Schematic of experimental methods to generate chromosome bridges. Left: TRF2-DN or Chr4 CRISPR methods produce uncapped chromosome ends, which can be fused by the DNA repair machinery to generate dicentric fusions. Note that Chr4 CRISPR can cut both ends of both homologs of Chr4, so like TRF2-DN, chromosome fusions can involve either or both termini of the chromosome. Right: low-dose topoisomerase II inhibition (ICRF-193) or partial knockdown of condensin subunit SMC2 interferes with chromosome decatenation during mitosis.

(B) Control western blots for experimental methods to used induce chromosome bridges. Top: Western blot of RPE-1 cell lines showing doxycycline-inducible expression of Myc-tagged TRF2-DN (α-Myc), or Cas9 (α-Cas9). Note that there is no detectable DNA damage in either case (blot for γ-H2AX) ~40 hr post-induction. Bottom: Western blot showing that partial knockdown of condensin (α-SMC2, depleted to ~68% of control levels), or low-dose topoisomerase II inhibition (ICRF-193) also did not elicit detectable DNA damage. In both panels, α-GAPDH serves as a loading control, and RPE-1 cells treated with etoposide (eto) are a positive control for γ-H2AX.

(C) Terminal Chr4 CRISPR-mediated DNA breaks generate chromosome bridges containing Chr4. Left: Representative image showing that Cas9 induction produces Chr4 bridges as observed by fluorescence in situ hybridization with chromosome 4 "paint" (red) and chromosome 4-specific centromere (CEN4, green) probes, counterstained with DAPI (blue). Right: quantification of images as shown at left. Bars show the frequency of anaphase bridges that were positive (red shading) or negative (gray) for Chr4 paint. Error bars indicate uncertainty based on counting statistics; n = 200 anaphase figures examined per condition.

(D) Optimized conditions for transient expression of TRF2-DN and live-cell imaging minimize previously reported effects on cell cycle progression. Cells were untreated (-Dox, No bridge), or treated with doxycycline for 14 hours, followed by washout, to induce TRF2-DN expression and chromosome bridge formation (+Dox, Bridge). Live-cell imaging was performed using GFP-BAF to visualize cell nuclei and chromosome bridges, and an mCherry-tagged fragment of Geminin was used to monitor S phase onset (*88*). Cells were imaged by widefield fluorescence microscopy using a 20× objective, with four z-focal plane sections acquired every 10 min. Plot shows time from the first mitosis until S phase initiation, and from the first mitosis until the second mitosis (each dot represents one cell). No significant difference was observed in the timing of S phase initiation, and there was a minor ~1.2-fold delay in total cell cycle duration (*p*-values calculated by Mann-Whitney test).

Note: these cell cycle transit times in our imaging conditions contrast sharply with those reported in a prior study that utilized the same cells (*26*). In this prior study (48 hr TRF2-DN induction; 60× objective confocal imaging with ~10-35 z-focal plane sections acquired every 10 min), only 20% of cells entered S phase within 20 hr after mitosis. Our use of the GFP-BAF reporter rather than GFP-H2B, which is lost from bridges under mechanical force, enabled us to employ lower light exposure conditions.

**Figure S2. Neither TREX1 deletion nor nuclear envelope disruption significantly affects the timing of chromosome bridge breakage.**

(A) Top: Western blot showing TREX1 knockout (α-TREX1, green) in published clones generated by transfection of Cas9/sgRNA ribonucleoprotein complex (*26*). α-tubulin (red) is used as a loading control. Bottom: Bridge lifetime curves for control relative to TREX1 knockout clone 2.2 ($p$ = 0.62) and clone 2.25 ($p$ = 0.098). For all lifetime experiments, widefield imaging was performed with a 20× objective at 10 min intervals, acquiring a z-stack of four images at 2-μm spacing.

(B) As in (A), Western blot showing TREX1 knockout clones generated in this study by Cas9/sgRNA transfection. Bridge lifetime curves for control and two TREX1 knockout clones are shown (control versus 1a, $p$ = 0.59; control versus clone 1b, $p$ = 0.90).

(C) A second method to generate TREX knockout clones in RPE-1 cells expressing inducible Cas9. As in (A), Western blot showing TREX1 knockout clones generated in this study by constitutive sgRNA expression (lentiviral delivery) with transient, doxycycline-inducible expression of Cas9 (*84*). Rather than by telomere crisis (A-B), chromosome bridges were induced with ICRF-193. Bridge lifetime curves for control and two TREX1 knockout clones are shown (control versus 2a, $p$ = 0.38; control versus 2b, $p$ = 0.75).

(D) Nuclear envelope rupture is neither required for nor accelerates chromosome bridge breakage. Left: Comparison of bridge lifetime for cells that experienced nuclear envelope disruption, versus cells where no nuclear envelope disruption was detected prior to bridge breakage ($p$ = 0.67). Middle line denotes median bridge lifetime and whiskers indicate interquartile range. Right: Plot showing no significant correlation between bridge lifetime and cumulative nuclear envelope (NE) rupture duration ($p$ = 0.54). Bridges were induced by transient TRF2-DN expression and visualized by GFP-BAF; nuclear envelope integrity was monitored using RFP-NLS.

**Figure S3. Cell motility and bridge extension correlates with bridge breakage.**

(A) Top: plot showing chromosome bridge lifetime in BJ hTert foreskin fibroblast cells induced to form bridges with low-dose topoisomerase II inhibition (ICRF-193). Bottom: representative time-lapse series showing bridge breakage in BJ cells (cells of interest indicated by red arrowheads). Timestamps show time in hours from completion of the first mitosis where the bridge formed.

(B) As in (A) for Hela cells. In time-lapse images, the cell of interest divided to form a bridge (-1.0 to 3.3 hours); the bridge failed to break in interphase and remained intact as the left daughter cell entered the next mitosis (17.3 to 19.2 hours).

(C) As in (A-B), for U2OS cells.

**Figure S4. Increased NE rupture time does not shorten bridge lifetime.**

Left: Plot showing cumulative NE rupture time, in hours, assessed by RFP-NLS, comparing Lamin B1 knockdown cells and controls. “No” micropattern indicates unrestricted cells migrating freely in 2D culture; “L” or “S” refer to cells plated on long or short micropatterns, respectively. N values are number of cells analyzed per condition. Bars represent the mean ± standard error of the mean. No micropattern ± siLB1, $p < 0.0001$; Long micropattern ± siLB1, $p = 0.22$; Short micropattern ± siLB1, $p < 0.0001$.

Right: Violin plot showing distribution of bridge lifetimes (time from mitosis until bridge breakage or entry into the next mitosis), with or without Lamin B1 knockdown, in the conditions described above. Middle solid line shows median bridge lifetime, and dashed lines show interquartile range. N values are number of cells analyzed per condition. No micropattern ± siLB1, $p = 0.69$; Long micropattern ± siLB1, $p = 0.65$; Short micropattern ± siLB1, $p = 0.22$. Note that the observed decrease in bridge lifetime on long micropatterns relative to 2D culture is due to the effects of fibronectin on increasing cell contractility (see Fig. 1I).

**Figure S5. Like RPE-1 cells, bridge breakage in BJ hTert cells requires actin and myosin II-dependent contractility.**

(A) As in Fig. 2A, robust actin concentration at the base of a chromosome bridge prior to breakage in BJ hTert cells. Time-lapse images showing actin dynamics (RFP-Utr261, red) during chromosome bridge breakage (GFP-BAF, green). A cell (cyan arrowhead) divided to form a bridge between the two daughters (-1.1 to 1.8 hours). Large actin accumulations formed at the “base” of the bridge in each daughter cell (5.0 hours), whose apparent contraction preceded bridge breakage (7.9 hours).

(B) Left: schematic of actin and myosin II inhibition experiments. During live-cell imaging, cells were allowed to divide and form bridges prior to exchange into drug medium. Bridges that were intact at the time of drug addition were analyzed. Right: plot of bridge lifetime for BJ hTert cells treated with DMSO (control), Latrunculin A, or ML7, after bridge formation and extension, as in Fig. 1H. Bridges were induced by low-dose ICRF-193. Control data are from Fig. S3A.

**Figure S6. LINC complex components SUN1 and SUN2 localize to chromosome bridges.**

(A) Top: Immunofluorescence images showing localization of SUN1 (green) to the base of chromosome bridges near cytoplasmic actin filaments (phalloidin, red). Bottom: Immunofluorescence images showing similar localization pattern for SUN2. For both SUN1 and SUN2, the staining patterns were validated to be on-target because the signal was absent from knockout clones.

(B) Western blots showing CRISPR knockout of SUN1 (left, red) and/or SUN2 (right, red) in RPE-1 cells; α-tubulin serves as a loading control (green). Dotted white lines indicate digital merging of non-adjacent lanes from the same blot.

**Figure S7. Chromosome bridges generate reciprocal copy number alterations between daughter cells (all examples from this study).**

Genome-wide copy number and structural variant analysis for all single cells (daughter cell dpairs and grand-daughter lineages) that were sequenced after the breakage of a chromosome bridge. For each cell, panel (A) shows CIRCOS plots for both daughter cells. Haplotype copy number was estimated independently for both haplotypes (red and blue dots). Intra- and inter-chromosomal rearrangements are represented as green and purple curves, respectively. Panel (B) shows haplotype copy number (red and blue dots) for each chromosome. Chromosomes were inferred to be in the micronucleus based on an odd-number copy number state relative to the non-micronucleated haplotype. Above each copy number plot, intra- and inter-chromosomal rearrangements are represented as black and magenta curves, respectively.

**Figure S8. Mechanical bridge breakage generates local fragmentation of bridge chromosomes (additional examples).**

(A) Chromosome bridges broken mechanically with a glass capillary (left) display localized reciprocal copy number changes distributed between the bridged cells. Binned copy number (CN) analysis and inferred segmental CN are shown for each chromosome with reciprocal copy number alterations, as described (Fig. 4A). Note that Daughter pairs 1, 2 and 3 are also shown in Fig. 4A.

(B) TREX1 knockout cells undergo bridge breakage with localized fragmentation as seen in control cells. Bridges were induced with transient TRF2-DN expression. Whole-chromosome CN/SV plots (as in Fig. 4B) are shown for each chromosome in the TREX1 knockout bridge pair samples with reciprocal copy number alterations. Note that Chr17 from ΔTREX1 daughter pair 3, and Chr21 from ΔTREX1 daughter pair 4 are also shown in Fig. 4B.

**Figure S9. Kataegis in single cells after chromosome bridge breakage.**

Clusters of strand-coordinated mutations in the TpC dinucleotide context (“kataegis”) detected by single-cell sequencing. “Rainfall” plots representing inter-mutation distance (i.m.d.) between adjacent TpC *de novo* single nucleotide variants (SNVs) are shown for T-2 daughter (a) and 4-8 daughter (b). Each SNV is shown by a point colored to indicate the class of mutation (C>A, C>G, C>T), assuming the alteration occurred on the C of the C:G base pair. In order to show the relative strand coordination of the kataegis clusters, points are filled according to whether the mutated C is present on the top (“Watson”) or bottom (“Crick”) strand of the reference genome. Clusters at least 4 SNVs with a minimum i.m.d. of 2 kb (dotted line) are highlighted with red vertical bars.

**Figure S10. Defective DNA replication within chromosome bridges with controls for detection sensitivity.**

(A) Additional representative example showing poor DNA replication of bridge chromatin. Top: scheme of the experiment. EdU pulse-labeling was performed to detect S phase replication in cells induced to form bridges (TRF2-DN). Bottom: representative images as in Fig. 5C.

(B) Control experiment demonstrating that EdU can be detected within chromosome bridges despite their small amount of DNA. Top: scheme of the control experiment. EdU pulse-labeling was performed in S phase prior to bridge induction (TRF2-DN), so that all chromosomes were labeled within a normal primary nucleus. Pre-labeled bridges then formed in the next mitosis and were imaged in the following interphase. Bottom: images as in Fig. 5C.

**Figure S11. Frequent chromothripsis after cells with broken bridges pass through mitosis.**
Apparent high frequency of complex rearrangement affecting bridge chromosomes after passing through the next mitosis. Top left: Schematic of the two-generation Look-Seq experiment. A cell divides to form a bridge between daughter cells in the first generation. After the bridge breaks, the daughter cells with broken bridge stubs divide, generating four "grand-daughters" in the second generation that were isolated for whole-genome sequencing. Top right and bottom panels: DNA copy number plots (gray dots: 1-Mb bins) with chromosome rearrangements for sets of four grand-daughter cells. In each case, yellow shading indicates the region of the bridge chromosome with chromothripsis.

**Figure S12. DNA damage coincident with a burst of DNA replication on broken bridge chromosomes upon entry into mitosis.**
Mitotic replication on chromosomes from bridges in BJ hTert cells. Top: scheme of the experiment. Cells were treated with ICRF-193 to generate bridges, followed by G2 synchronization with RO-3306. Cells were then released into mitosis in the presence of EdU, fixed, and then stained to detect EdU, γ-H2AX, and RPA1. Bottom: Representative images showing mitotic replication (EdU and RPA1) and associated massive DNA damage (γ-H2AX) on the bridge chromosome.

**Figure S13. CENP-A depletion in bridges leads to a high rate of micronucleation in the cell division after the formation of a chromosome bridge.**

(A) A high rate of micronucleation is common in the second generation after chromosome bridge formation. Scheme of the live-cell imaging experiment is depicted in Fig. 8A. Shown are bar graph plots of the results from additional experiments in RPE-1 and BJ hTert cells. In RPE-1 cells, bridges were generated by transient TRF2-DN expression; control cells lacking chromosome bridges (n = 85) and cells with broken bridges (n = 87) were present in the same imaging dish and treated identically. In BJ hTert cells, bridges were generated with ICRF-193; as above, both control cells lacking chromosome bridges (n = 74) and cells with broken bridges (n = 62) were analyzed from the same imaging experiments. Error bars indicate uncertainty based on counting statistics.

(B) Representative images showing an apparent severe reduction in CENP-A levels in bridge-derived micronuclei.

(C) Nuclear transport defects of micronuclei account for only ~2-fold reduction of CENP-A on micronuclear chromosomes. Left: Fluorescence in situ hybridization to detect centromeres (CEN FISH, red) was used as a fiducial to measure CENP-A levels (CENP-A-Halo, green) in primary nuclei and in micronuclei generated by a nocodazole block and release protocol (*17*). Right: Quantification of fluorescence intensity ratio for a centromere in the primary nucleus (PN*) or in the micronucleus (MN) relative to ten randomly selected centromeres in the primary nucleus (PN); n = 64 cells analyzed, $p < 0.0001$, paired $t$-test. Middle red line: median intensity ratio; whiskers: interquartile range. Note that the ~ 1.5-fold reduction in CENP-A in this experiment, due to defective import into micronuclei (*19, 53*), is a significantly smaller reduction in CENP-A levels than is observed when micronuclei are derived from broken chromosome bridges (Fig. 8C, $p < 0.0001$, Mann-Whitney test).

**Figure S14. DNA damage coincident with a burst of DNA replication on micronuclear chromosomes upon entry into mitosis.**

(A) Representative time-lapse images showing a burst of mitotic DNA replication specifically on a chromosome from an intact micronucleus. Micronuclei were induced by nocodazole washout. Mitotic replication was visualized by GFP-RPA2; chromosomes with H2B-RFP. Confocal imaging was performed with a 100× objective, 7 z-slices at 1-μm spacing, acquired every 2 min.

(B) Mitotic replication on chromosomes from micronuclei in BJ hTert and HeLa cells. Top: scheme of the experiment. Micronuclei were induced by MPS1 inhibition, followed by G2 synchronization with RO-3306. Cells were then released into mitosis in the presence of EdU, fixed, and then stained to detect EdU, γ-H2AX, and RPA1. Bottom: Representative images showing mitotic replication (EdU and RPA1) and associated massive DNA damage (γ-H2AX) on the micronuclear chromosome.

**Figure S15. Chromosomes from intact micronuclei undergo chromothripsis after passing through the next mitosis (all examples from this study).**

Genome-wide copy number and structural variant analysis for all daughter cell pairs generated by division of mother cells containing intact micronuclei. For each cell, panel (A) shows CIRCOS plots for both daughter cells. Haplotype copy number was estimated independently for both haplotypes (red and blue dots). Intra- and inter-chromosomal rearrangements are represented as green and purple curves, respectively. Panel (B) shows haplotype copy number (red and blue dots) for each chromosome. Chromosomes were inferred to be in the micronucleus based on an odd-number copy number state relative to the non-micronucleated haplotype. Above each copy number plot, intra- and inter-chromosomal rearrangements are represented as black and magenta curves, respectively.

**Figure S16. Daughter cells derived from the breakage of a CRISPR-generated Chr4 bridge produced primary clones with extensive Chr4 copy number alterations (bulk sequencing of all 12 primary clones).**

After bridge breakage, daughter cells were separated, grown into primary clones, and then analyzed by whole-genome sequencing. Plots showing copy number data for the two homologous copies of each chromosome (red or blue circles for the different haplotypes). In most cases, both daughter cells that were isolated after breakage grew into primary clones (PC); pairing information for these populations is indicated by cladograms at left. For the PC 5a and PC 7a samples, the corresponding "b" sister cell failed to grow into a clone. Gain of 10q blue haplotype is clonal in RPE-1 cells, and iso(12p) red haplotype is a subclonal event in some populations (based on copy number and karyotype data).

**Figure S17. Subclonal karyotype aberrations in primary clones derived from single cells after the breakage of CRISPR-generated Chr4 bridges.**

Left: Karyotypes were determined by Giemsa staining (G-banding) of metaphase spreads from Primary Clone 1a. Solid arrows indicate near-clonal aberrations of Chr4, such as dicentric fusion of chromosomes 4 and 14, and inverted duplications of 4p. Hollow arrows and arrowheads indicate subclonal aberrations: isochromosome 13q (upper panel), and dicentric fusion of chromosomes 13 and 14 (lower panel).

Right: Karyotypes for Primary Clone 6b. Subclonal aberrations (hollow arrows) of Chr4, including dicentric fusion with Chr22, as well as possible inverted duplications of Chr4q are indicated. Additional subclonal aberrations were observed on Chr22: fusion of unidentified material to the p-arm, or large deletions of material from the p- and q-arms.

Note that translocations involving Chr 4 involved fusions with acrocentric chromosomes more frequently than expected by random chance ($p$ = 0.024, Fisher's exact test). Other translocations, such as the 13:14 fusion in Primary Clone 1a may be derived from an original Chr4:14 fusion, although we cannot exclude that some of these events not involving Chr 4 could be due to CRISPR off-target DNA cleavage. It was previously reported that TRF2-DN induces a high frequency of acrocentric fusions in RPE-1 cells (*57*), supporting the idea that fusion to acrocentric chromosomes is a general mechanism for the resolution of DNA breaks caused by chromosome bridge breakage, irrespective of how the bridge is formed.

**Figure S18. Gradual copy number transitions indicate ongoing genome instability within populations of cells.**

(A) Cartoon showing that cell-to-cell heterogeneity in the size of segmental copy number alterations (left) results in a gradual, sloping copy number transition when the population is analyzed by bulk DNA sequencing (right).

(B) Four examples of gradual sloping copy number transitions. The profiles are from bulk DNA sequencing data of primary clones derived from single cells after the breakage of CRISPR-mediated Chr4 bridges (see Fig. S16). Shown is the relevant Chr 4 haplotype.

**Figure S19. Copy number heterogeneity among single cells within primary clones derived from cells after the breakage of CRISPR-generated Chr4 bridges.**

Paired plots showing genome-wide (top) and Chr4 (bottom) haplotype copy number profiles obtained from low-pass single-cell sequencing. Normalized log-adjusted haplotype coverage (haplotype copy number) was rounded to the nearest integer and plotted as shaded rectangles in 1-Mb intervals. Each row of the grid represents a single cell from the pooled sequencing library. Rows are ordered based on unsupervised clustering of haplotype CN. Each set of single cells derived from a primary clone is plotted on its own page.

**Figure S20. Kataegis in single-cell subclones derived from Primary Clone 1a.**

Multiple subclonal kataegis clusters in Primary Clone 1a. Top: as in Fig. S9, rainfall plot of TpC SNVs on Chr4p with at least two supporting reads in any subclone. Clusters of at >2 SNVs are highlighted. Bottom: dot plot representing the number of TpC SNVs detected in a given subclone from each kataegis cluster. Left: Dendrogram representing the phylogeny of subclones inferred from shared copy number breakpoints. Groups of subclones with similar copy number patterns were readily apparent (solid lines) whereas the hierarchical relationship between more divergent subclones could not be unambiguously resolved from copy number alone (dashed lines).

**Figure S21. Model for the generation of cycles of chromothripsis from either chromosome bridges or micronuclei.**

(A) Chromothripsis from chromosome bridges. **First mitosis**: Chromosome bridges result from the segregation of dicentric or catenated chromosomes. Nuclear envelope (NE) assembly on bridge DNA is abnormal, such that only one subset of NE proteins ("core" proteins) assemble on the bridge, while the remaining NE proteins, including nuclear pore complexes ("non-core") do not assemble (not shown; (*19*)). **First interphase**: Bridges extend during interphase because of cell migration. The dearth of nuclear pore complexes, combined with the fact that the long, thin geometry of the bridge will act as a barrier to diffusion (*79*), likely results in a local nucleoplasm deficient for DNA replication and repair proteins (*14, 17, 19*). Bridge DNA is therefore poorly replicated. Furthermore, during the extension of the bridge, local contractile forces may stretch chromatin and eject histones (*26, 68*), including the centromeric histone, CENP-A. Next, the bridge breaks in a manner that requires actomyosin mechanical forces. This event generates simple DNA breakage or local DNA fragmentation, which can lead to "local jumps" and chromothripsis-like rearrangements by DNA end-joining. In addition, these broken DNA ends may undergo error prone replicative repair (e.g., by MMBIR), generating the TST jump signature and chromothripsis. Overall, chromothripsis occurs at a relatively low frequency in the first interphase. **Second mitosis**. Upon entry into the second mitosis, under-replicated DNA from the broken chromosome bridge stub undergoes DNA replication coupled to heavy DNA damage, which increases the frequency of chromothripsis. Presumably due to the loss of CENP-A during the prior interphase, chromosomes from broken bridges frequently mis-segregate during the second mitosis. **Second interphase**. Mis-segregation of the broken chromosome generates micronuclei, which then amplifies the frequency and extent of chromothripsis (*17*). DNA damage acquired during the initial bridge breakage event, during mitotic replication, and/or within subsequently forming micronuclei, can all lead to more chromosome bridges (BFB cycles) and more micronuclei during subsequent cell divisions. Altogether, these events generate ongoing complex genome evolution from a single cell division error—the formation of a chromosome bridge.

(B) Parallels between chromothripsis from micronuclei and from chromosome bridges. **First mitosis.** Lagging chromosomes result from mitotic errors or because of the inability to segregate acentric chromosome fragments. **First interphase.** Nuclear envelope (NE) assembly on the lagging chromosome is abnormal, as described above for chromosome bridges (not shown; (*19*)). Accordingly, there is poor DNA replication and an initial low frequency of chromothripsis. Due to defects in the import of proteins required to maintain nuclear envelope stability, the micronuclear envelope can also undergo spontaneous rupture (not shown; (*18*)), leading to further DNA damage by an unknown mechanism. Note the nuclear envelope surrounding chromosome bridges may also undergo rupture, but the geometry of the bridge precludes cytological assessment by monitoring for loss of nuclear integrity. **Second mitosis**. Like chromosome bridges, under-replicated chromosomes from micronuclei undergo a burst of aberrant DNA replication, leading to additional DNA damage and further promoting chromothripsis.

#### SUPPLEMENTAL MOVIES

Movie S1. Bridge breakage can occur in the absence of detectable nuclear envelope rupture. The bridge was visualized with GFP-BAF (green) nuclear envelope integrity was monitored with the import reporter, RFP-NLS (red). The time interval between frames is five minutes; total video duration is 10.8 hours. Widefield imaging was performed with a 20x objective. Related to Fig. S2D.

Movie S2. An example of a bridge breakage event in BJ hTert foreskin fibroblasts (GFP-BAF is grayscale). Timestamp shows relative time in hours. Widefield imaging was performed with a 20x objective. Related to Fig. S3A.

Movie S3. An example of U2OS cells with a bridge (GFP-BAF is grayscale) that remains intact throughout interphase. The time interval between frames is 10 minutes; total video duration is 31 hours. Widefield imaging was performed with a 20x objective. Related to Fig. S3B.

Movie S4. An example of Hela cells with a bridge (GFP-BAF is grey scale) that remains intact throughout interphase. The time interval between frames is 10 minutes; total video duration is 23.7 hours. Widefield imaging was performed with a 20x objective. Related to Fig. S3C.

Movie S5. A pair of RPE-1 daughter cells with a chromosome bridge (GFP-BAF is grey scale) on a long fibronectin micropattern. Broken bridge ends are indicated by blue arrowheads. Timestamp shows relative time in hours. Widefield imaging was performed with a 20x objective. Related to Fig. 1B, left.

Movie S6. A pair of RPE-1 daughter cells with a chromosome bridge (GFP-BAF is grey scale) on a short fibronectin micropattern. Timestamp shows relative time in hours. Widefield imaging was performed with a 20x objective. Related to Fig. 1B, right.

Movie S7. Non-uniform stretching of a chromosome bridge and subsequent breakage in the taut region (RPE-1 cells; GFP-BAF is grey scale). The time interval between frames is 10 minutes; total video duration is 3.7 hours. Imaging was performed on a spinning disk confocal microscope with a 40x objective. Related to Fig. 1D.

Movie S8. Actin dynamics associated with chromosome bridge breakage in RPE-1 cells. Left panel: Merge overlay of GFP-BAF (green) with the actin reporter, RFP-Utr261 (red). Middle panel: GFP-BAF (monochrome). Right panel: RFP-Utr261 (monochrome). Contraction of the actin-rich structure (red arrowheads, 10 to 35 min) immediately precedes bridge breakage (35 min), and then the actin structure rapidly disassembles (40 to 55 min). Cyan arrowheads indicate broken bridge ends. Timestamp shows relative time in minutes. Widefield imaging was performed with a 20x objective. Related to Fig. 1E.

Movie S9. Actin dynamics associated with chromosome bridge breakage in BJ hTert cells. Left panel: Merge overlay of GFP-BAF (green) with the actin reporter, RFP-Utr261 (red). Middle panel: GFP-BAF (monochrome). Right panel: RFP-Utr261 (monochrome). Timestamp shows relative time in hours. Widefield imaging was performed with a 20x objective. Related to Fig. S5A.

Movie S10. Representative video showing that disruption of the actin cytoskeleton (Latrunculin A) blocks chromosome bridge breakage (GFP-BAF is grayscale). Red arrowheads mark the daughter cells of interest. At the start of the video, cells connected by a long bridge are in normal growth medium ("No drug") and are later exchanged into drug medium ("Latrunculin A"). Near the end of the video, the upper cell passes through the next mitosis as indicated (but fails cytokinesis due to actin inhibition). The time interval between frames is 10 minutes; total video duration is 11 hours. Widefield imaging was performed with a 20x objective. Related to Fig. 1F.

Movie S11. Example video from a Look-Seq experiment (C-4 daughter cell pair, condensin knockdown; GFP-BAF is grayscale). Timestamp shows relative time in hours. Widefield imaging was performed with a 20x objective. Related to Fig. 2A-B.

Movie S12. Example video from a Look-Seq experiment (T-1 daughter cell pair, TRF2-DN; GFP-BAF is grayscale). Timestamp shows relative time in hours. Widefield imaging was performed with a 20x objective. Related to Fig. 2A-B.

Movie S13. Example video from a Look-Seq experiment (4-2 daughter cell pair, Chr 4 CRISPR; GFP-BAF is grayscale). Timestamp shows relative time in hours. Widefield imaging was performed with a 20x objective. Related to Fig. 2A-B.

Movie S14. Sudden onset of DNA replication on a broken bridge stub after entry into mitosis. Left panel: Merge overlay, GFP-RPA2 (green) is a marker for mitotic replication; SNAP-BAF (red) labels the broken bridge stub. Middle panel: GFP-RPA2 (monochrome). Right panel: SNAP-BAF (monochrome). Imaging was performed on a spinning disk confocal microscope with a 40x objective. Timestamp shows relative time in minutes. Related to Fig. 7C.

Movie S15. Sudden onset of DNA replication (GFP-RPA2, green) on a micronucleated chromosome (H2B-RFP, red) after entry into mitosis. Imaging was performed on a spinning disk confocal microscope with a 100x objective. Time interval between frames is two minutes; total video duration is 92 minutes. Related to Fig. S14A.

Movie S16. Example of apparent CENP-A depletion when the centromere is trapped inside of a chromosome bridge. Video shows endogenous Halo-tagged CENP-A as a cell divides (0 to 35 min) to form a bridge between the two daughters. Left panel: merge overlay of GFP-BAF (green) and CENP-A-Halo (magenta). Middle panel: GFP-BAF (monochrome). Right panel: CENP-A-Halo (monochrome). Fluorescence intensity for the CENP-A-Halo focus in the bridge (magenta arrowhead), but not a nearby focus present in the main nucleus (cyan arrowhead) decreases sharply upon stretching of the bridge (from ~100 min), and does not recover over time. Timestamp shows relative time in minutes. Imaging was performed on a spinning disk confocal microscope with a 40x objective.

**Table S1. Cytogenetic analysis of primary clones derived from single cells after bridge breakage.**

| Sample | Karyotype | Metaphases counted | Karyotyped | Aberrations of Chromosome 4 | Aberrations involving all other chromosomes |
|---|---|---|---|---|---|
| Primary Clone 1a | 45~46,X,add(X)(q28),add(4)(p16),?psu dic(4)(4;14)(p11;p11.2),i(13)(q10)[cp5] | 10 | 5 | Homolog 1: add(4)(p16) in 5/5 karyotypes, with subclonal variation in the amount of additional material fused to 4p<br>Homolog 2: ?psu dic(4)(4;14)(p11;p11.2) in 3/5 karyotypes; non-clonal aberrations in remaining 2/5 karyotypes | ?psu dic(4)(4;14)(p11;p11.2) in 3/5 karyotypes<br>i(13)(q10) in 3/5 karyotypes<br>?psu dic(4)(4;13) together with der(14) in 1/5 karyotypes |
| Primary Clone 1b | 79~85,XX,add(X)(q28)x2,add(1)(q23.1)x2,-4,-4,?psu dic(4)(4;14)(p11;p11.2)x2,i(4)(q10),-14[cp4] | 10 | 4 | Homolog 1: i(4)(q10) in 4/4 karyotypes<br>Homolog 2: ?psu dic(4)(4;14)(p11;p11.2) in 3/4 karyotypes<br>None of the aberrant Chr4s were seen in duplicate in any karyotype | Loss of one copy of Chr14 in 4/4 karyotypes<br>?psu dic(4)(4;14)(p11;p11.2) in 3/4 karyotypes<br>Two copies of add(1)(q23.1) in 4/4 karyotypes |
| Primary Clone 2a | 44~45,X,add(X)(q28),add(4)(p12),?psu dic(4)(4;14)(p11;p11.2)[cp5] | 10 | 5 | Homolog 1: ?psu dic(4)(4;14)(p11;p11.2) in 5/5 karytoypes<br>Homolog 2: add(4)(p12) in 3/5 karyotypes | ?psu dic(4)(4;14)(p11;p11.2) in 5/5 karytoypes<br>add(22)(p) in 4/5 karyotypes |
| Primary Clone 2b | 44~46,X,add(X)(q28),?psu dic(4)(4;14)(p11;p11.2)[cp8] | 10 | 8 | Homolog 1: ?psu dic(4)(4;14)(p11;p11.2) in 8/8 karyotypes, but with subclonal variation in the amount of additional material fused to 4p<br>Homolog 2: add(4)(p16) in 2/8 karyotypes | ?psu dic(4)(4;14)(p11;p11.2) in 8/8 karyotypes |
| Primary Clone 3a | 43~45,X,add(X)(q28),-4,der(14;22)(q10;q10)[cp5] | 11 | 5 | Homolog 1: Loss of Chr4 in 5/5 karyotypes<br>Homolog 2: Apparently normal, except for unusual banding pattern in pericentromere region | der(14;22)(q10;q10) with aberrant banding near the centromere in 5/5 karyotypes |
| Primary Clone 3b | 43~44,X,add(X)(q28),-4,-10,der(14;22)(q10;q10)[cp4] | 10 | 4 | Homolog 1: Loss of Chr4 in 4/4 karyotypes<br>Homolog 2: Apparently normal | der(14;22)(q10;q10) with aberrant banding on 14q (possibly Chr4 material) in 4/4 karyotypes |
| Primary Clone 4a | 45~46,X,add(X)(q28),-4,?psu dic(4)(4;14)(p11;p11.2),add(12)(q15),+r[cp4] | 10 | 4 | Homolog 1: ?psu dic(4)(4;14)(p11;p11.2) in 3/4 karyotypes<br>Homolog 2: ring chromosome containing 4q in 4/4 karyotypes | ?psu dic(4)(4;14)(p11;p11.2) in 3/4 karyotypes<br>add(12)(q15) in 4/4 karyotypes |
| Primary Clone 4b | 43~45,X,add(X)(q28),der(4)add(4)(p16)add(4)(q35),add(12)(q15),der(14;22)(q10;q10), +19,-21[cp7] | 10 | 7 | Homolog 1: der(4)add(4)(p16)add(4)(q35) in most karyotypes with subclonal variation in the amount of additional material fused to 4p<br>Homolog 2:<br>?psu dic(4)(4;14)(p11;p11.2) in 1/5 karyotypes | der(14;22)(q10;q10) in 3/7 karyotypes<br>add(12)(q15) in 7/7 karyotypes, which in 1 case contains additional material compared to the others |
| Primary Clone 5a | 42~46,X,add(X)(q28),-4,add(4)(p14),-7,-22,+r[cp6] | 10 | 6 | Homolog 1: add(4)(p14) in 6/6 karyotypes<br>Homolog 2: ring chromosome containing 4q in 6/6 karyotypes | Loss of one copy of Chr7<br>Loss of one copy of Chr22<br>Material from Chrs 7 and/or 22 could be present in the ring chromosome |
| Primary Clone 6a | 43~45,X,add(X)(q28),-4,add(4)(p16),-22,add(22)(p11.2),+r[cp4] | 10 | 4 | Homolog 1: add(4)(p16) in 4/4 karyotypes, with subclonal variation in the amount of additional material fused to 4p<br>Homolog 2: ring chromosome containing 4q in 2/4 karyotypes | Loss of one copy of Chr22 in 4/4 karyotypes<br>add(22)(p11.2) in 2/4 karyotypes |
| Primary Clone 6b | 44~46,X,add(X)(q28),add(4)(q35),-22,add(22)(p11.2)[cp7] | 10 | 7 | Homolog 1: add(4)(q35) in most karyotypes | add(22)(p11.2) in 7/7 karyotypes |
| Primary Clone 7a | | | | | |

### Supplemental Figure 1

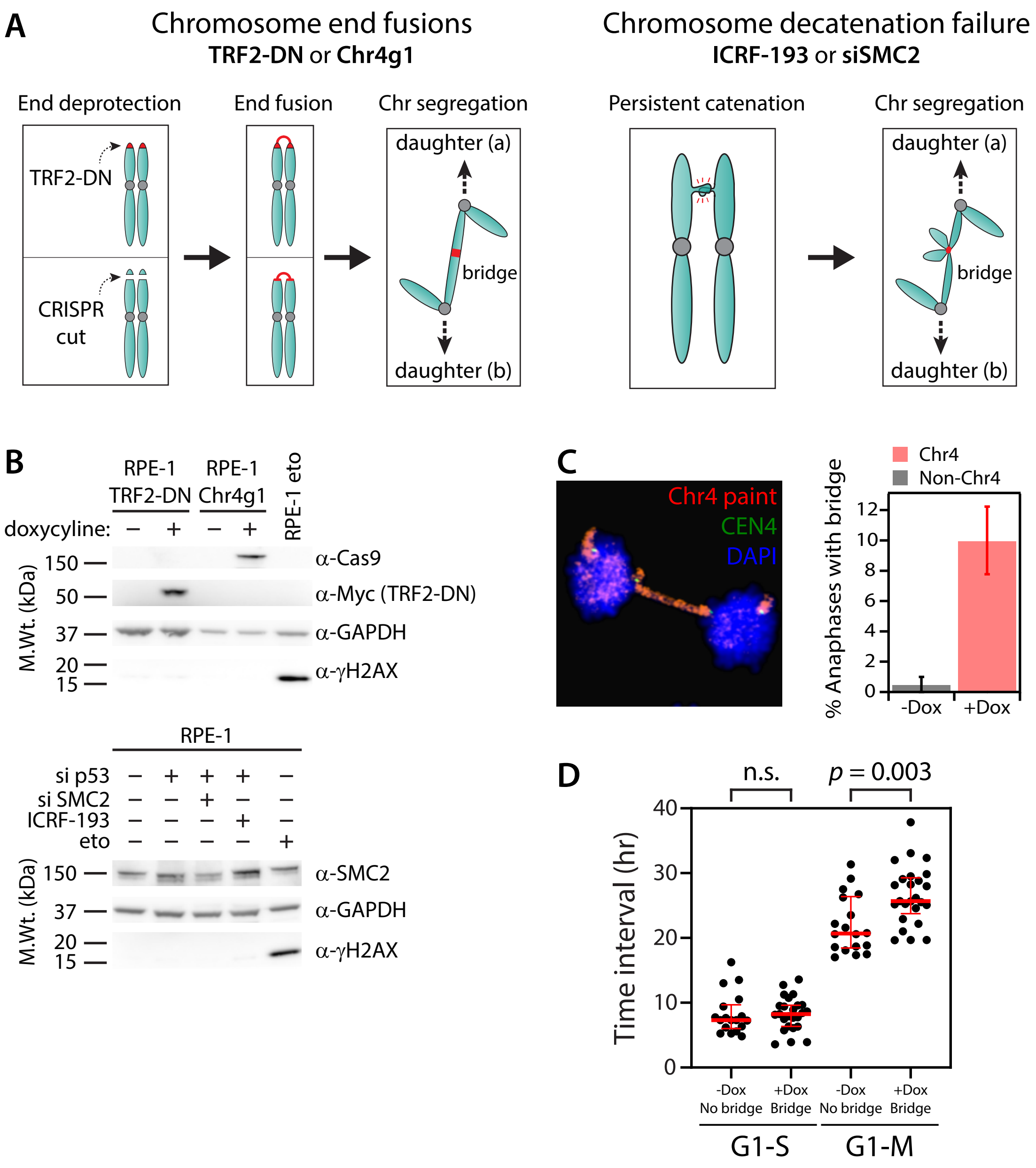

### Supplemental Figure 2

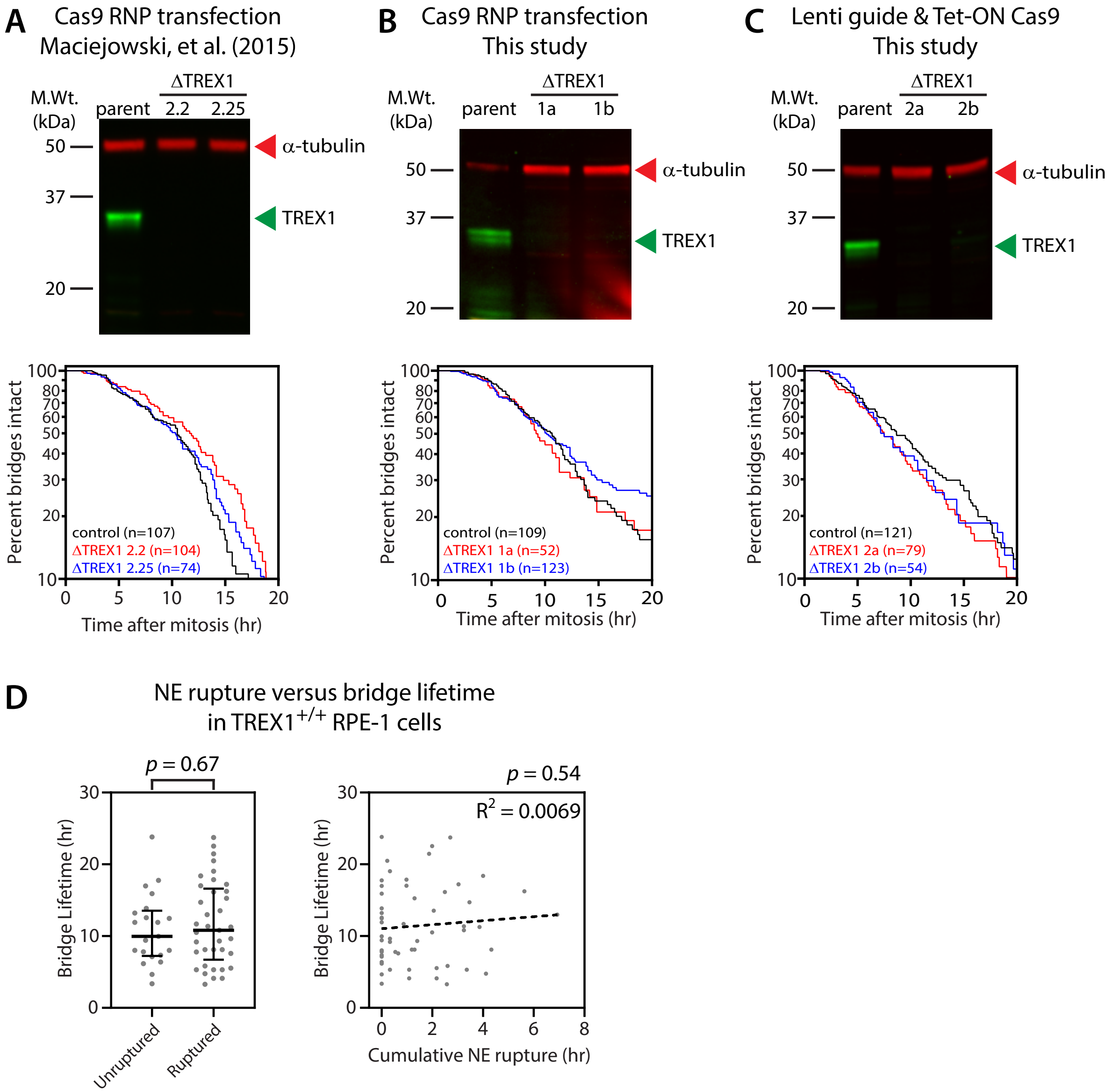

### Supplemental Figure 3

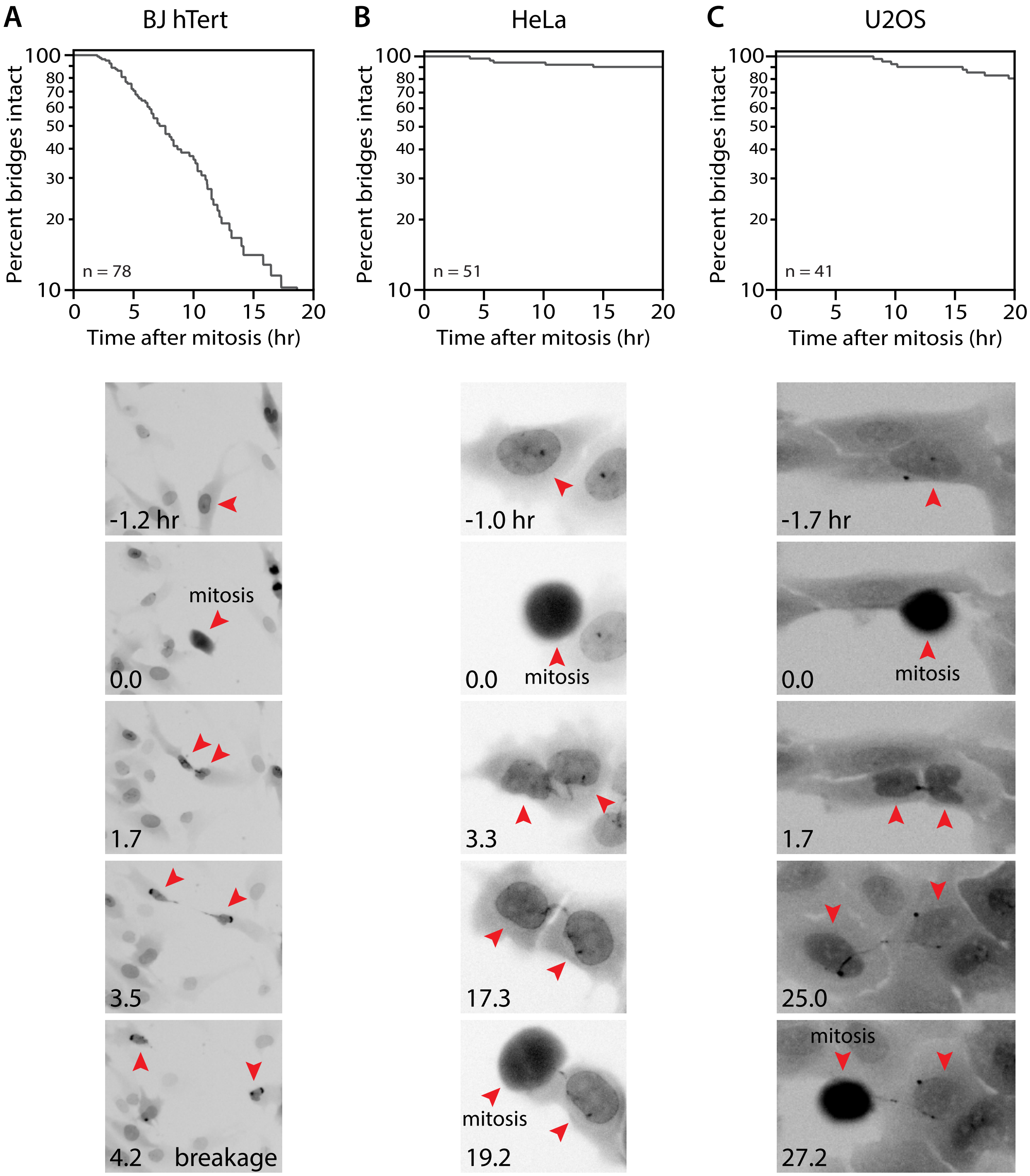

Supplemental Figure 4

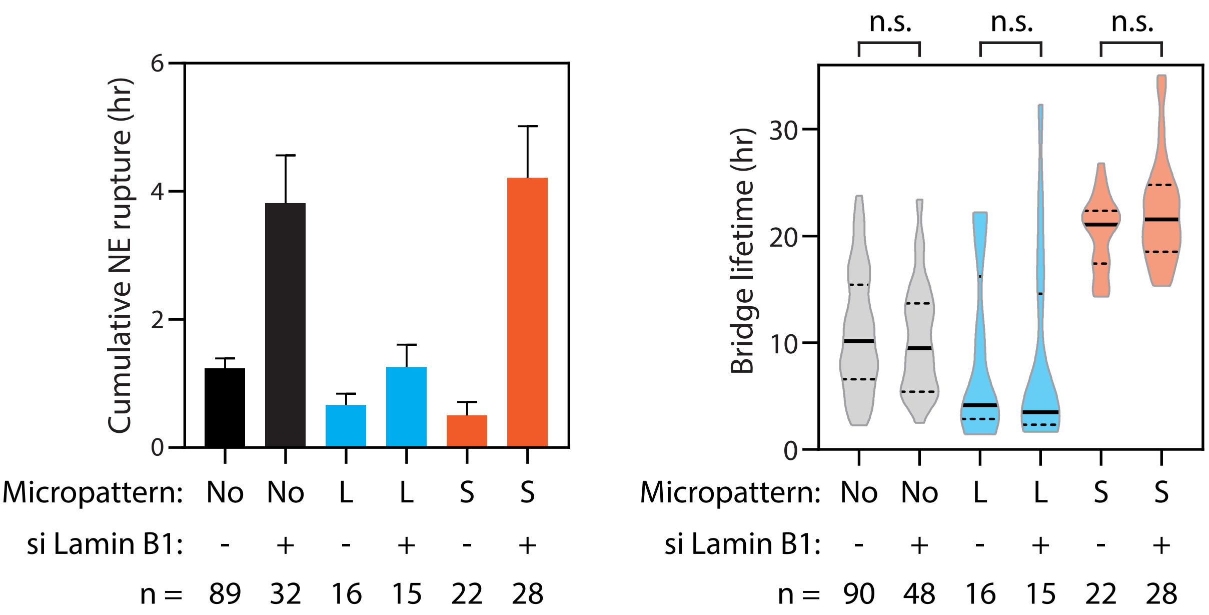

### Supplemental Figure 5

**BJ hTert cells with chromosome bridges induced by ICRF-193**

**A** Actin dynamics associated with chromosome bridge breakage

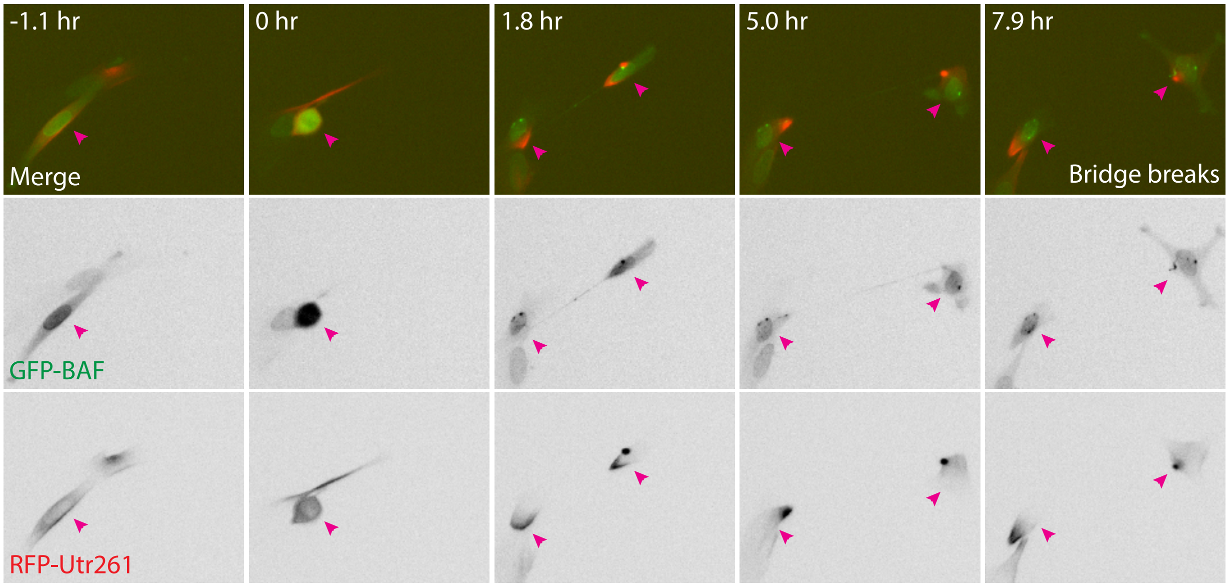


**B** Actomyosin contractility is required for bridge breakage

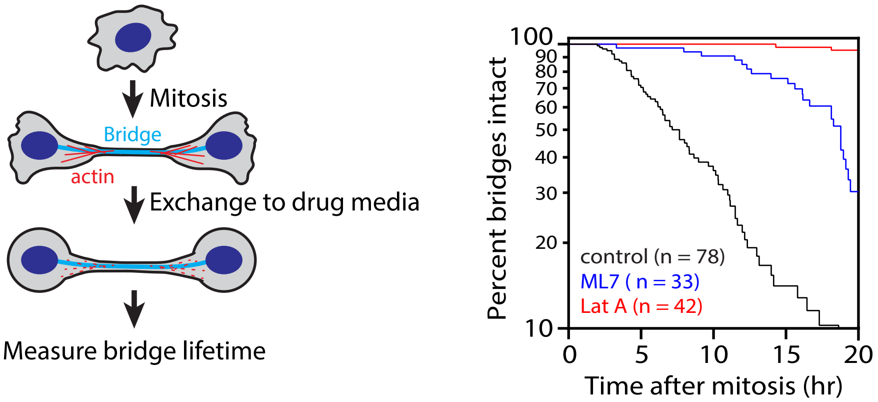

Supplemental Figure 6

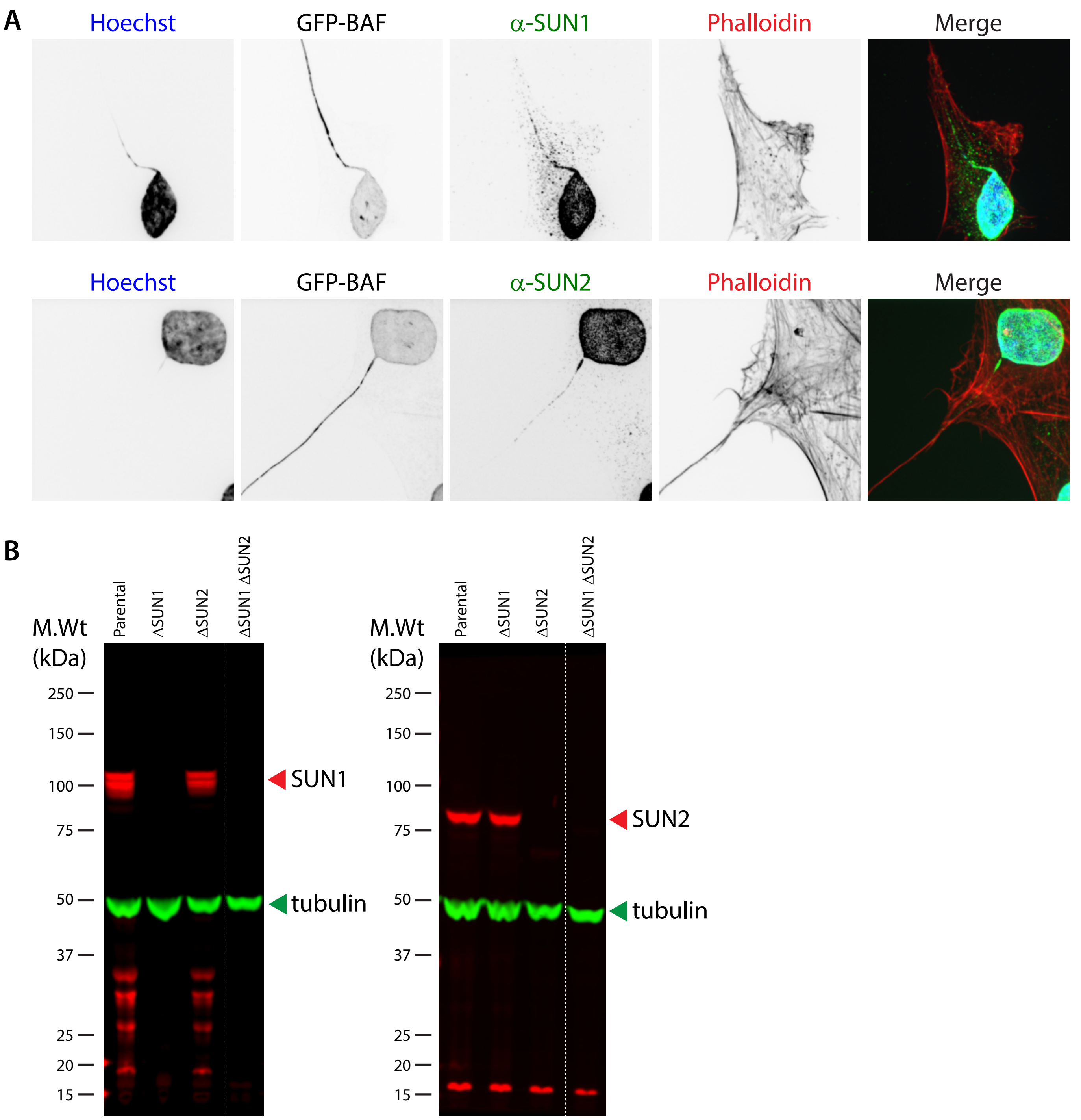

#### Supplemental Figure 7

Chromosome bridges generate reciprocal copy number alterations between daughter cells (all examples from this study).


C-1 Condensin knockdown: Chromosomes 7B, q-arm; 13A,q-arm; XB, q-ter. . . . . . . . . . 1

C-2 Condensin knockdown: **Chromosomes 4B, 5A, 6A, all on q-ter.**; Chr. Xq segment likely in a micronucleus. 4

C-3 Condensin knockdown: Chromosome 1A, p-arm internal. . . . . . . . . . 7

C-4 Condensin knockdown: **Chromosomes 16A and 18B, p-ter.** . . . . . . . . . . 10

T-1 Transient TRF2-DN expression: **Chromosome 3A, q-ter.**. . . . . . . . . . 13

T-2 Transient TRF2-DN expression: Chromosome 2B, q-ter. . . . . . . . . . . 16

T-3 Transient TRF2-DN expression: Chromosomes 7B, p-ter.; Chromosome 12B, p-arm; 19B, p-arm. . . . . . . 19

T-4 Transient TRF2-DN expression: Chromosome 18B, q-arm internal; incomplete loss of Chromosomes 17B and 21A, q-ter. . . . . . . . . . . 22

T-5 Transient TRF2-DN expression: Chromosomes 1B, p-ter.; 6B, q-ter.; 8A, q-ter. . . . . . . . . . . 25

T-6 Transient TRF2-DN expression: Chromosomes 3B, p-ter.; 11A, q-ter.; 12A, p-ter.; 19B, p-arm; incomplete loss of Chromosome 20B on q-ter. . . . . . . . . . . 28

T-7 Transient TRF2-DN expression: Chromosomes 12B, p-arm; 17B, q-ter. . . . . . . . . . . 31

T-8 Transient TRF2-DN expression: Chromosomes 3A, p-ter.; 16B, q-arm. . . . . . . . . . . 34

4-1 CRISPR: Chromosome 4A q-ter. . . . . . . . . . . 37

4-2 CRISPR: Chromosome 4A q-ter. . . . . . . . . . . 40

4-3 CRISPR: Chromosome 4A p-arm internal. . . . . . . . . . . 43

4-4 CRISPR: Chromosome 4A q-ter. . . . . . . . . . . 46

4-5 CRISPR: Chromosome 4A q-ter. . . . . . . . . . . 49

4-6 CRISPR: Chromosomes 4A (p- and q-ter.) and 4B (p-ter). . . . . . . . . . . 52

4-7 CRISPR: Chromosome 4B p-ter. . . . . . . . . . . 55

4-8 CRISPR: **Chromosome 4A and B (p-ter)**. . . . . . . . . . . 58

TREX1-1 Transient TRF2-DN expression + TREX1 knockout: Chromosome 21A q-ter. . . . . . . . . . . 61

TREX1-2 Transient TRF2-DN expression + TREX1 knockout: Chromosome 4B q-ter. . . . . . . . . . . 64

TREX1-3 Transient TRF2-DN expression + TREX1 knockout: Chromosome 7A, p-arm internal; Chromosome 12A, p-arm. . . . . . . . . . . 67

TREX1-4 Transient TRF2-DN expression + TREX1 knockout: Chromosome 16A, q-arm. . . . . . . . . . . 70

TREX1-5 Transient TRF2-DN expression + TREX1 knockout: Chromosome 2A, q-arm; XB, q-ter. . . . . . . . . . 73

TREX1-6 Transient TRF2-DN expression + TREX1 knockout: Chromosome 1B, p-arm internal; 5A, q-ter.; 16A, q-ter. . . . . . . . . . . 76

TREX1-7 Transient TRF2-DN expression + TREX1 knockout: Chromosomes 4A and 4B, p-ter.; 11B, q-ter.; 17B, p-ter. . . . . . . . . . . 79

TREX1-8 Transient TRF2-DN expression + TREX1 knockout: Chromosome 3B, p-ter. with pre-existing terminal loss; 19B, p-arm. . . . . . . . . . . 82

**Bridge Pair C-1** | Condensin knockdown: Chromosomes 7B, q-arm; 13A,q-arm; XB, q-ter.

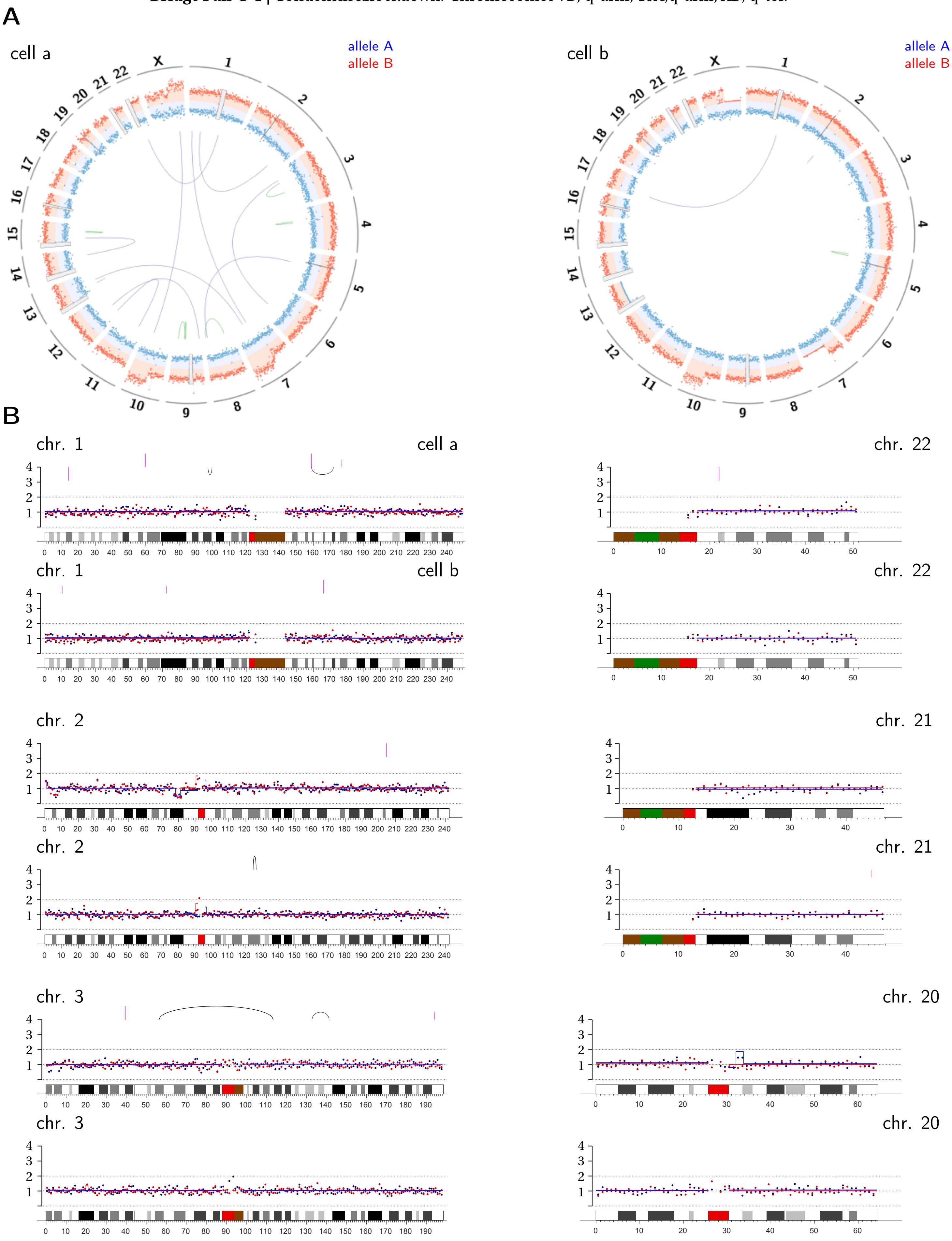

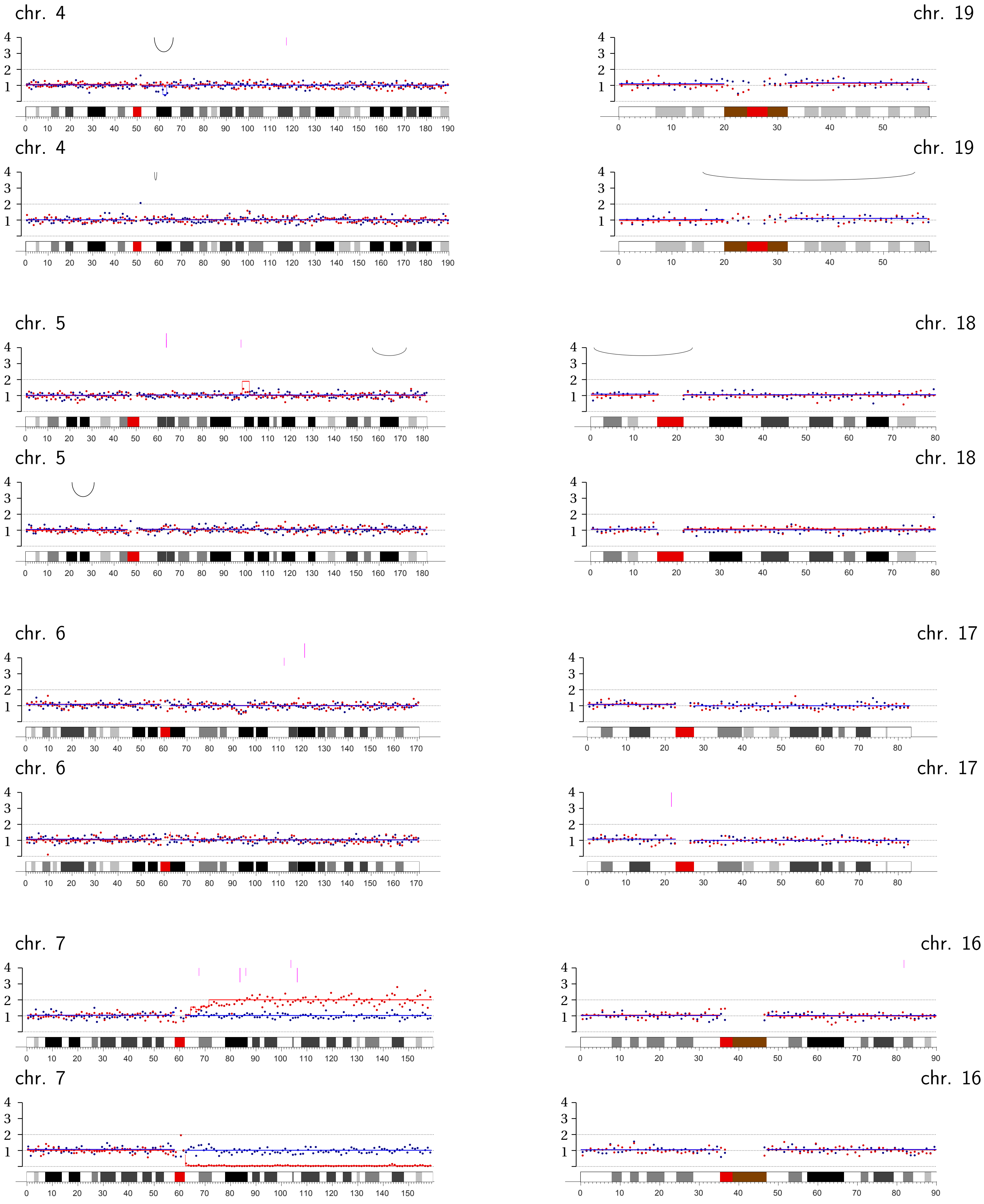
chr. 4
chr. 19
chr. 4
chr. 19
chr. 5
chr. 18
chr. 5
chr. 18
chr. 6
chr. 17
chr. 6
chr. 17
chr. 7
chr. 16
chr. 7
chr. 16

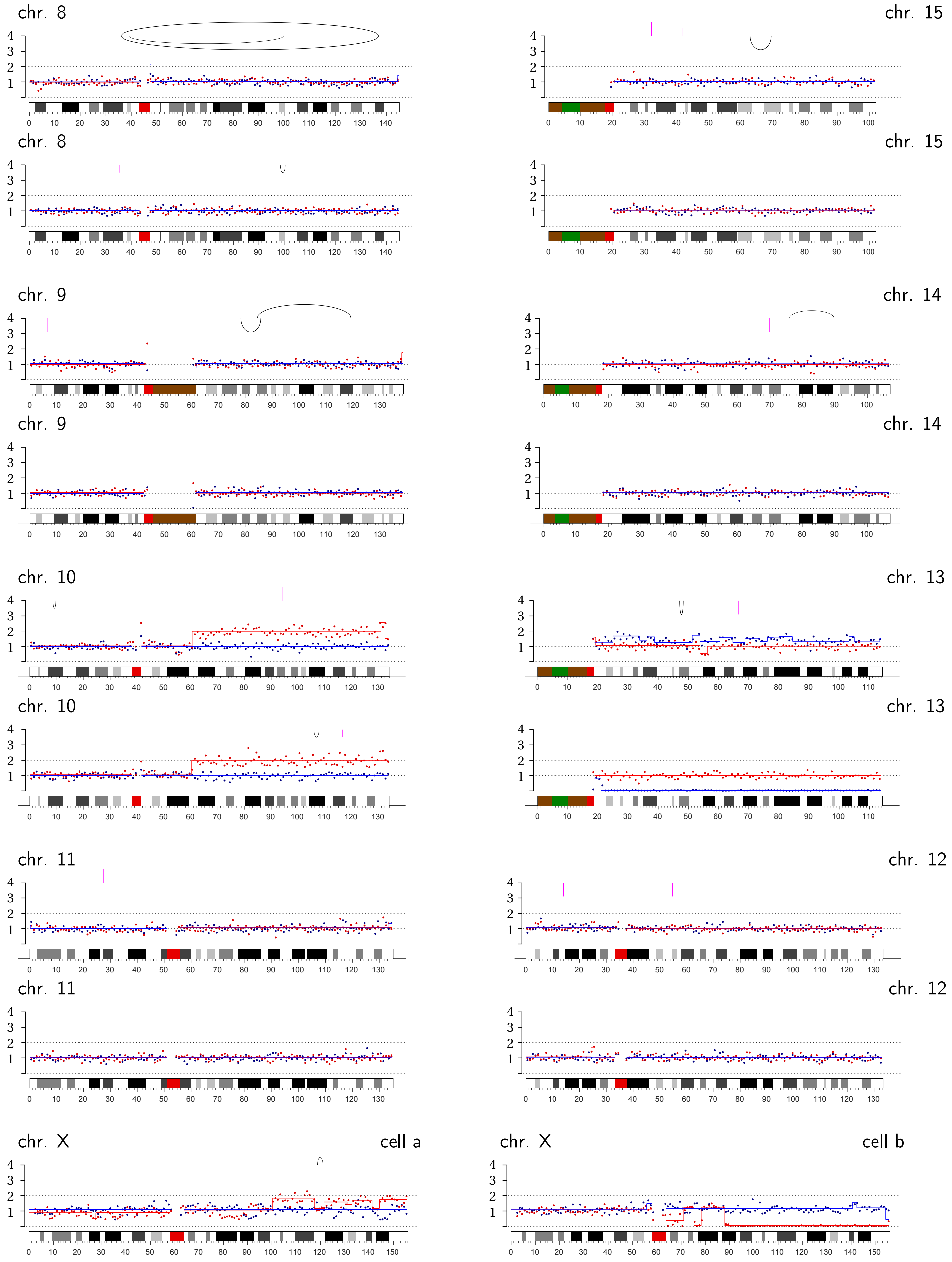
chr. 8
chr. 15
chr. 8
chr. 15
chr. 9
chr. 14
chr. 9
chr. 14
chr. 10
chr. 13
chr. 10
chr. 13
chr. 11
chr. 12
chr. 11
chr. 12
chr. X
cell a
chr. X
cell b

**Bridge Pair C-2 |** Condensin knockdown: **Chromosomes 4B, 5A, 6A, all on q-ter.**; Chr. Xq segment likely in a micronucleus.

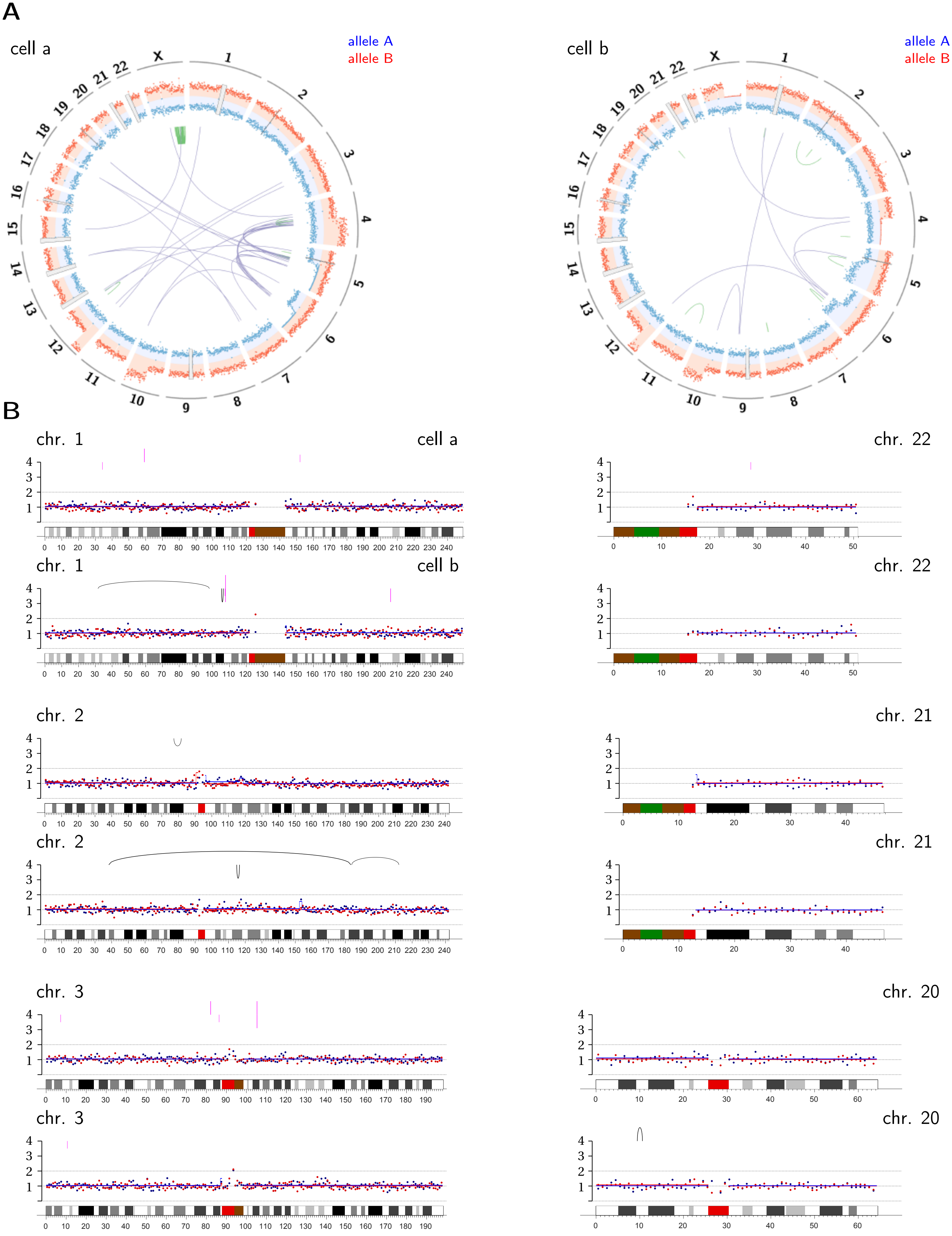

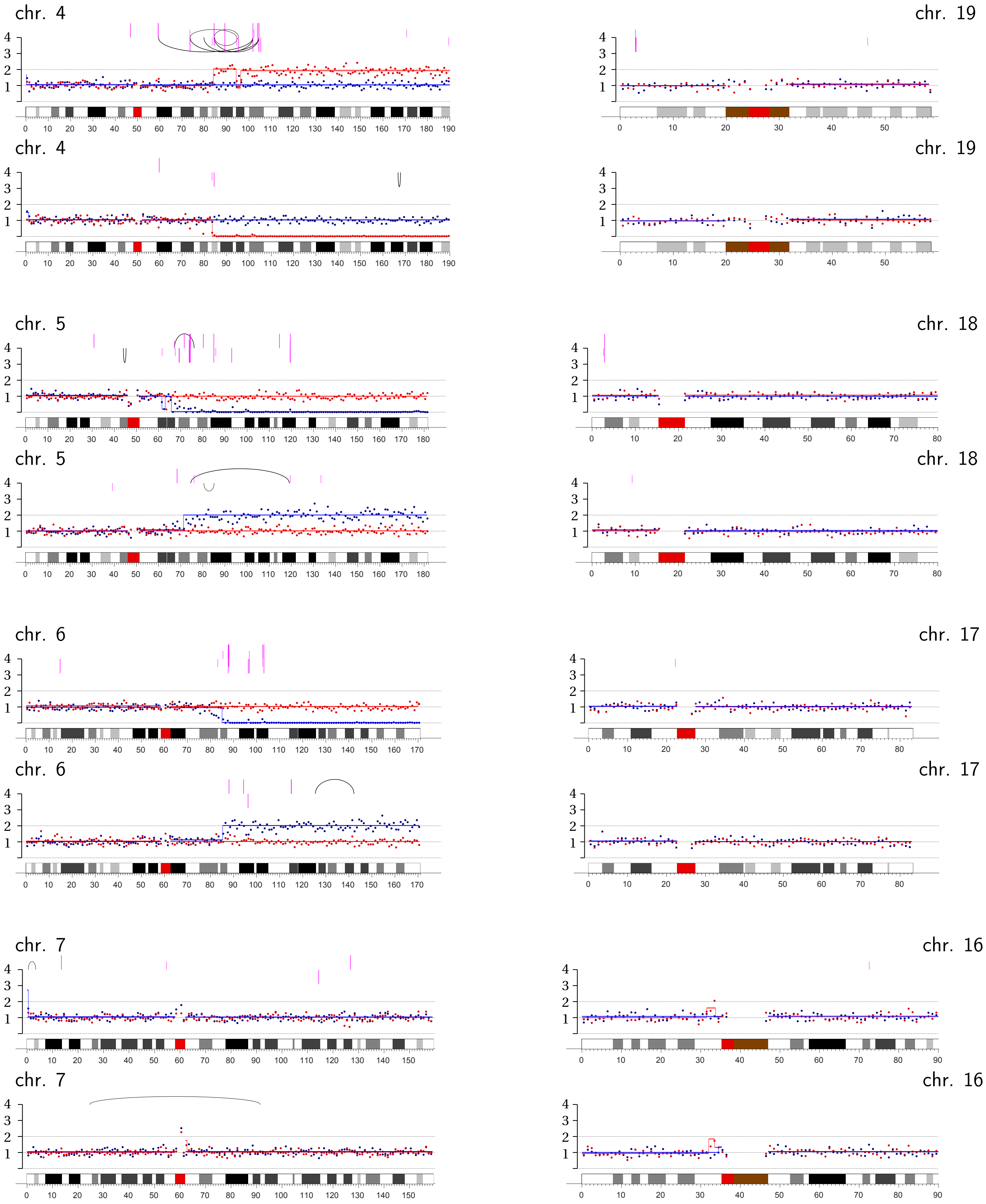
chr. 4
chr. 19
chr. 4
chr. 19
chr. 5
chr. 18
chr. 5
chr. 18
chr. 6
chr. 17
chr. 6
chr. 17
chr. 7
chr. 16
chr. 7
chr. 16

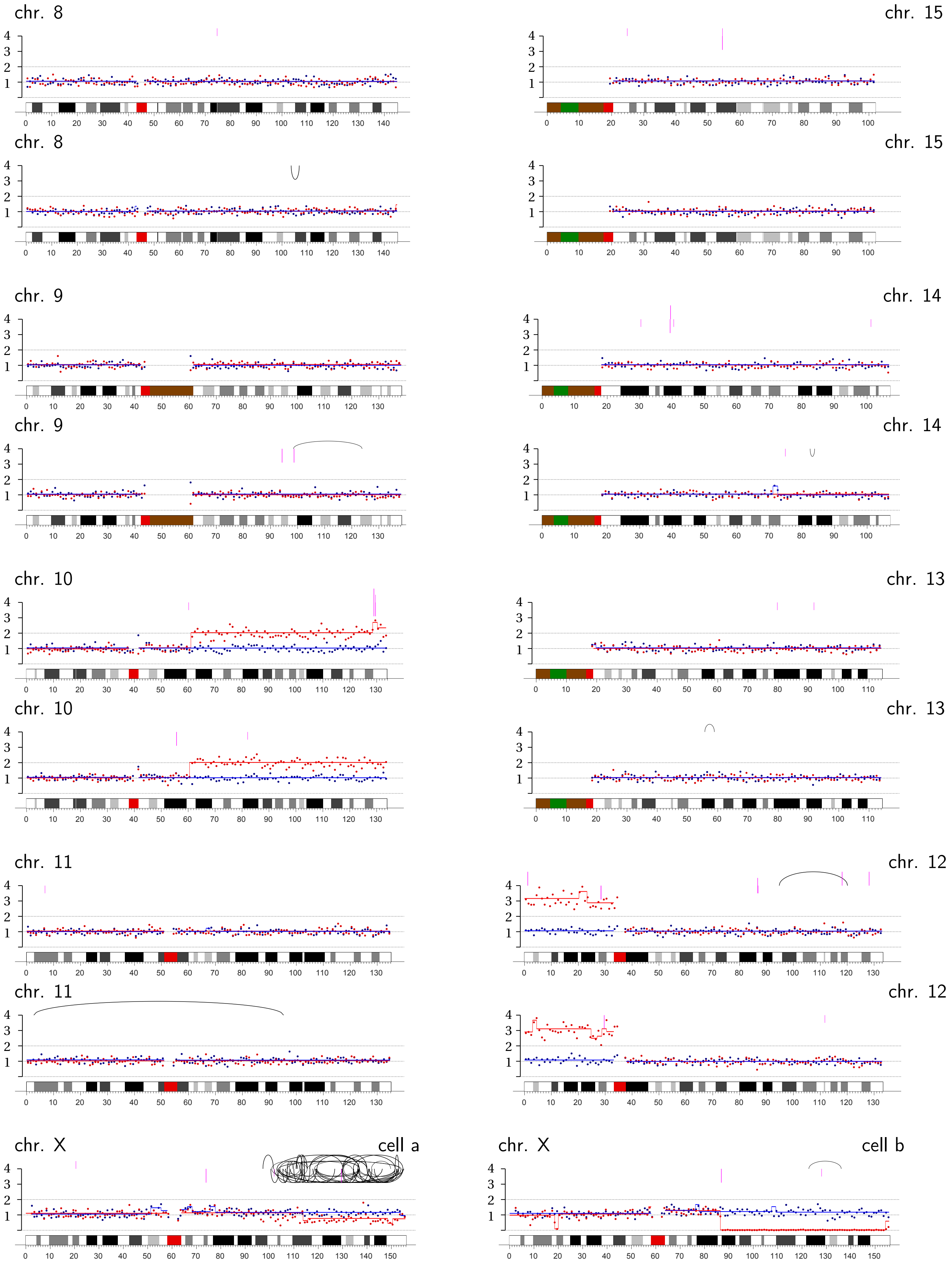
chr. 8
chr. 15
chr. 8
chr. 15
chr. 9
chr. 14
chr. 9
chr. 14
chr. 10
chr. 13
chr. 10
chr. 13
chr. 11
chr. 12
chr. 11
chr. 12
chr. X
cell a
chr. X
cell b

**Bridge Pair C-3** | Condensin knockdown: Chromosome 1A, p-arm internal.

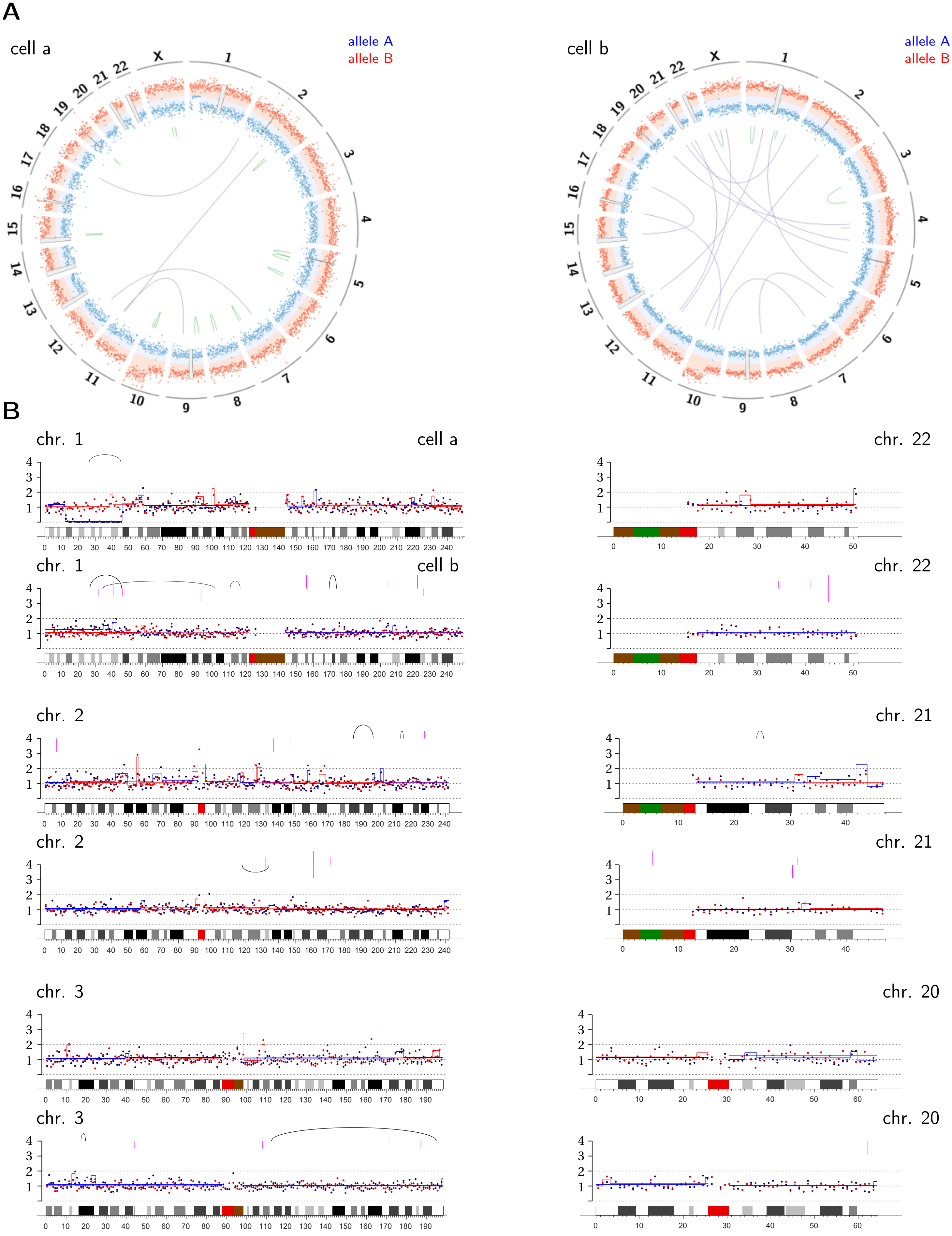

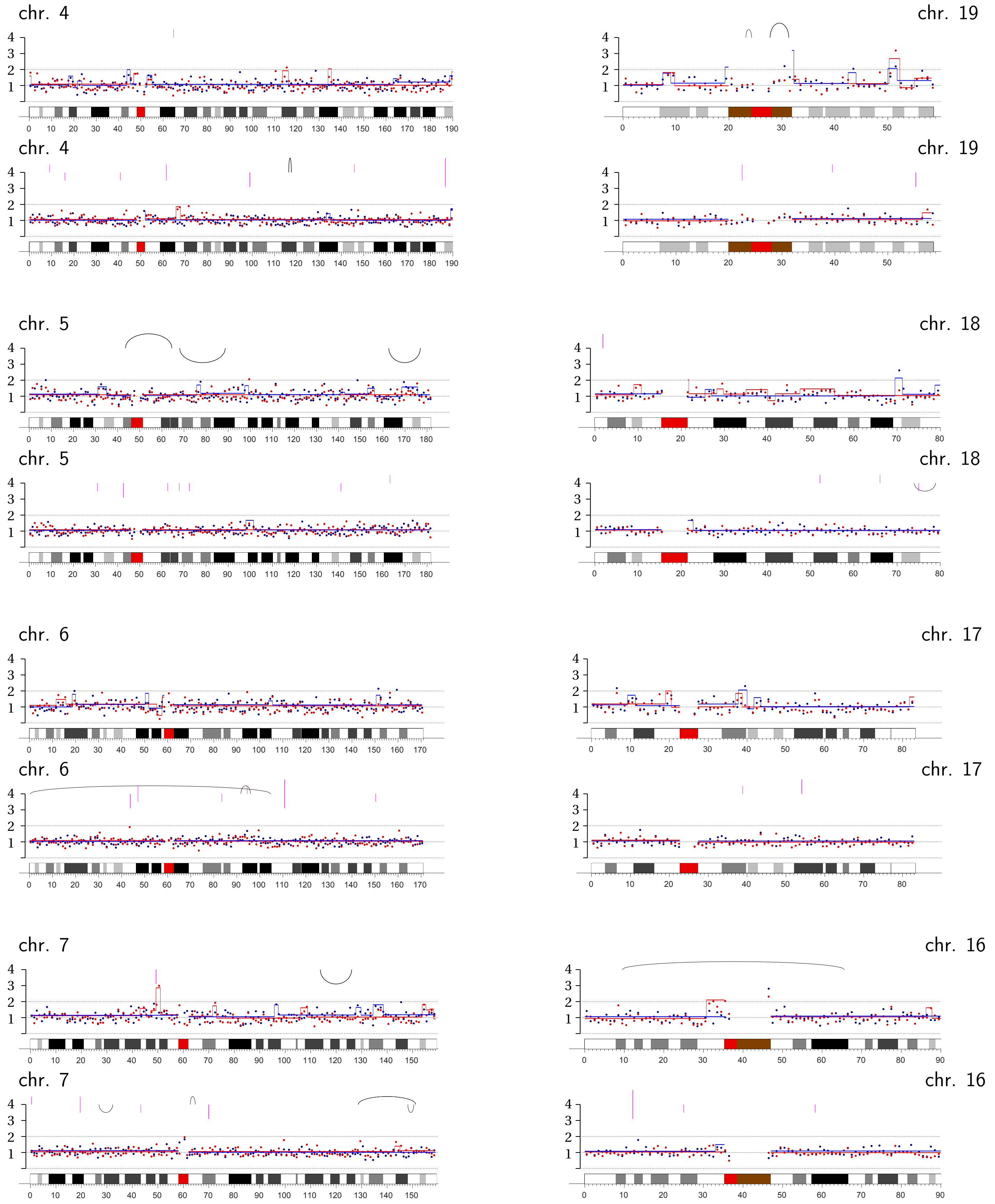
chr. 4
chr. 19
chr. 4
chr. 19
chr. 5
chr. 18
chr. 5
chr. 18
chr. 6
chr. 17
chr. 6
chr. 17
chr. 7
chr. 16
chr. 7
chr. 16

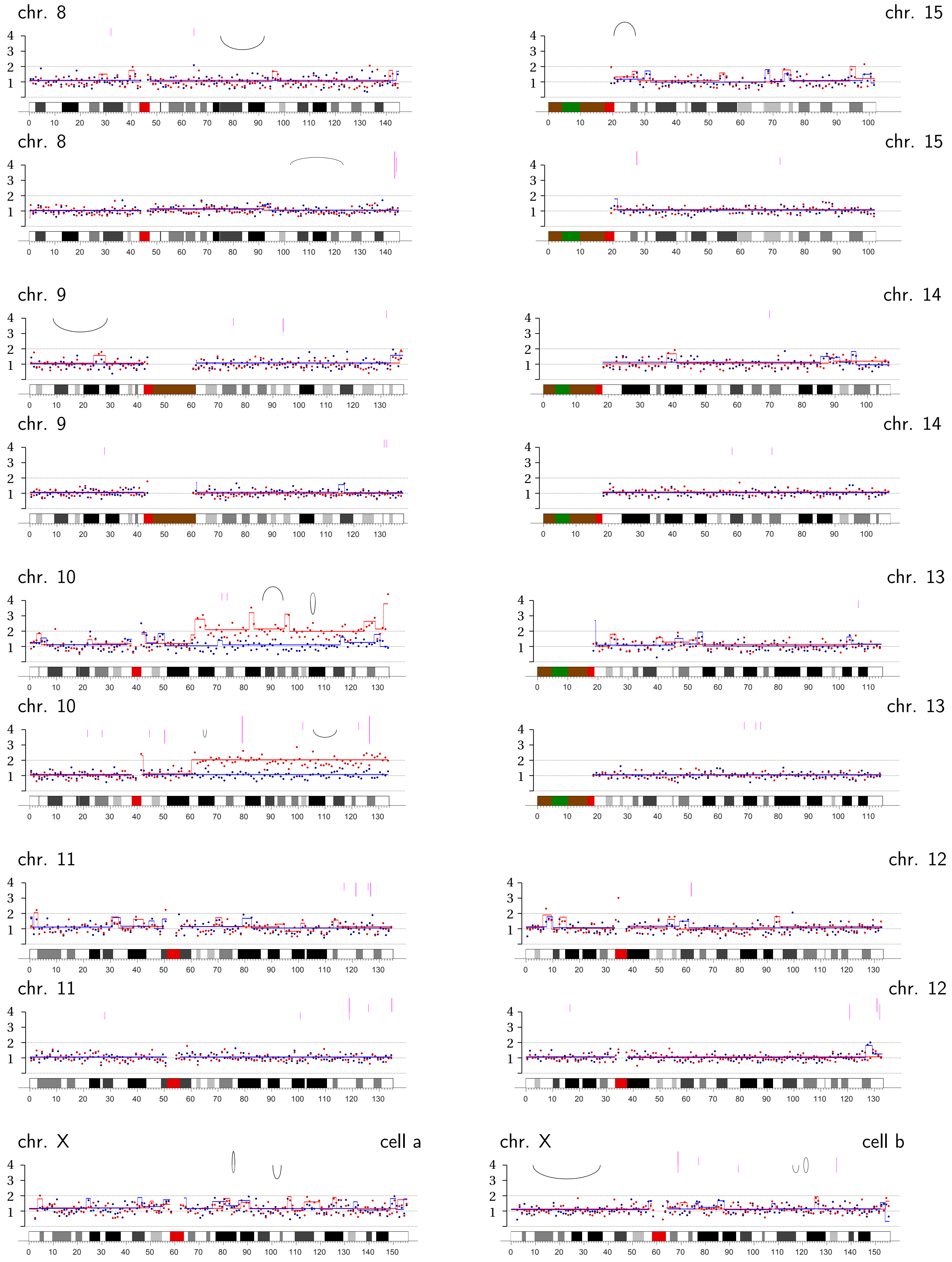
chr. 8
chr. 15
chr. 8
chr. 15
chr. 9
chr. 14
chr. 9
chr. 14
chr. 10
chr. 13
chr. 10
chr. 13
chr. 11
chr. 12
chr. 11
chr. 12
chr. X
cell a
chr. X
cell b

**Bridge Pair C-4** | Condensin knockdown: **Chromosomes 16A and 18B, p-ter.**

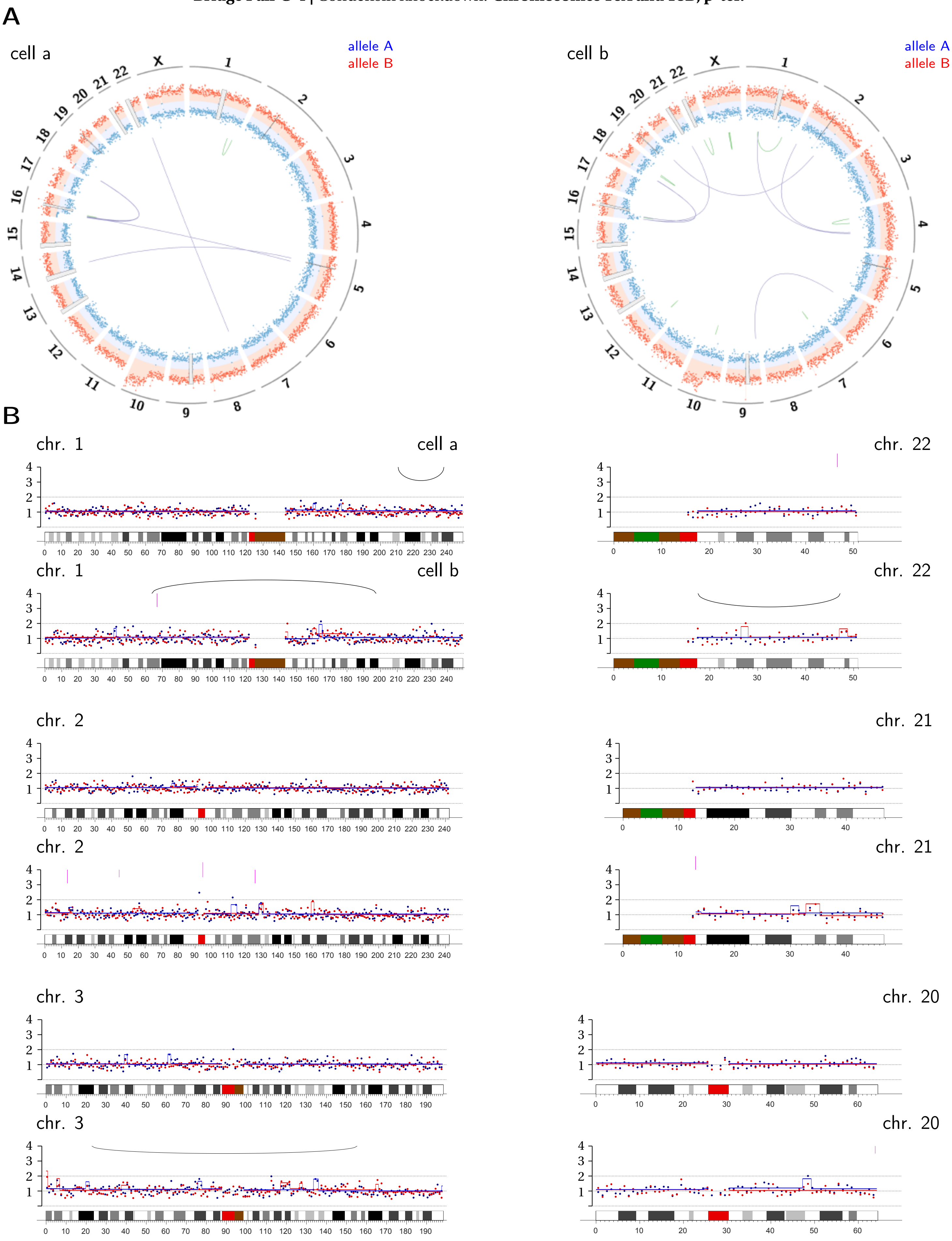

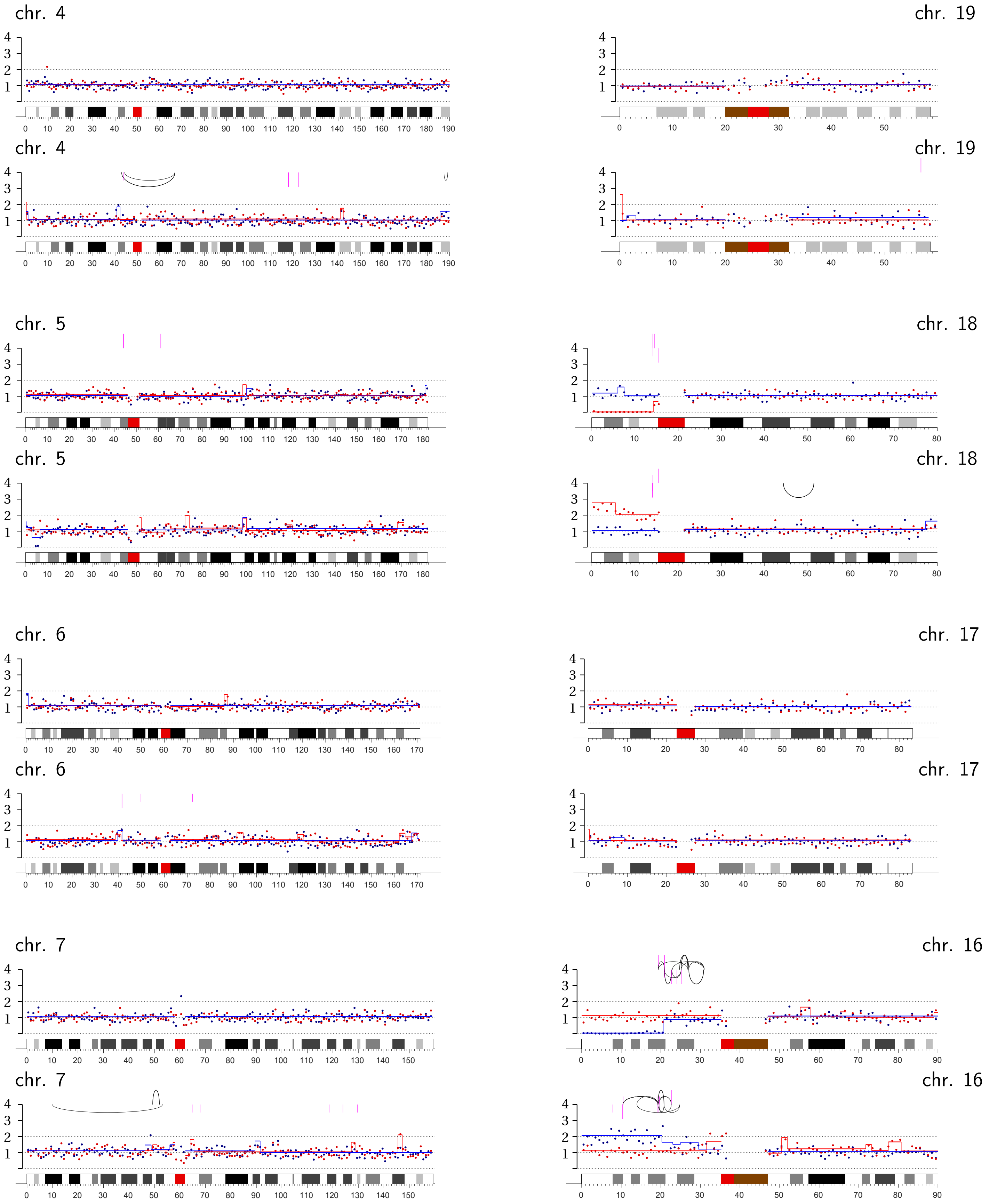
chr. 4
chr. 19
chr. 4
chr. 19
chr. 5
chr. 18
chr. 5
chr. 18
chr. 6
chr. 17
chr. 6
chr. 17
chr. 7
chr. 16
chr. 7
chr. 16

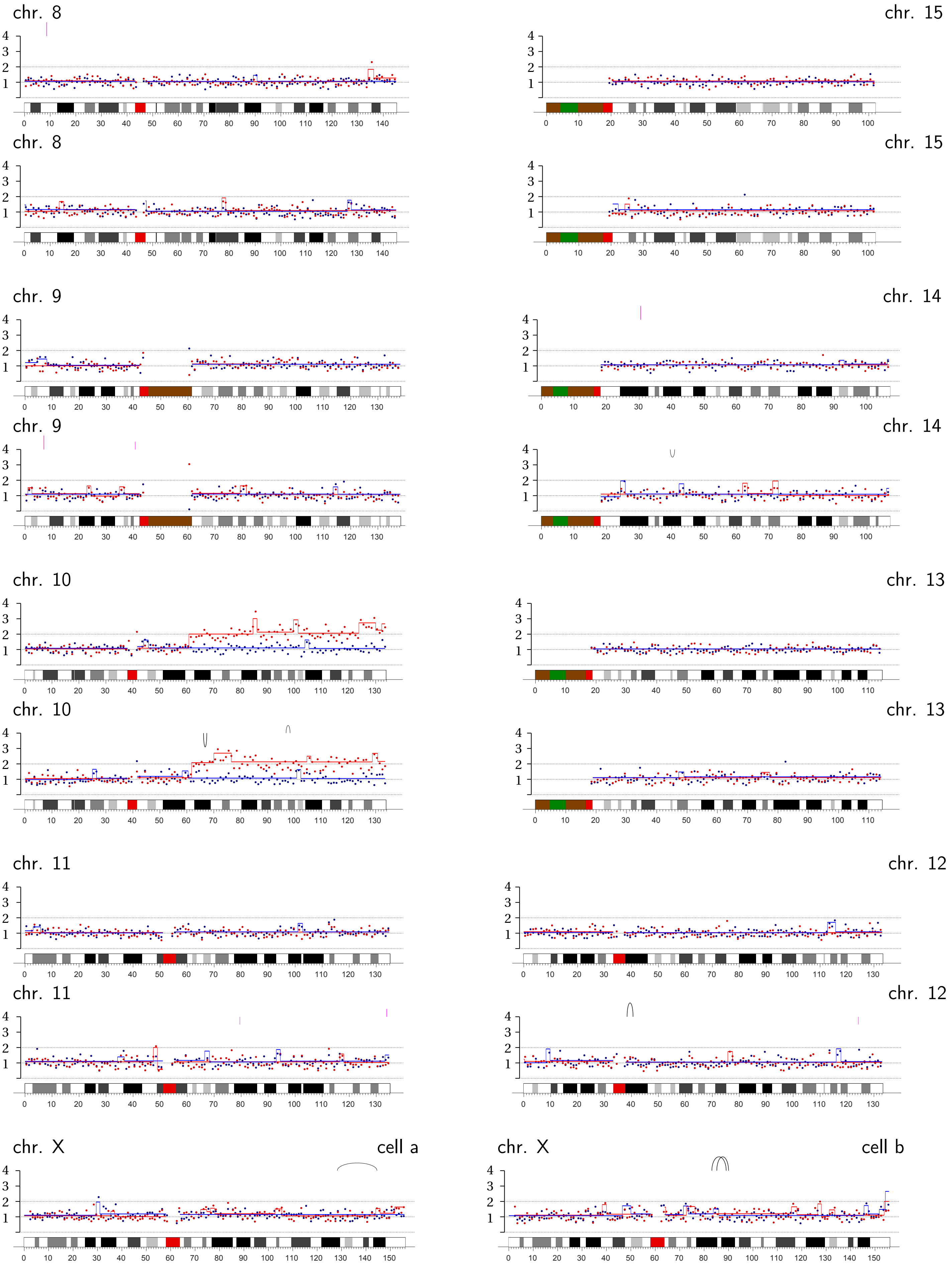
chr. 8
chr. 15
chr. 8
chr. 15
chr. 9
chr. 14
chr. 9
chr. 14
chr. 10
chr. 13
chr. 10
chr. 13
chr. 11
chr. 12
chr. 11
chr. 12
chr. X
cell a
chr. X
cell b

**Bridge Pair T-1** | Transient TRF2-DN expression: **Chromosome 3A, q-ter.**.

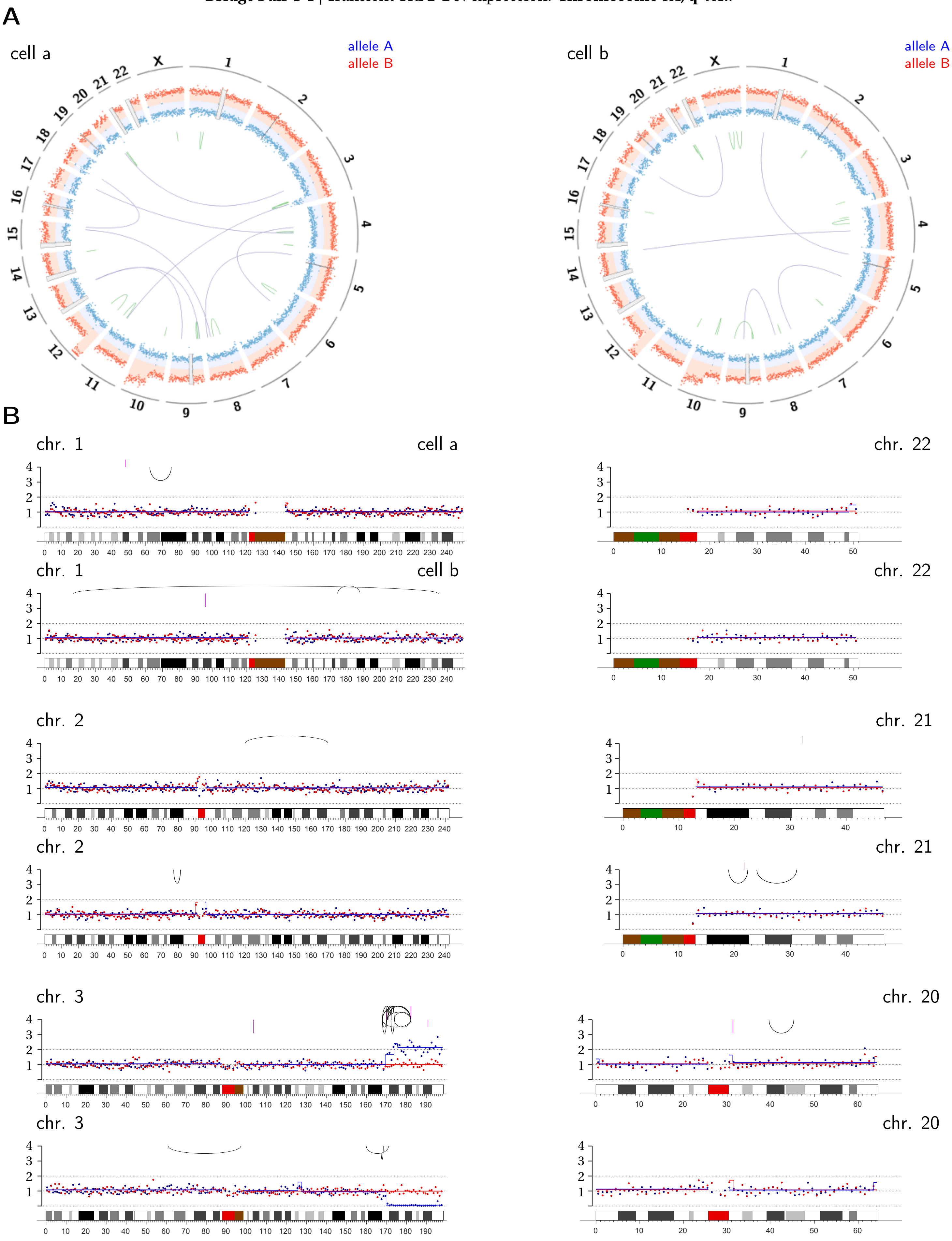

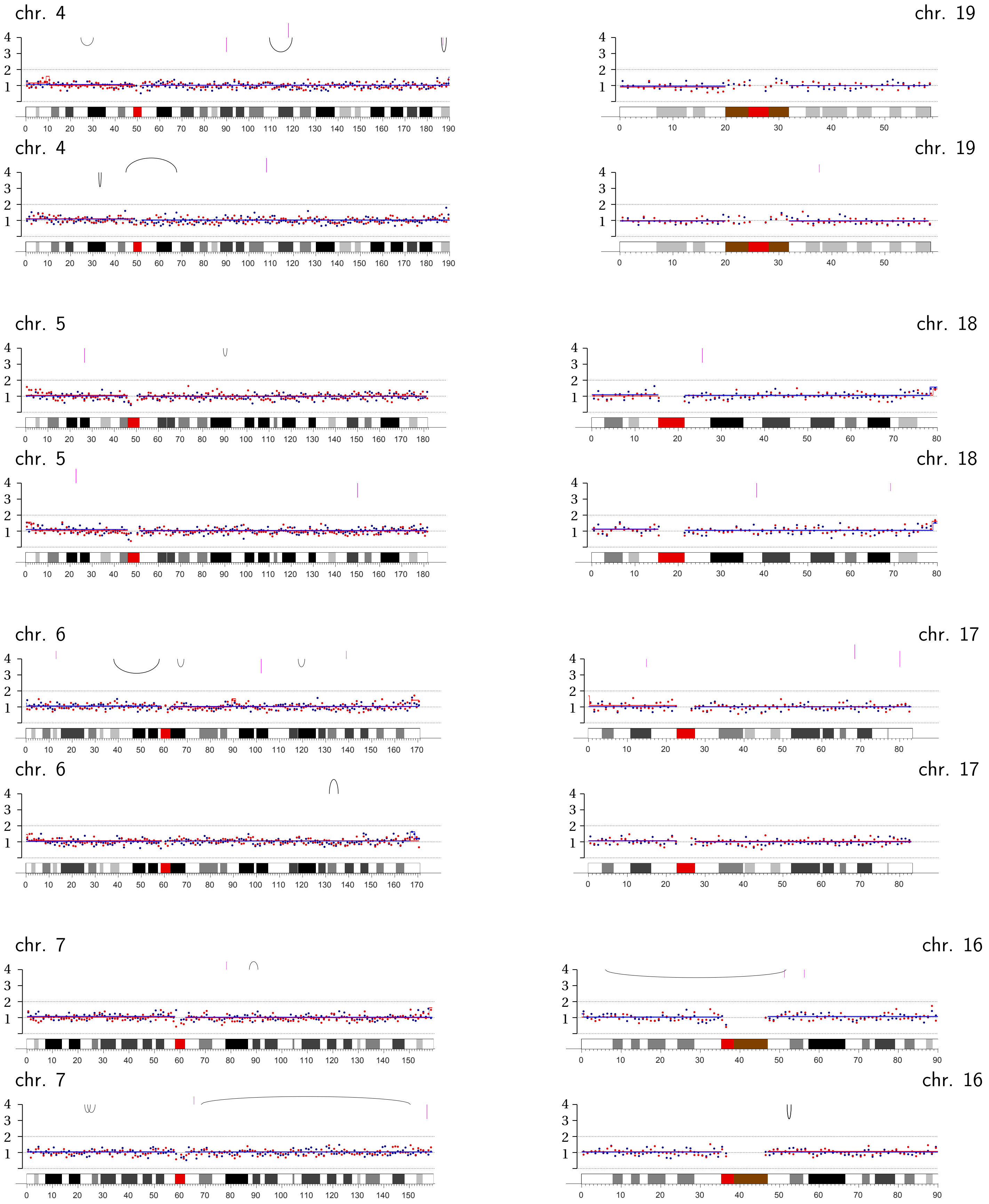
chr. 4
chr. 19
chr. 4
chr. 19
chr. 5
chr. 18
chr. 5
chr. 18
chr. 6
chr. 17
chr. 6
chr. 17
chr. 7
chr. 16
chr. 7
chr. 16

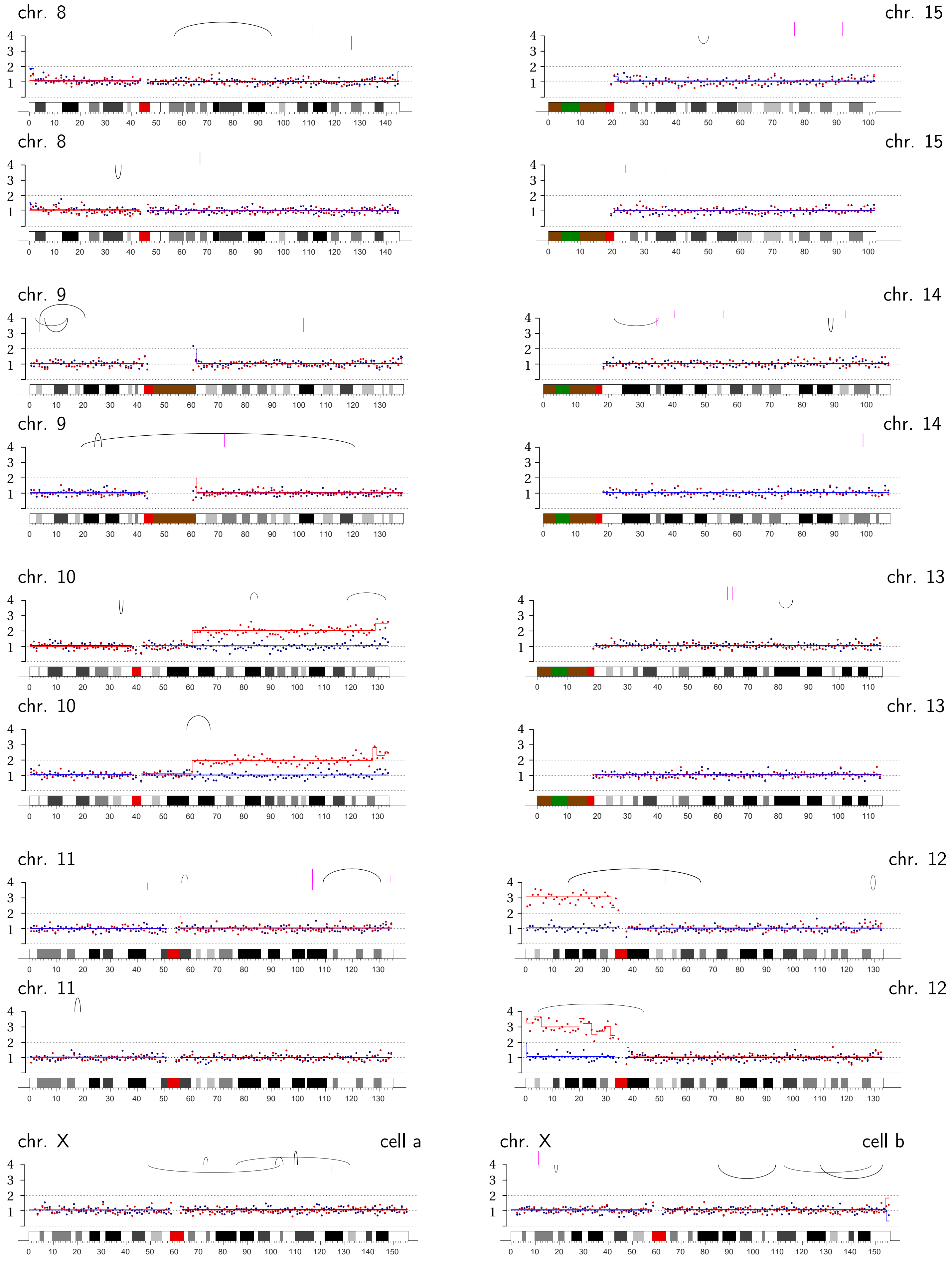
chr. 8
chr. 15
chr. 8
chr. 15
chr. 9
chr. 14
chr. 9
chr. 14
chr. 10
chr. 13
chr. 10
chr. 13
chr. 11
chr. 12
chr. 11
chr. 12
chr. X
cell a
chr. X
cell b

**Bridge Pair T-2** | Transient TRF2-DN expression: Chromosome 2B, q-ter.

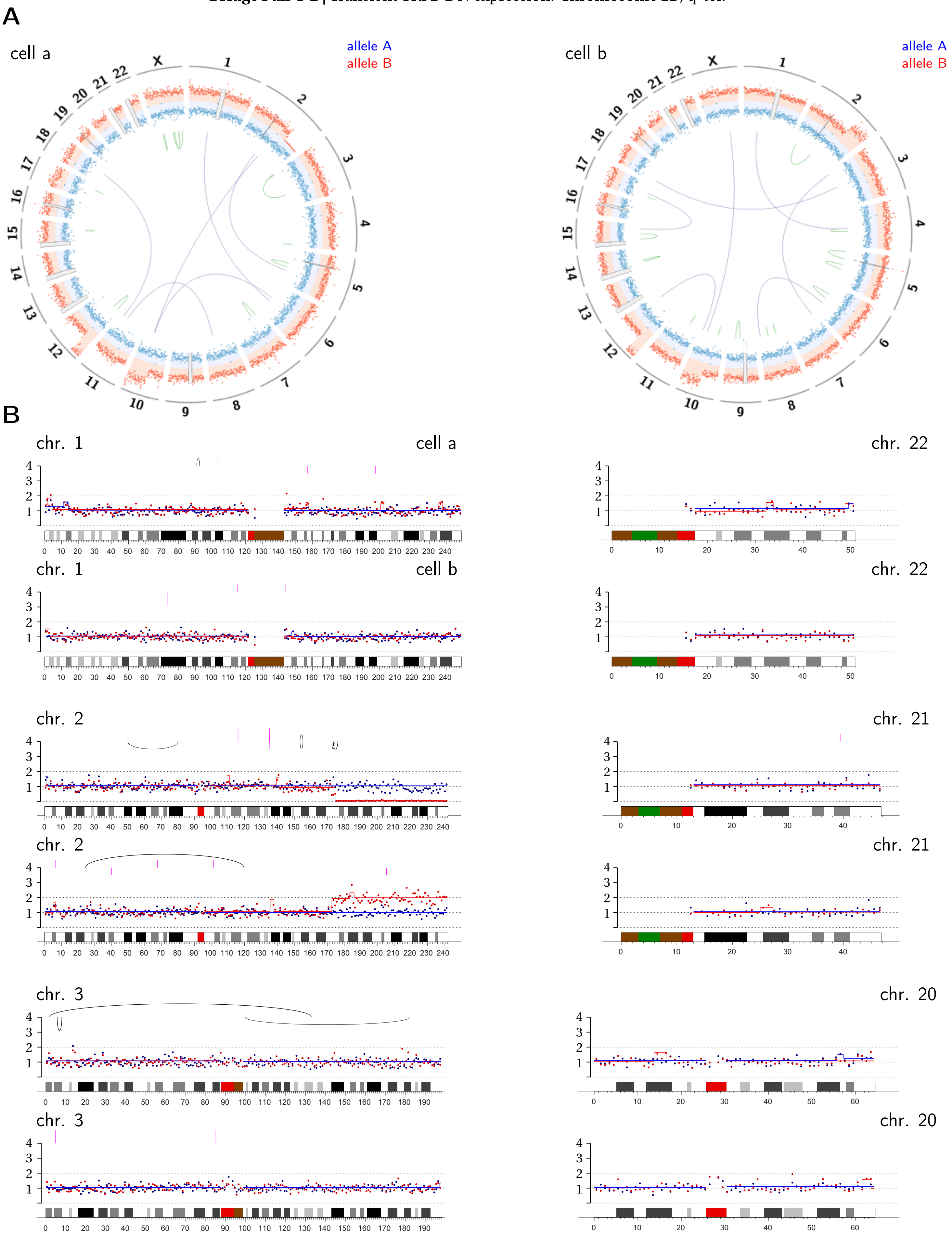

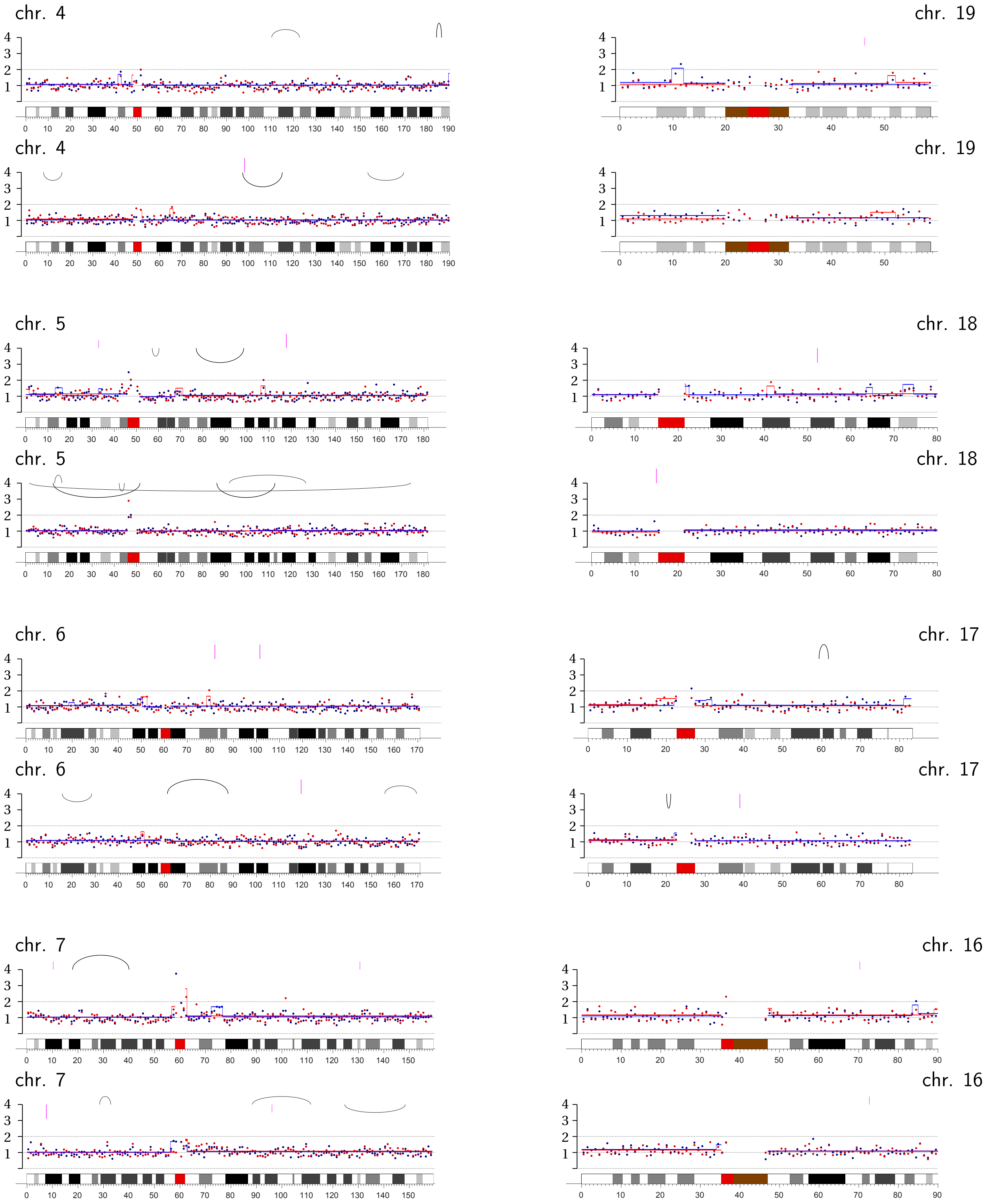
chr. 4
chr. 19
chr. 4
chr. 19
chr. 5
chr. 18
chr. 5
chr. 18
chr. 6
chr. 17
chr. 6
chr. 17
chr. 7
chr. 16
chr. 7
chr. 16

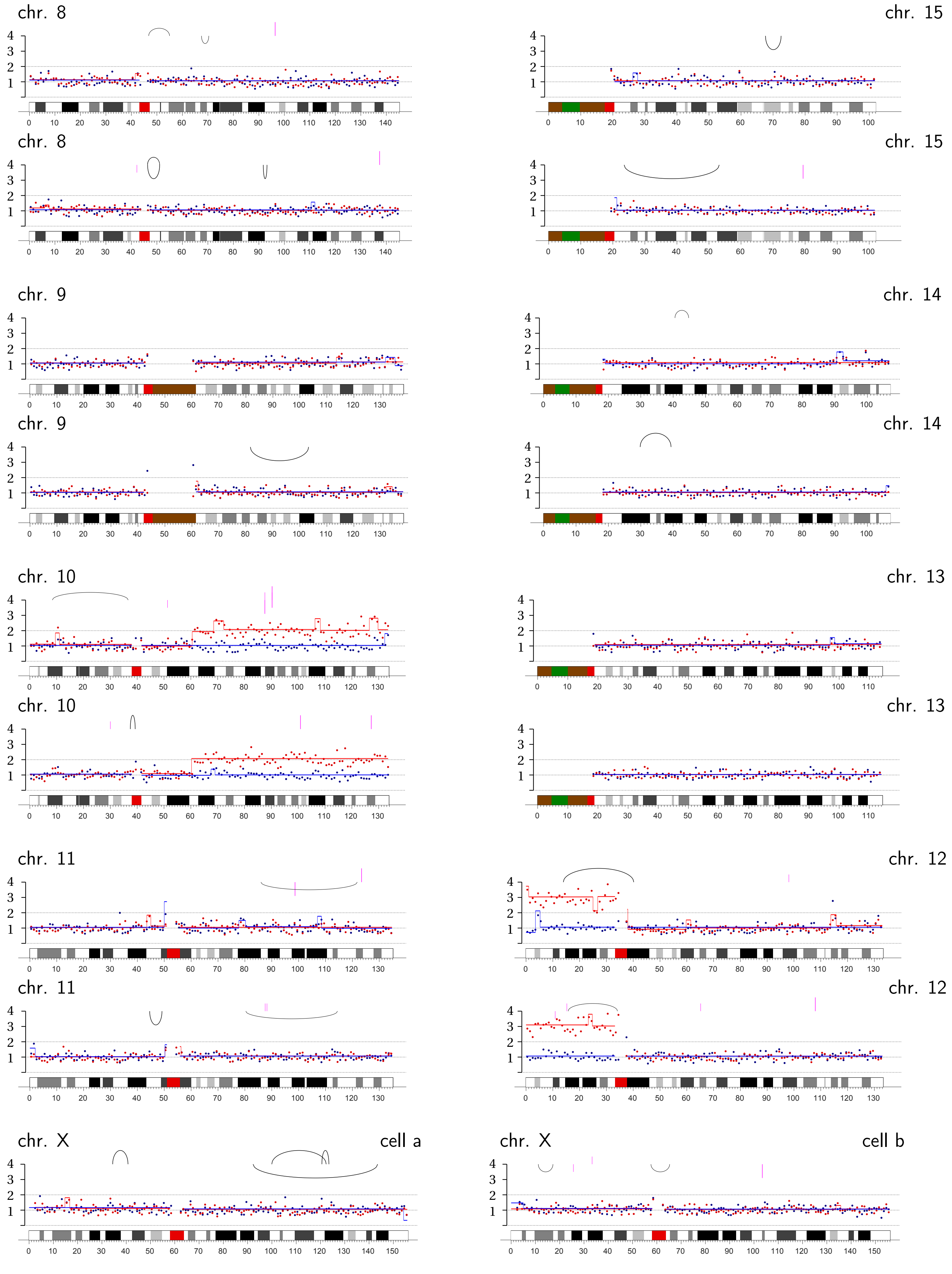
chr. 8
chr. 15
chr. 8
chr. 15
chr. 9
chr. 14
chr. 9
chr. 14
chr. 10
chr. 13
chr. 10
chr. 13
chr. 11
chr. 12
chr. 11
chr. 12
chr. X
cell a
chr. X
cell b

**Bridge Pair T-3** | Transient TRF2-DN expression: Chromosomes 7B, p-ter.; Chromosome 12B, p-arm; 19B, p-arm.

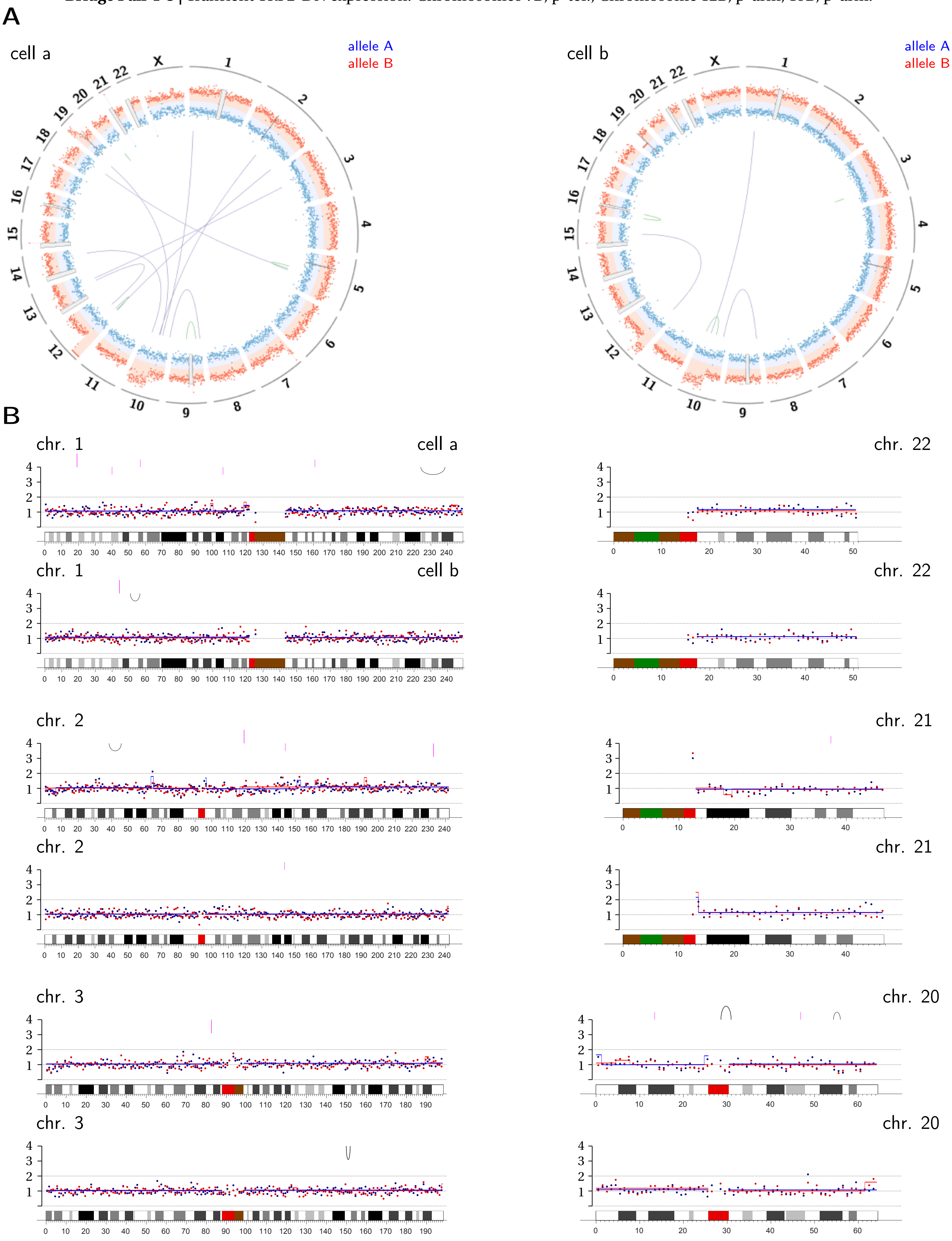

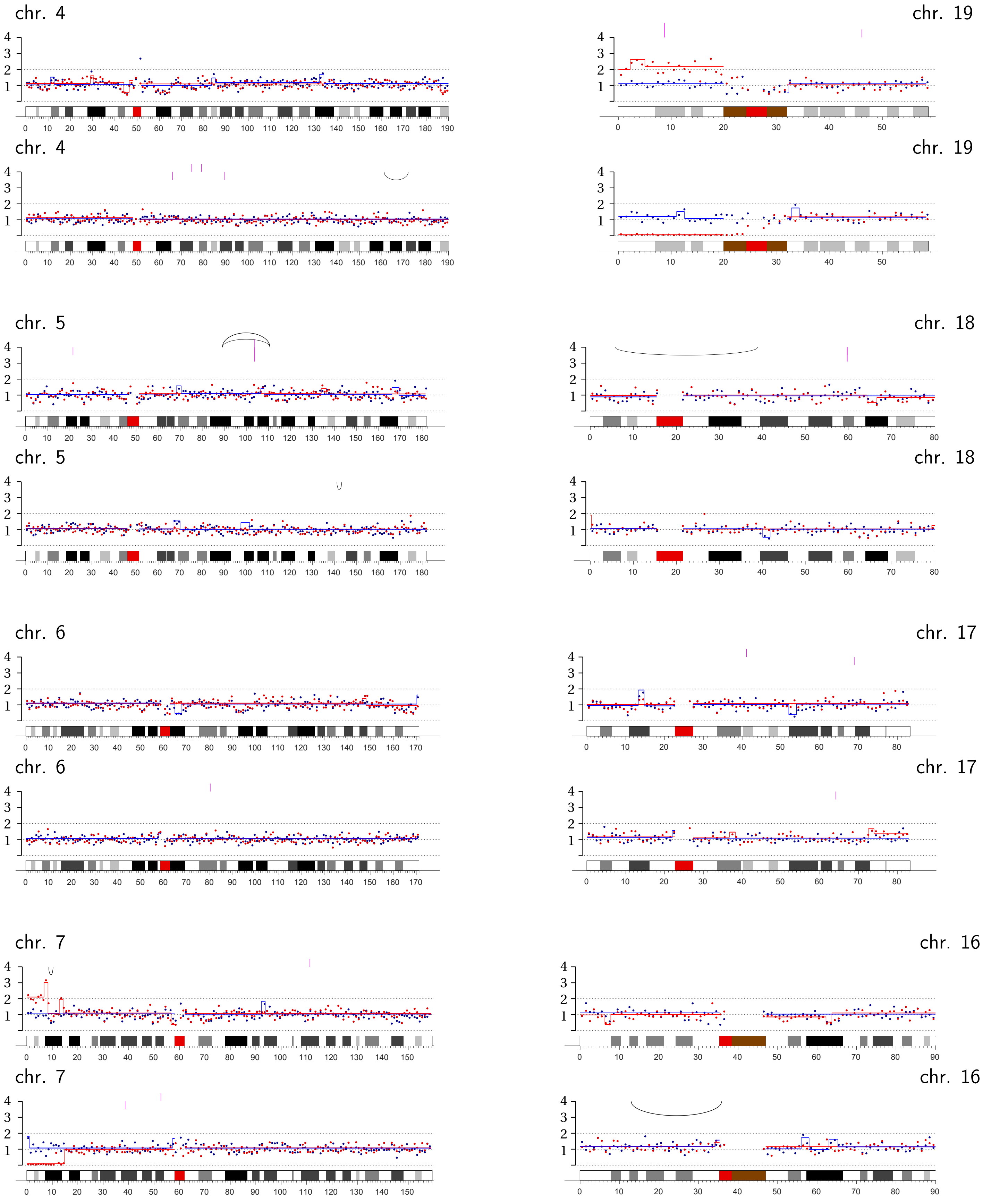
chr. 4
chr. 19
chr. 4
chr. 19
chr. 5
chr. 18
chr. 5
chr. 18
chr. 6
chr. 17
chr. 6
chr. 17
chr. 7
chr. 16
chr. 7
chr. 16

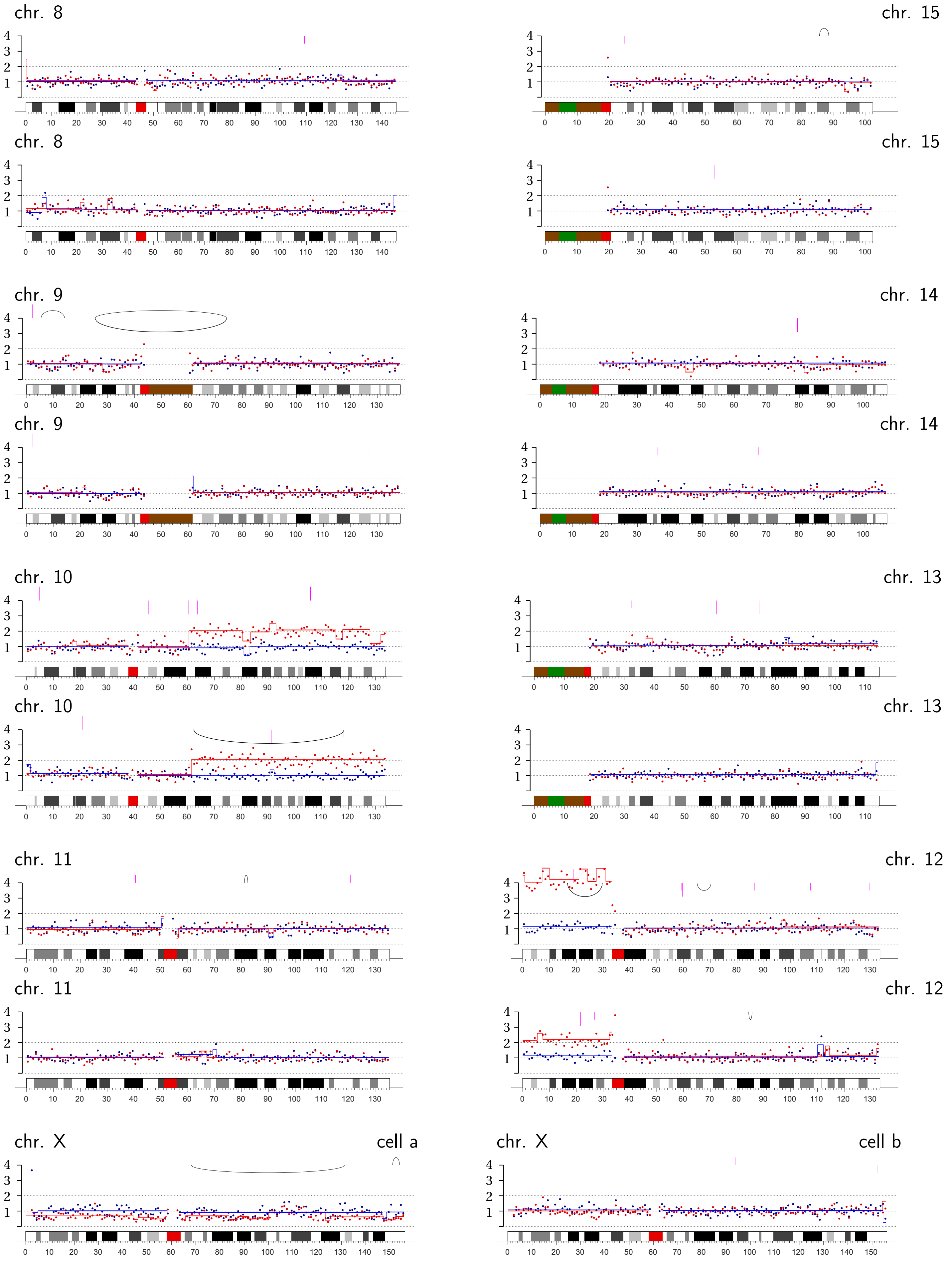
chr. 8
chr. 15
chr. 8
chr. 15
chr. 9
chr. 14
chr. 9
chr. 14
chr. 10
chr. 13
chr. 10
chr. 13
chr. 11
chr. 12
chr. 11
chr. 12
chr. X
cell a
chr. X
cell b

**Bridge Pair T-4 |** Transient TRF2-DN expression: Chromosome 18B, q-arm internal; incomplete loss of Chromosomes 17B and 21A, q-ter.

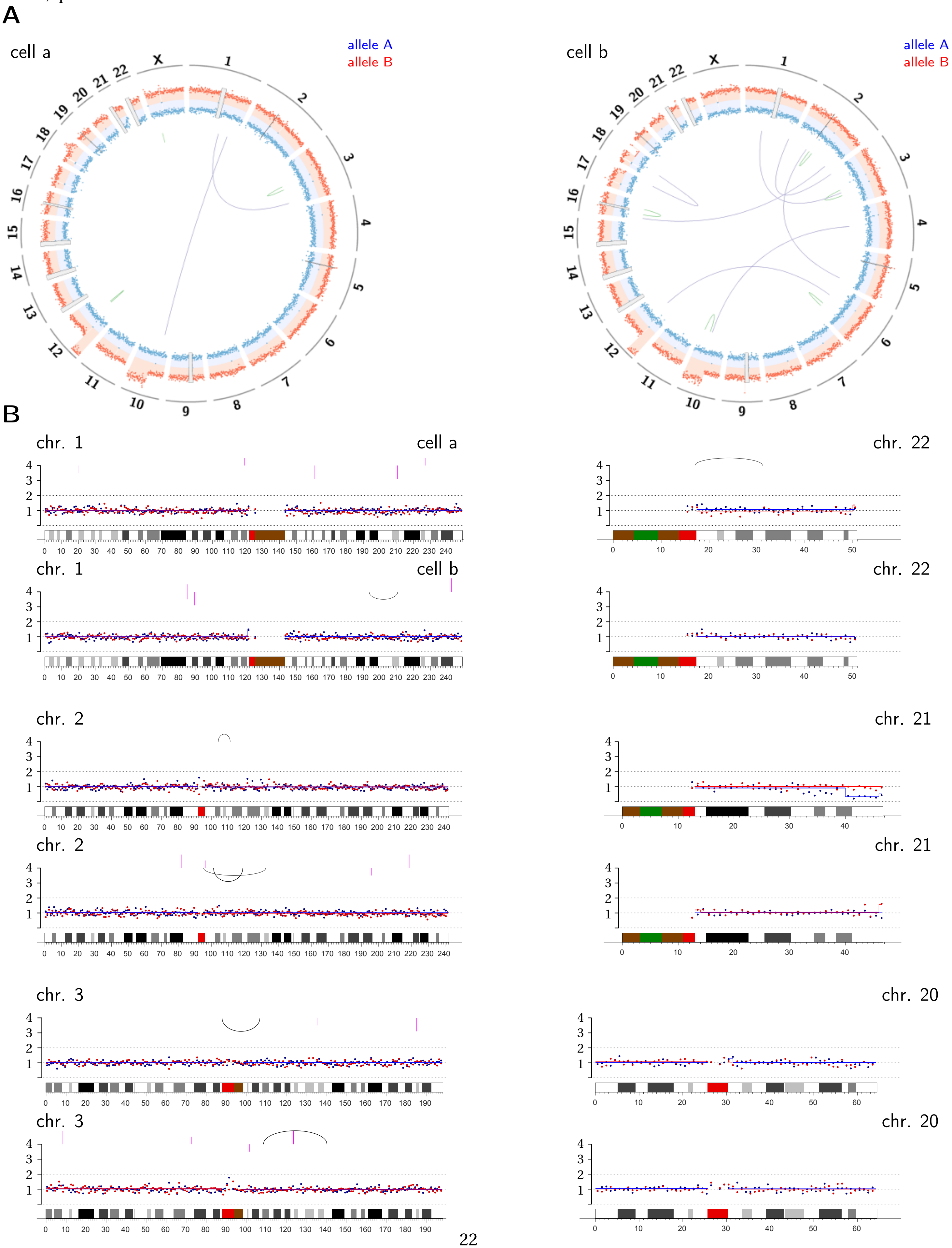

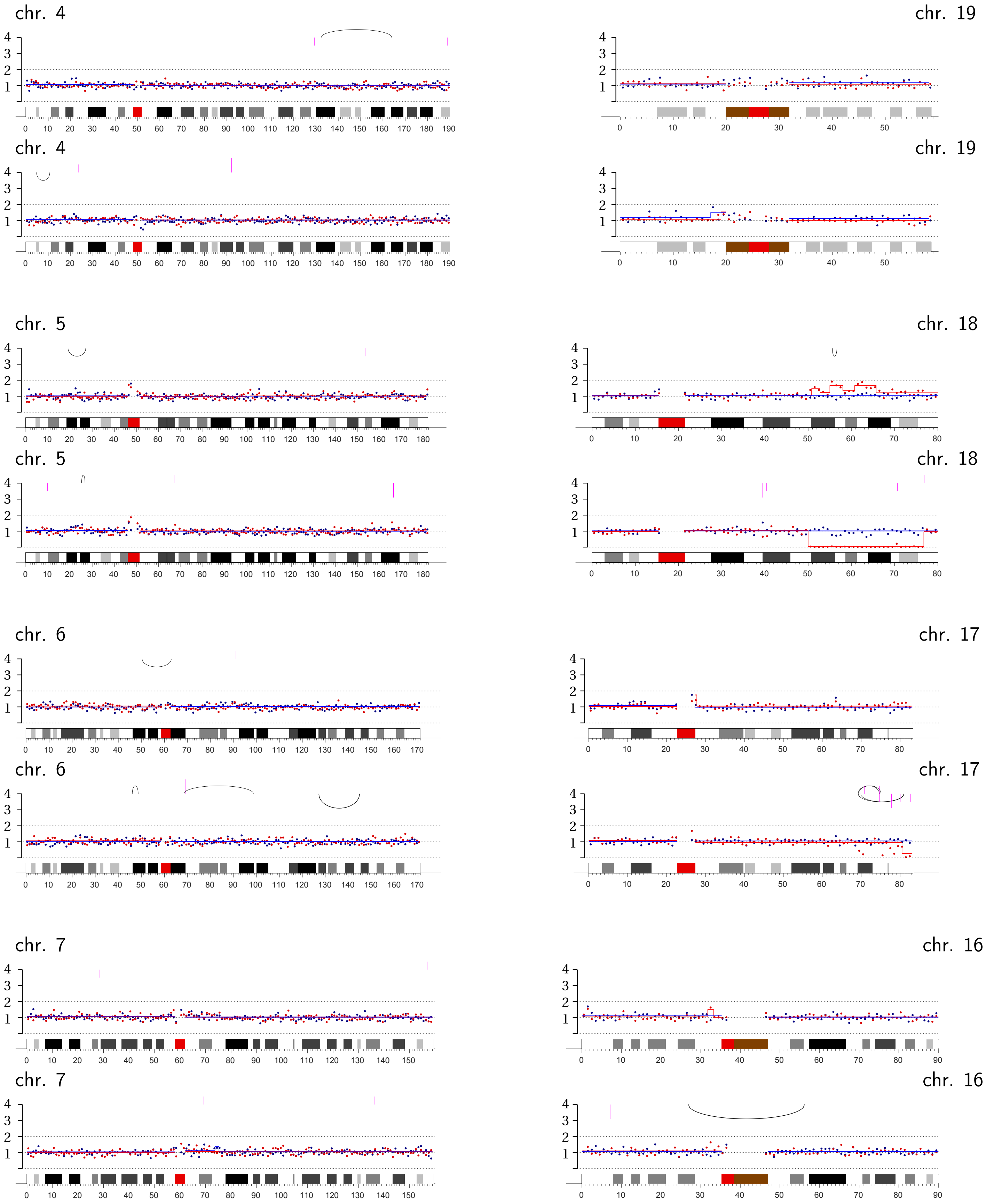
chr. 4
chr. 19
chr. 4
chr. 19
chr. 5
chr. 18
chr. 5
chr. 18
chr. 6
chr. 17
chr. 6
chr. 17
chr. 7
chr. 16
chr. 7
chr. 16

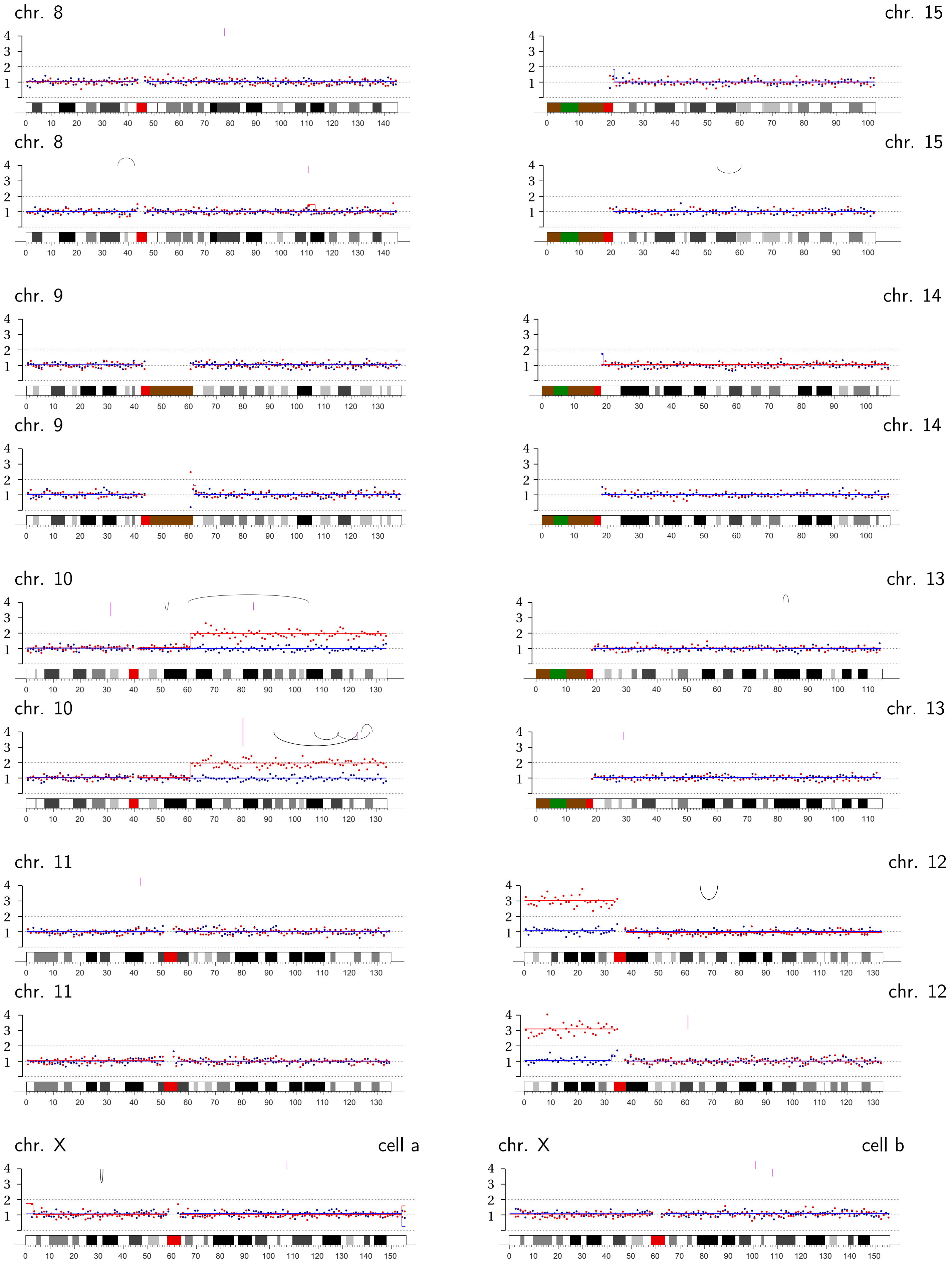
chr. 8
chr. 15
chr. 8
chr. 15
chr. 9
chr. 14
chr. 9
chr. 14
chr. 10
chr. 13
chr. 10
chr. 13
chr. 11
chr. 12
chr. 11
chr. 12
chr. X
cell a
chr. X
cell b

**Bridge Pair T-5 |** Transient TRF2-DN expression: Chromosomes 1B, p-ter.; 6B, q-ter.; 8A, q-ter.

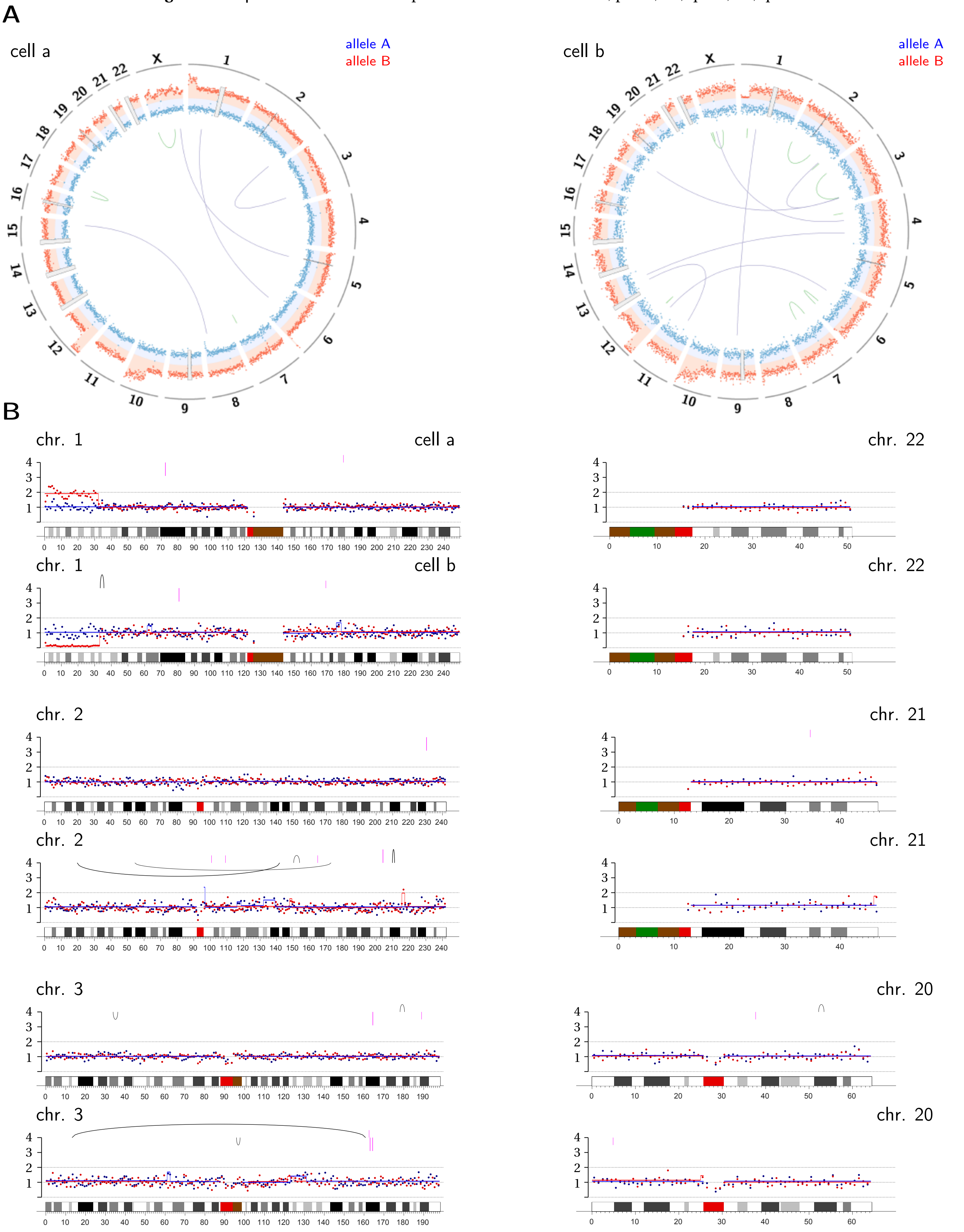

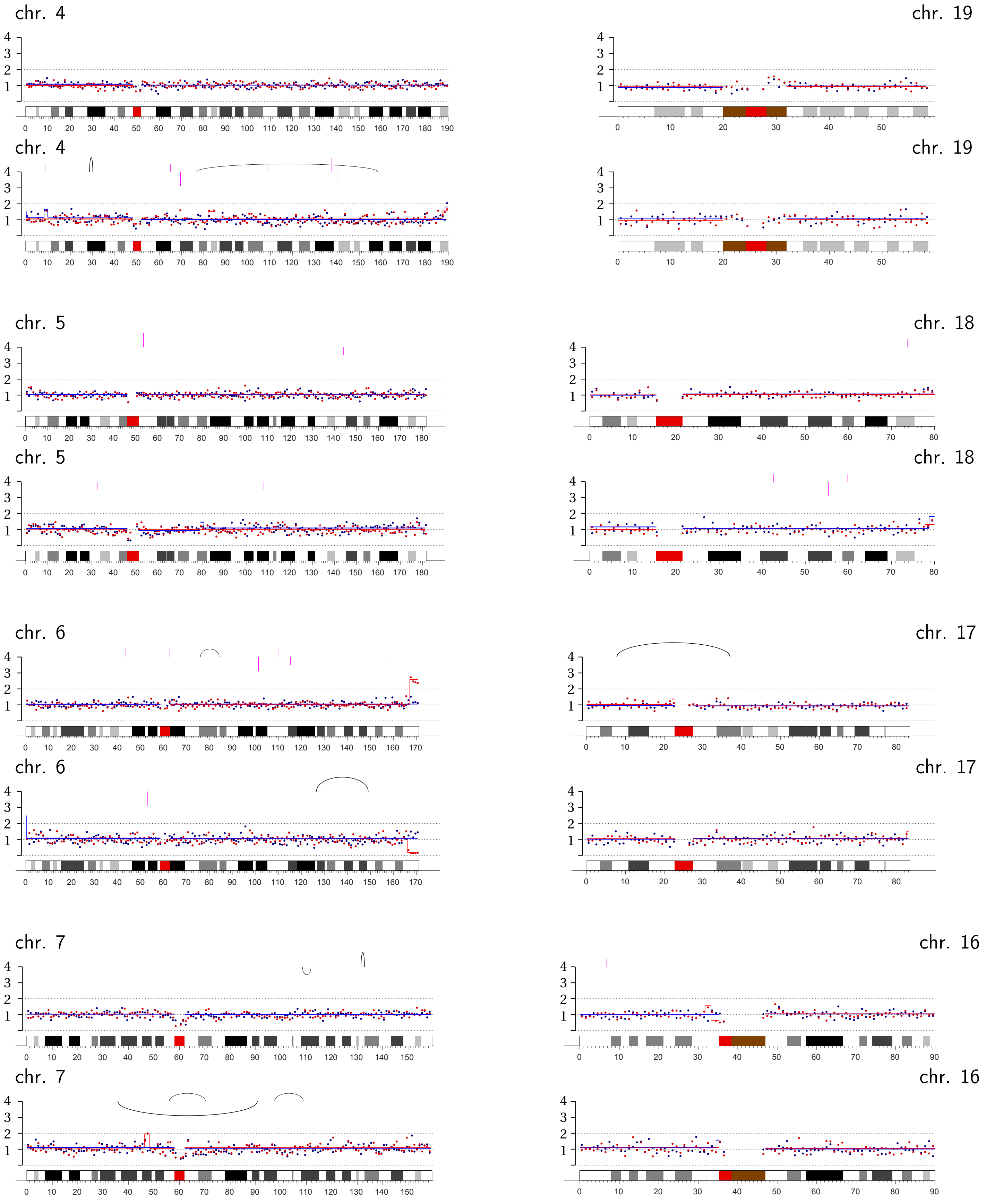
chr. 4
chr. 19
chr. 4
chr. 19
chr. 5
chr. 18
chr. 5
chr. 18
chr. 6
chr. 17
chr. 6
chr. 17
chr. 7
chr. 16
chr. 7
chr. 16

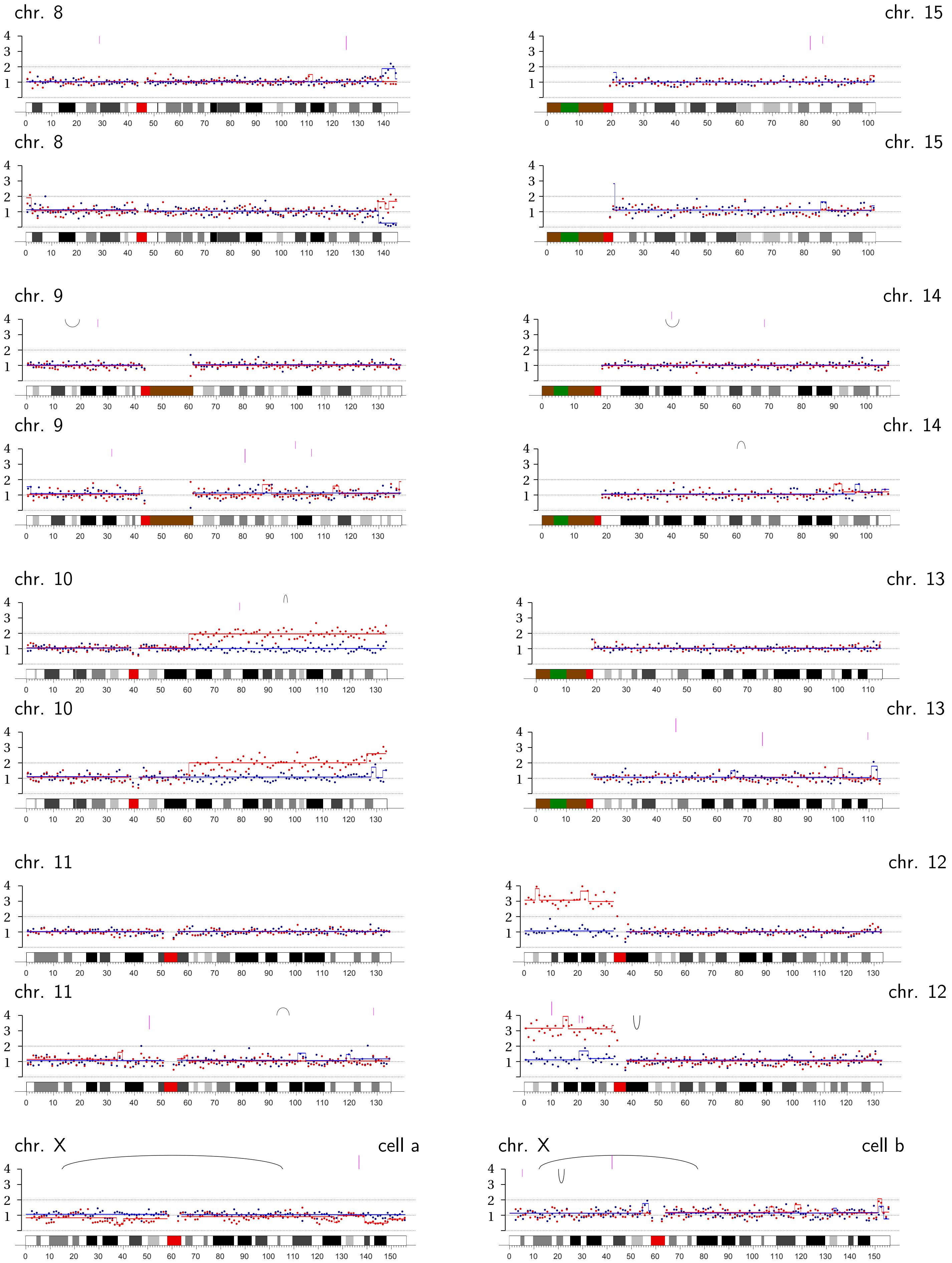
chr. 8
chr. 15
chr. 8
chr. 15
chr. 9
chr. 14
chr. 9
chr. 14
chr. 10
chr. 13
chr. 10
chr. 13
chr. 11
chr. 12
chr. 11
chr. 12
chr. X
cell a
chr. X
cell b

**Bridge Pair T-6 |** Transient TRF2-DN expression: Chromosomes 3B, p-ter.; 11A, q-ter.; 12A, p-ter.; 19B, p-arm; incomplete loss of Chromosome 20B on q-ter.

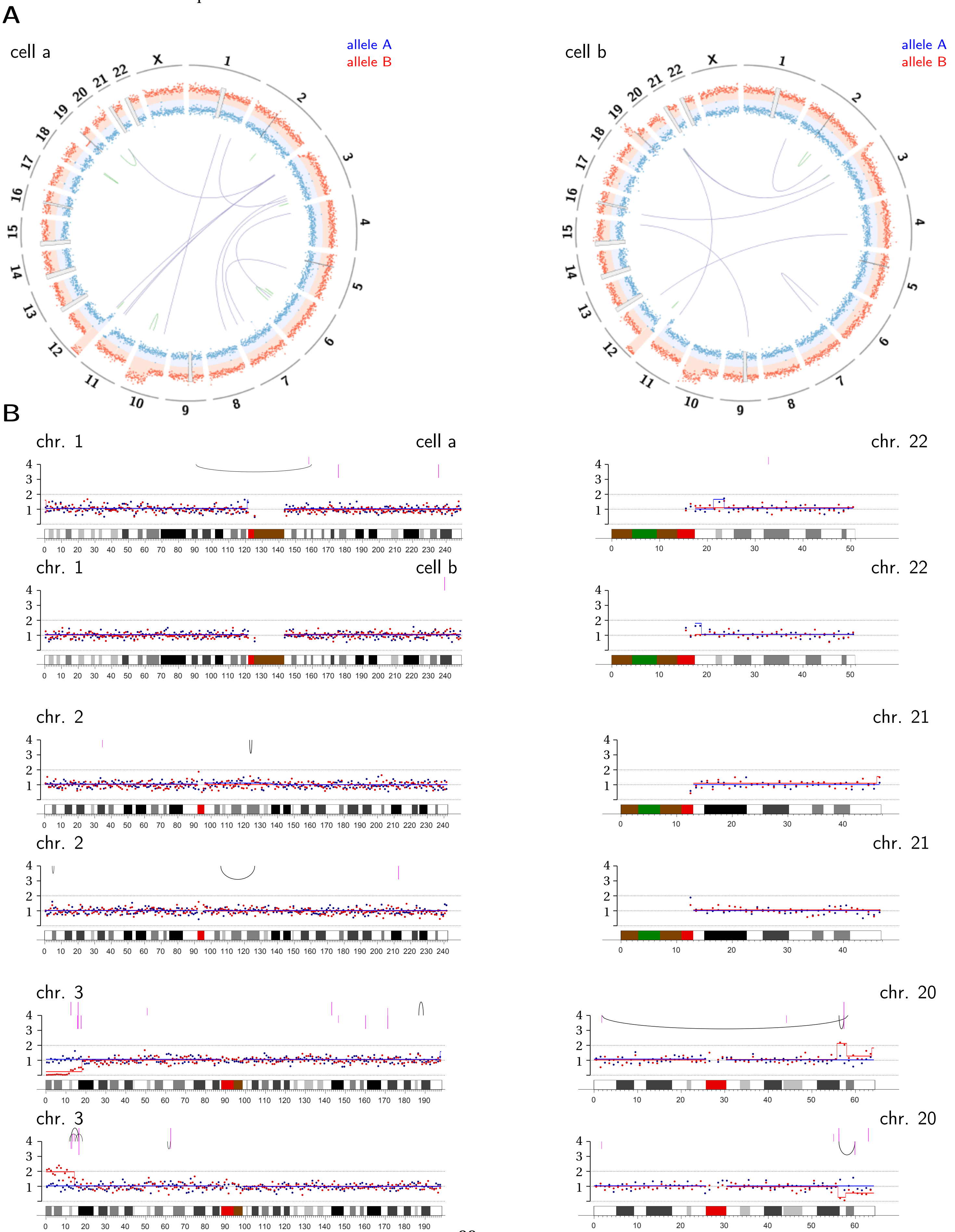

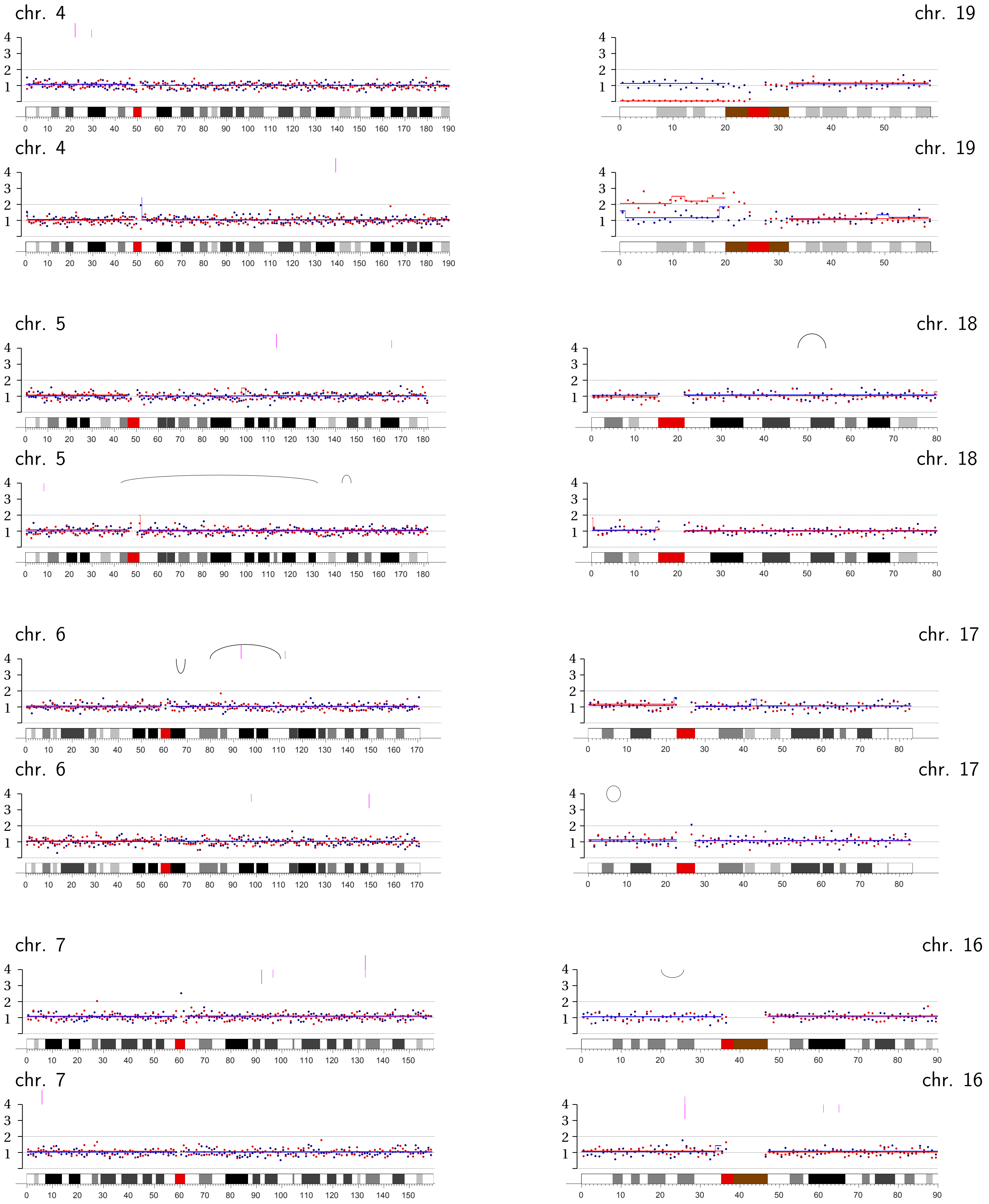

chr. 4
chr. 19
chr. 4
chr. 19
chr. 5
chr. 18
chr. 5
chr. 18
chr. 6
chr. 17
chr. 6
chr. 17
chr. 7
chr. 16
chr. 7
chr. 16

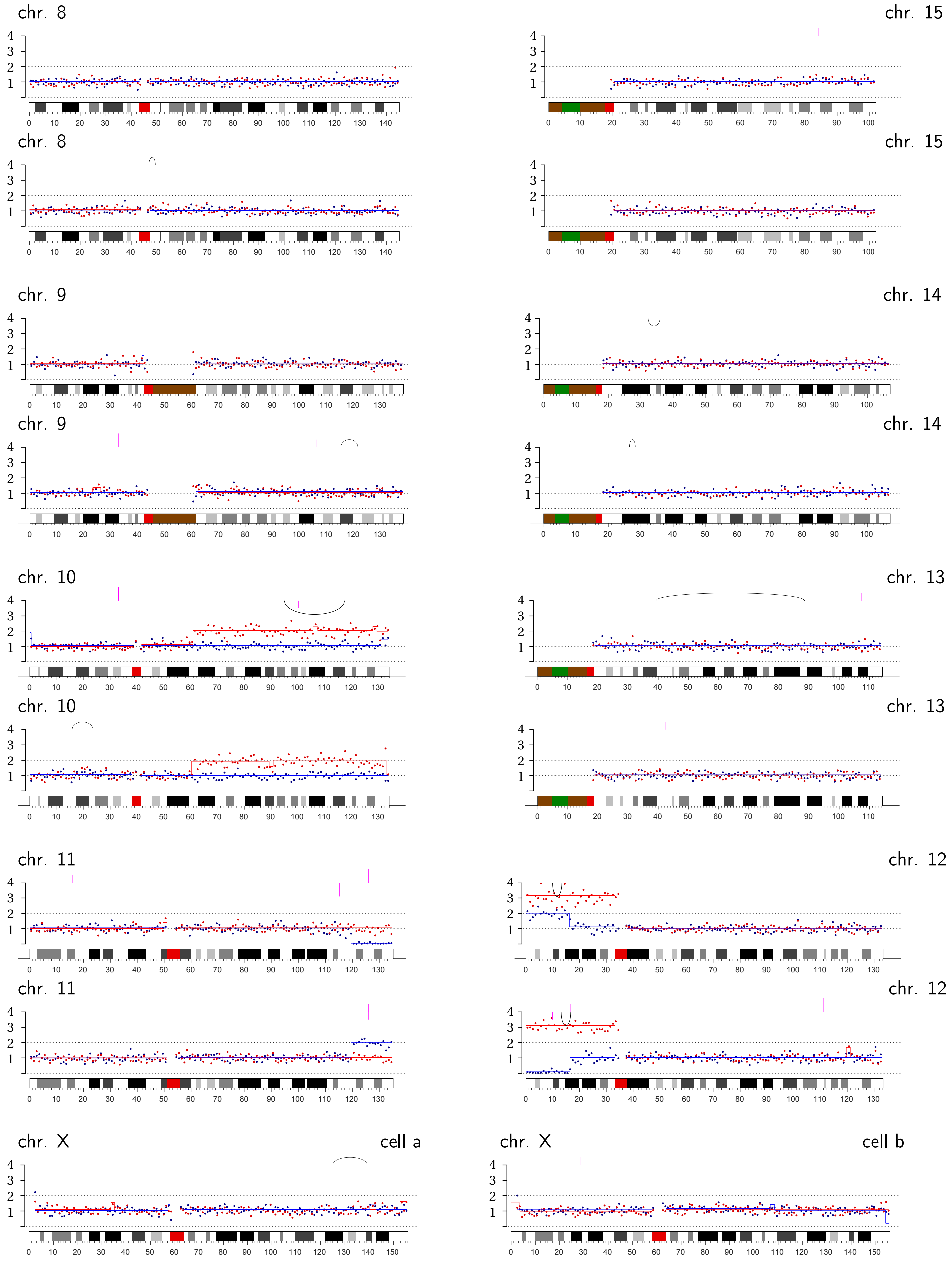
chr. 8
chr. 15
chr. 8
chr. 15
chr. 9
chr. 14
chr. 9
chr. 14
chr. 10
chr. 13
chr. 10
chr. 13
chr. 11
chr. 12
chr. 11
chr. 12
chr. X
cell a
chr. X
cell b

**Bridge Pair T-7** | Transient TRF2-DN expression: Chromosomes 12B, p-arm; 17B, q-ter.

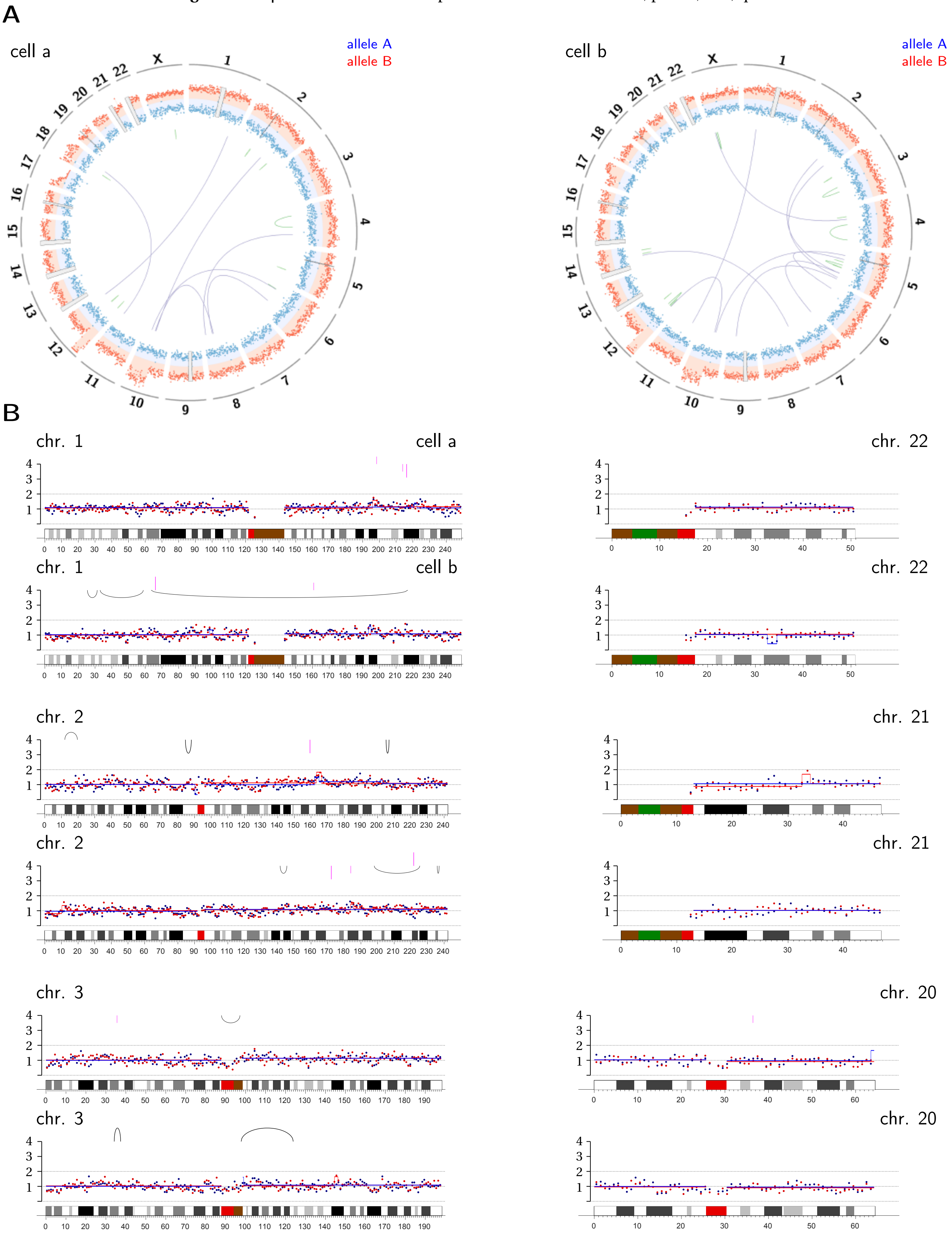

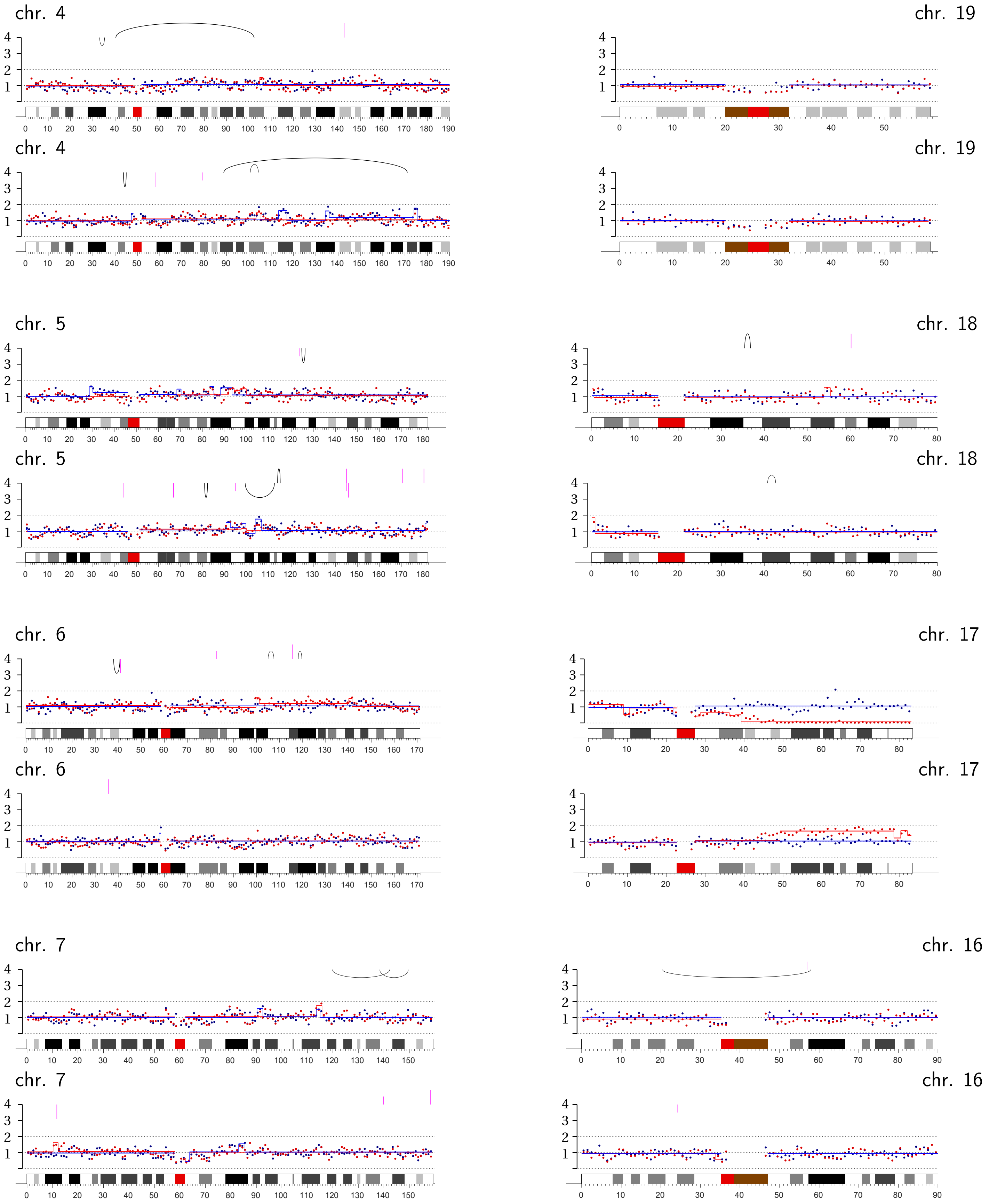
chr. 4
chr. 4
chr. 19
chr. 19
chr. 5
chr. 5
chr. 18
chr. 18
chr. 6
chr. 6
chr. 17
chr. 17
chr. 7
chr. 7
chr. 16
chr. 16

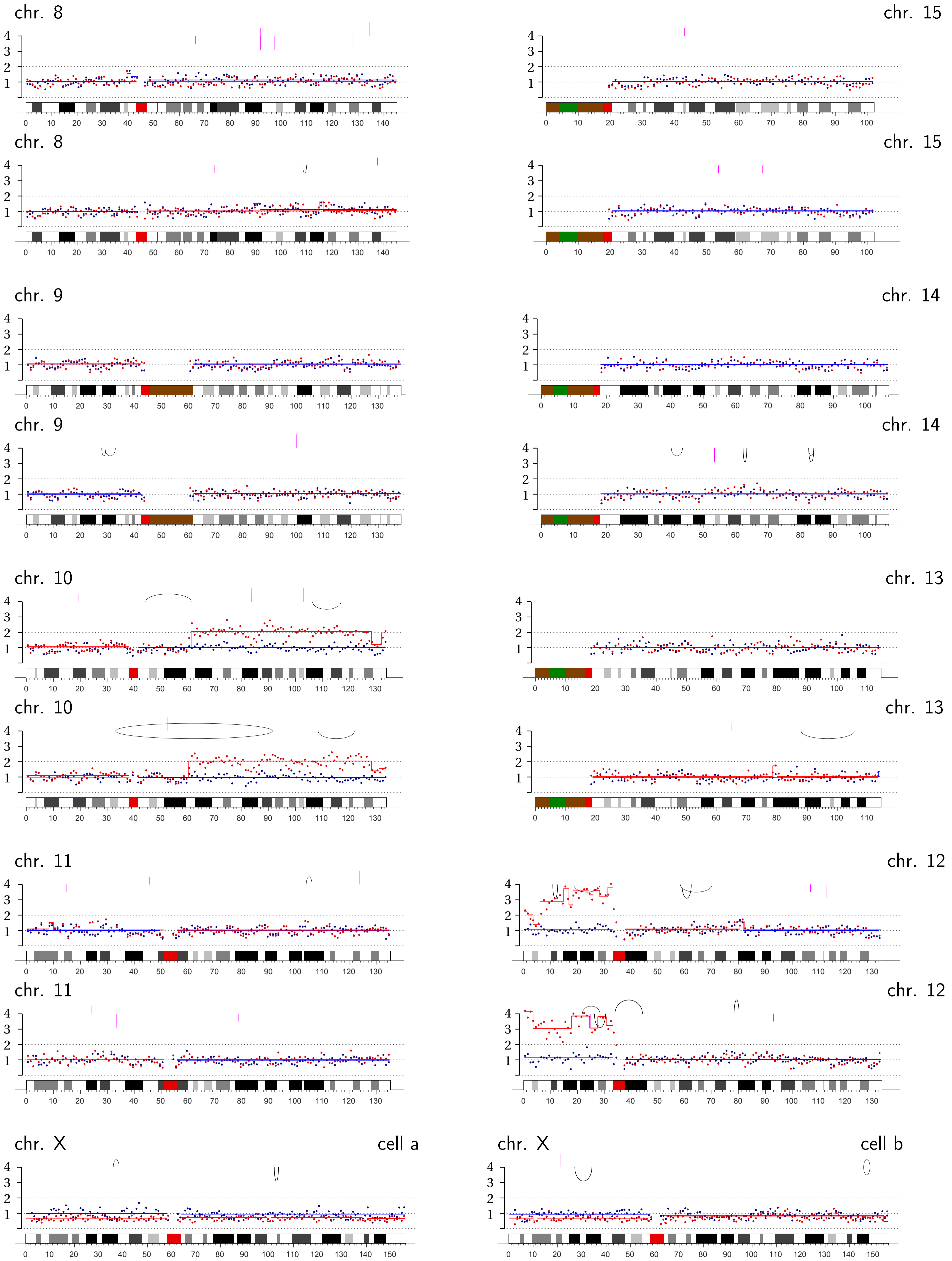
chr. 8
chr. 15
chr. 8
chr. 15
chr. 9
chr. 14
chr. 9
chr. 14
chr. 10
chr. 13
chr. 10
chr. 13
chr. 11
chr. 12
chr. 11
chr. 12
chr. X
cell a
chr. X
cell b

**Bridge Pair T-8 |** Transient TRF2-DN expression: Chromosomes 3A, p-ter.; 16B, q-arm.

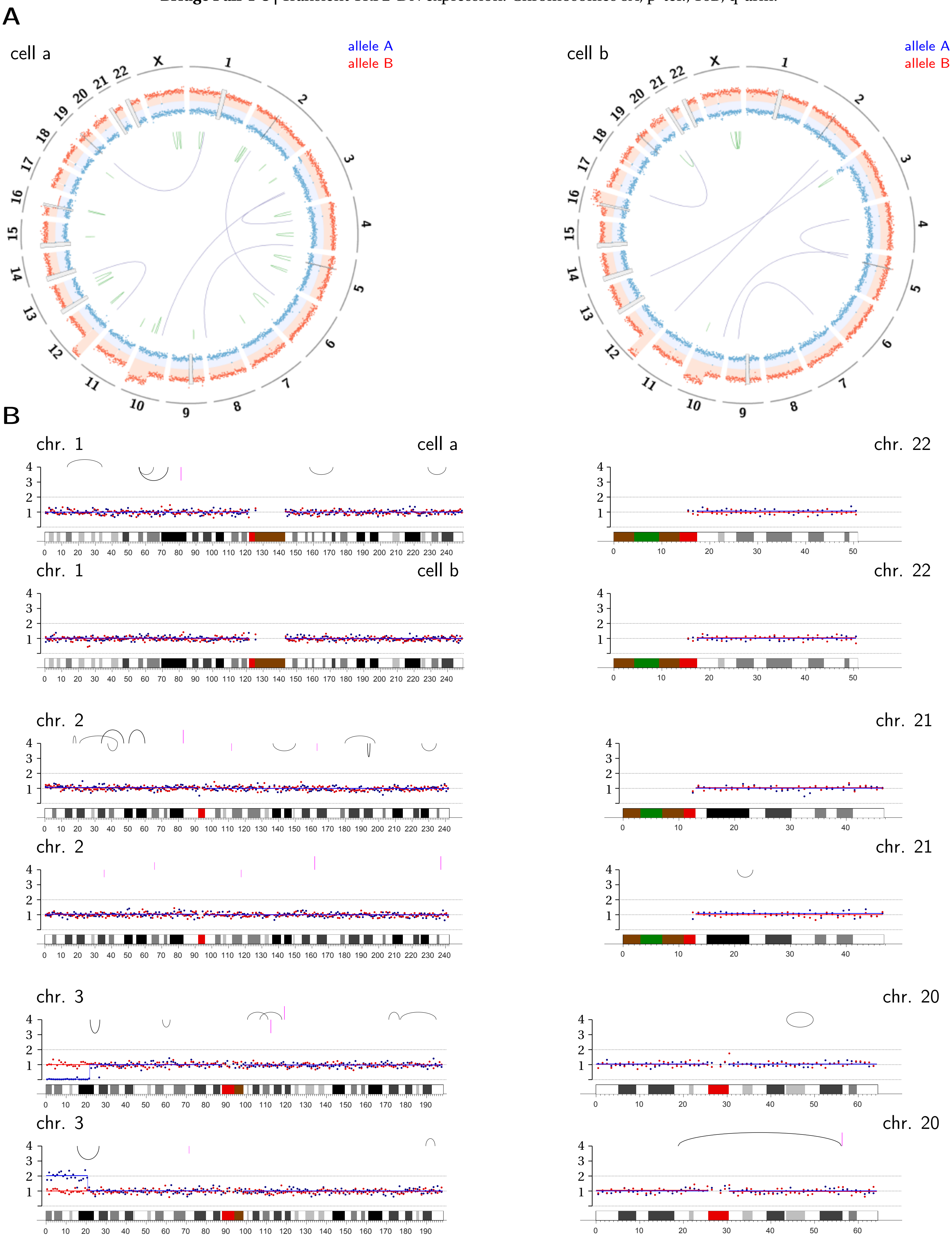

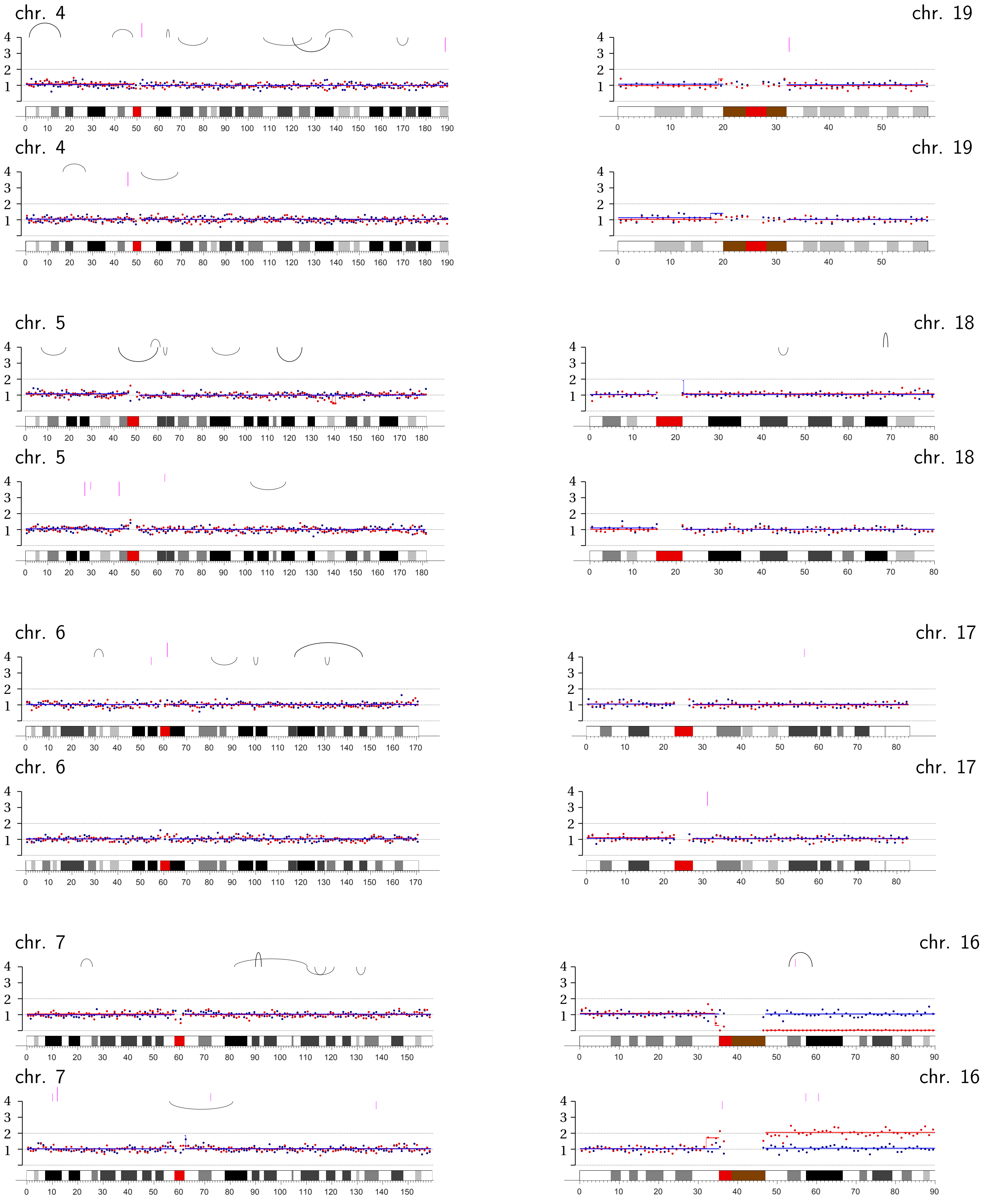
chr. 4
chr. 19
chr. 4
chr. 19
chr. 5
chr. 18
chr. 5
chr. 18
chr. 6
chr. 17
chr. 6
chr. 17
chr. 7
chr. 16
chr. 7
chr. 16

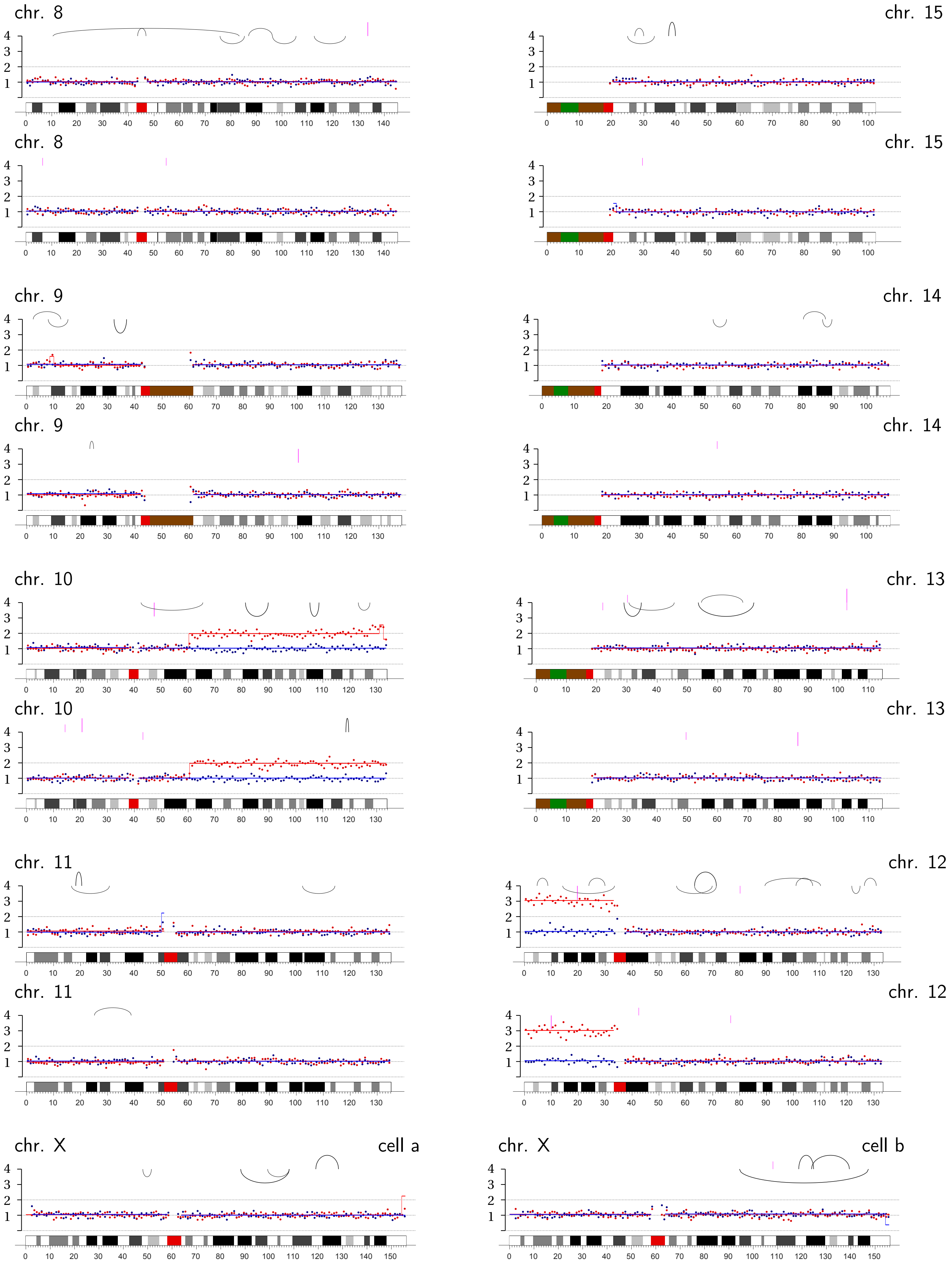
chr. 8
chr. 15
chr. 8
chr. 15
chr. 9
chr. 14
chr. 9
chr. 14
chr. 10
chr. 13
chr. 10
chr. 13
chr. 11
chr. 12
chr. 11
chr. 12
chr. X
cell a
chr. X
cell b

**Bridge Pair 4-1** | CRISPR: Chromosome 4A q-ter.

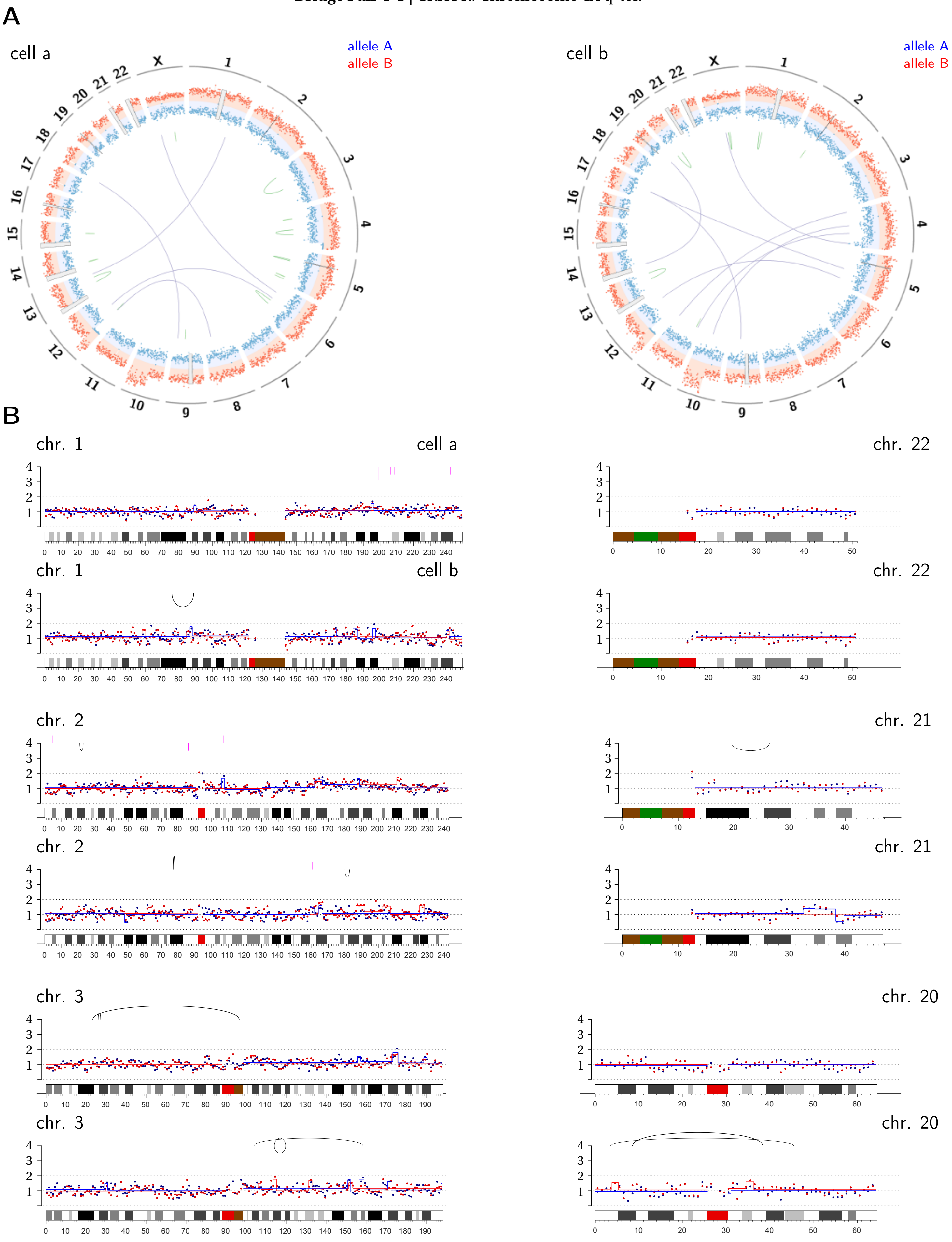

chr. 4

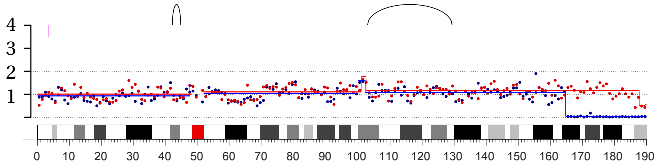

chr. 4

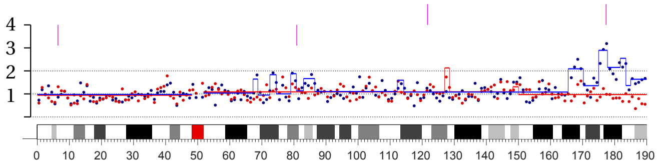

chr. 5

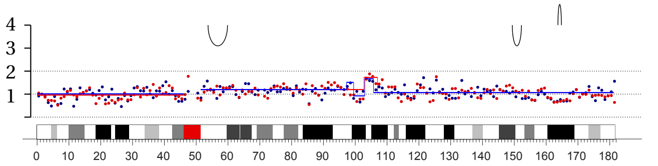

chr. 5

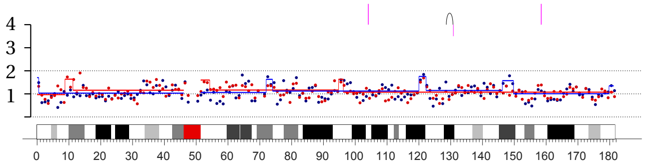

chr. 6

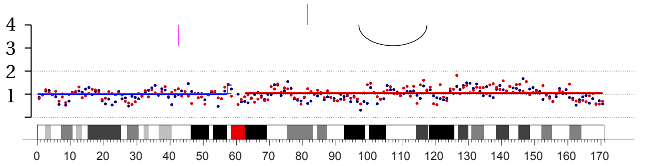

chr. 6

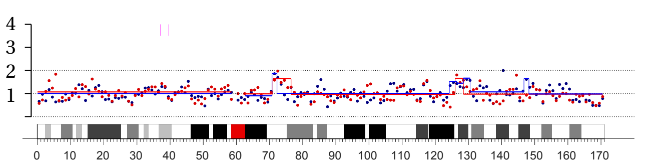

chr. 7

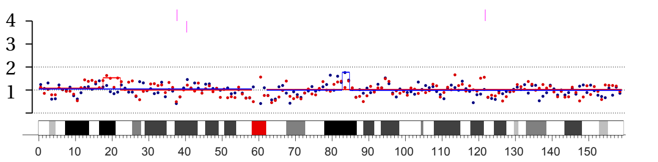

chr. 7

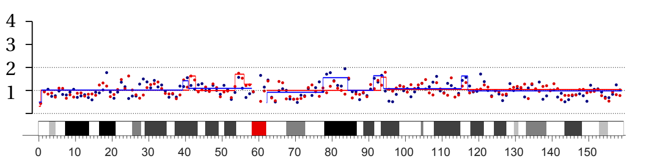

chr. 19

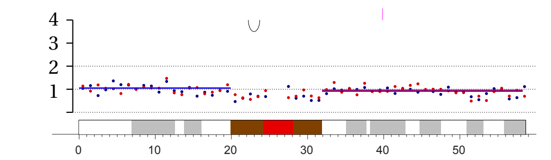

chr. 19

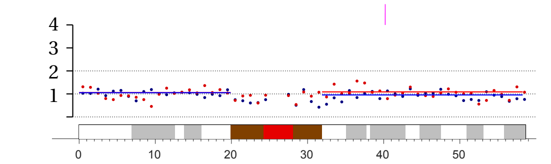

chr. 18

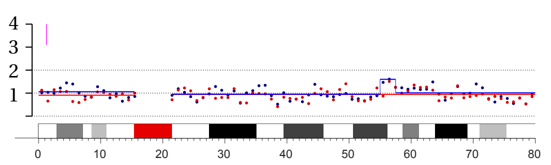

chr. 18

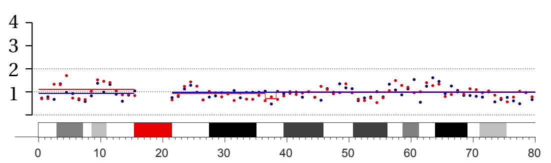

chr. 17

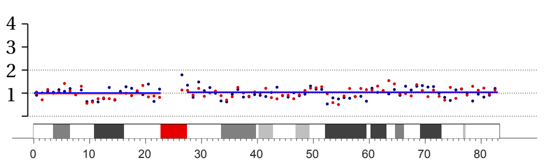

chr. 17

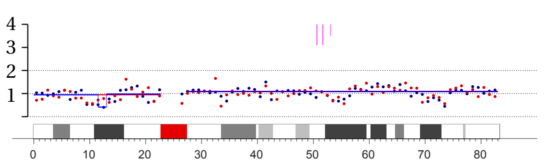

chr. 16

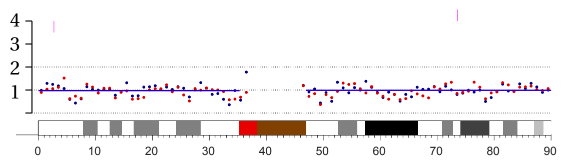

chr. 16

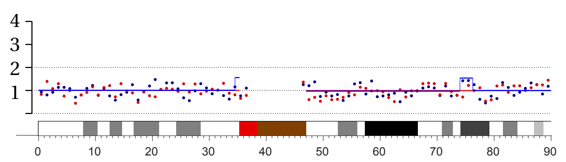

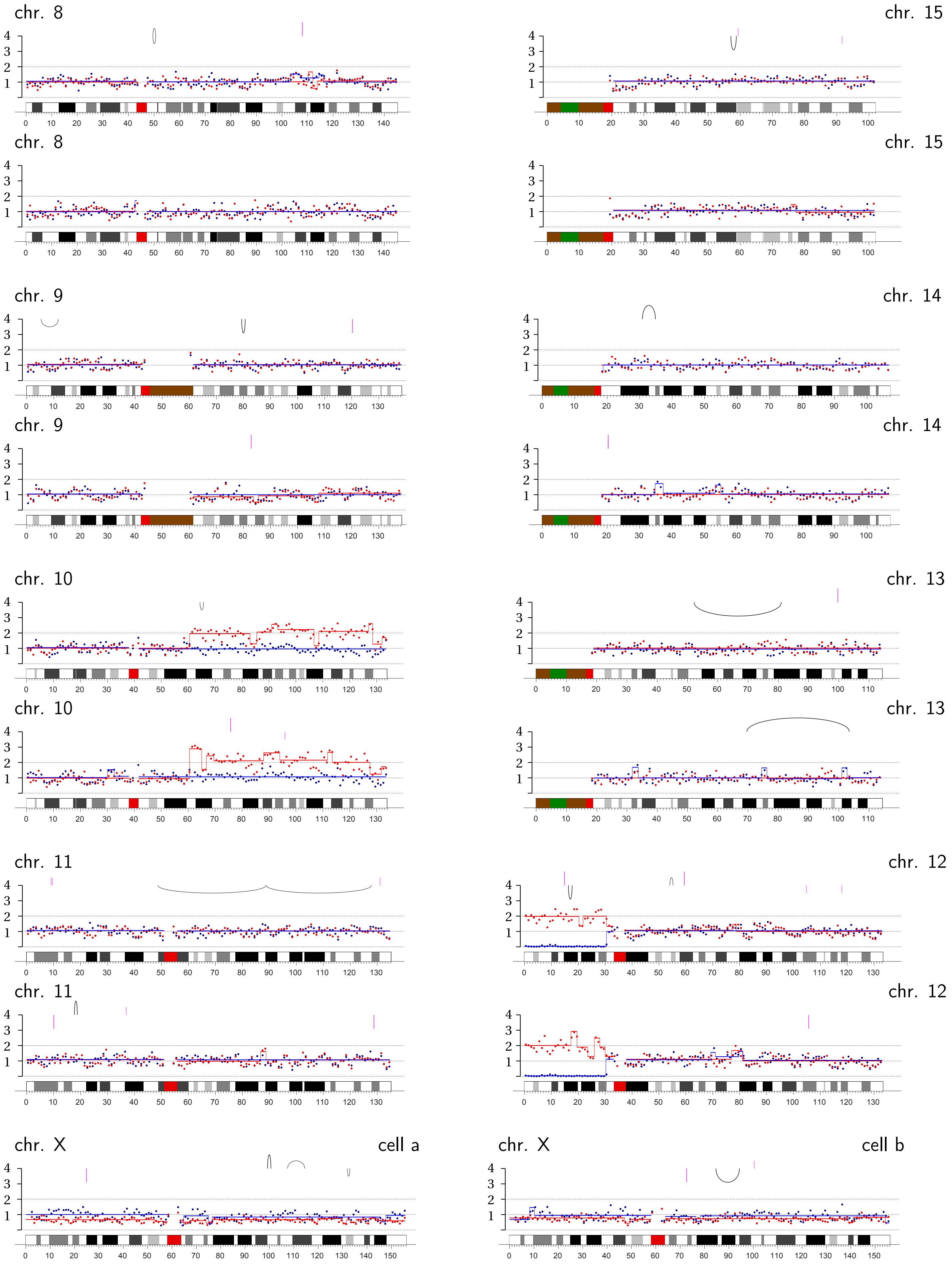
chr. 8
chr. 15
chr. 8
chr. 15
chr. 9
chr. 14
chr. 9
chr. 14
chr. 10
chr. 13
chr. 10
chr. 13
chr. 11
chr. 12
chr. 11
chr. 12
chr. X
cell a
chr. X
cell b

**Bridge Pair 4-2** | CRISPR: Chromosome 4A q-ter.

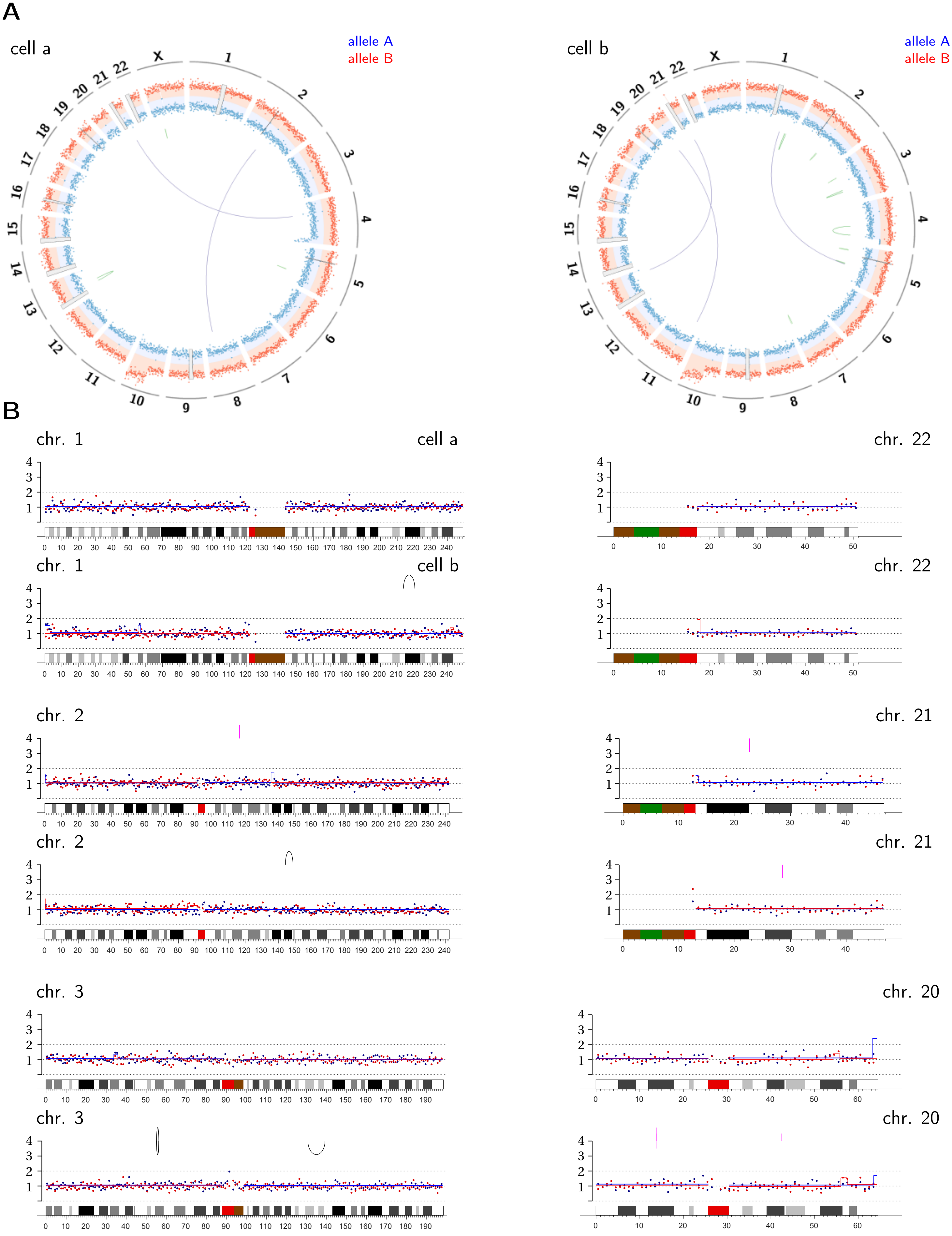

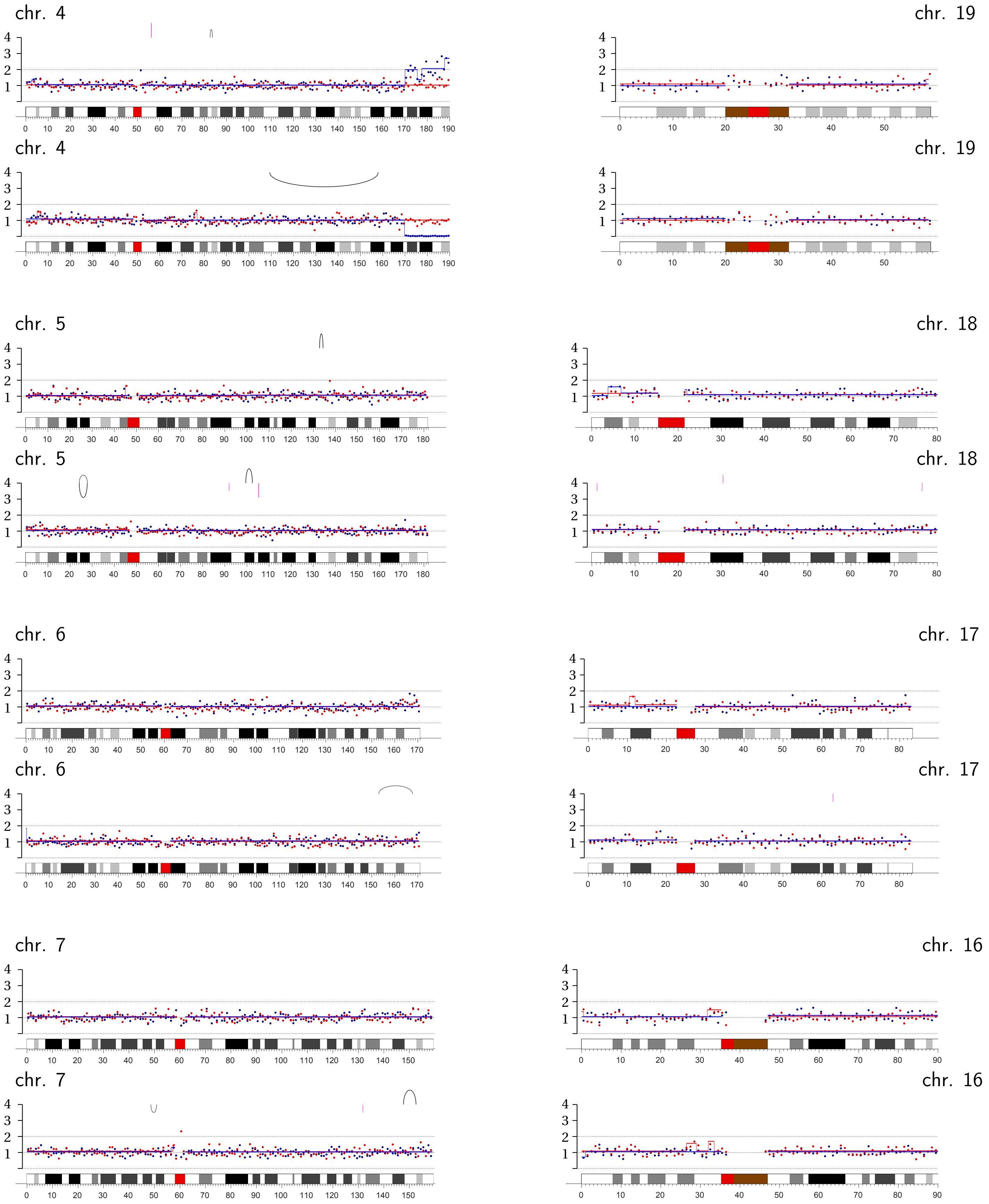
chr. 4
chr. 19
chr. 4
chr. 19
chr. 5
chr. 18
chr. 5
chr. 18
chr. 6
chr. 17
chr. 6
chr. 17
chr. 7
chr. 16
chr. 7
chr. 16

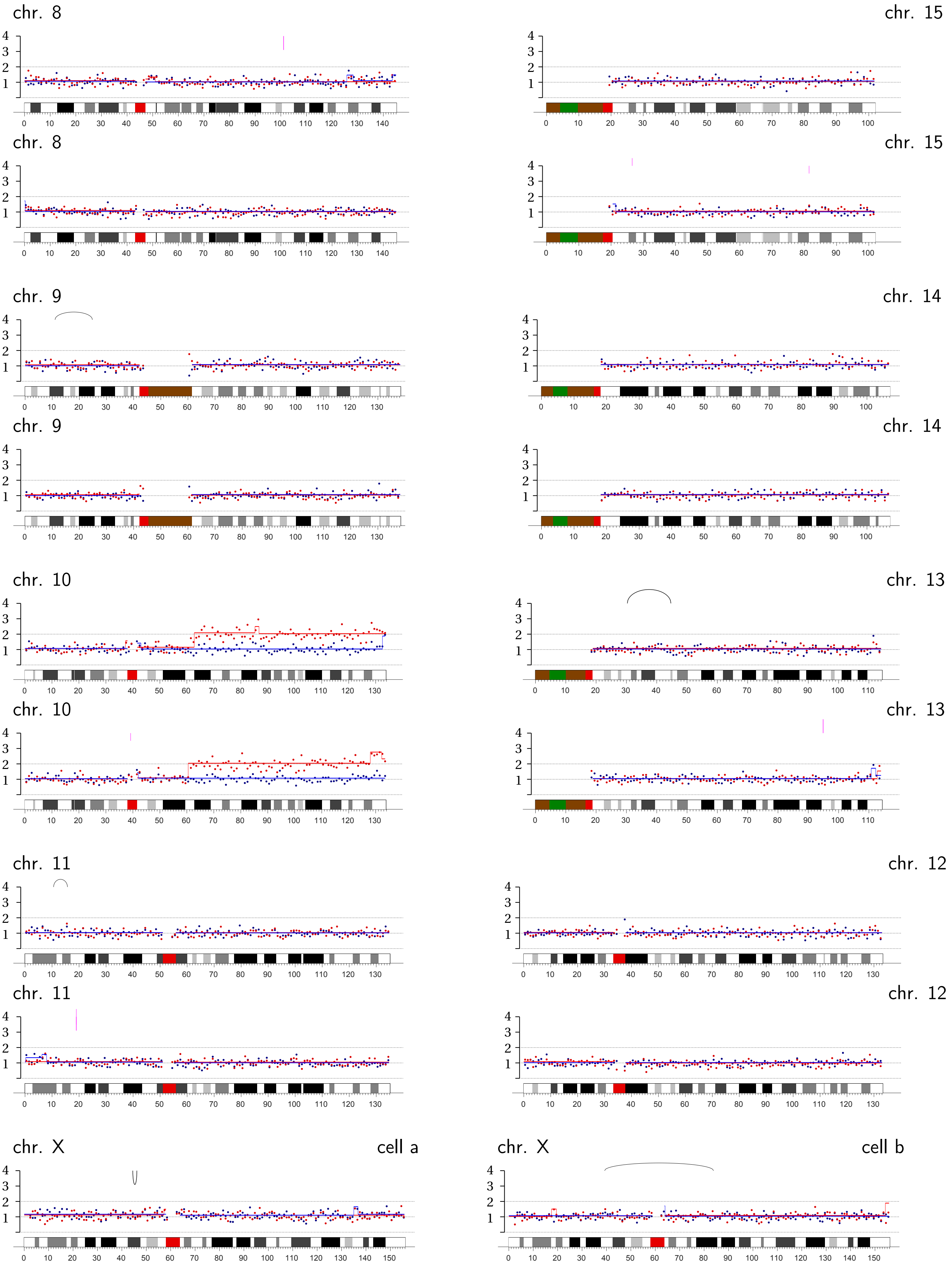
chr. 8
chr. 15
chr. 8
chr. 15
chr. 9
chr. 14
chr. 9
chr. 14
chr. 10
chr. 13
chr. 10
chr. 13
chr. 11
chr. 12
chr. 11
chr. 12
chr. X
cell a
chr. X
cell b

**Bridge Pair 4-3** | CRISPR: Chromosome 4A p-arm internal.

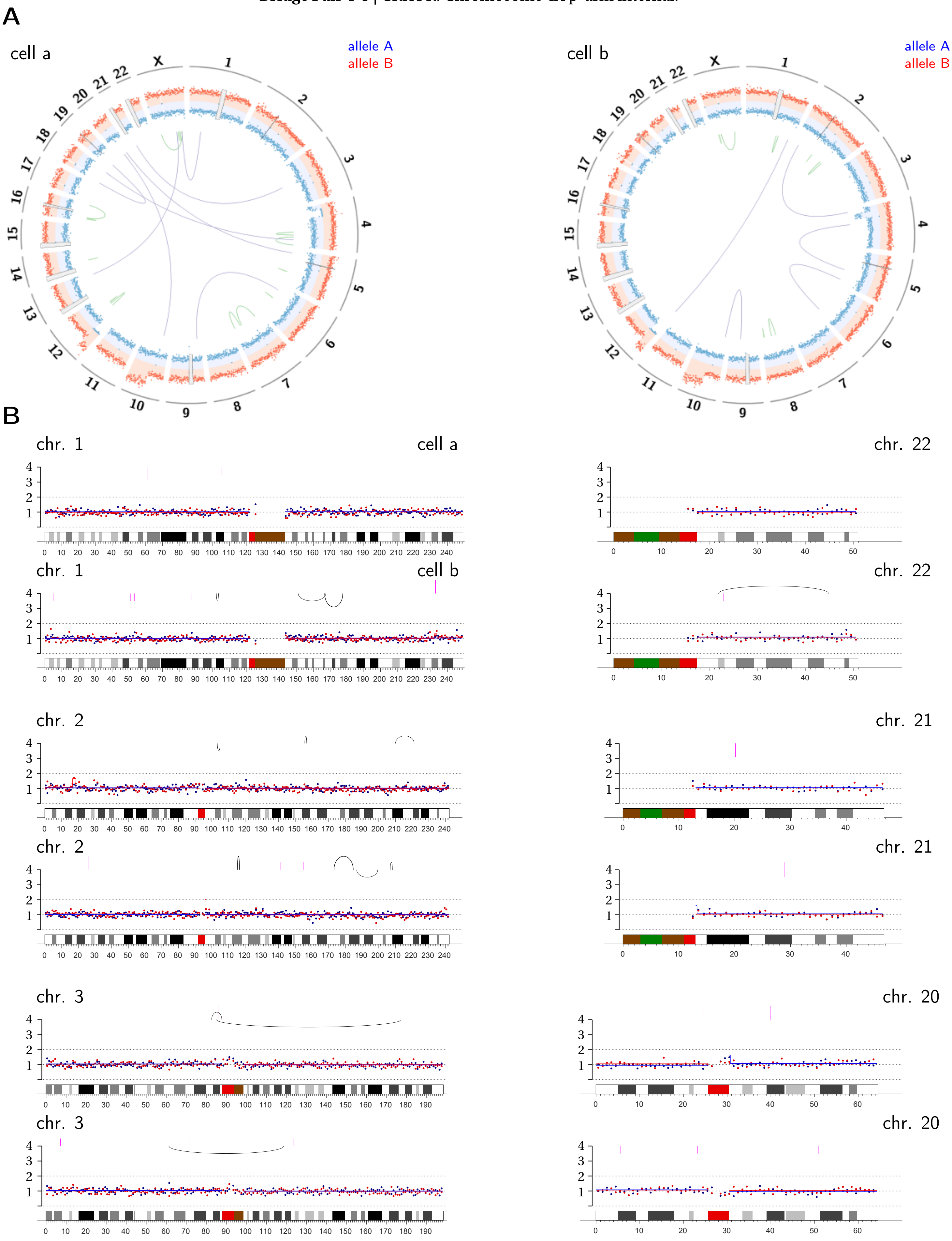

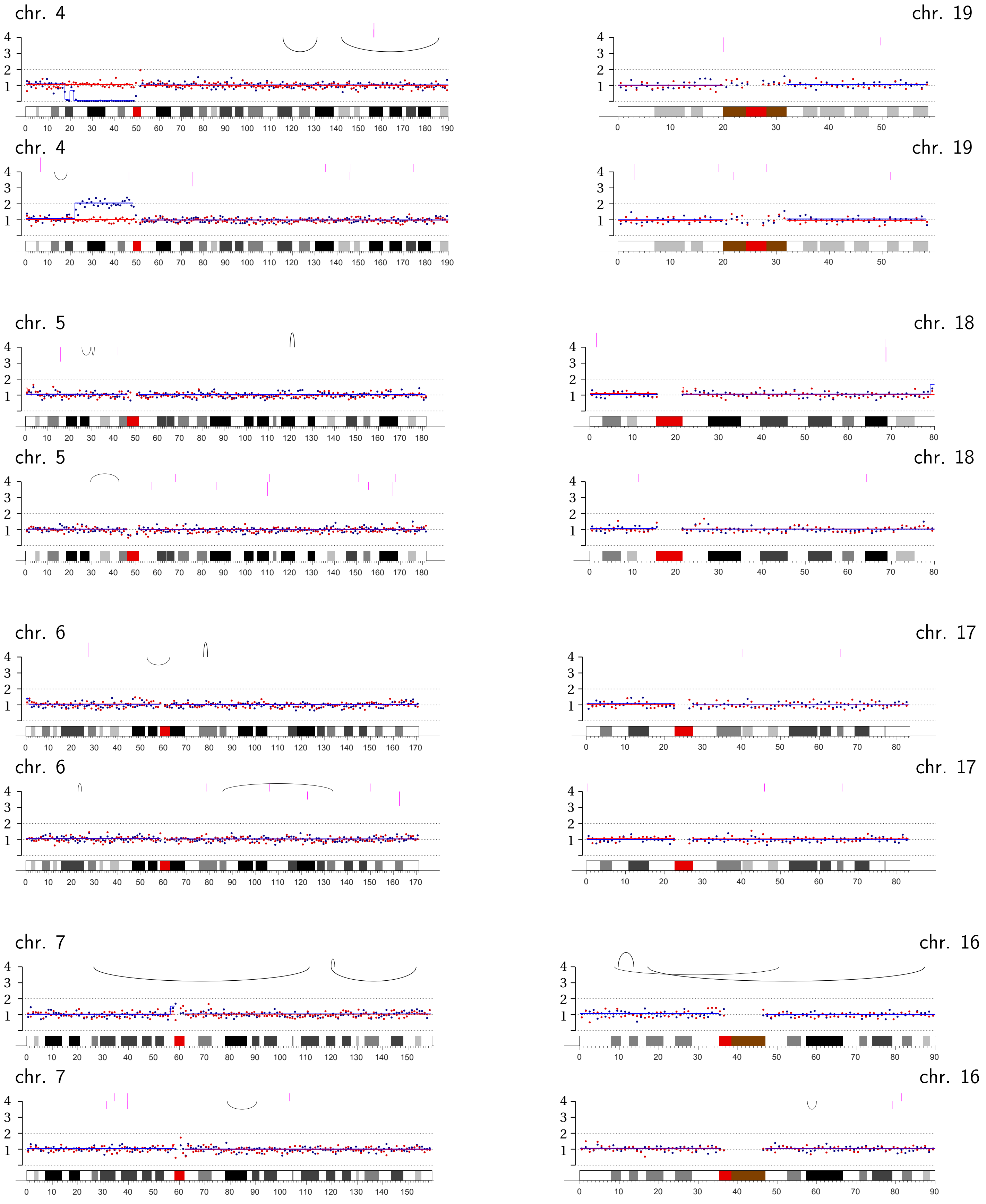
chr. 4
chr. 19
chr. 4
chr. 19
chr. 5
chr. 18
chr. 5
chr. 18
chr. 6
chr. 17
chr. 6
chr. 17
chr. 7
chr. 16
chr. 7
chr. 16

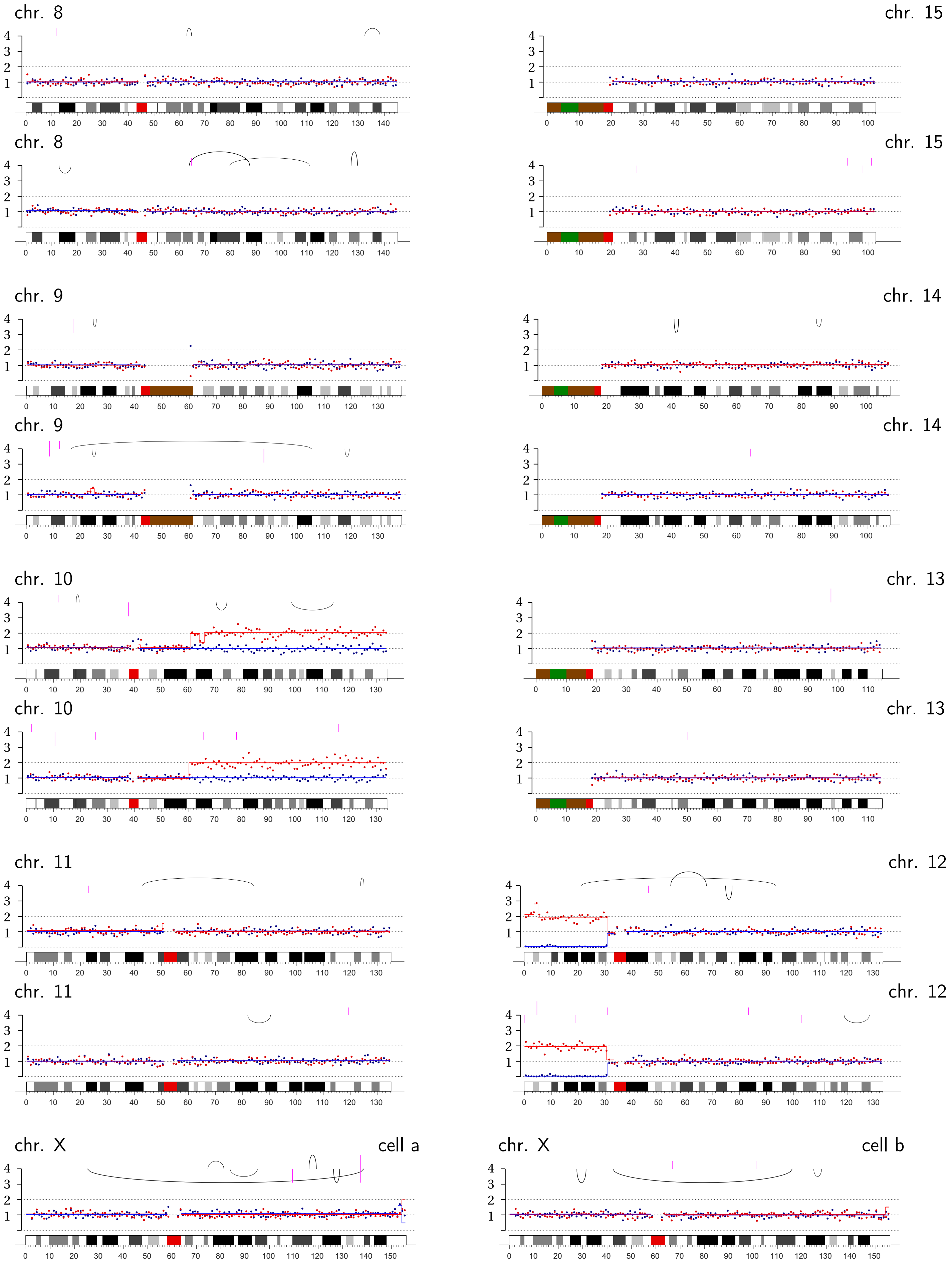
chr. 8
chr. 15
chr. 8
chr. 15
chr. 9
chr. 14
chr. 9
chr. 14
chr. 10
chr. 13
chr. 10
chr. 13
chr. 11
chr. 12
chr. 11
chr. 12
chr. X
cell a
chr. X
cell b

**Bridge Pair 4-4** | CRISPR: Chromosome 4A q-ter.

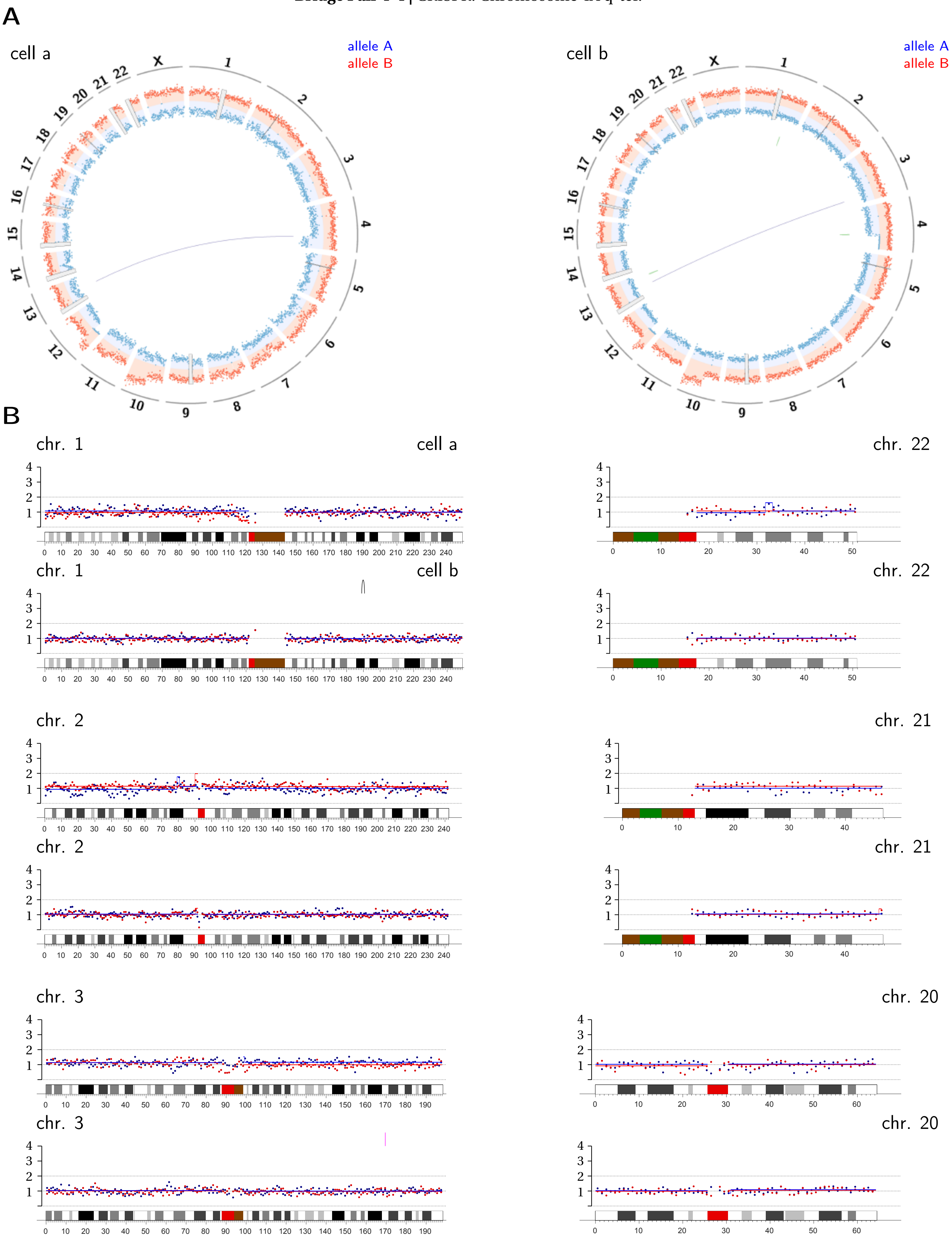

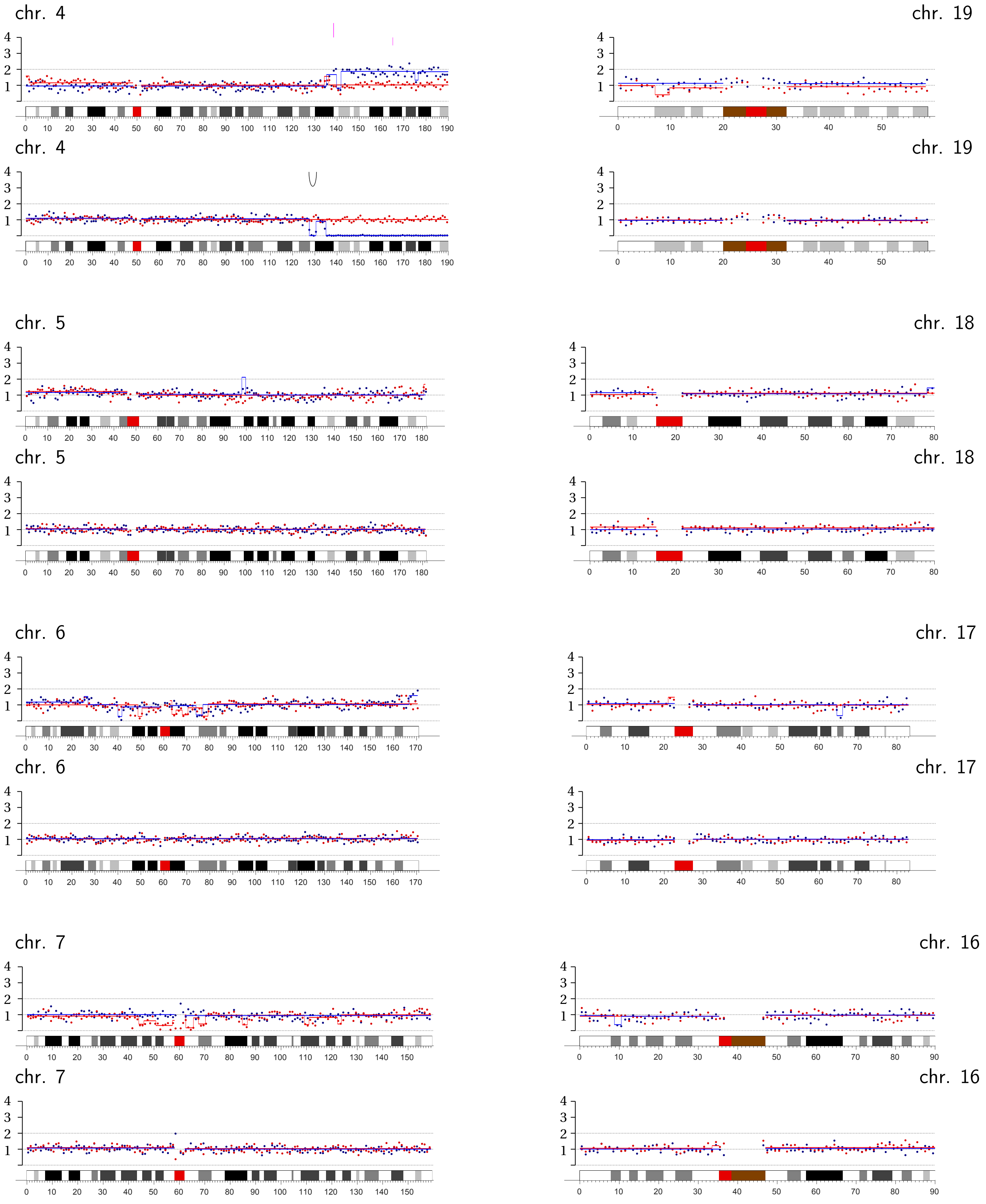
chr. 4
chr. 19
chr. 4
chr. 19
chr. 5
chr. 18
chr. 5
chr. 18
chr. 6
chr. 17
chr. 6
chr. 17
chr. 7
chr. 16
chr. 7
chr. 16

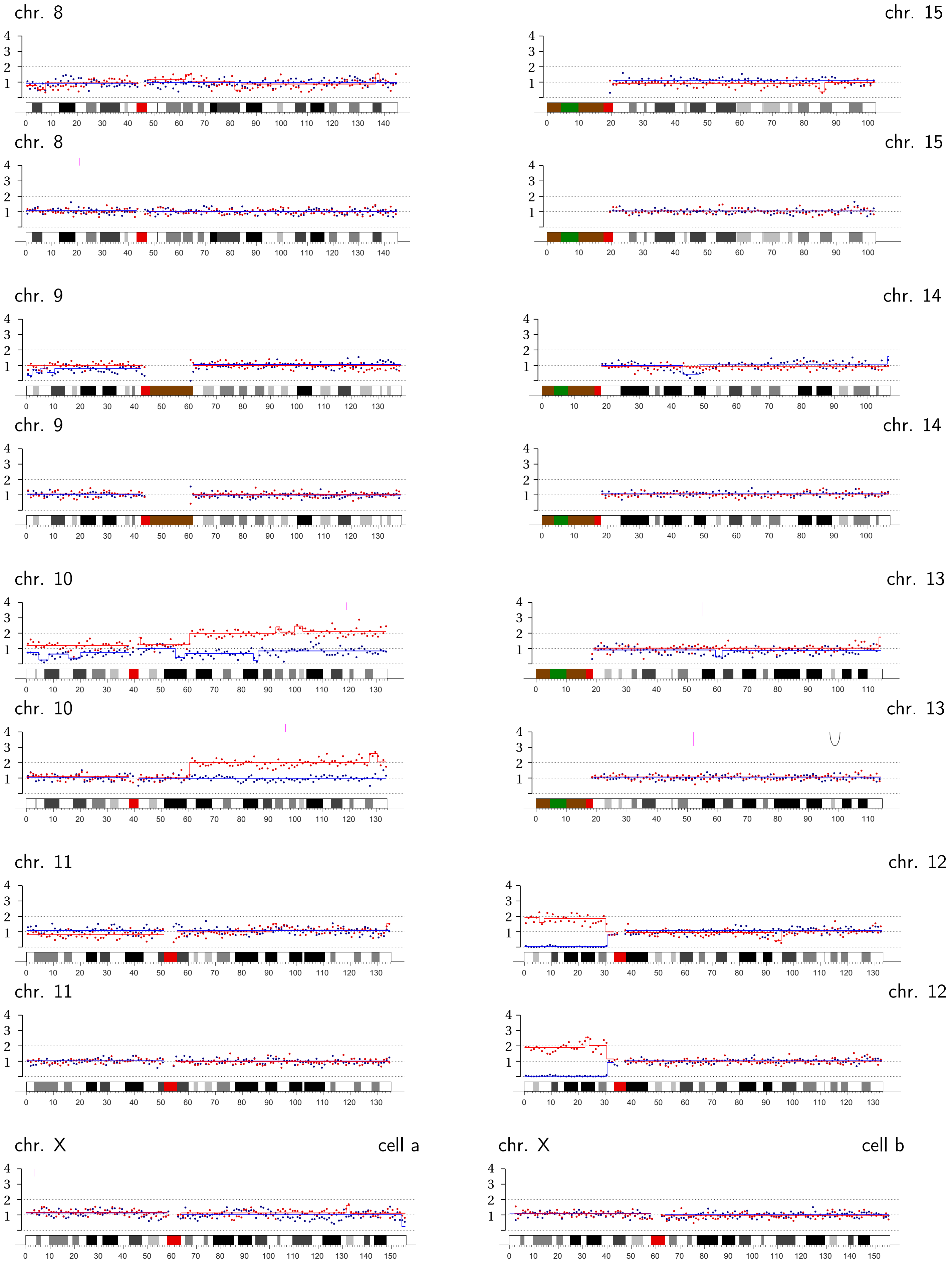
chr. 8
chr. 15
chr. 8
chr. 15
chr. 9
chr. 14
chr. 9
chr. 14
chr. 10
chr. 13
chr. 10
chr. 13
chr. 11
chr. 12
chr. 11
chr. 12
chr. X
cell a
chr. X
cell b

**Bridge Pair 4-5** | CRISPR: Chromosome 4A q-ter.

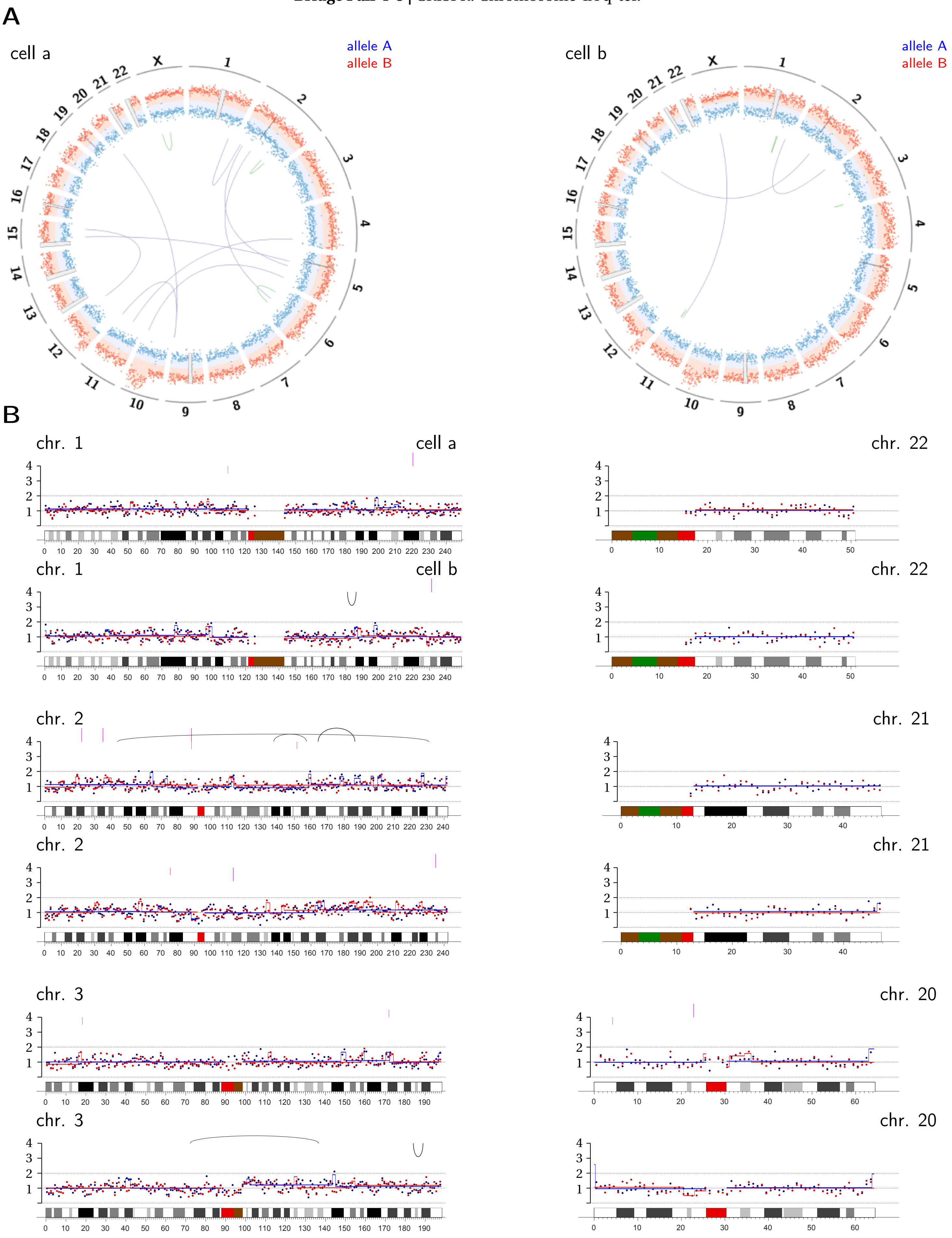

chr. 4

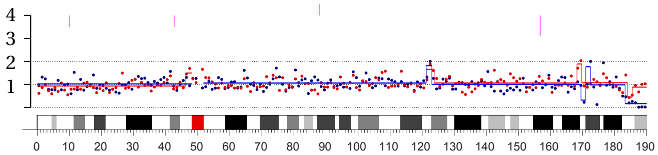


chr. 4

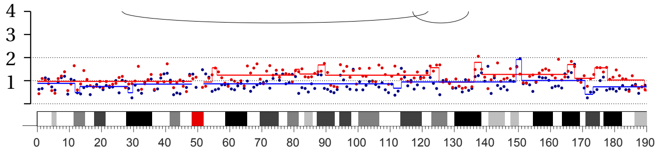


chr. 19

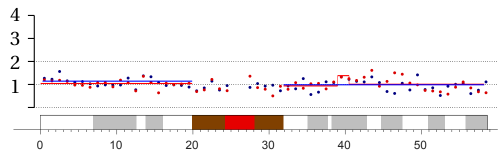


chr. 19

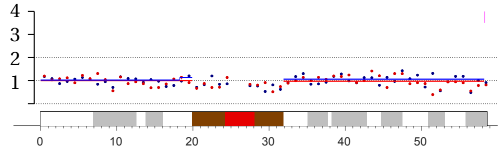


chr. 5

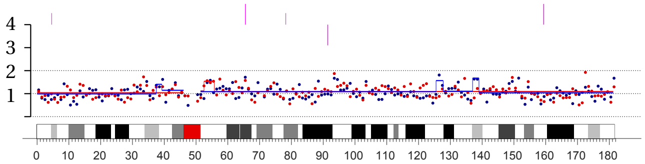


chr. 5

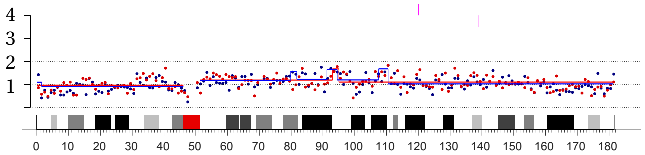


chr. 18

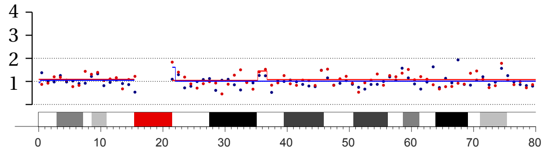


chr. 18

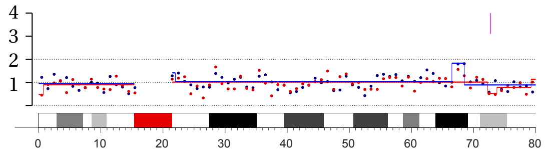


chr. 6

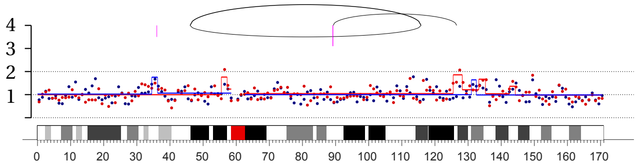


chr. 6

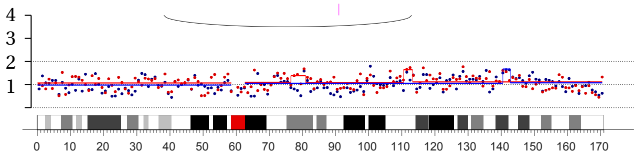


chr. 17

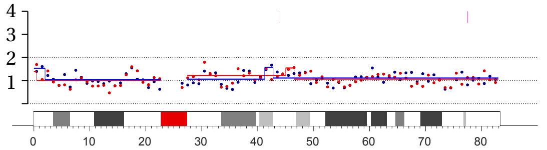


chr. 17

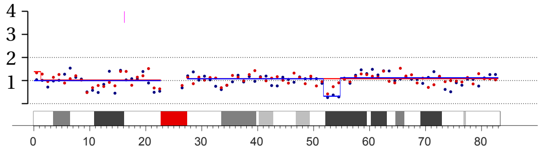


chr. 7

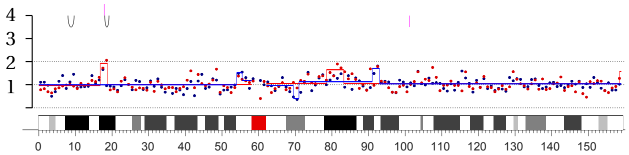


chr. 7

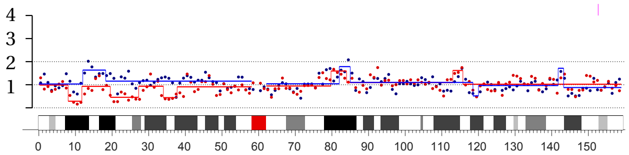


chr. 16

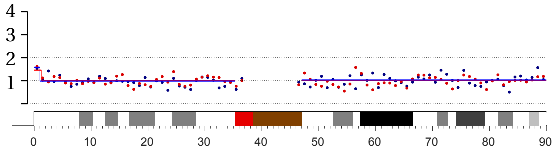


chr. 16

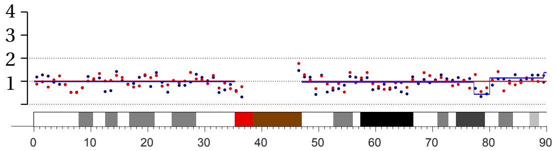

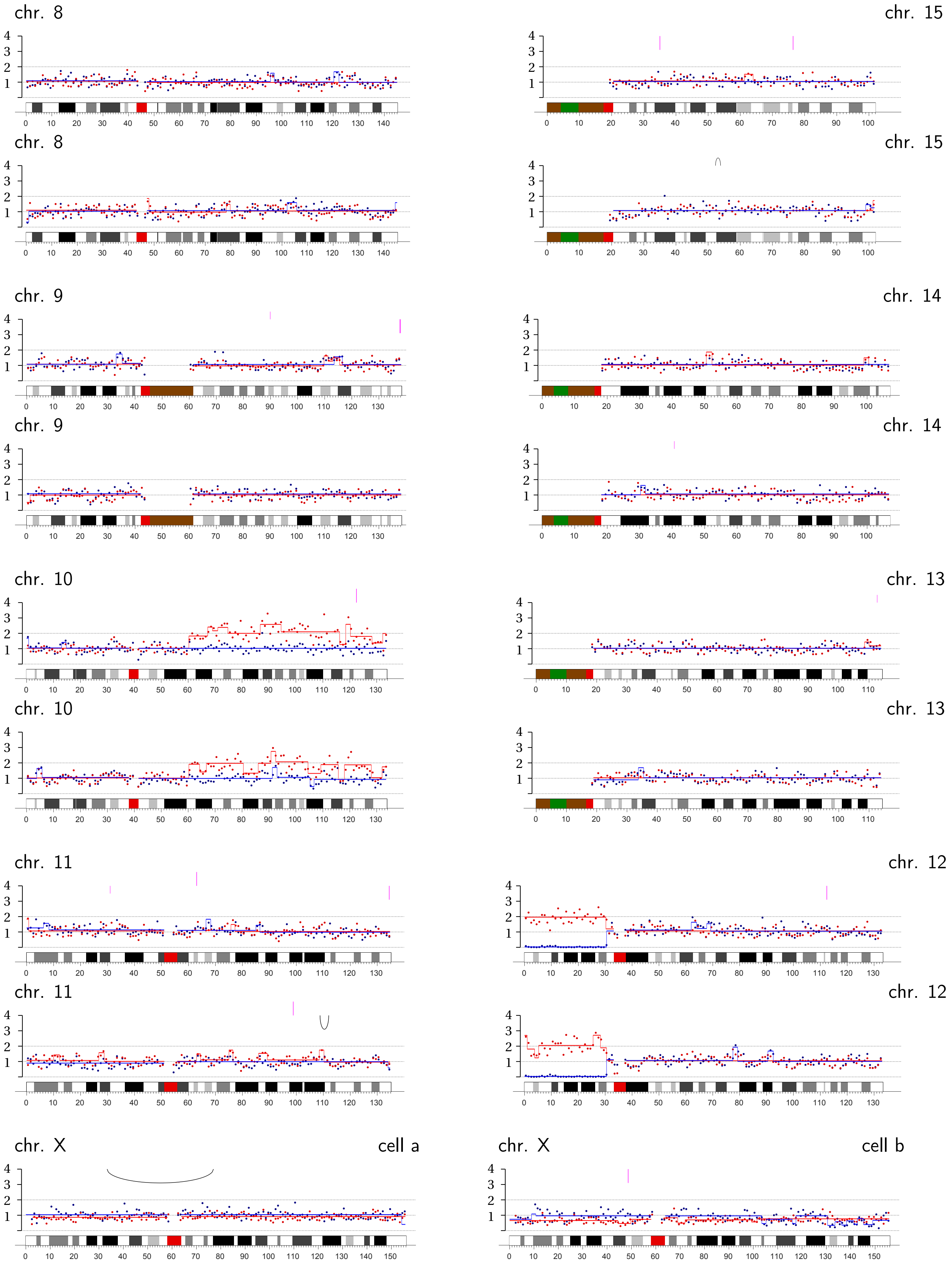
chr. 8
chr. 15
chr. 8
chr. 15
chr. 9
chr. 14
chr. 9
chr. 14
chr. 10
chr. 13
chr. 10
chr. 13
chr. 11
chr. 12
chr. 11
chr. 12
chr. X
cell a
chr. X
cell b

**Bridge Pair 4-6** | CRISPR: Chromosomes 4A (p- and q-ter.) and 4B (p-ter).

A

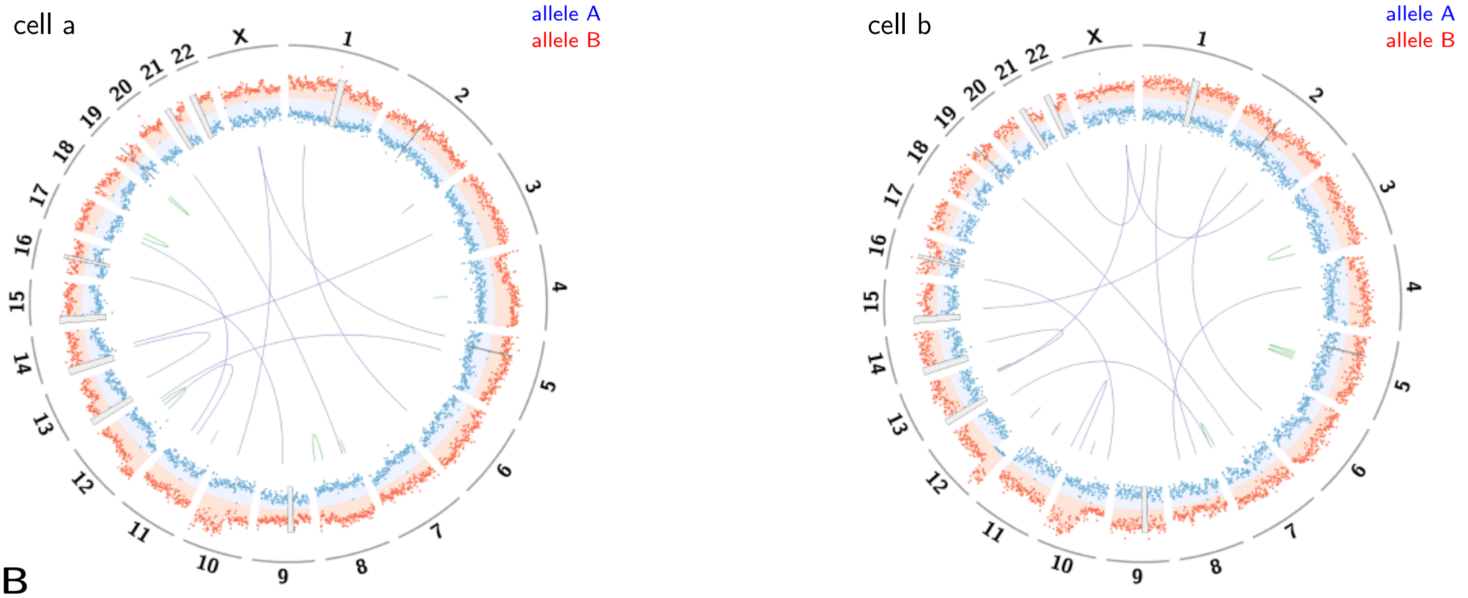


B

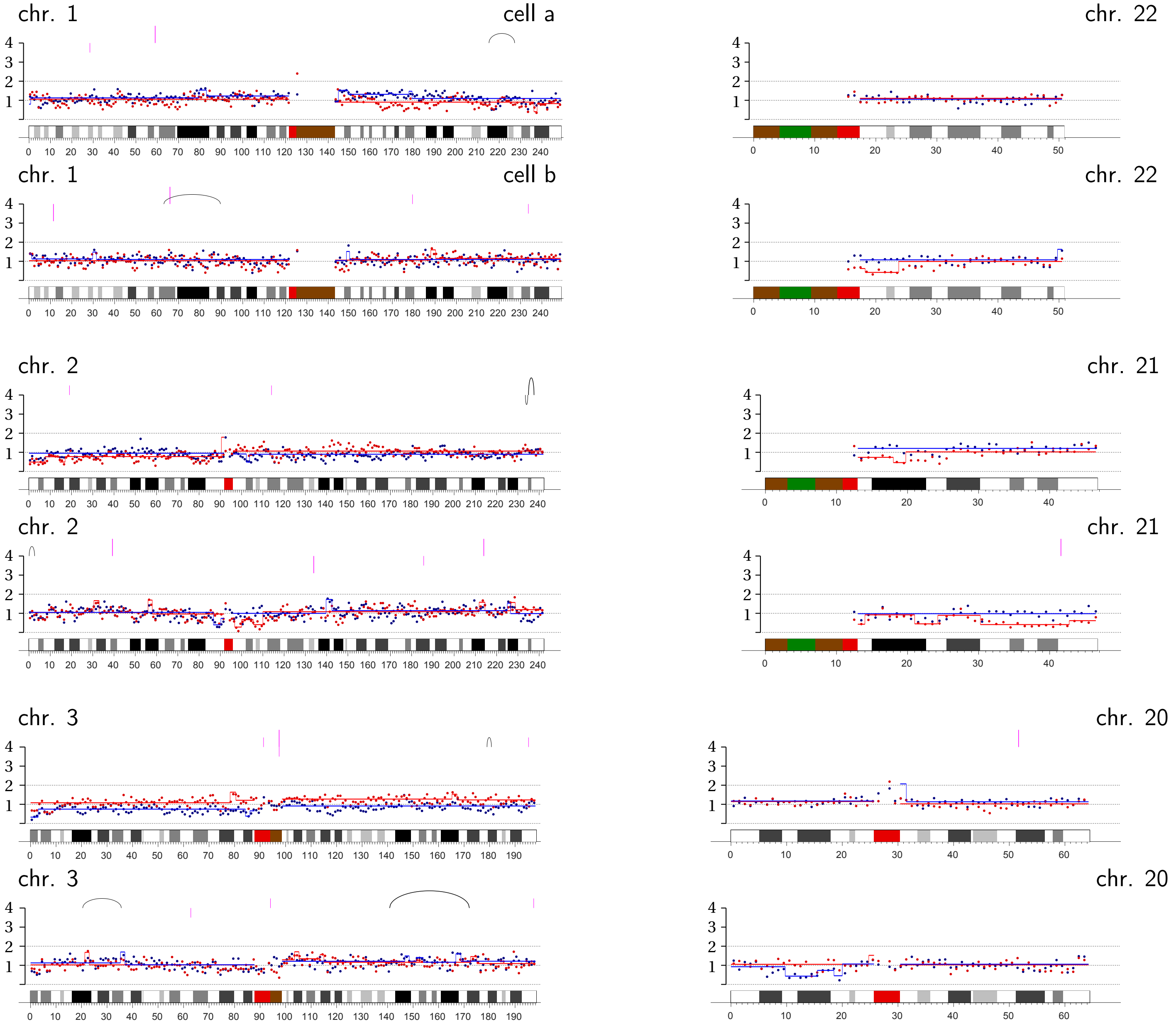

chr. 4

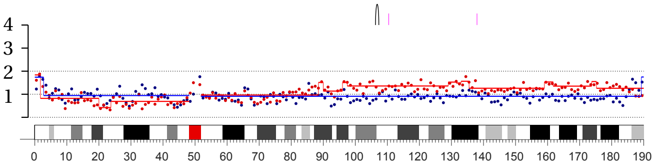


chr. 4

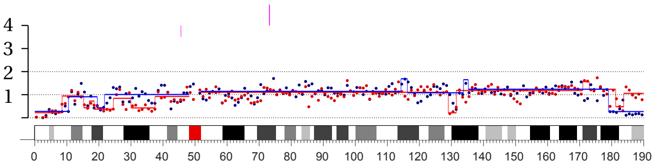


chr. 19

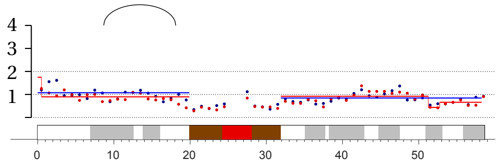


chr. 19

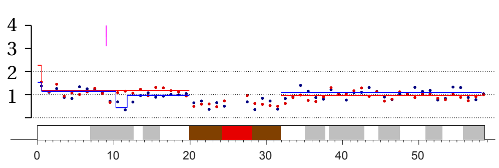


chr. 5

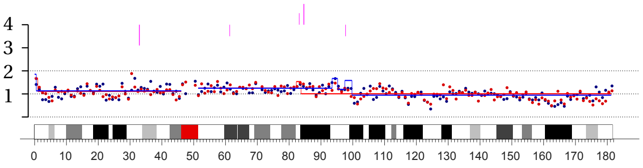


chr. 5

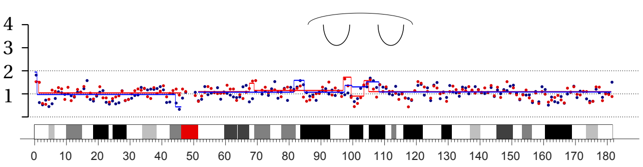


chr. 18

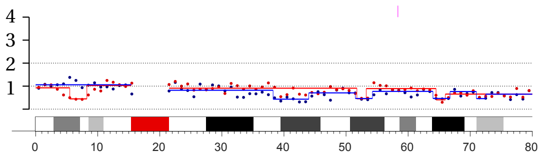


chr. 18

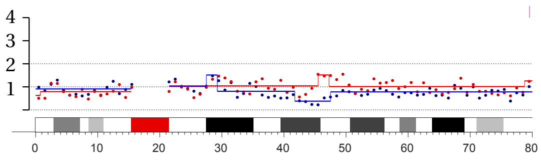


chr. 6

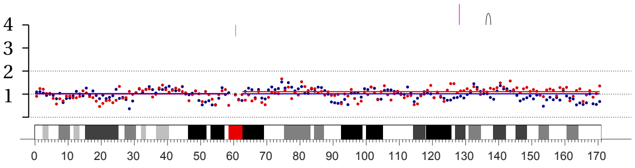


chr. 6

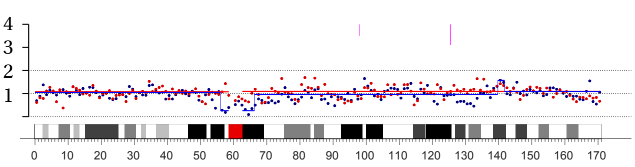


chr. 17

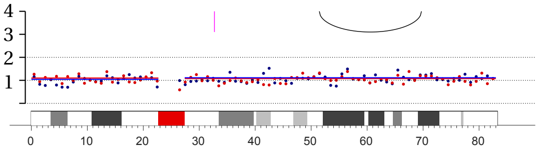


chr. 17

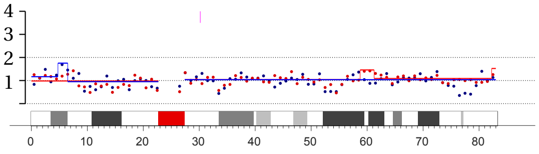


chr. 7

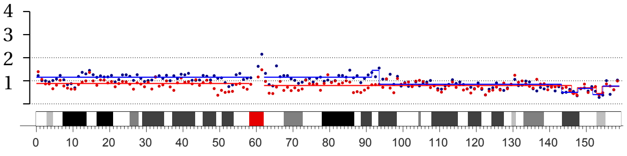


chr. 7

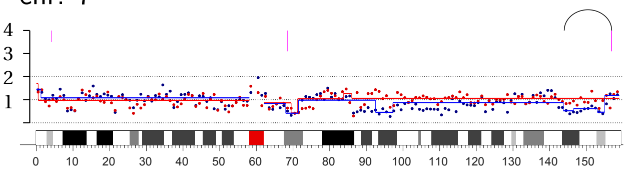


chr. 16

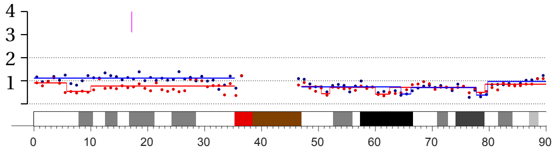


chr. 16

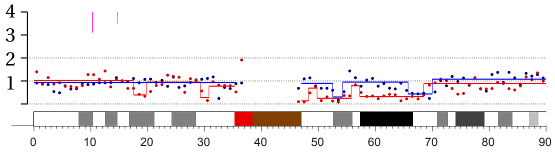

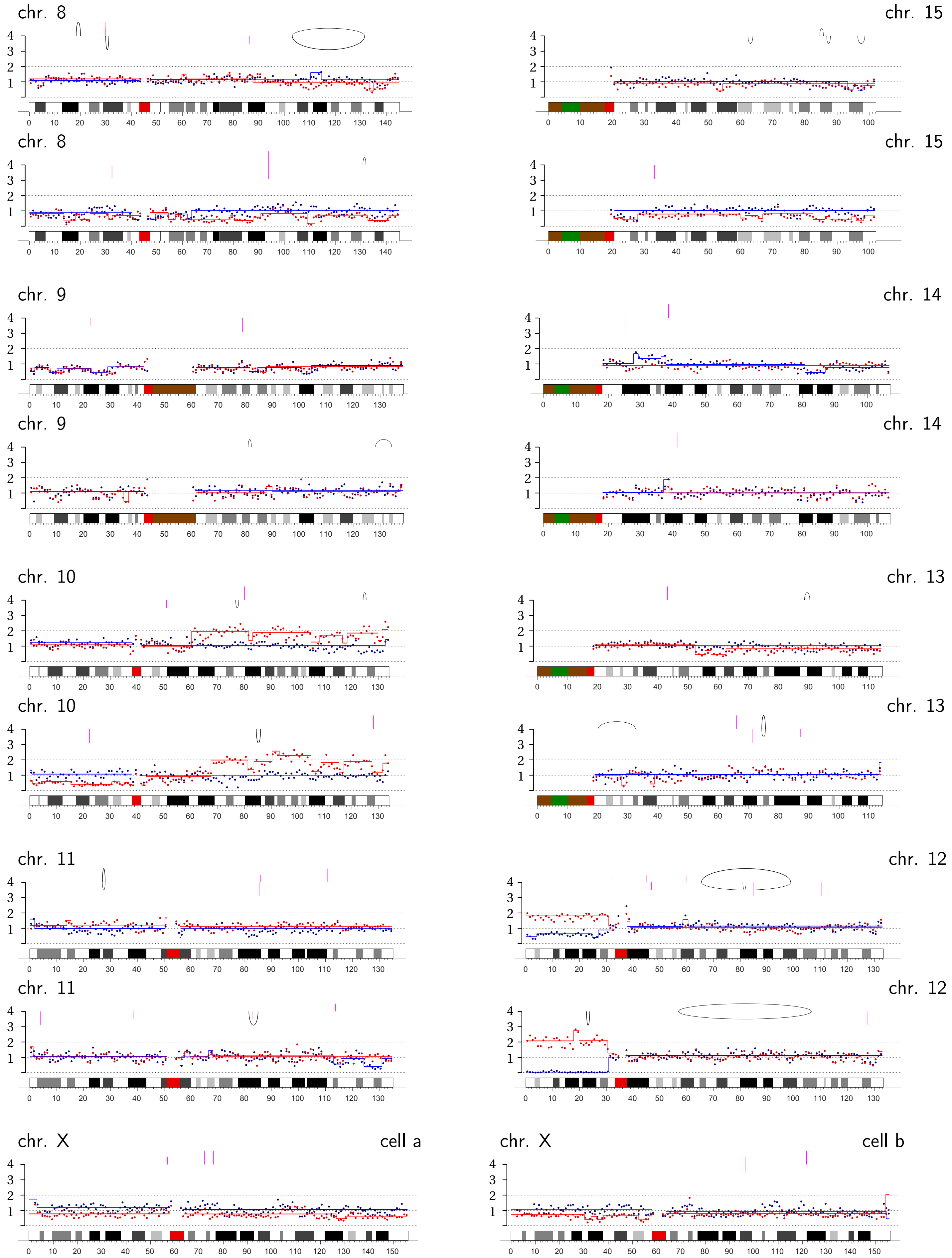
chr. 8
chr. 15
chr. 8
chr. 15
chr. 9
chr. 14
chr. 9
chr. 14
chr. 10
chr. 13
chr. 10
chr. 13
chr. 11
chr. 12
chr. 11
chr. 12
chr. X
cell a
chr. X
cell b

**Bridge Pair 4-7** | CRISPR: Chromosome 4B p-ter.

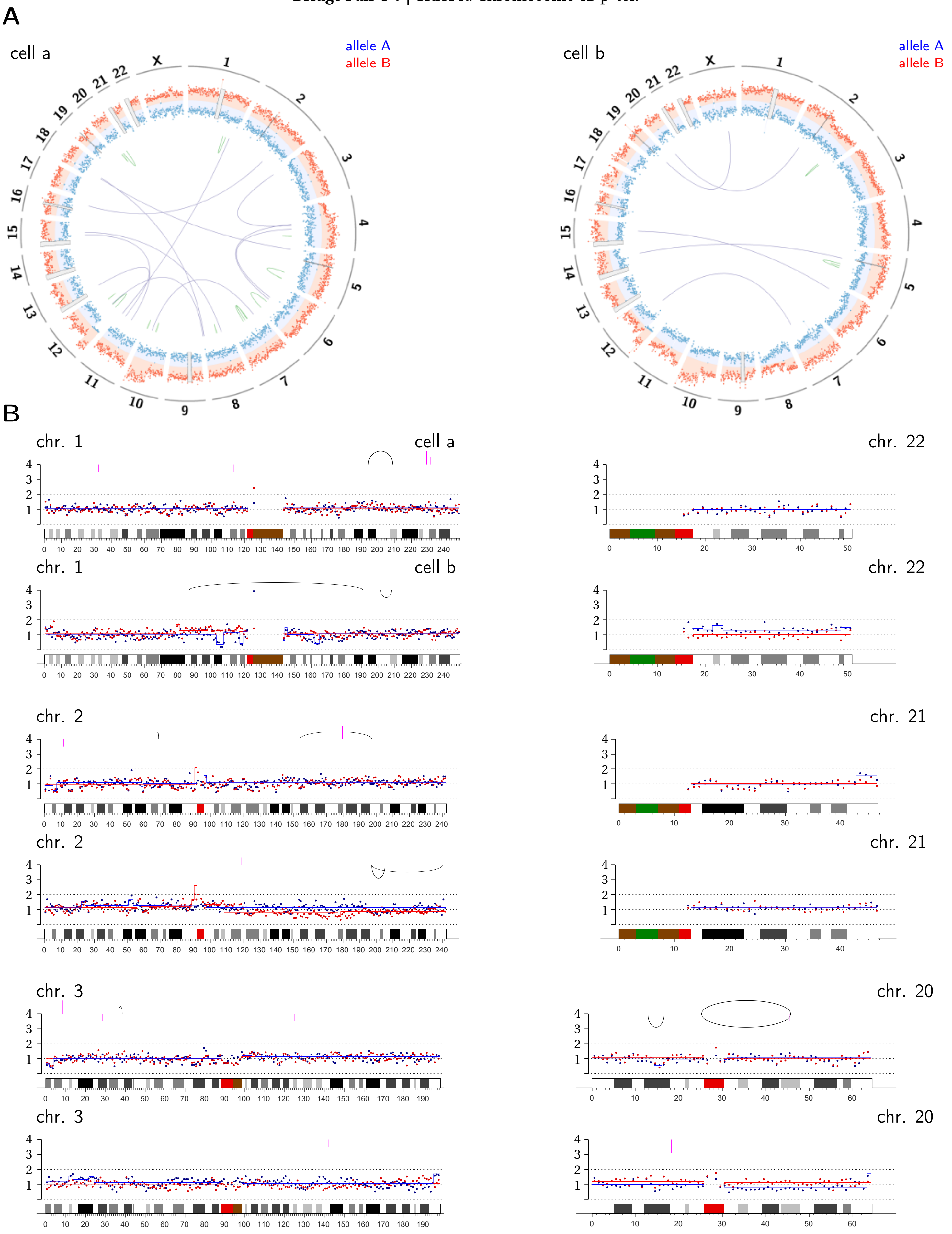

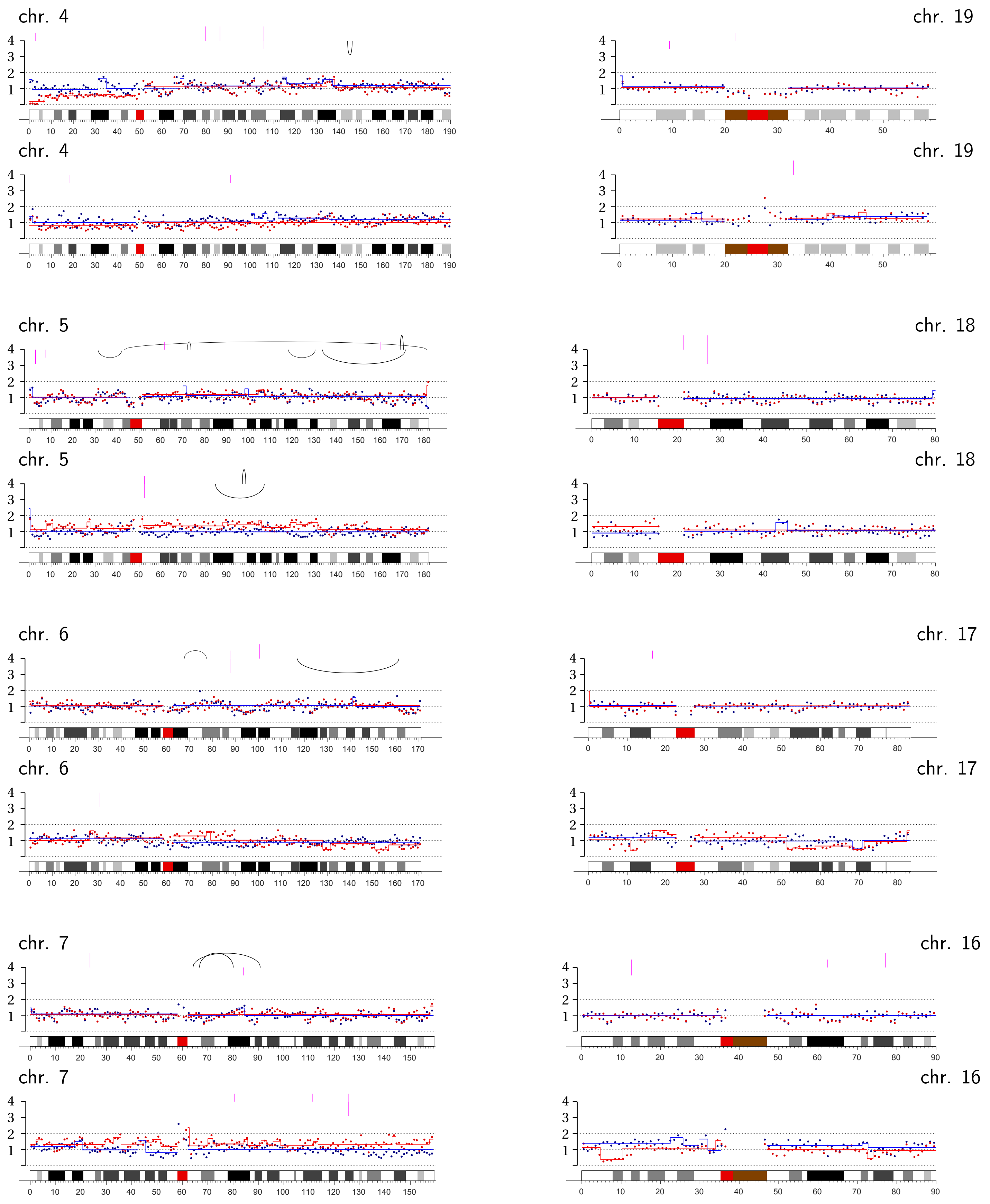
chr. 4
chr. 19
chr. 4
chr. 19
chr. 5
chr. 18
chr. 5
chr. 18
chr. 6
chr. 17
chr. 6
chr. 17
chr. 7
chr. 16
chr. 7
chr. 16

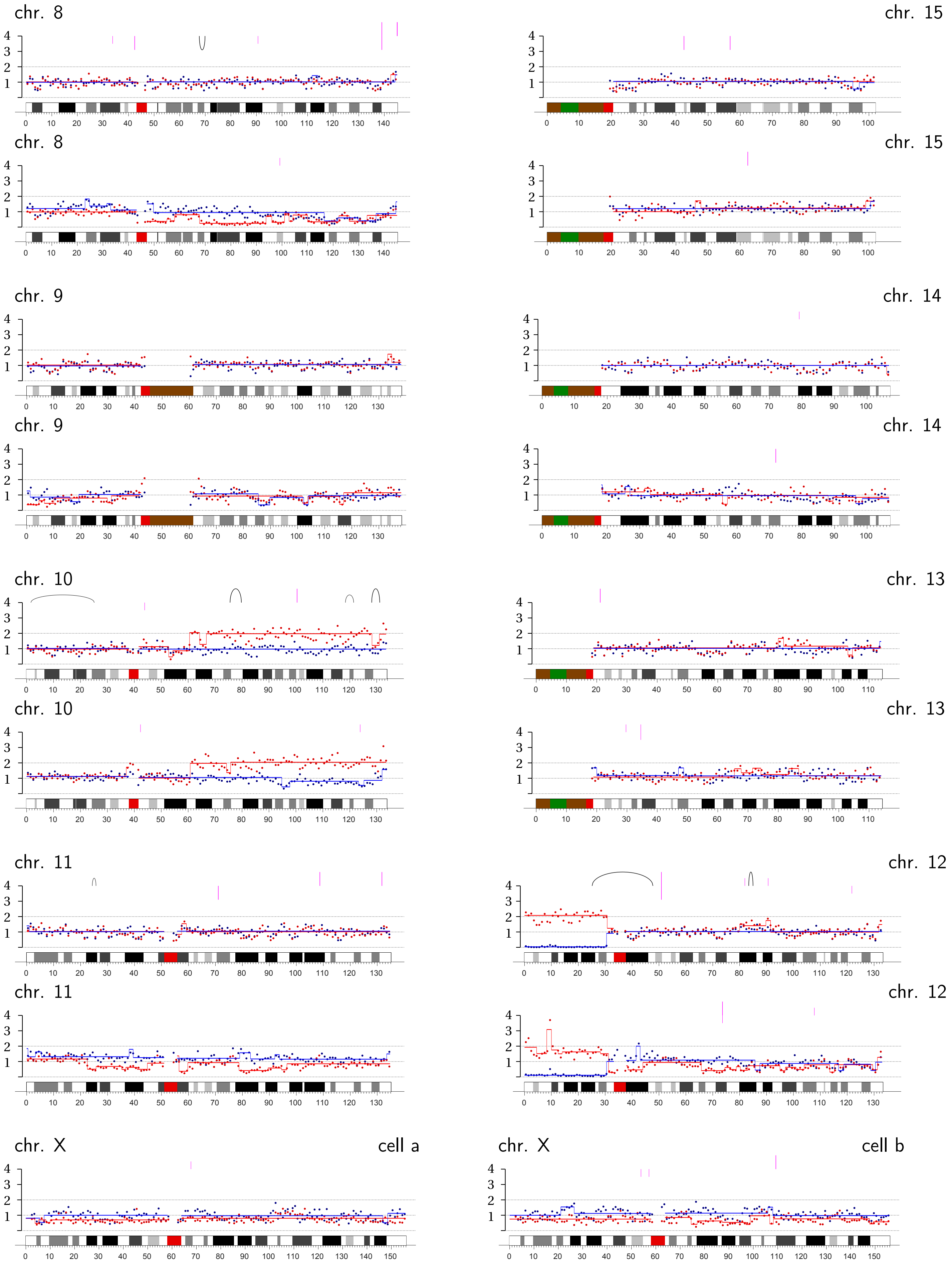
chr. 8
chr. 15
chr. 8
chr. 15
chr. 9
chr. 14
chr. 9
chr. 14
chr. 10
chr. 13
chr. 10
chr. 13
chr. 11
chr. 12
chr. 11
chr. 12
chr. X
cell a
chr. X
cell b

**Bridge Pair 4-8** | CRISPR: **Chromosome 4A and B (p-ter)**.

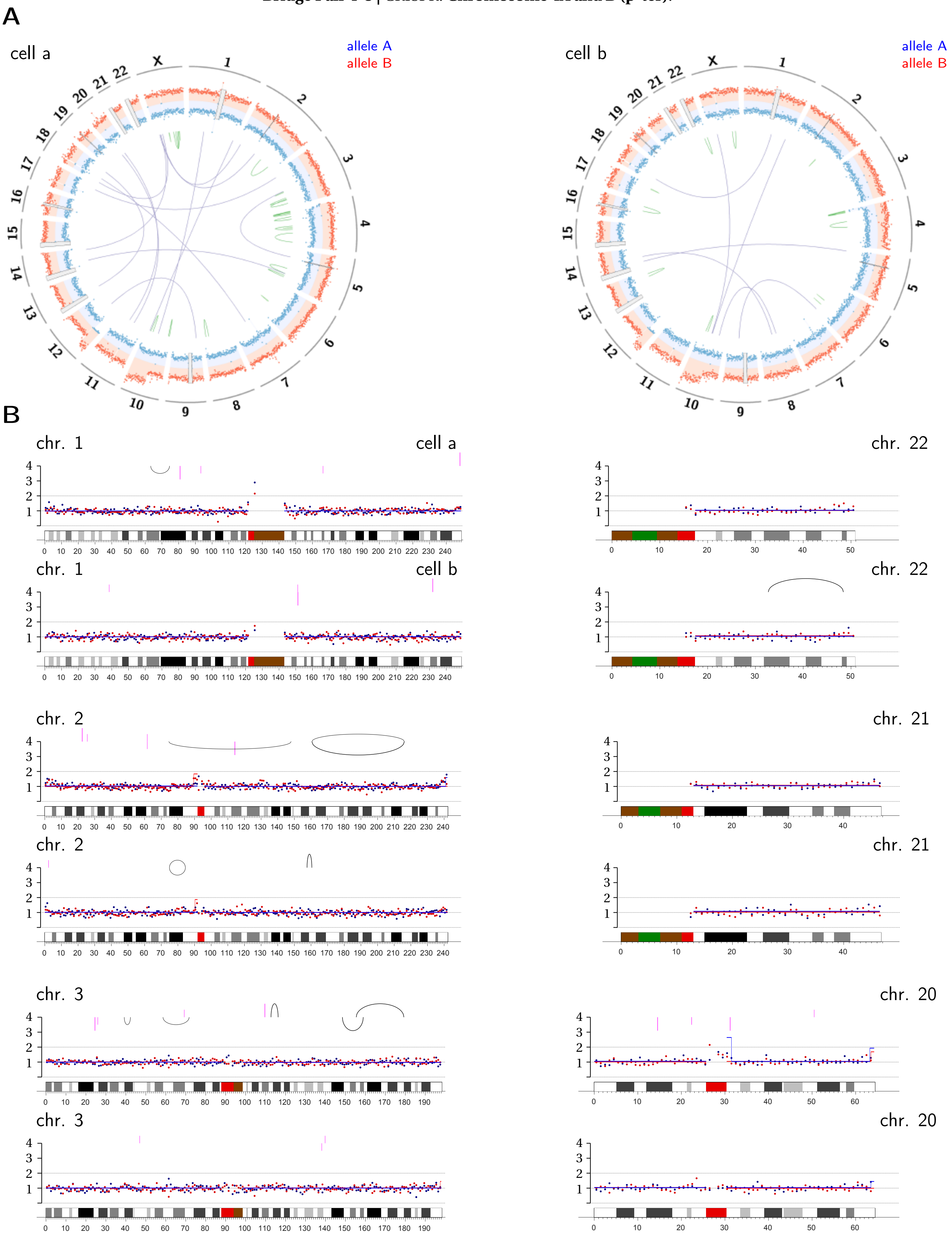

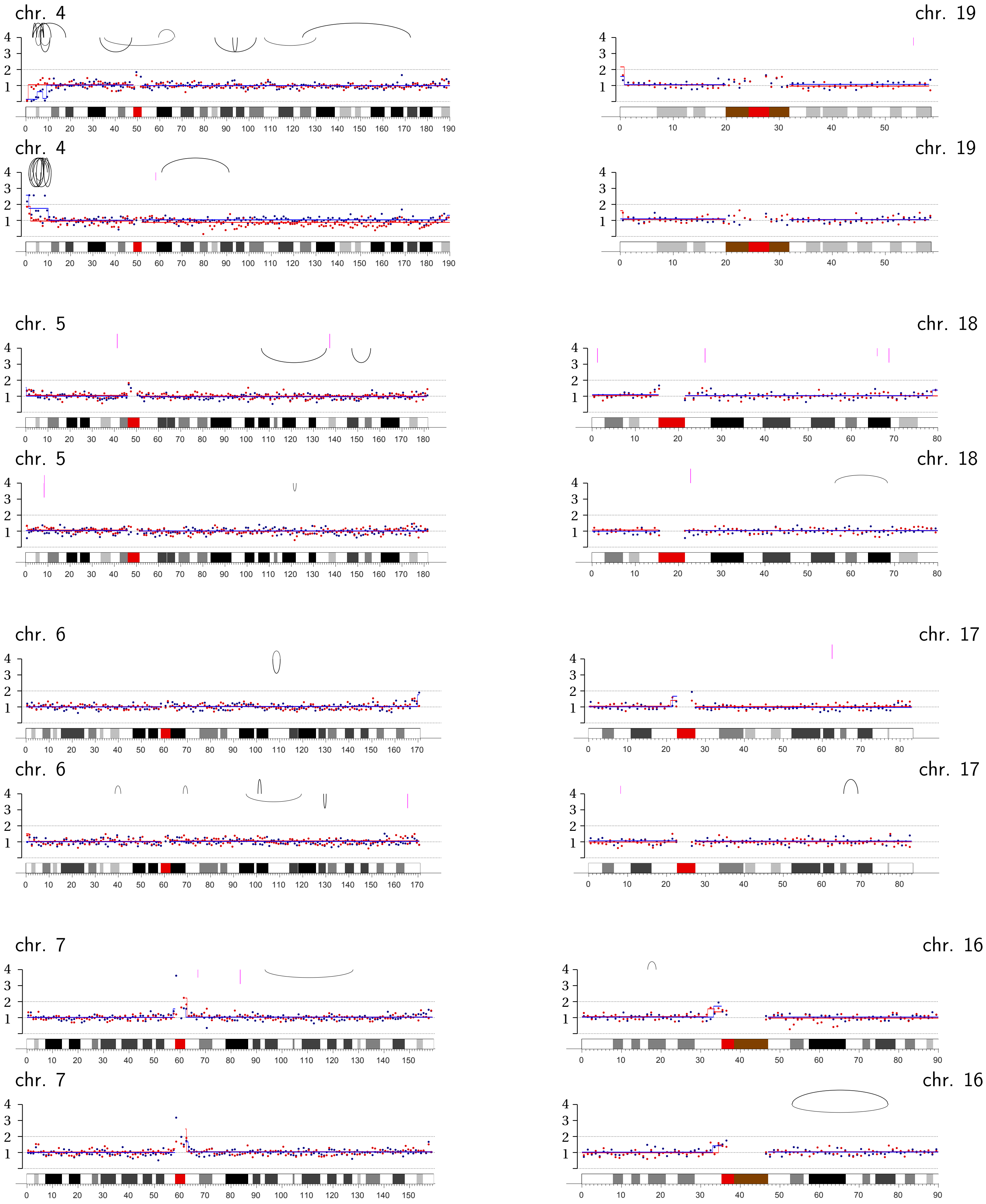
chr. 4
chr. 19
chr. 4
chr. 19
chr. 5
chr. 18
chr. 5
chr. 18
chr. 6
chr. 17
chr. 6
chr. 17
chr. 7
chr. 16
chr. 7
chr. 16

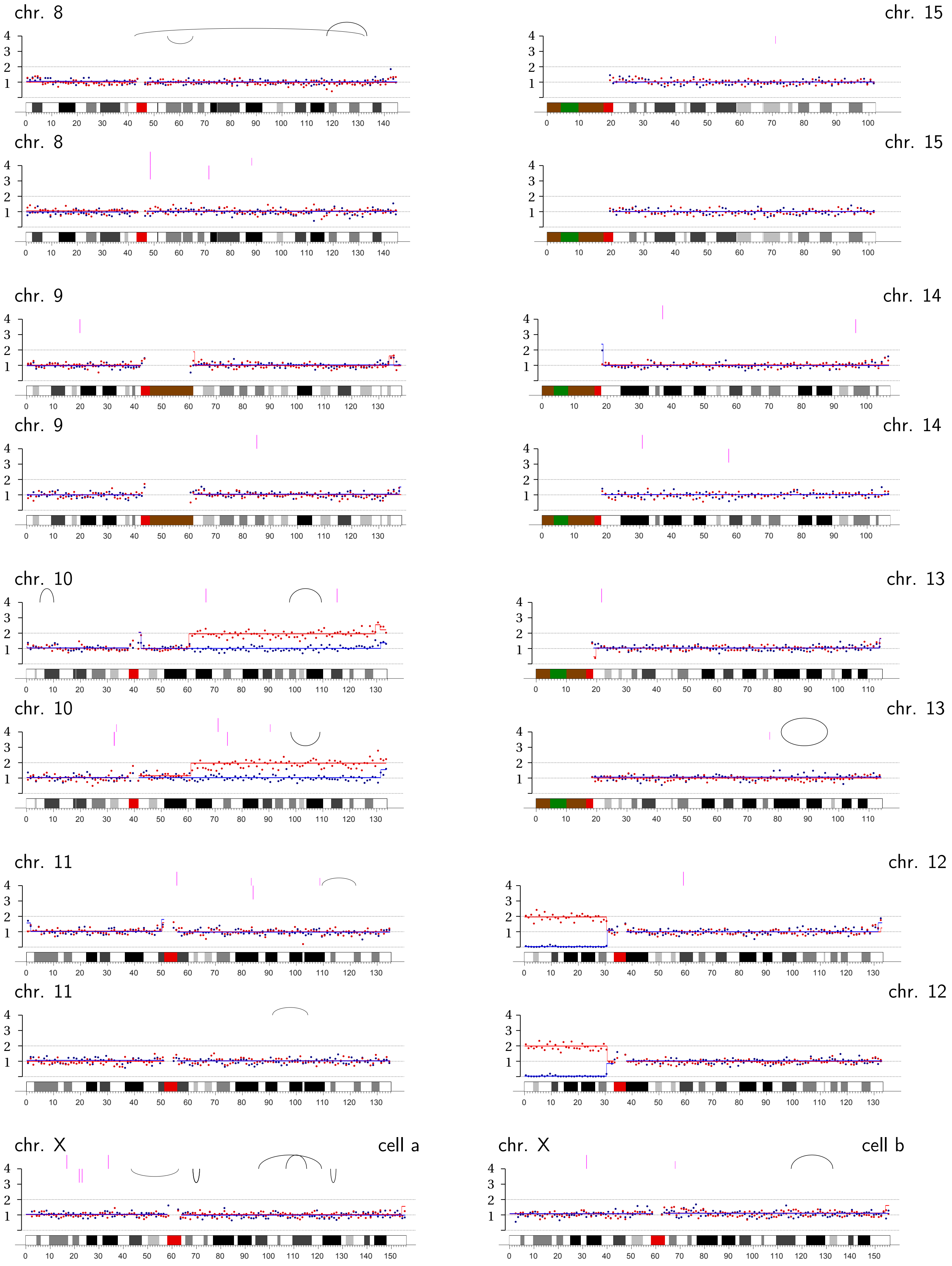
chr. 8
chr. 15
chr. 8
chr. 15
chr. 9
chr. 14
chr. 9
chr. 14
chr. 10
chr. 13
chr. 10
chr. 13
chr. 11
chr. 12
chr. 11
chr. 12
chr. X
cell a
chr. X
cell b

**Bridge Pair TREX1-1** | Transient TRF2-DN expression + TREX1 knockout: Chromosome 21A q-ter.

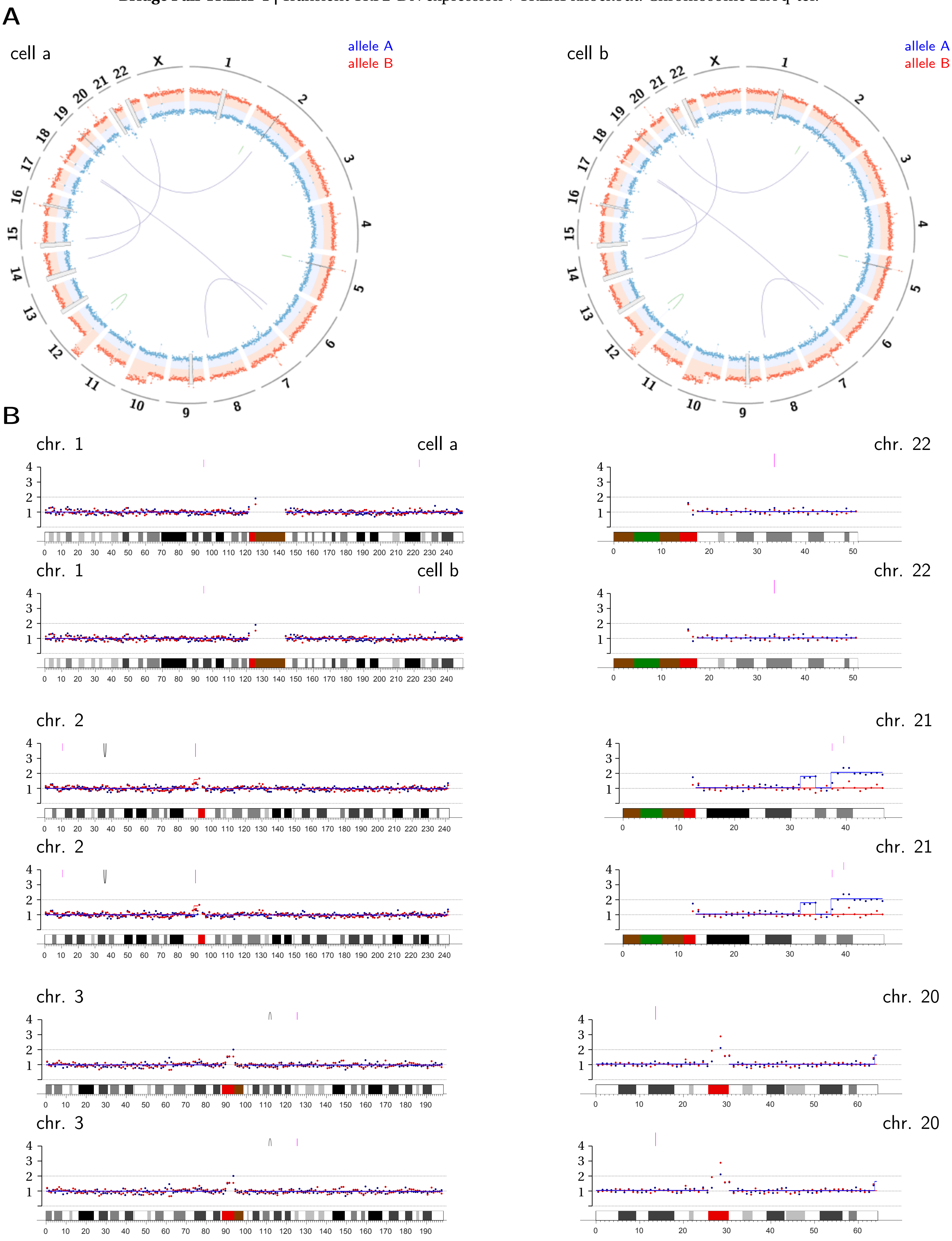

chr. 4

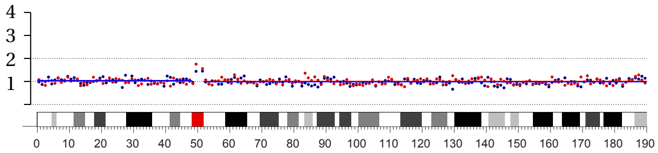

chr. 4

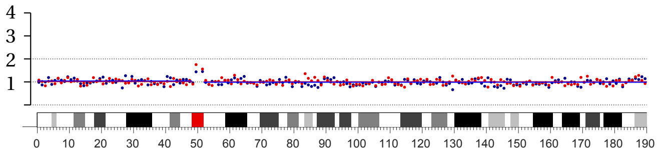

chr. 5

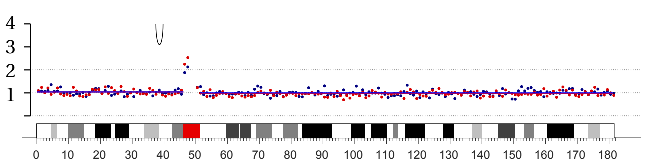

chr. 5

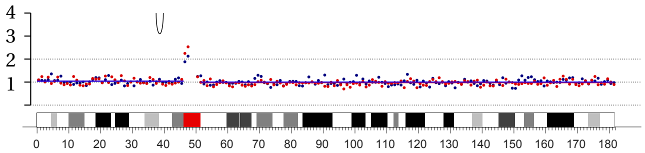

chr. 6

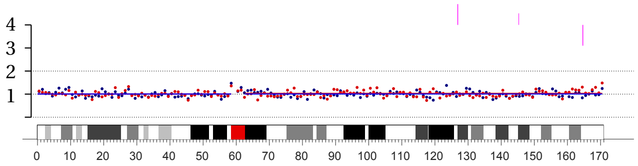

chr. 6

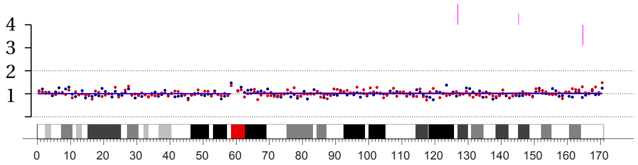

chr. 7

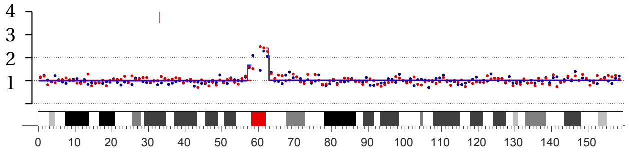

chr. 7

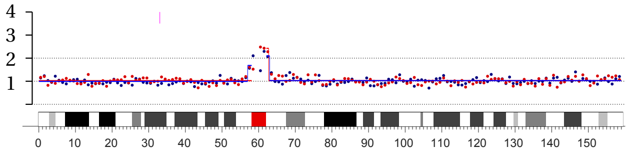

chr. 19

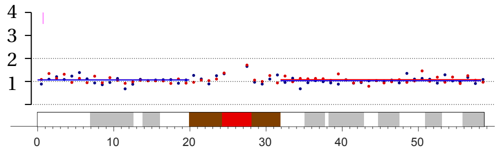

chr. 19

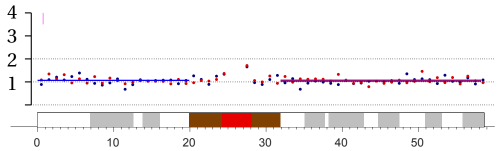

chr. 18

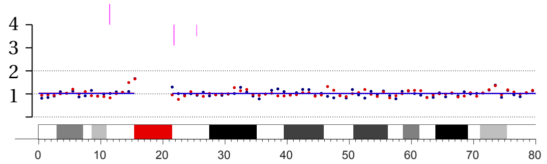

chr. 18

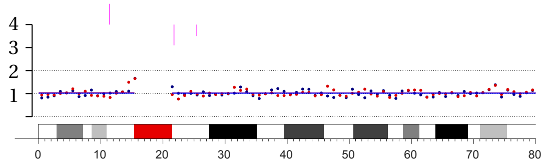

chr. 17

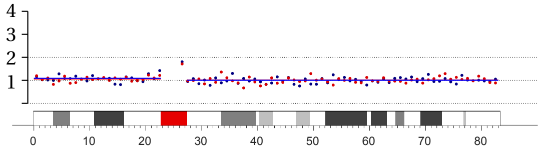

chr. 17

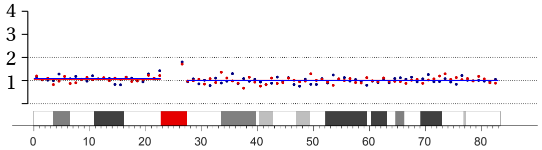

chr. 16

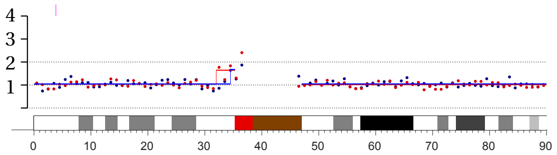

chr. 16

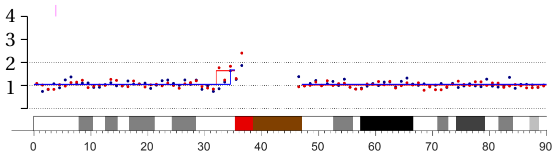

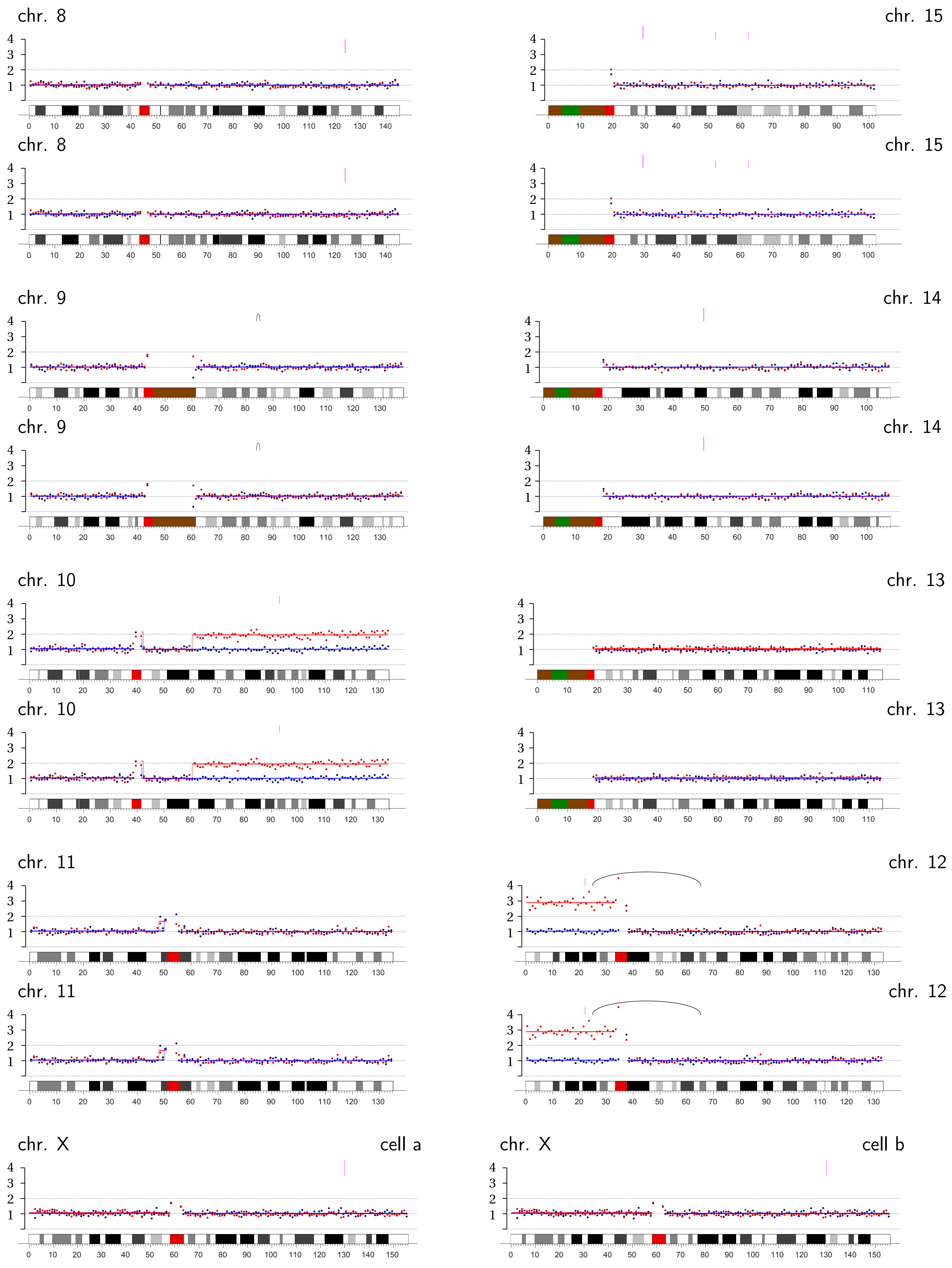
chr. 8
chr. 15
chr. 8
chr. 15
chr. 9
chr. 14
chr. 9
chr. 14
chr. 10
chr. 13
chr. 10
chr. 13
chr. 11
chr. 12
chr. 11
chr. 12
chr. X
cell a
chr. X
cell b

**Bridge Pair TREX1-2 |** Transient TRF2-DN expression + TREX1 knockout: Chromosome 4B q-ter.

A

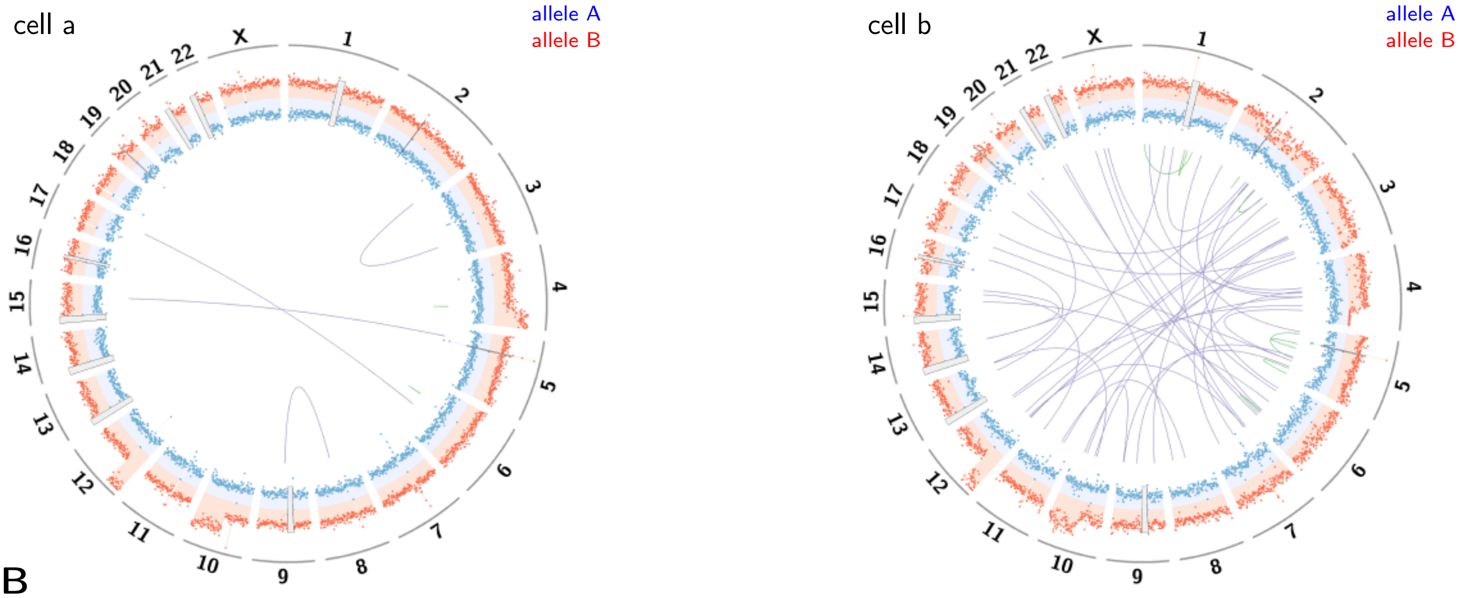


B

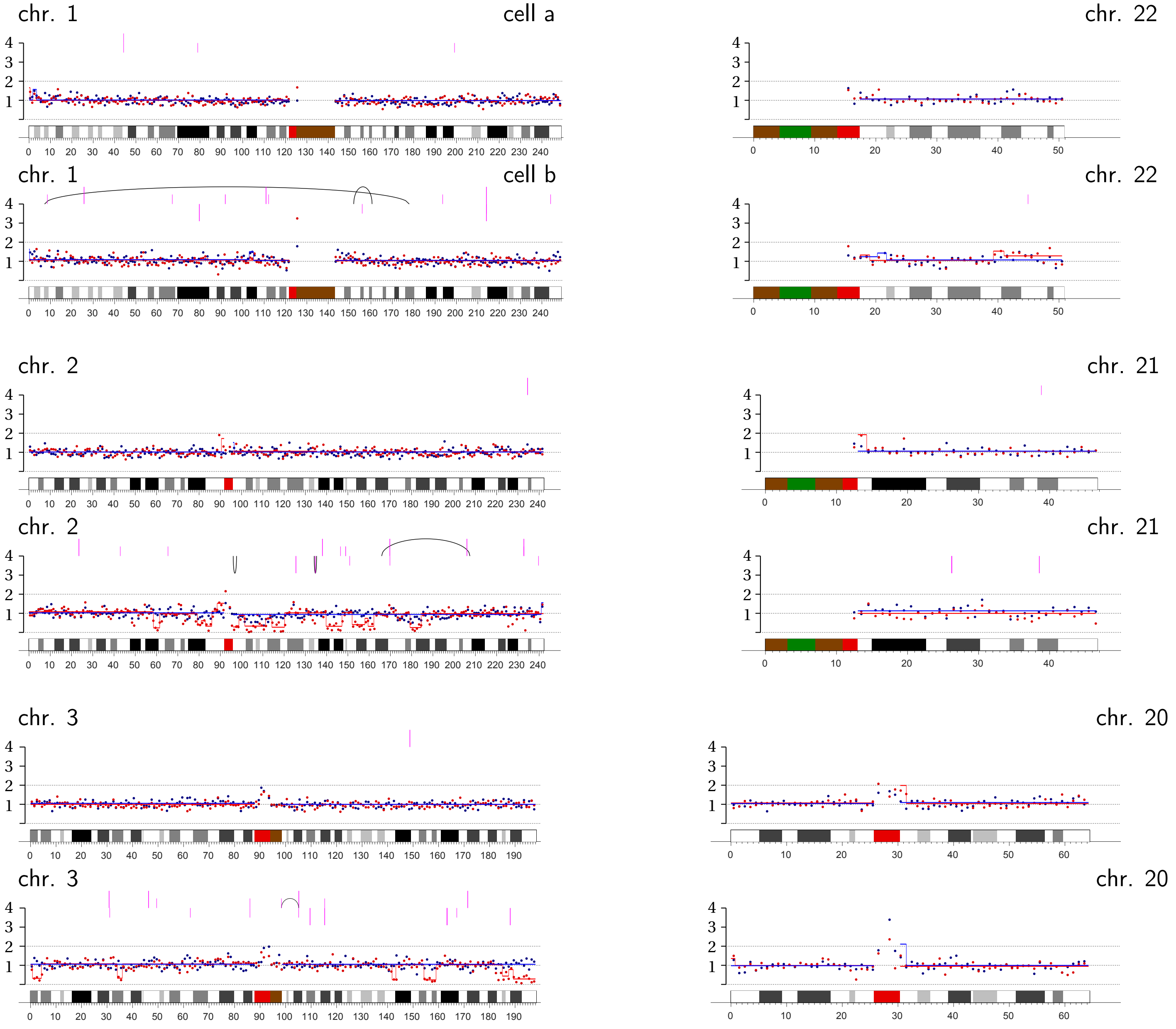

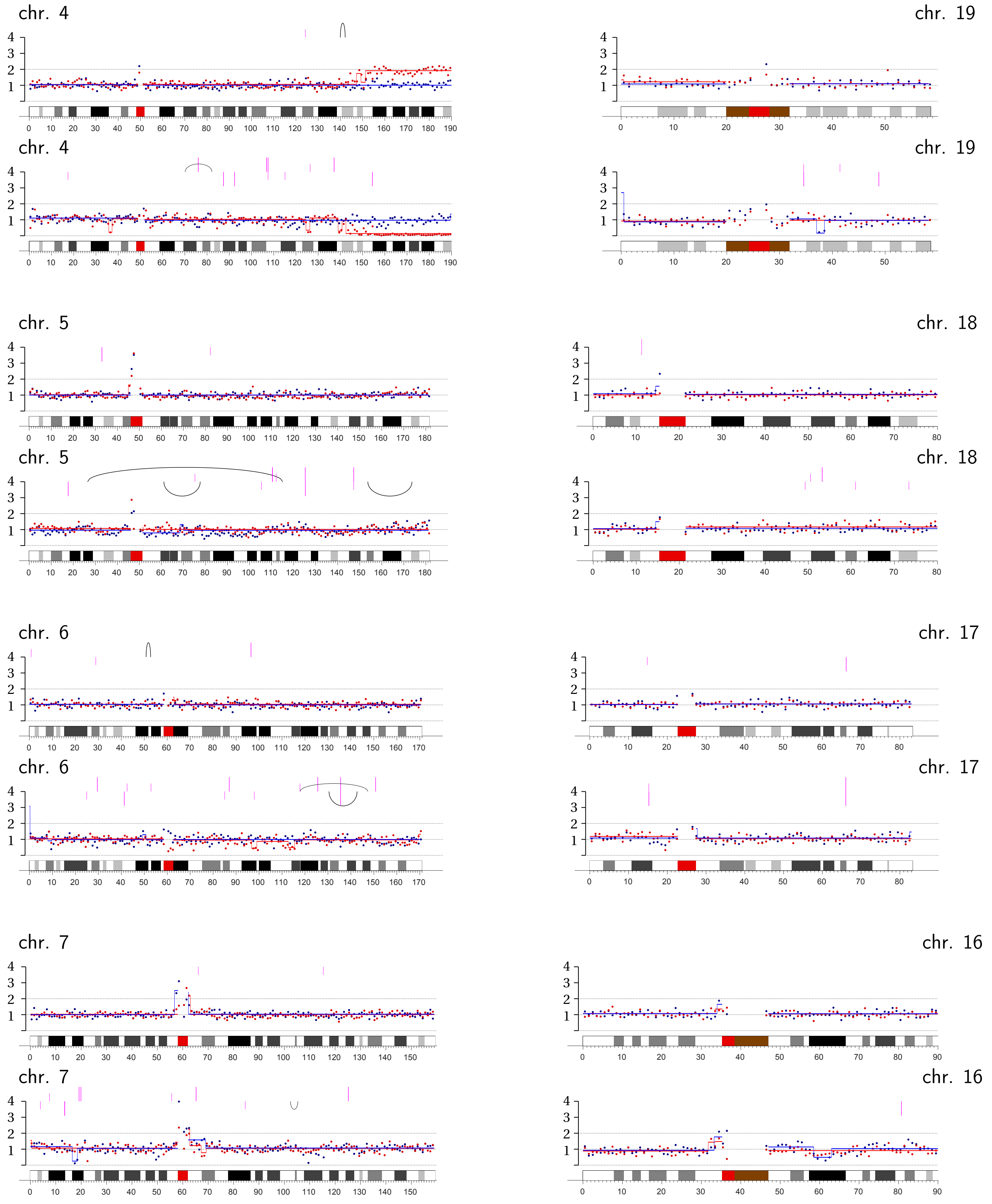
chr. 4
chr. 19
chr. 4
chr. 19
chr. 5
chr. 18
chr. 5
chr. 18
chr. 6
chr. 17
chr. 6
chr. 17
chr. 7
chr. 16
chr. 7
chr. 16

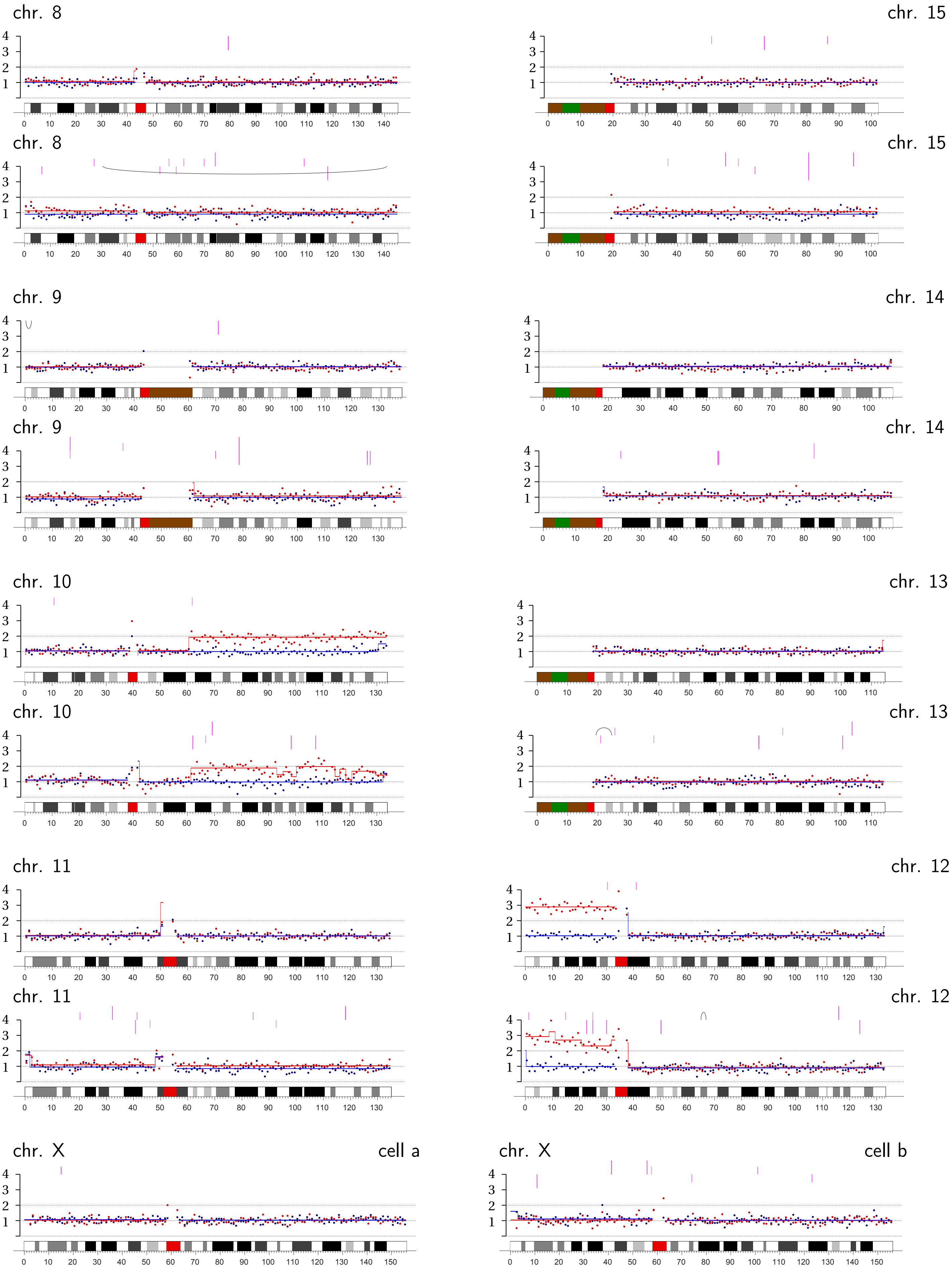
chr. 8
chr. 15
chr. 8
chr. 15
chr. 9
chr. 14
chr. 9
chr. 14
chr. 10
chr. 13
chr. 10
chr. 13
chr. 11
chr. 12
chr. 11
chr. 12
chr. X
cell a
chr. X
cell b

**Bridge Pair TREX1-3 |** Transient TRF2-DN expression + TREX1 knockout: Chromosome 7A, p-arm internal; Chromosome 12A, p-arm.

A

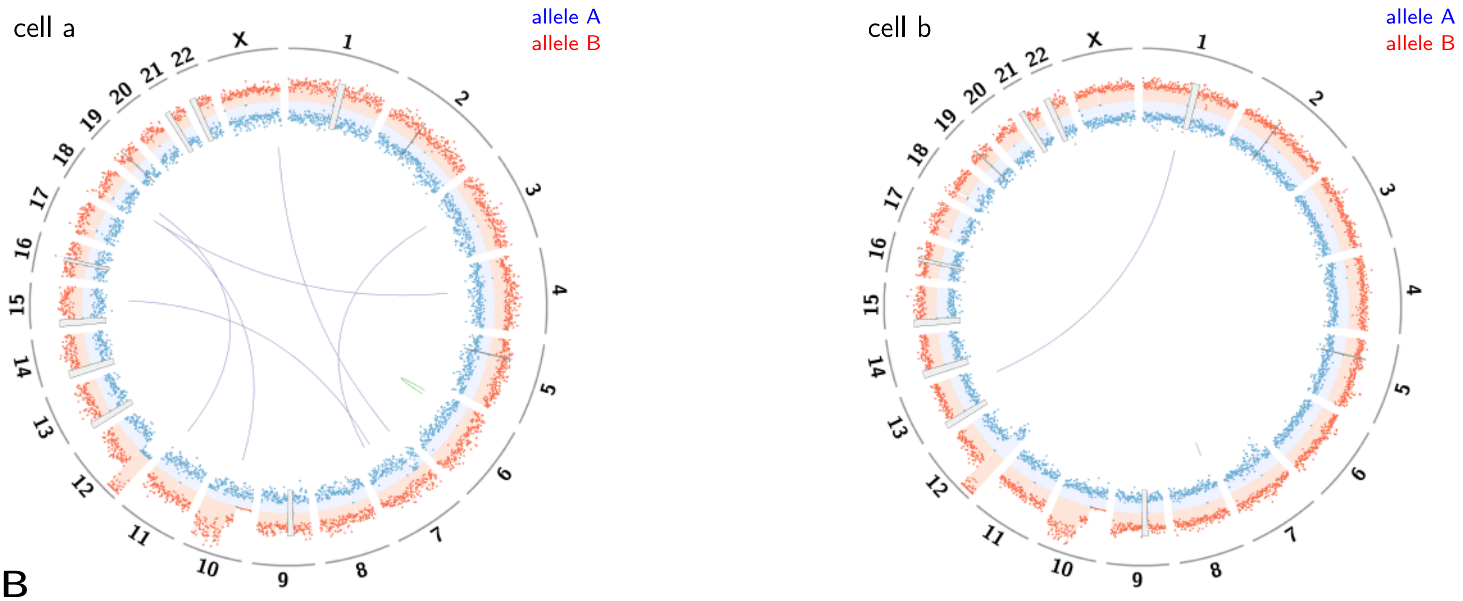


B

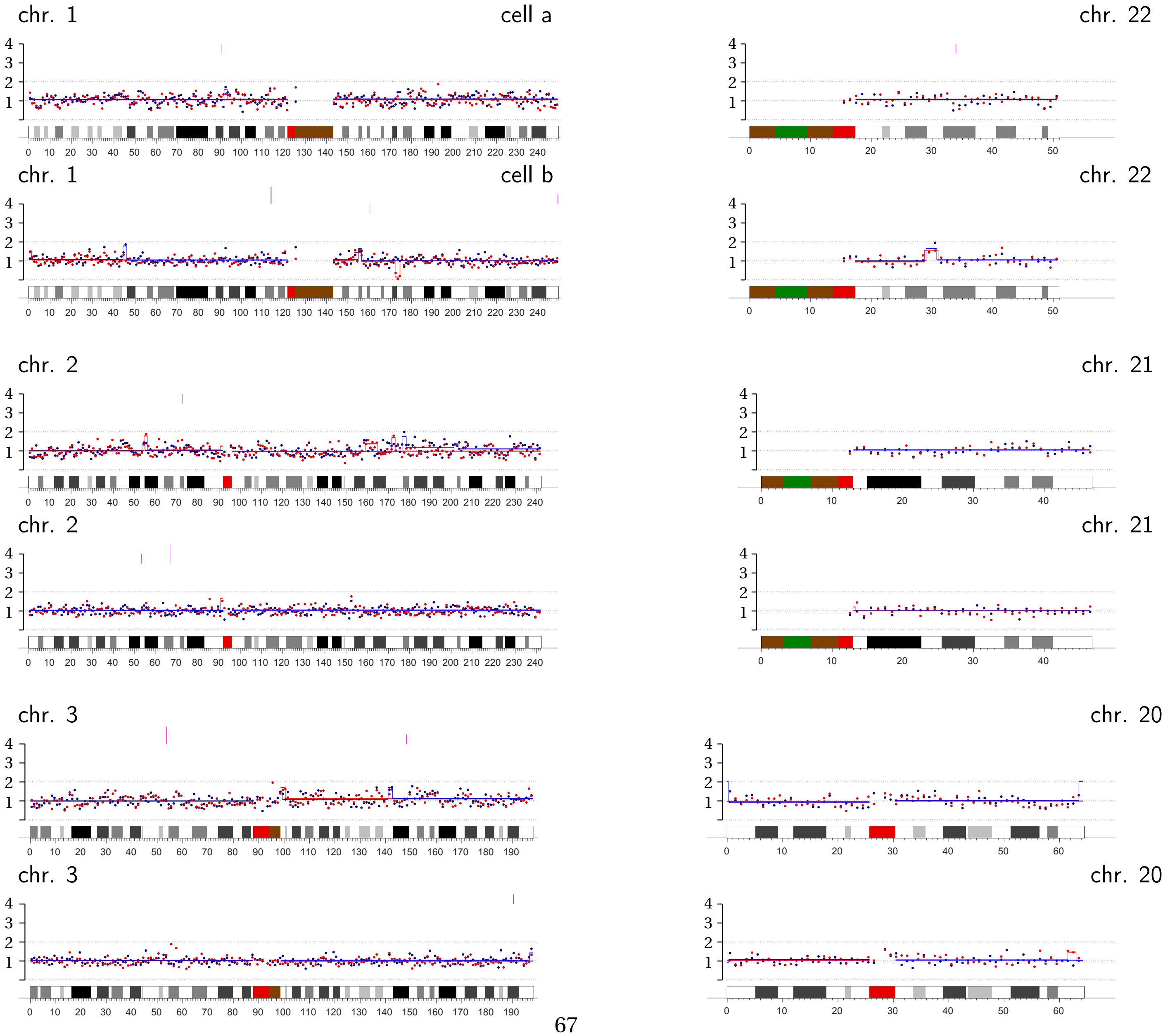

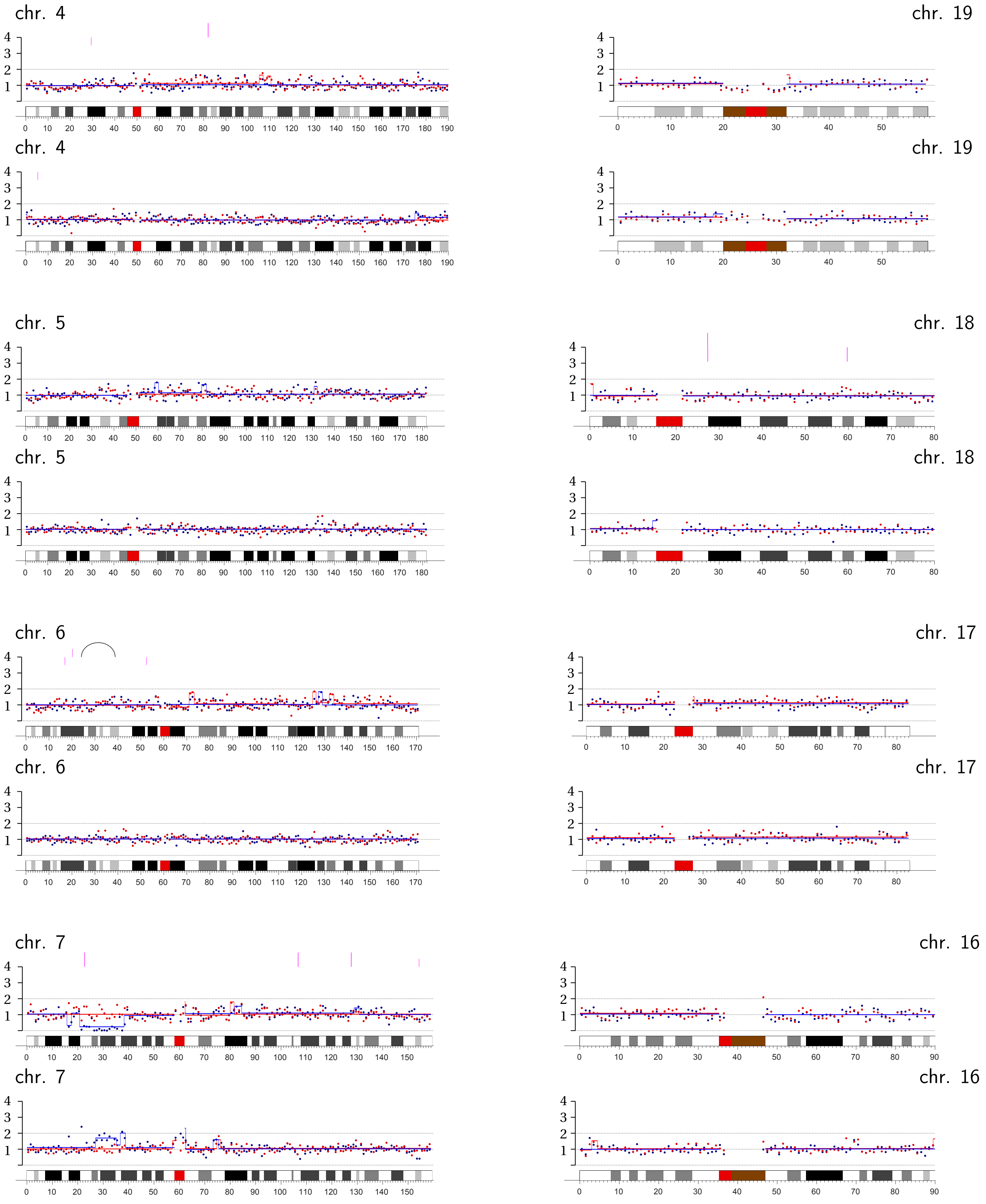
chr. 4
chr. 19
chr. 4
chr. 19
chr. 5
chr. 18
chr. 5
chr. 18
chr. 6
chr. 17
chr. 6
chr. 17
chr. 7
chr. 16
chr. 7
chr. 16

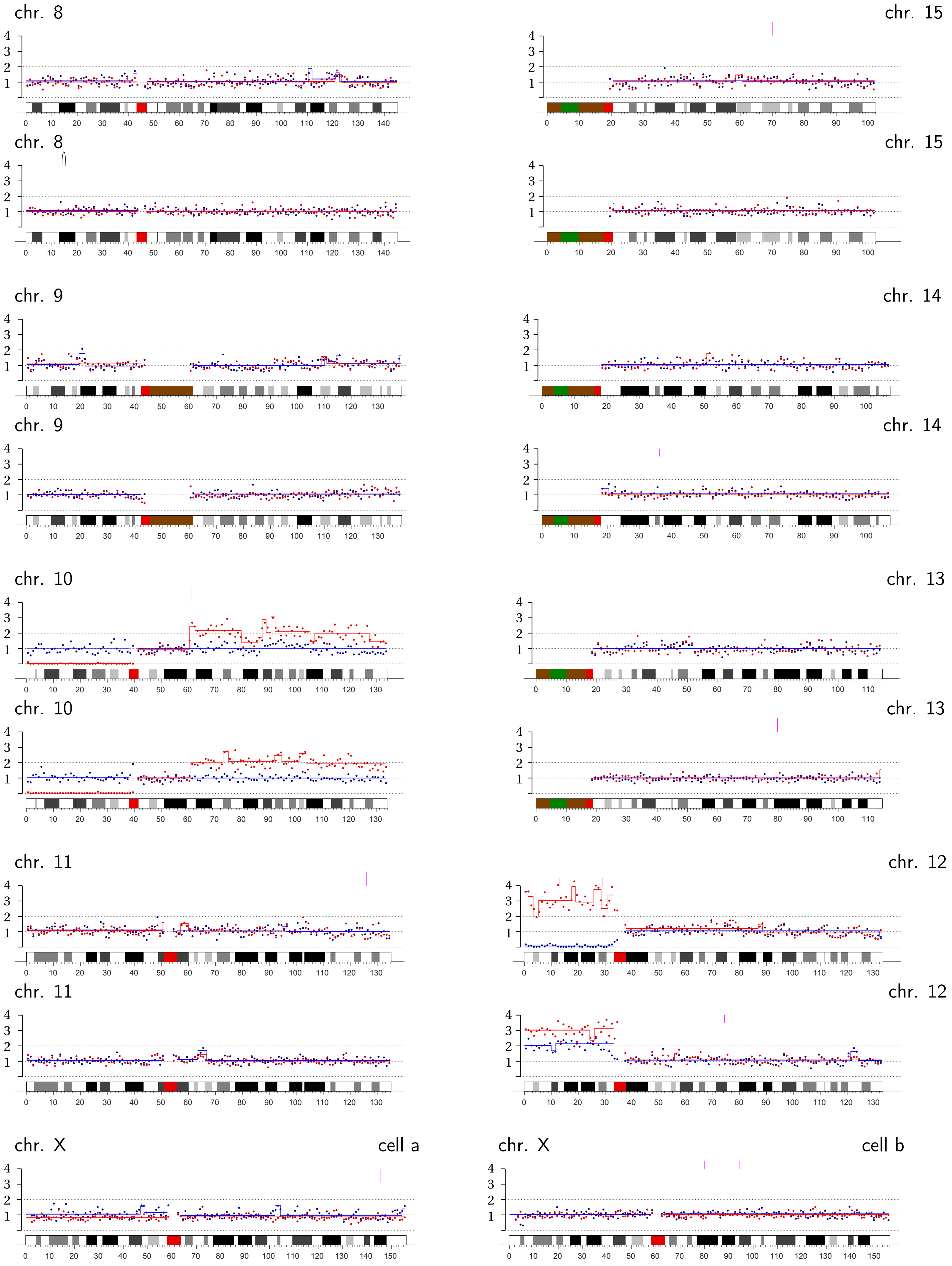
chr. 8
chr. 15
chr. 8
chr. 15
chr. 9
chr. 14
chr. 9
chr. 14
chr. 10
chr. 13
chr. 10
chr. 13
chr. 11
chr. 12
chr. 11
chr. 12
chr. X
cell a
chr. X
cell b

**Bridge Pair TREX1-4** | Transient TRF2-DN expression + TREX1 knockout: Chromosome 16A, q-arm.

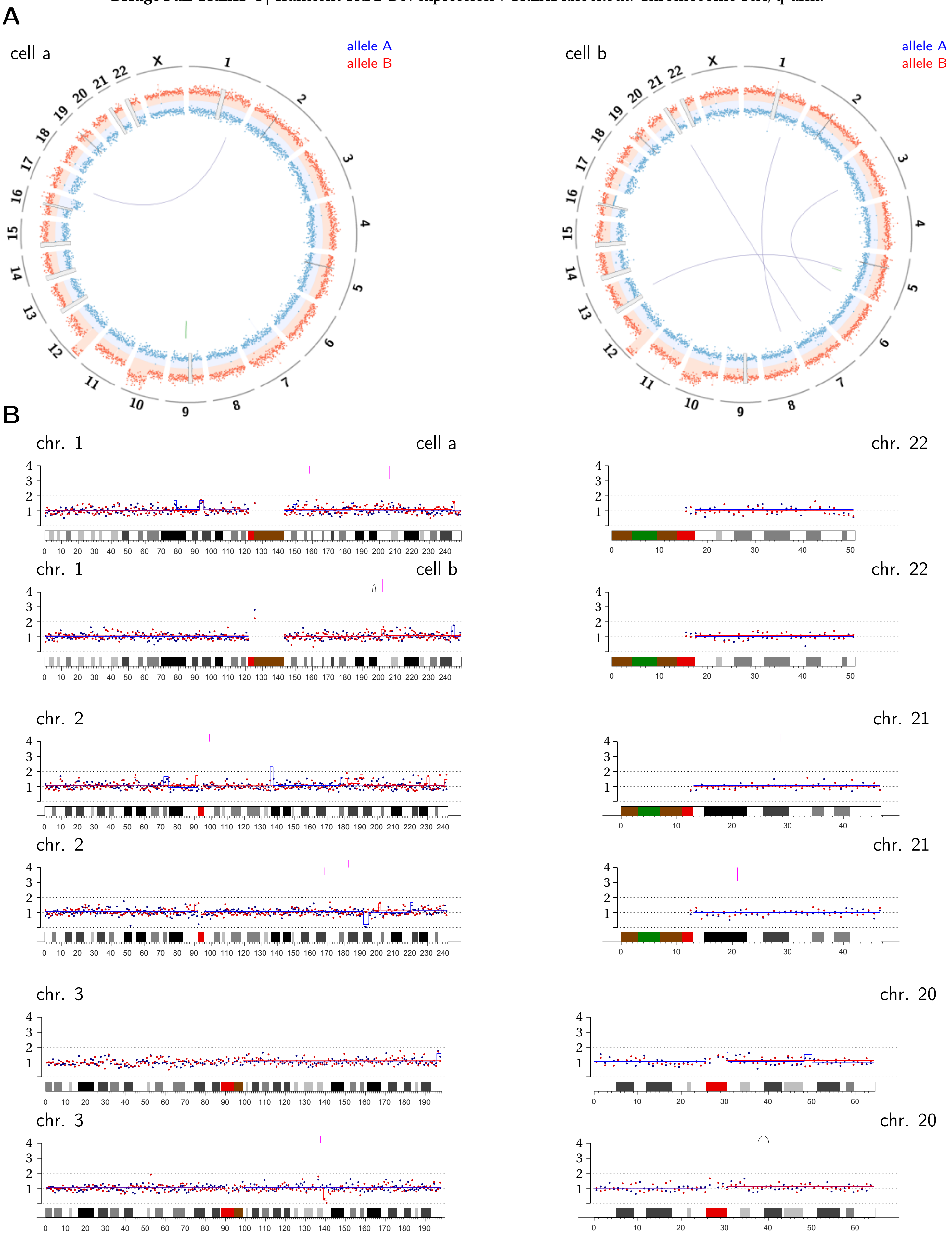

chr. 4

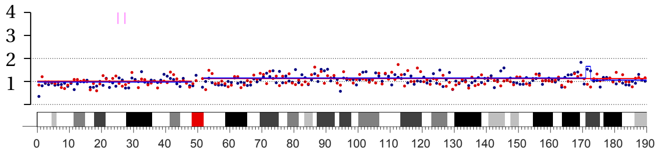

chr. 19

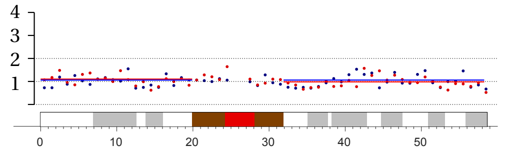

chr. 4

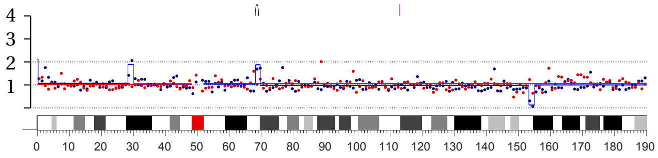

chr. 19

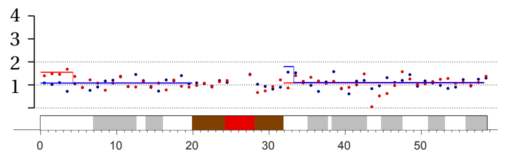

chr. 5

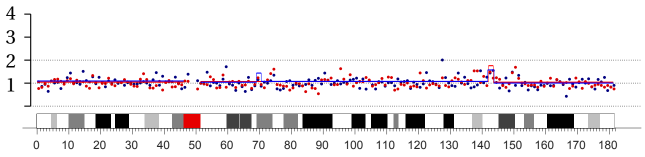

chr. 18

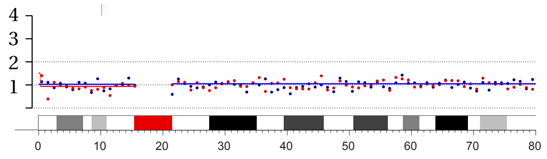

chr. 5

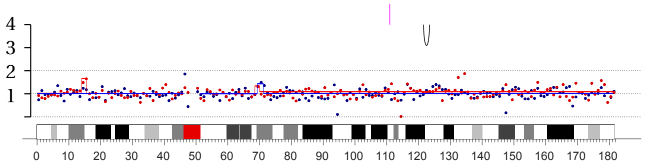

chr. 18

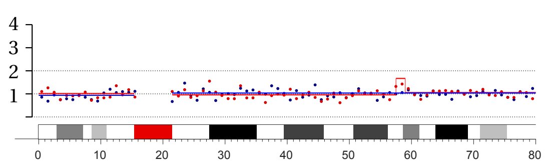

chr. 6

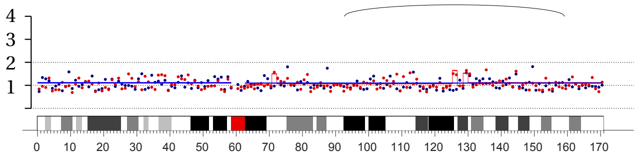

chr. 17

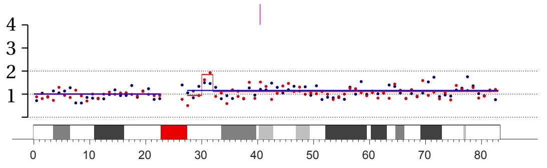

chr. 6

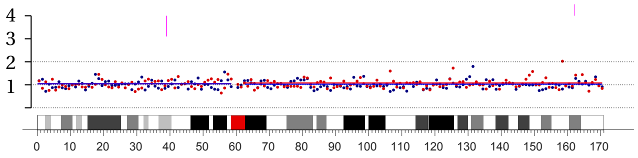

chr. 17

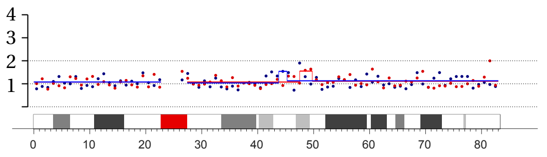

chr. 7

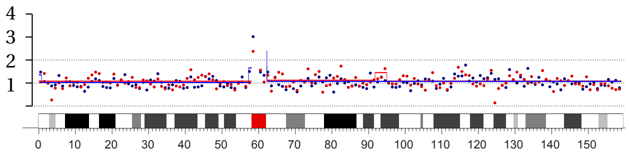

chr. 16

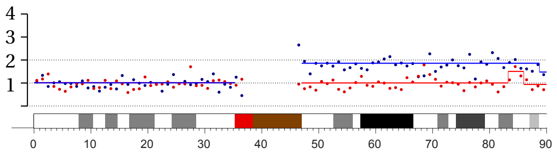

chr. 7

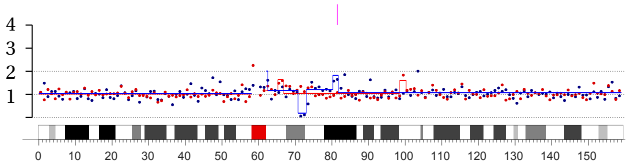

chr. 16

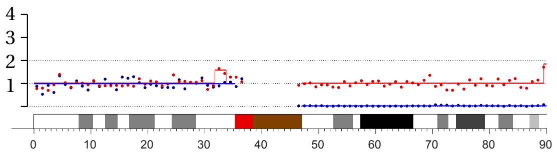

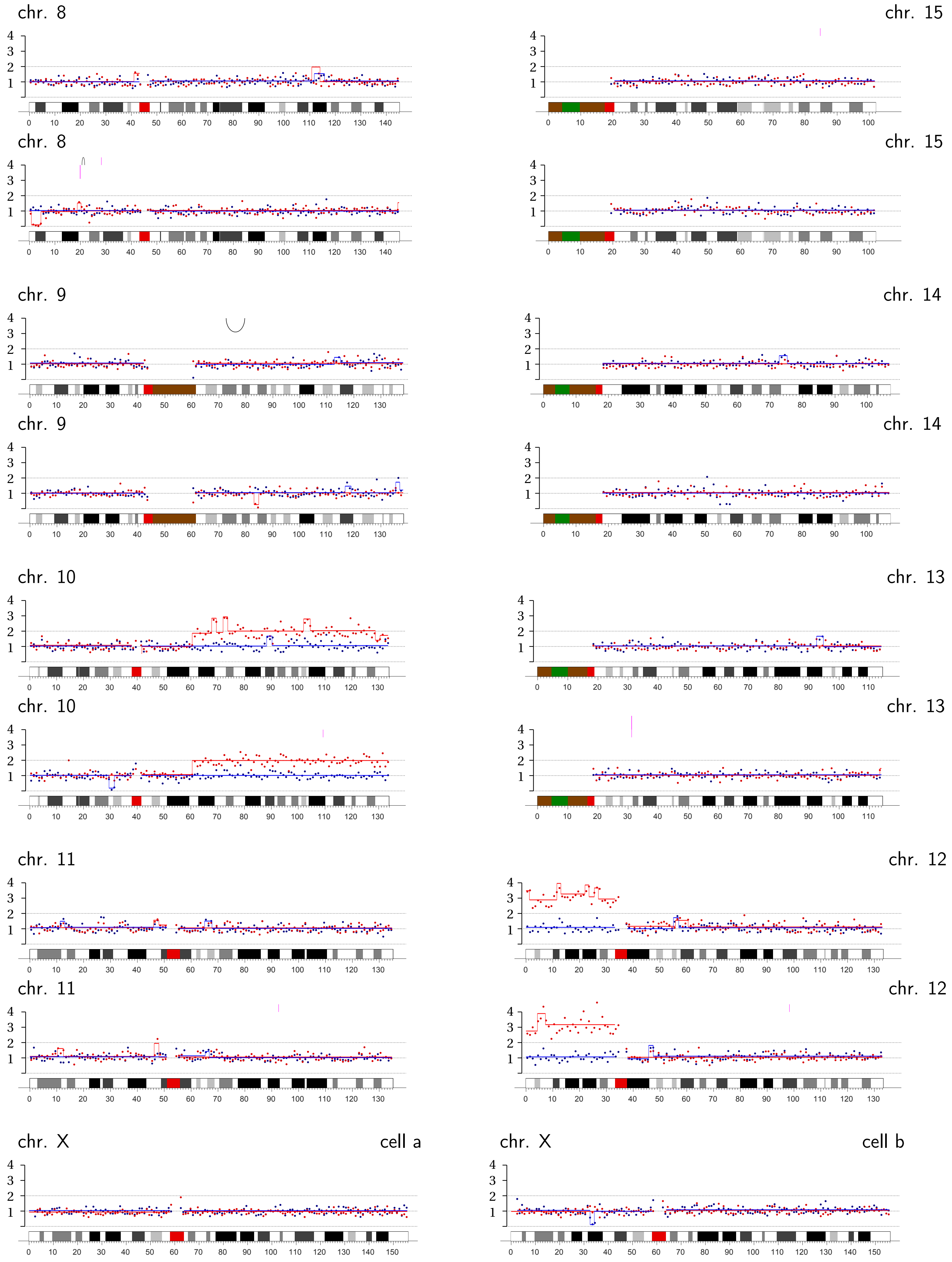
chr. 8
chr. 15
chr. 8
chr. 15
chr. 9
chr. 14
chr. 9
chr. 14
chr. 10
chr. 13
chr. 10
chr. 13
chr. 11
chr. 12
chr. 11
chr. 12
chr. X
cell a
chr. X
cell b

**Bridge Pair TREX1-5 |** Transient TRF2-DN expression + TREX1 knockout: Chromosome 2A, q-arm; XB, q-ter.

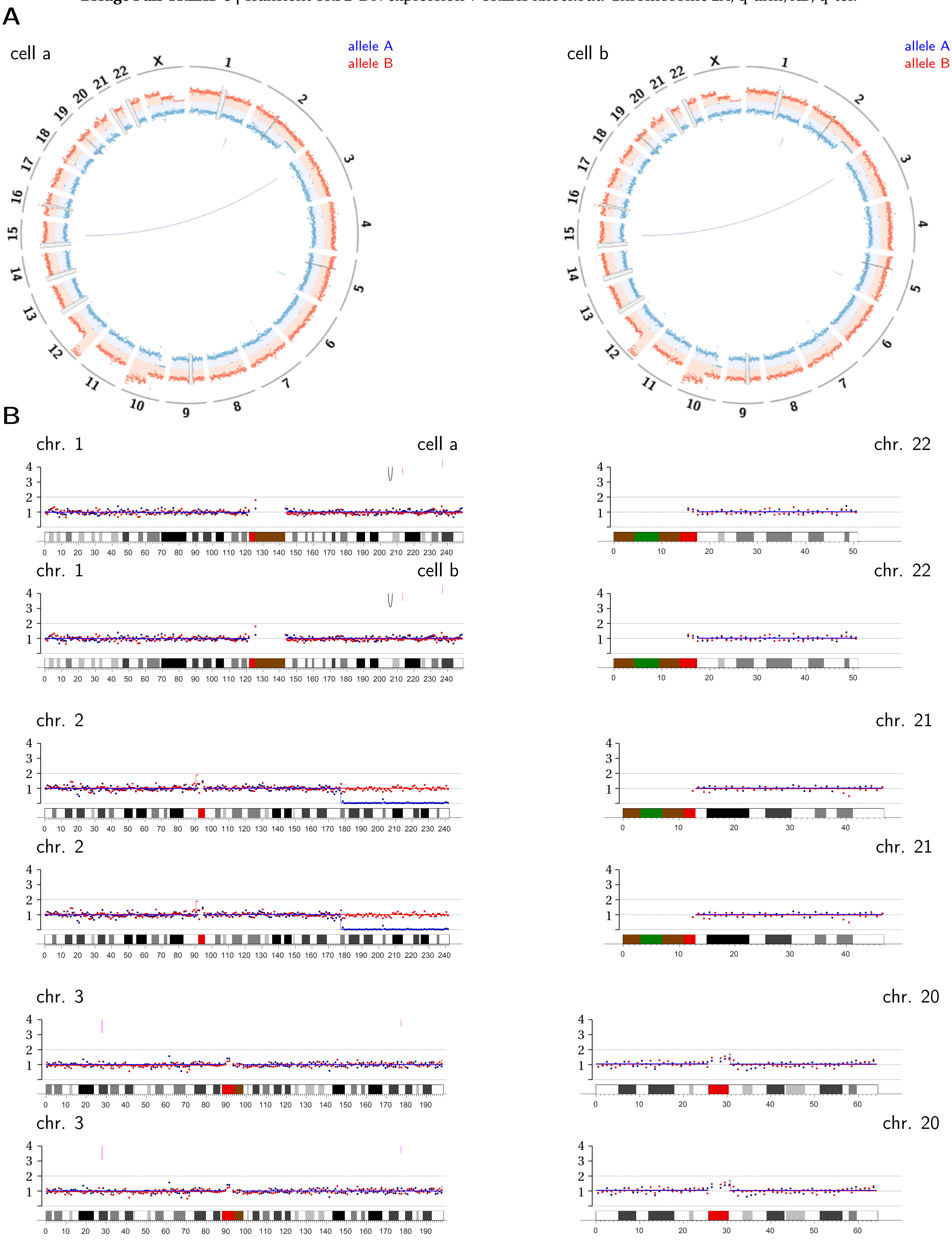

chr. 4

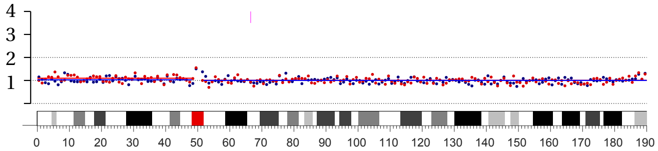


chr. 4

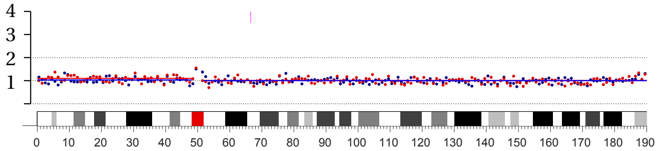


chr. 19

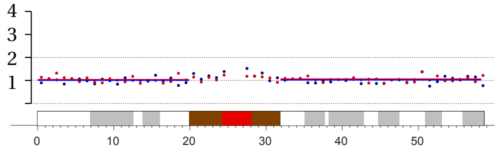


chr. 19

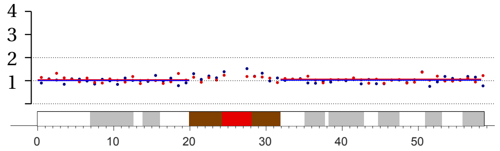


chr. 5

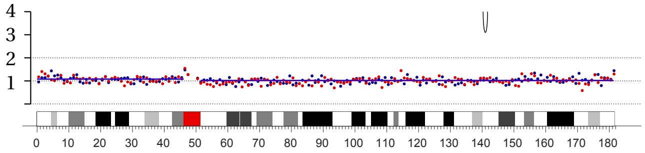


chr. 5

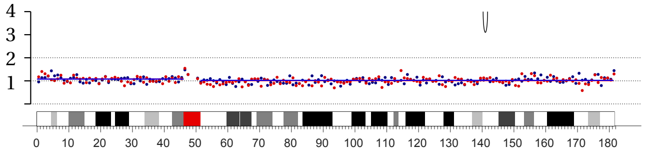


chr. 18

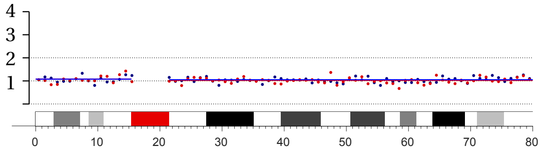


chr. 18

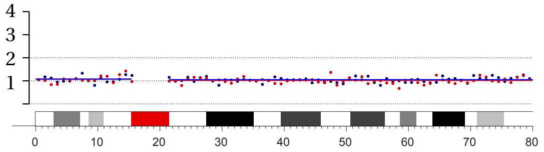


chr. 6

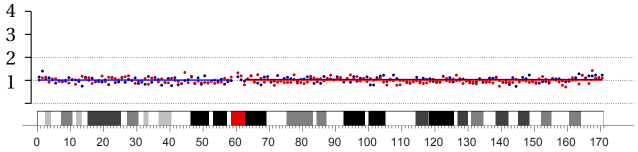


chr. 6

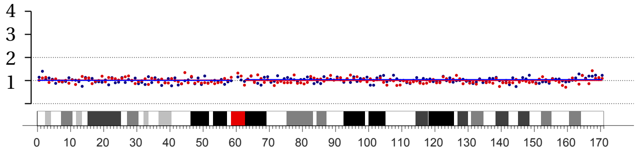


chr. 17

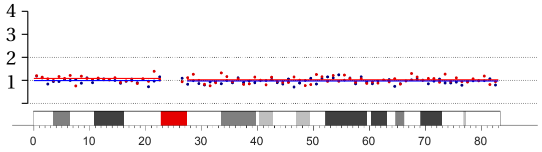


chr. 17

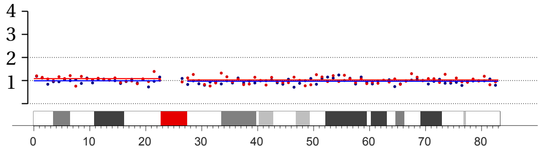


chr. 7

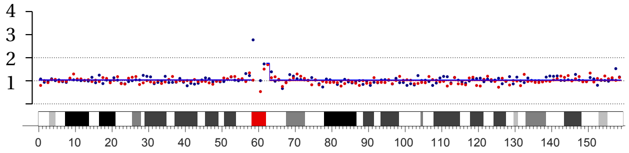


chr. 7

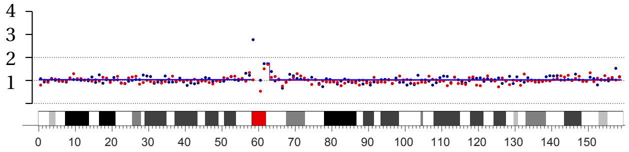


chr. 16

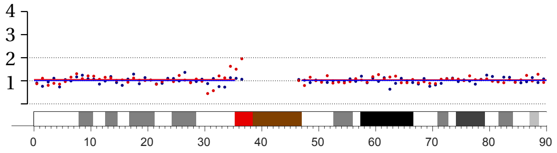


chr. 16

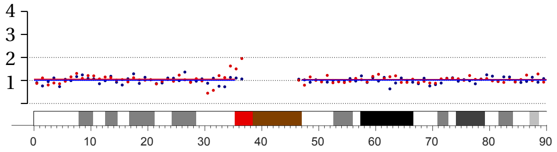

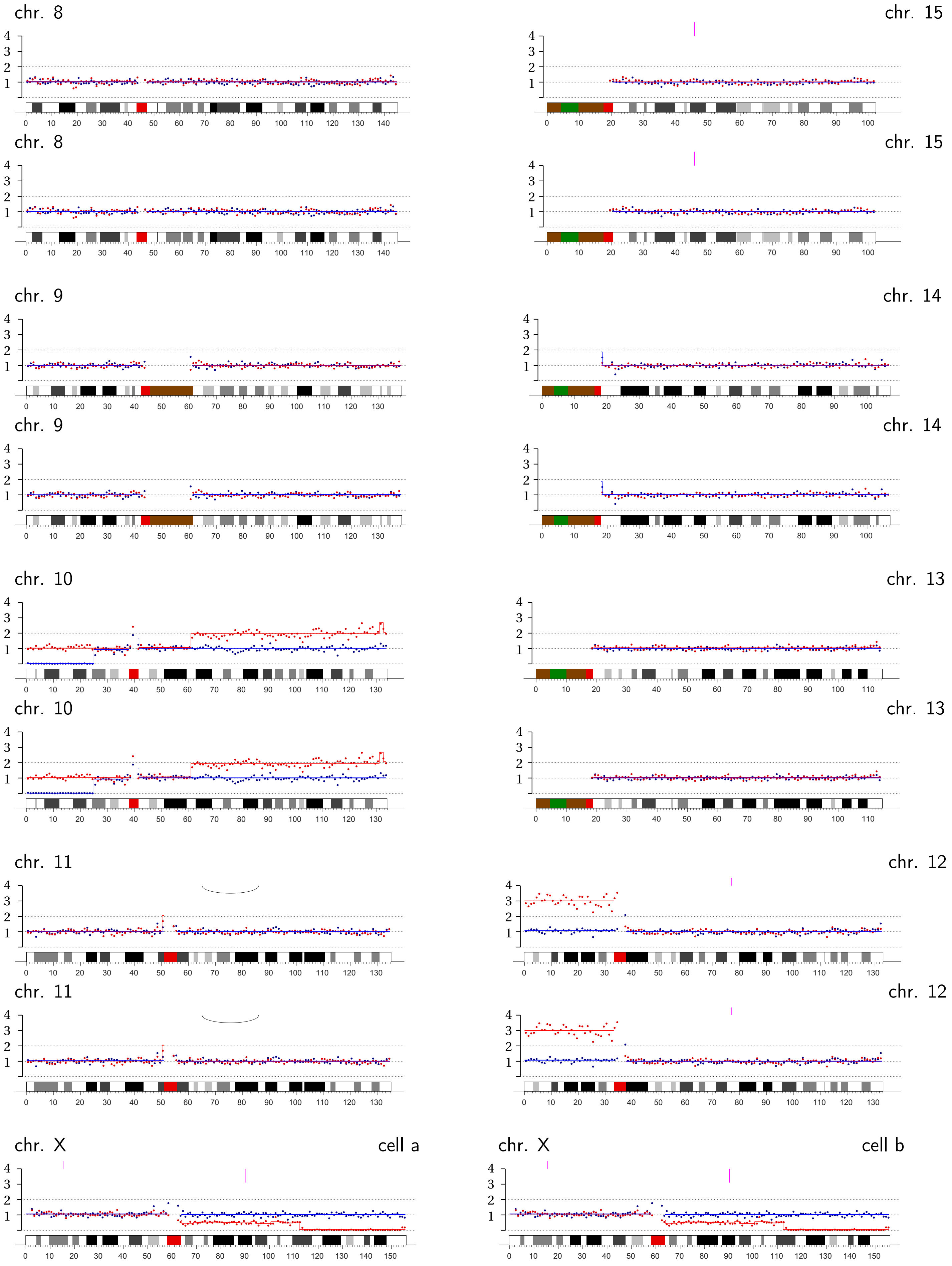
chr. 8
chr. 15
chr. 8
chr. 15
chr. 9
chr. 14
chr. 9
chr. 14
chr. 10
chr. 13
chr. 10
chr. 13
chr. 11
chr. 12
chr. 11
chr. 12
chr. X
cell a
chr. X
cell b

**Bridge Pair TREX1-6 |** Transient TRF2-DN expression + TREX1 knockout: Chromosome 1B, p-arm internal; 5A, q-ter.; 16A, q-ter.

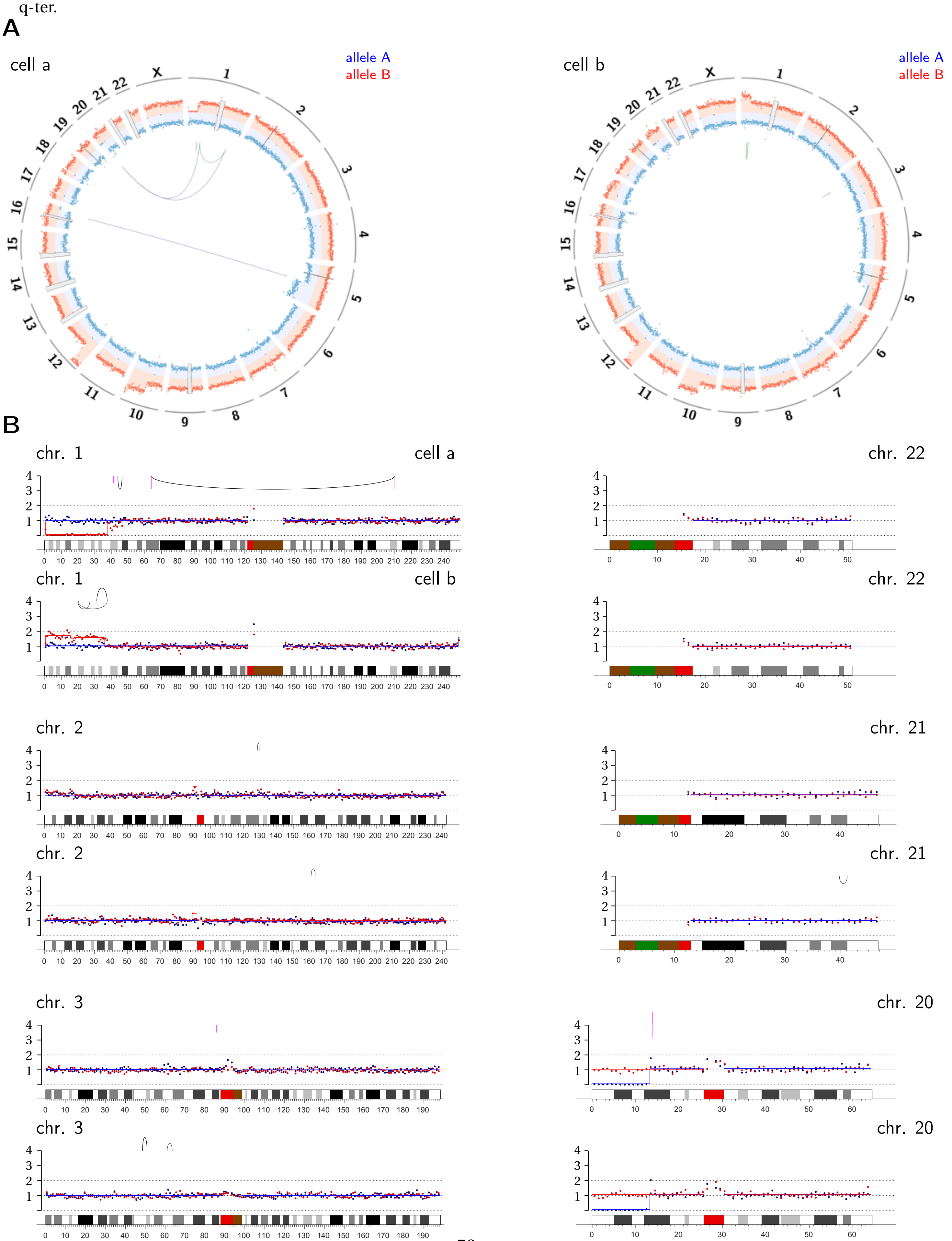

chr. 4

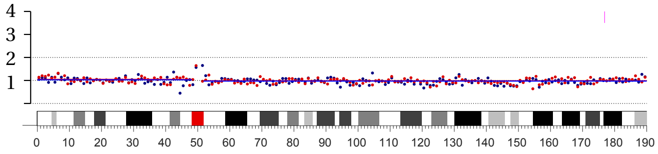

chr. 4

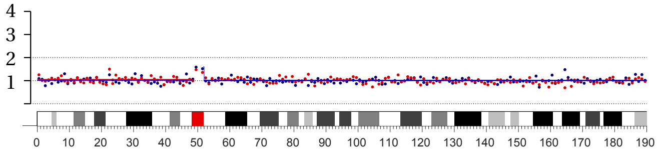

chr. 5

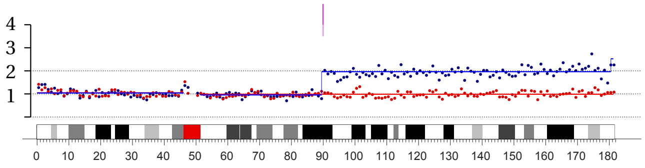

chr. 5

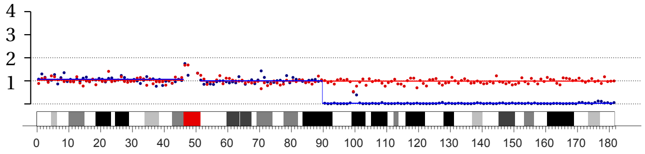

chr. 6

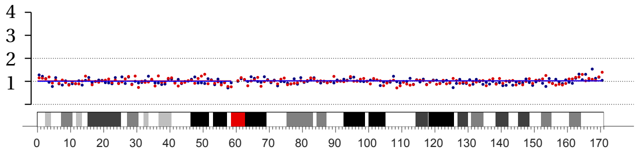

chr. 6

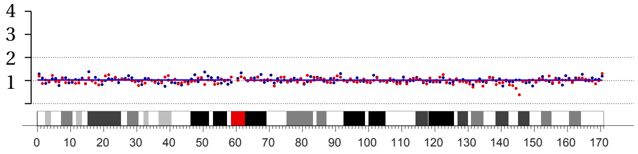

chr. 7

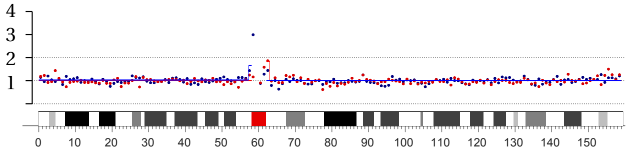

chr. 7

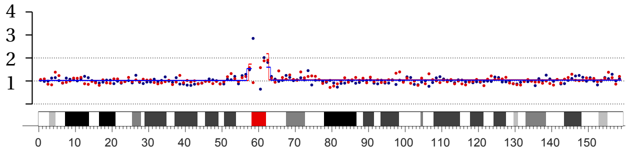

chr. 19

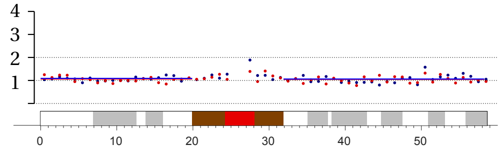

chr. 19

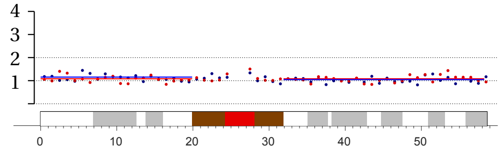

chr. 18

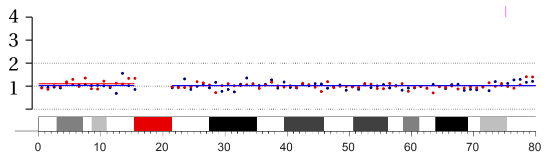

chr. 18

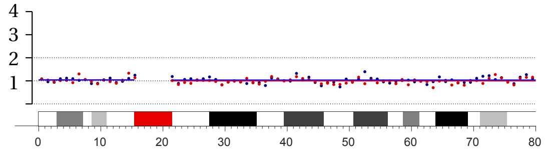

chr. 17

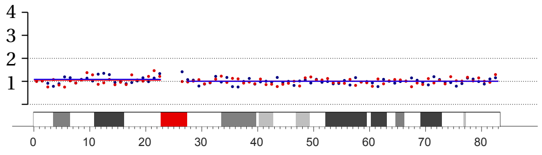

chr. 17

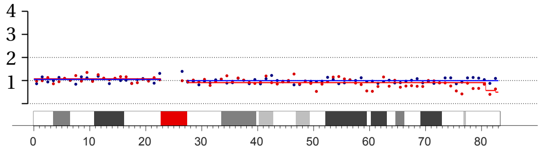

chr. 16

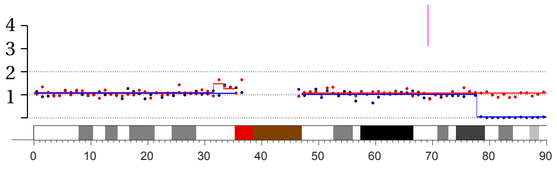

chr. 16

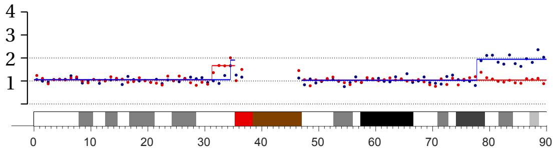

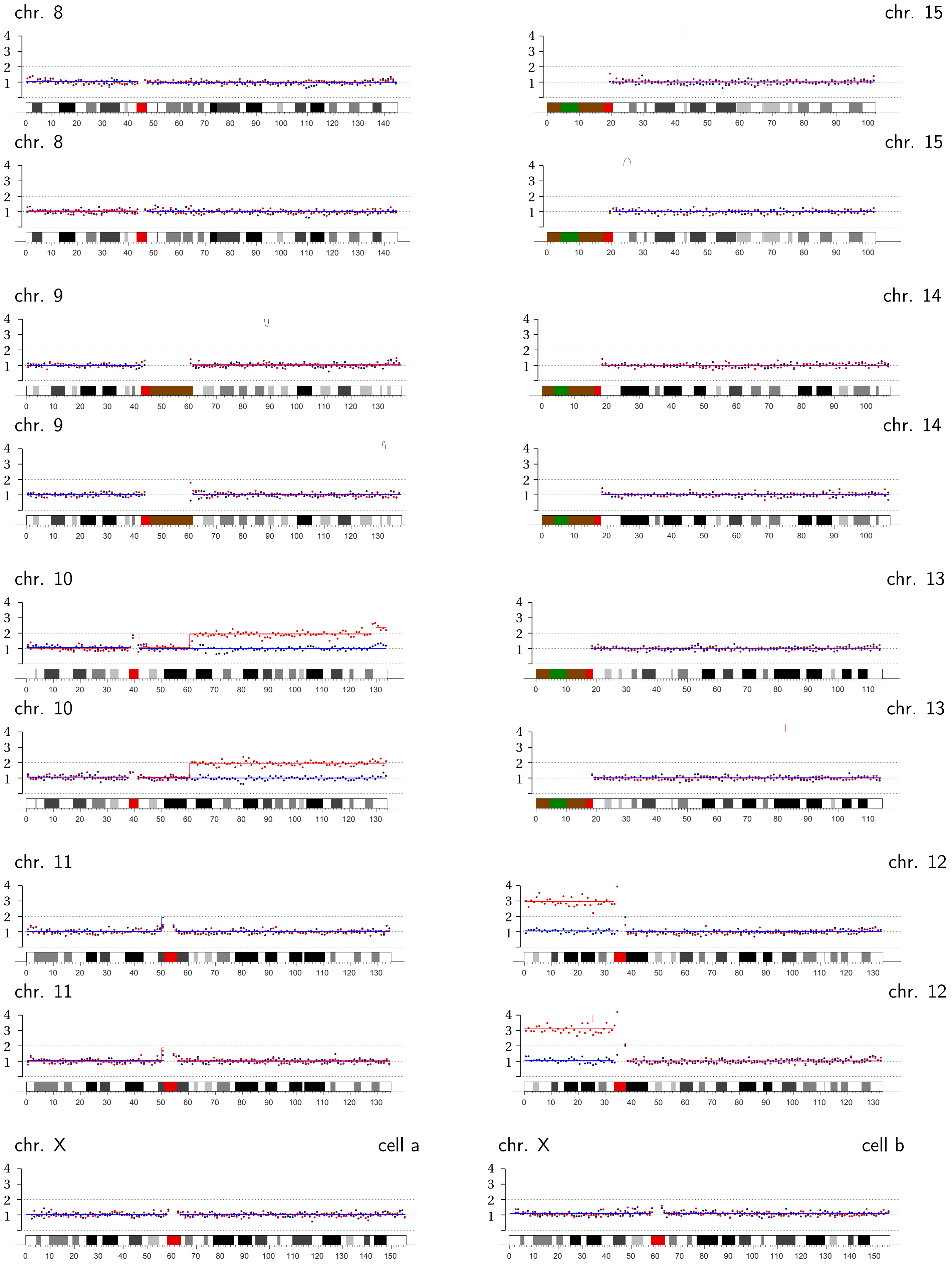
chr. 8
chr. 15
chr. 8
chr. 15
chr. 9
chr. 14
chr. 9
chr. 14
chr. 10
chr. 13
chr. 10
chr. 13
chr. 11
chr. 12
chr. 11
chr. 12
chr. X
cell a
chr. X
cell b

**Bridge Pair TREX1-7 |** Transient TRF2-DN expression + TREX1 knockout: Chromosomes 4A and 4B, p-ter.; 11B, q-ter.; 17B, p-ter.

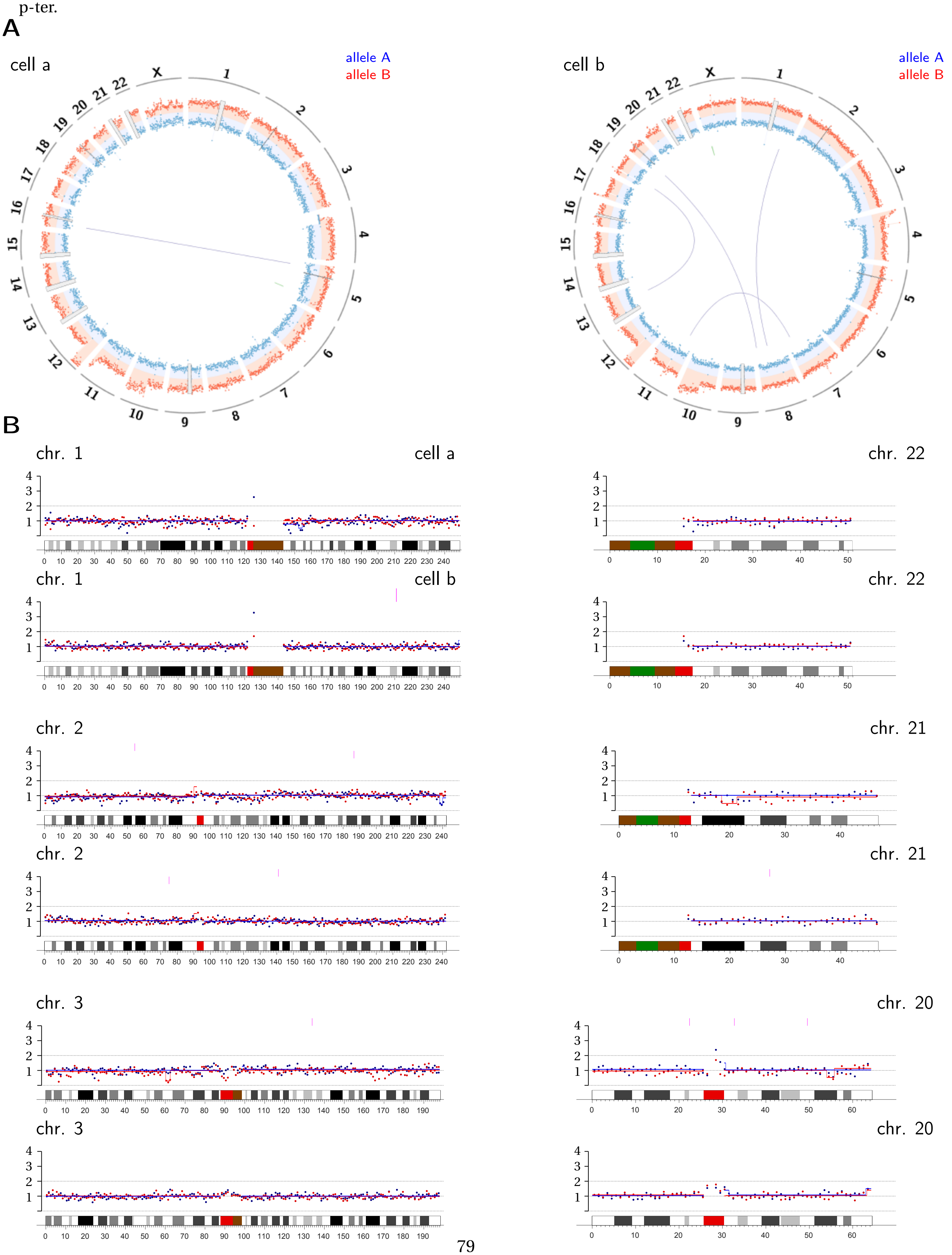

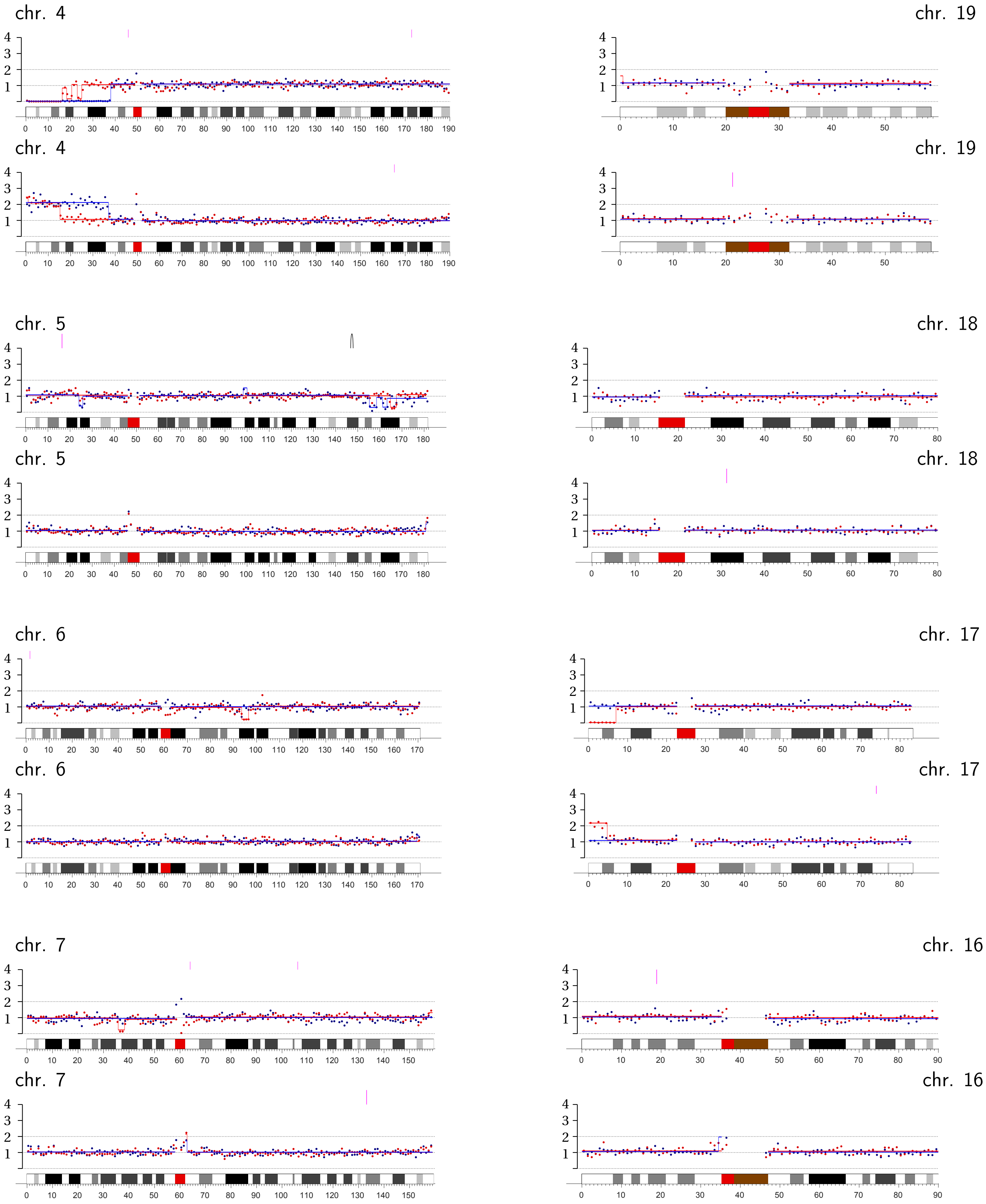
chr. 4
chr. 19
chr. 4
chr. 19
chr. 5
chr. 18
chr. 5
chr. 18
chr. 6
chr. 17
chr. 6
chr. 17
chr. 7
chr. 16
chr. 7
chr. 16

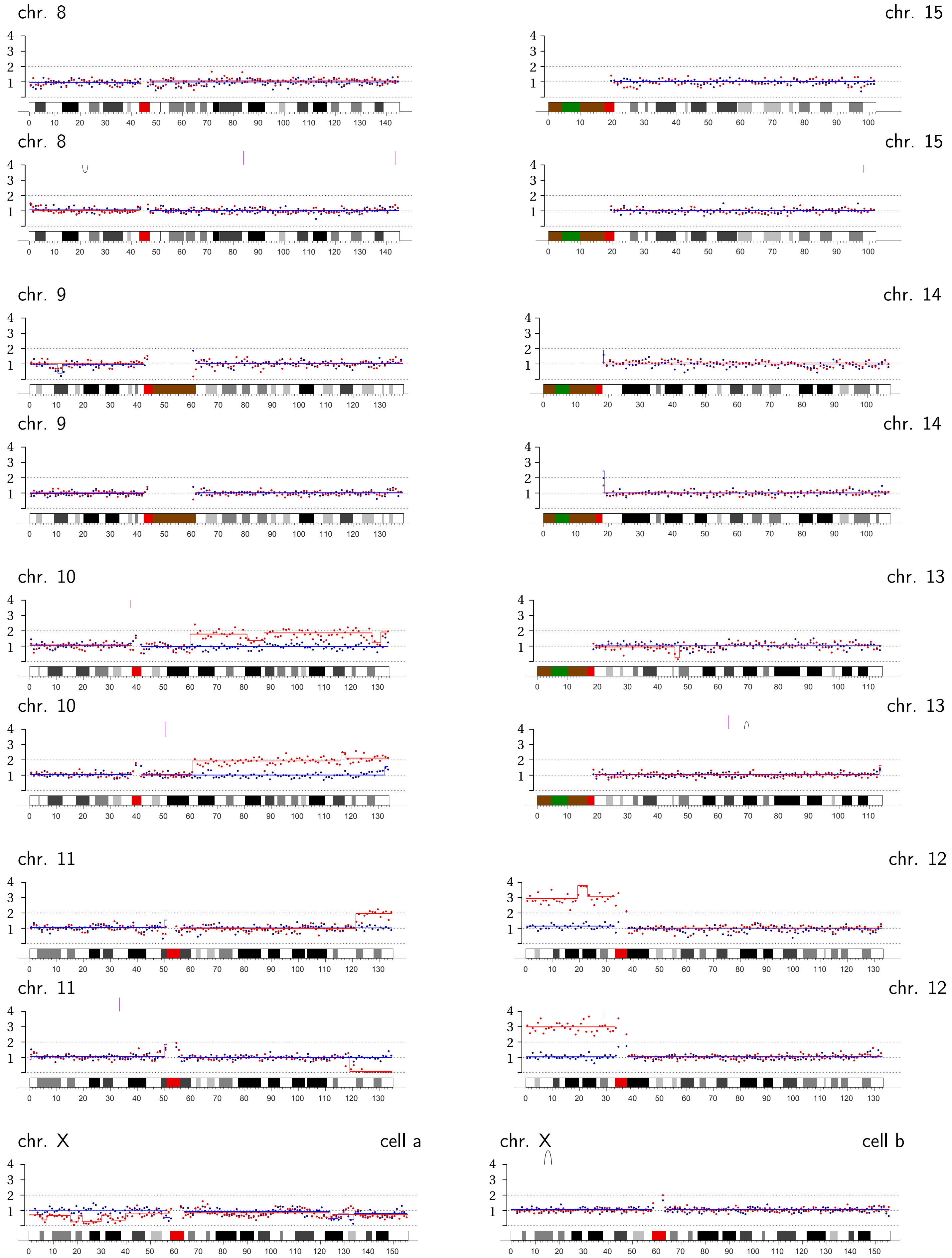
chr. 8
chr. 15
chr. 8
chr. 15
chr. 9
chr. 14
chr. 9
chr. 14
chr. 10
chr. 13
chr. 10
chr. 13
chr. 11
chr. 12
chr. 11
chr. 12
chr. X
cell a
chr. X
cell b

**Bridge Pair TREX1-8 |** Transient TRF2-DN expression + TREX1 knockout: Chromosome 3B, p-ter. with pre-existing terminal loss; 19B, p-arm.

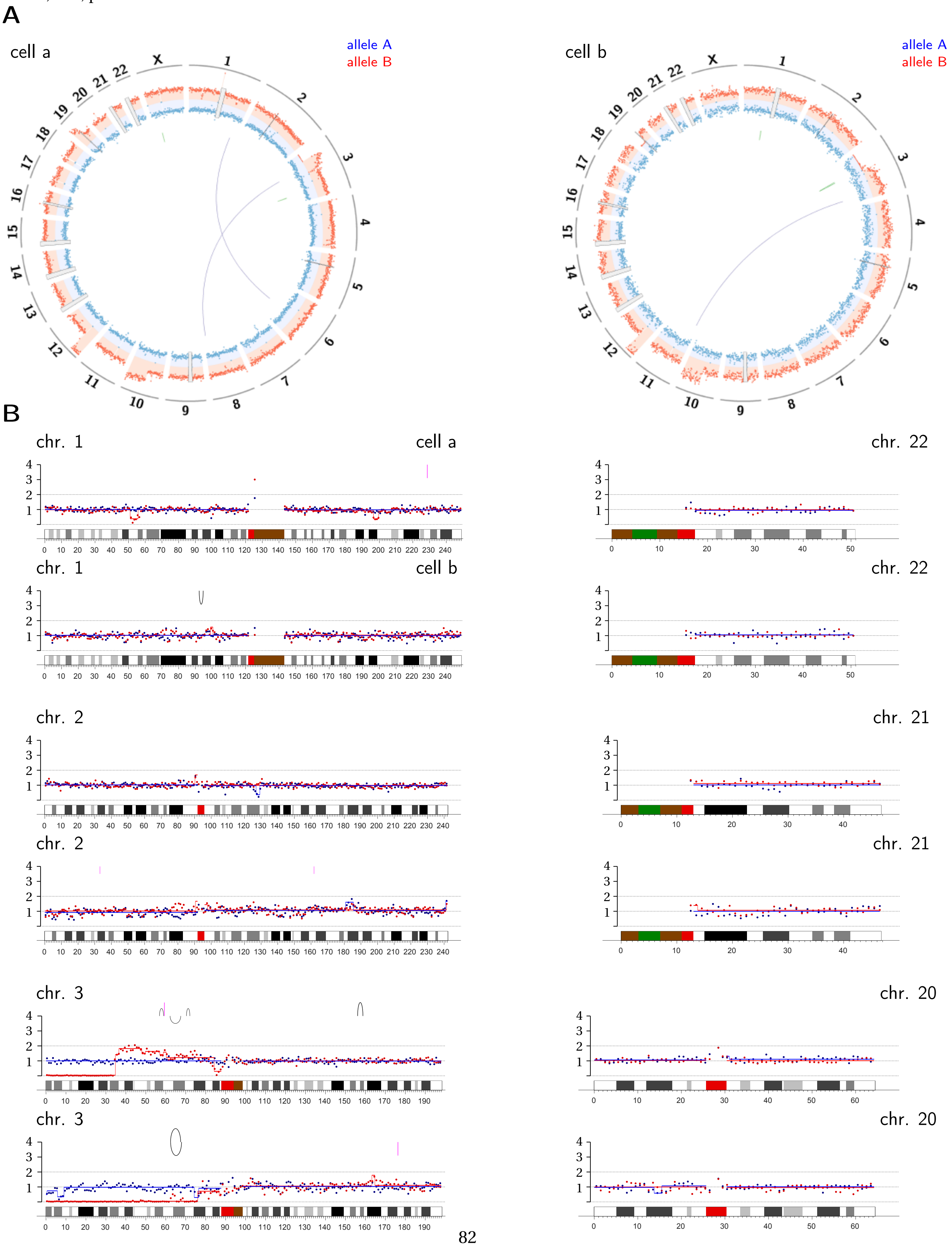

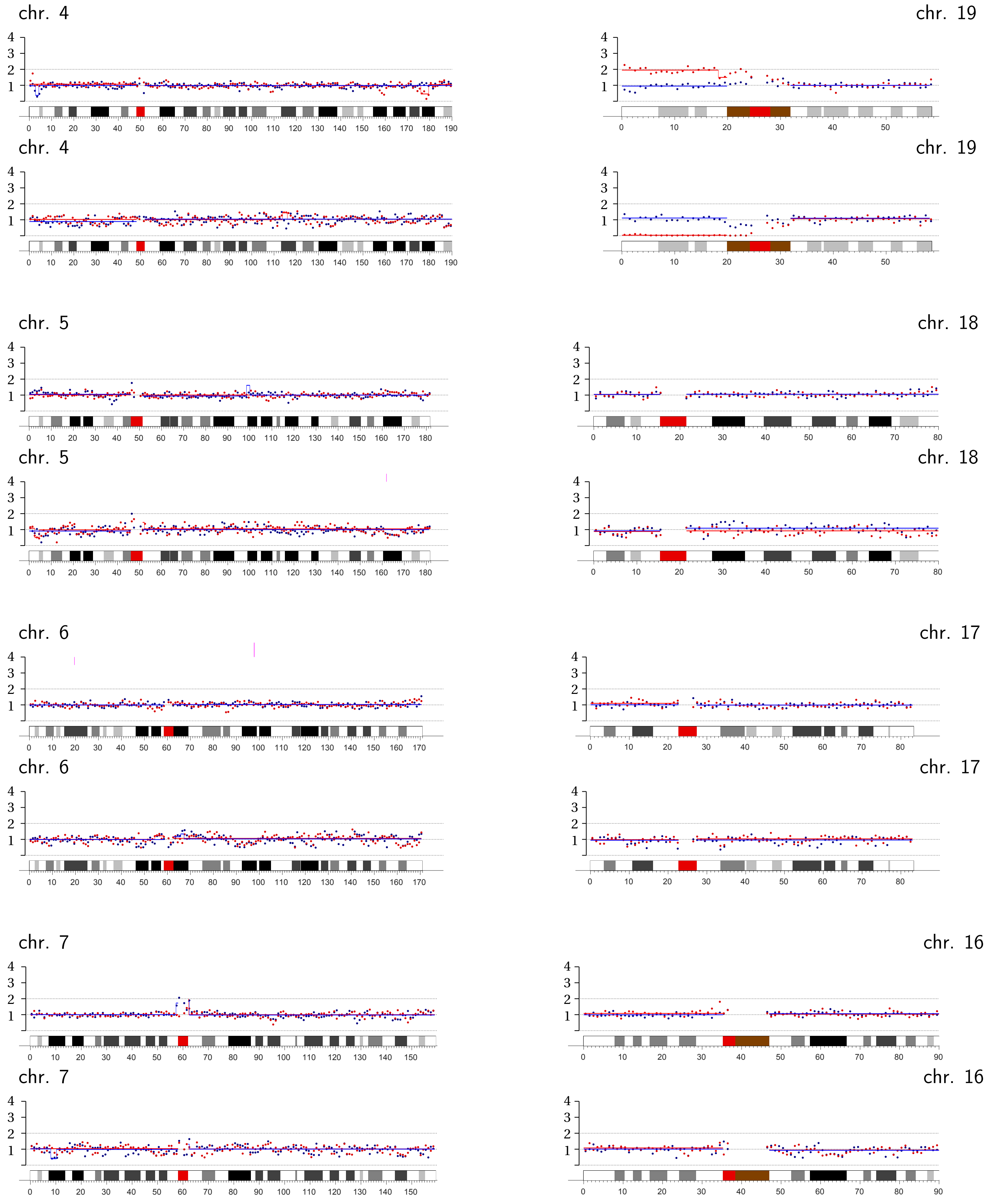
chr. 4
chr. 19
chr. 4
chr. 19
chr. 5
chr. 18
chr. 5
chr. 18
chr. 6
chr. 17
chr. 6
chr. 17
chr. 7
chr. 16
chr. 7
chr. 16

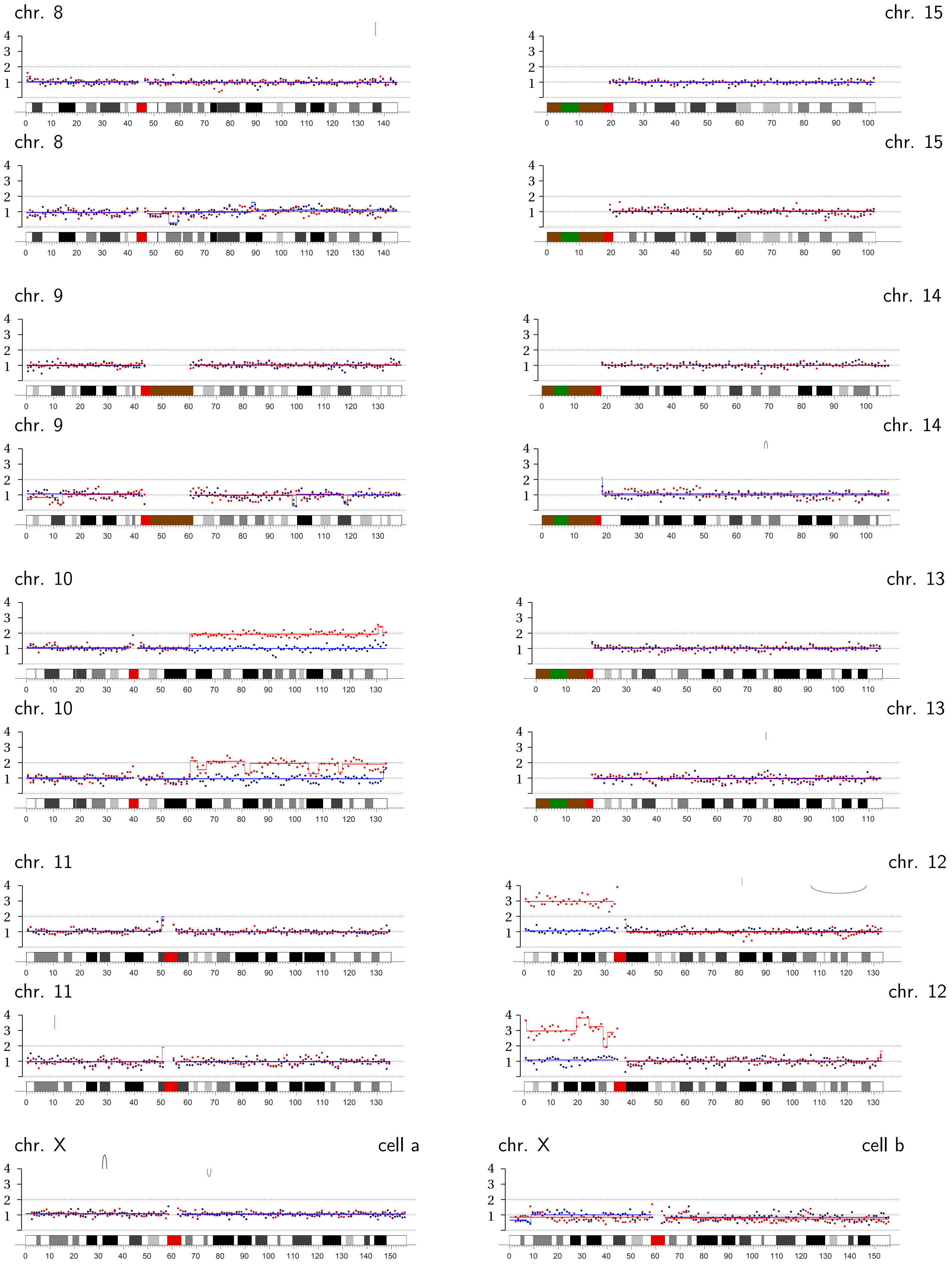
chr. 8
chr. 15
chr. 8
chr. 15
chr. 9
chr. 14
chr. 9
chr. 14
chr. 10
chr. 13
chr. 10
chr. 13
chr. 11
chr. 12
chr. 11
chr. 12
chr. X
cell a
chr. X
cell b

### Figure S8

**A** Mechanical breakage (bridges induced by TRF2-DN)

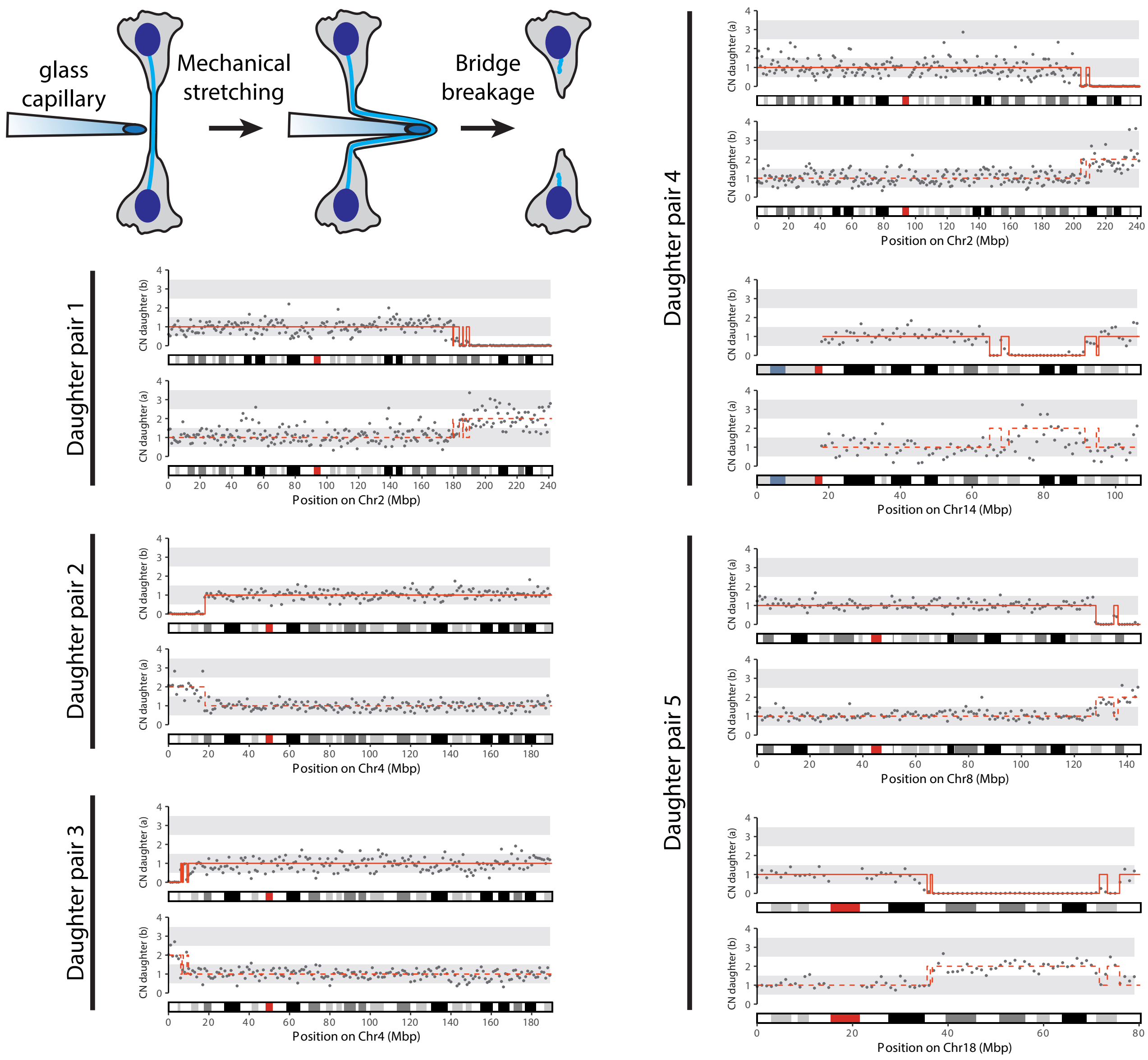


**B** ΔTREX1 (bridges induced by TRF2-DN)

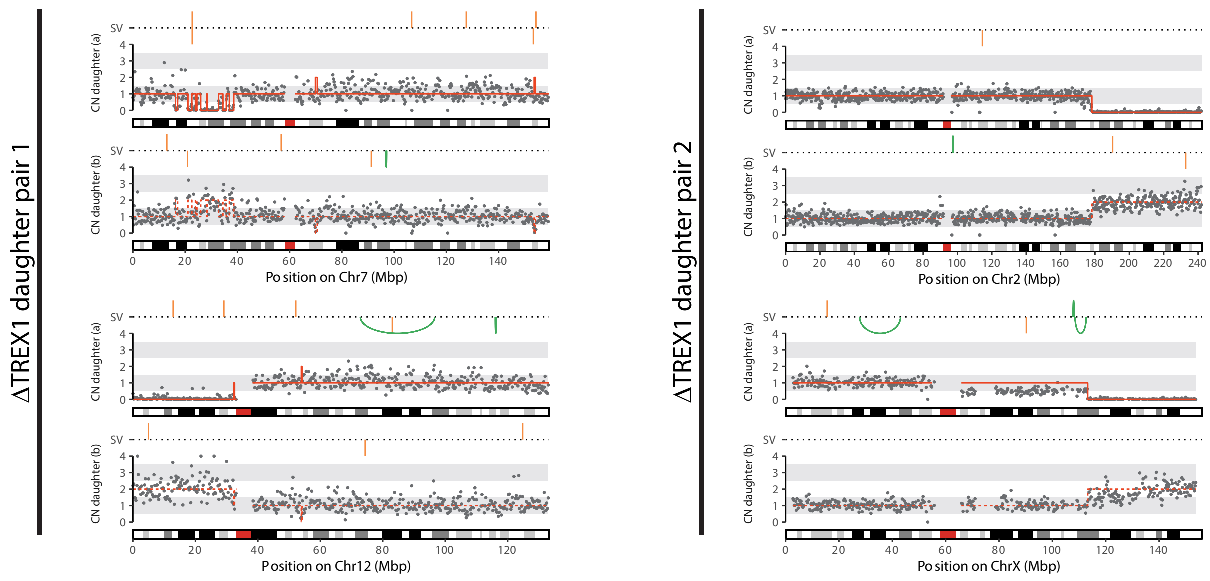

### Figure S8

**B**

ΔTREX1 (bridges induced by TRF2-DN)

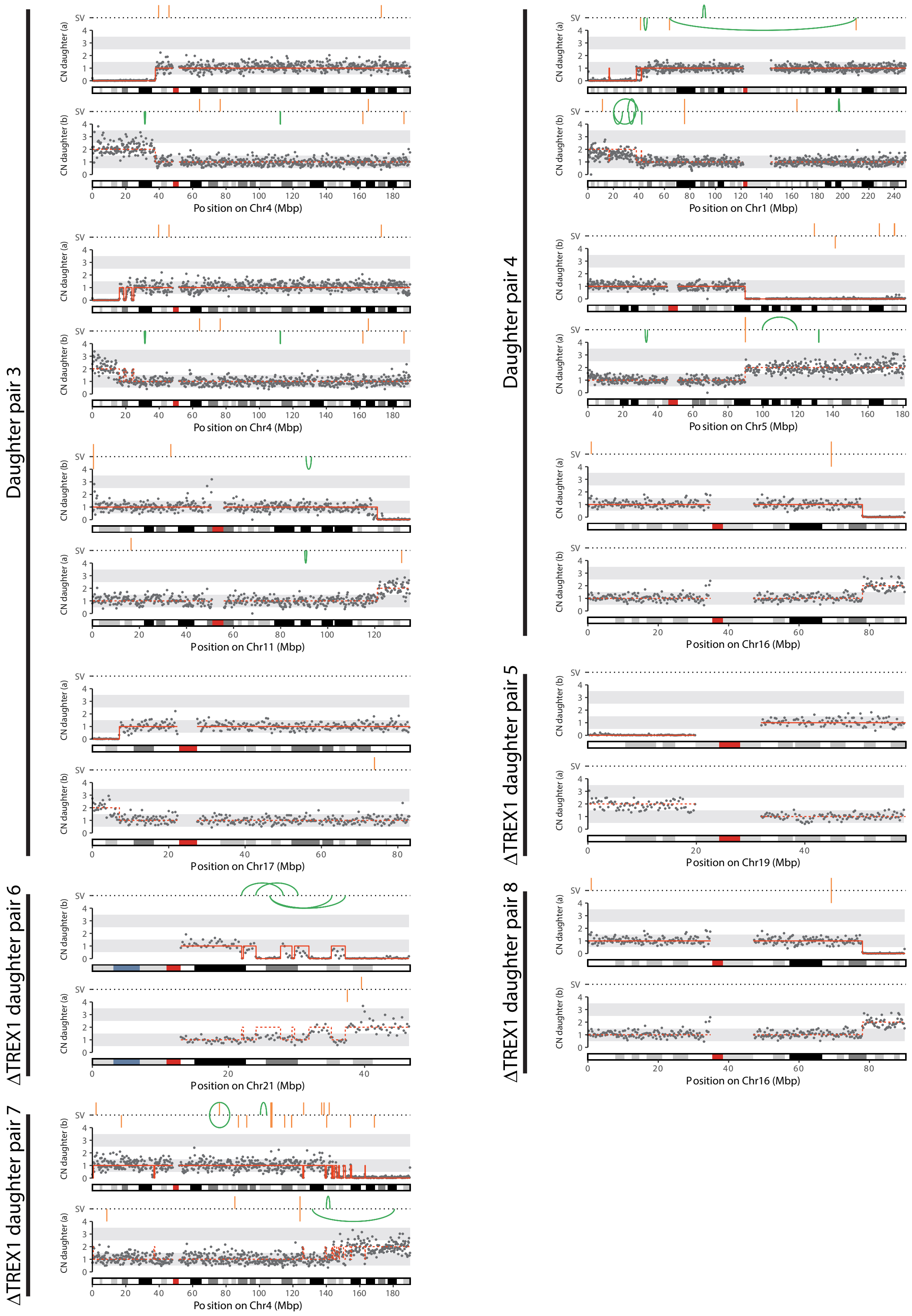

Supplemental Figure 9

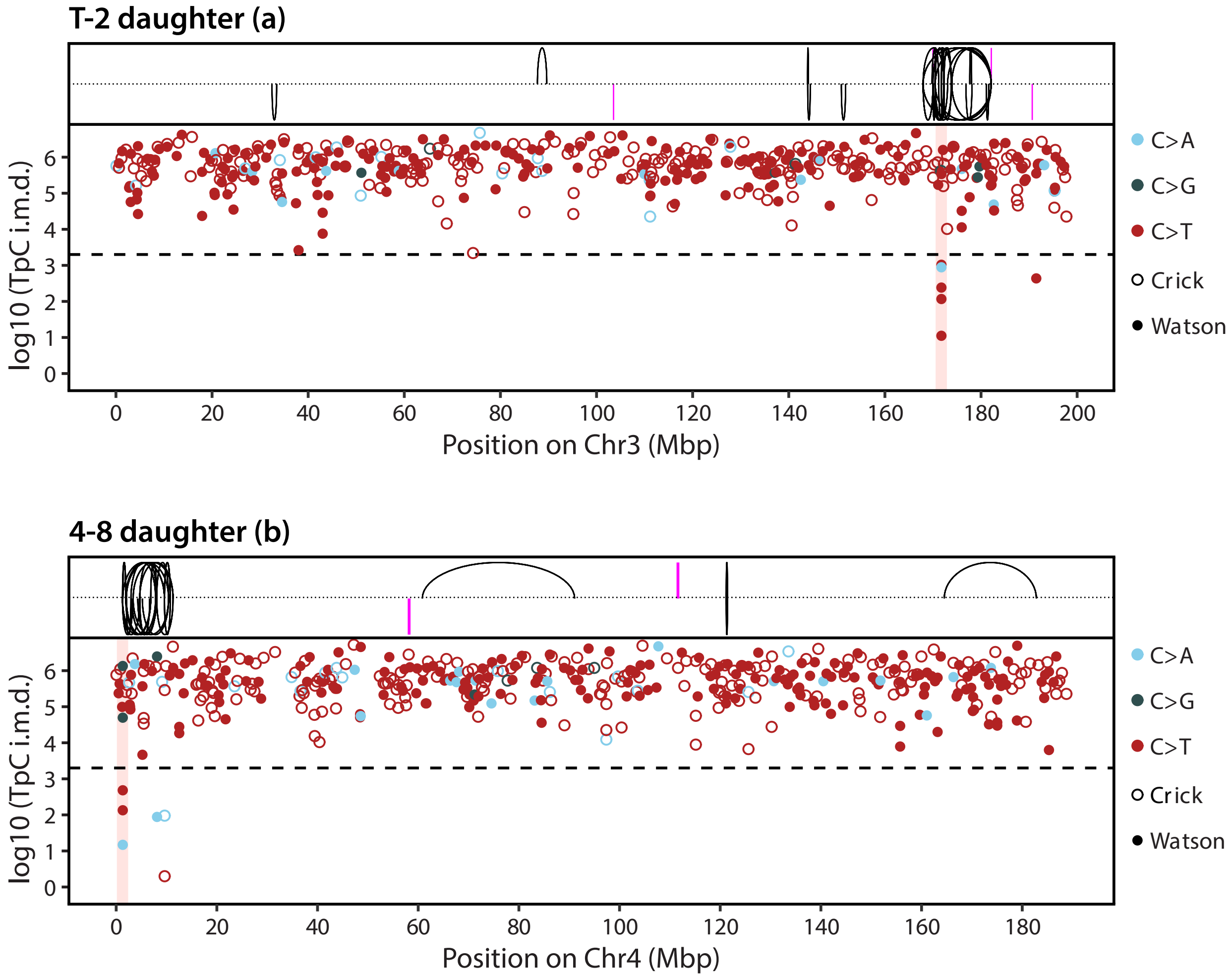

### Supplemental Figure 10

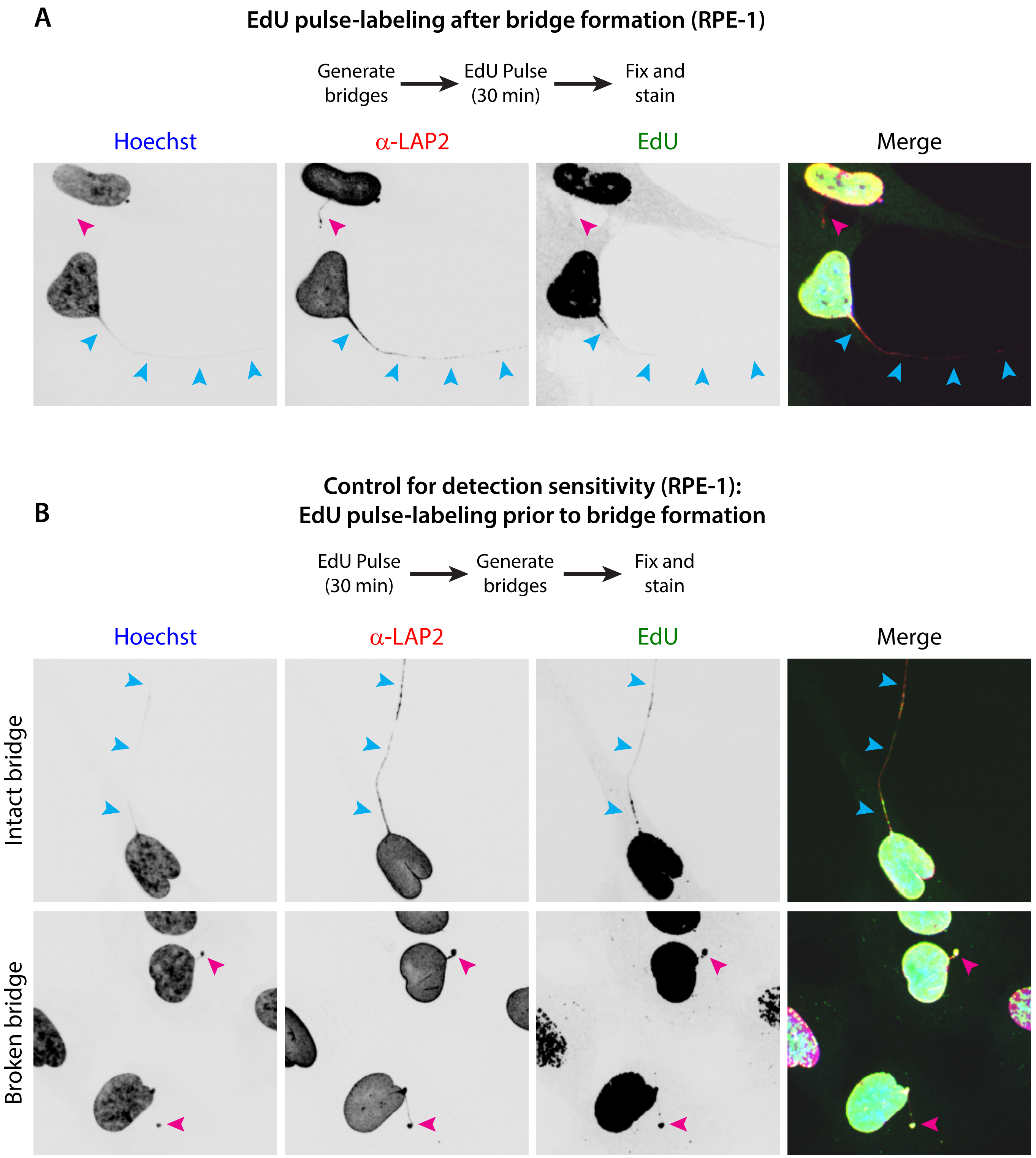

Supplemental Figure 11

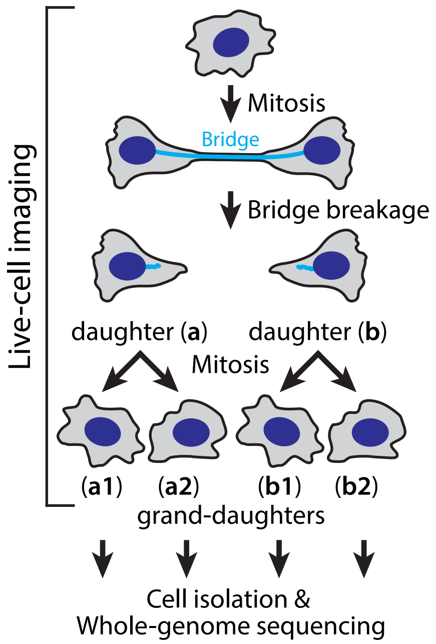


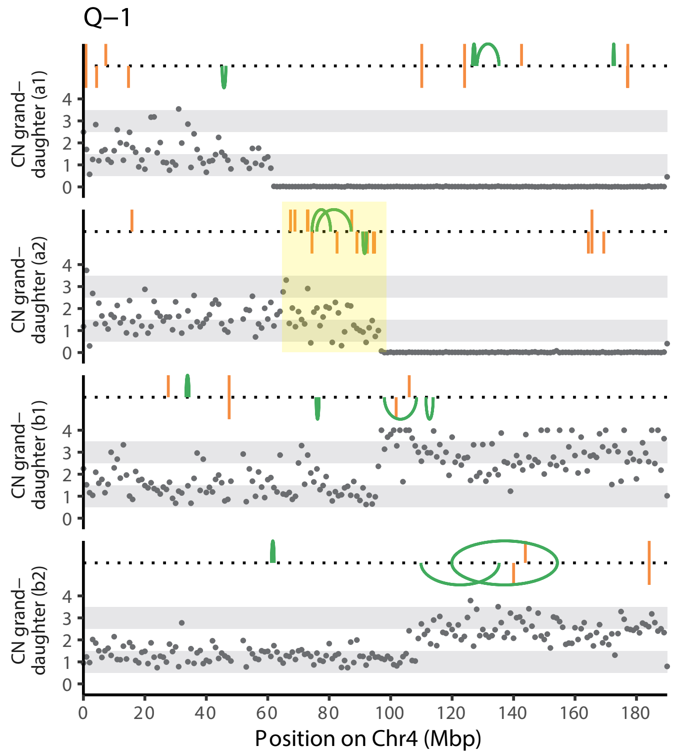


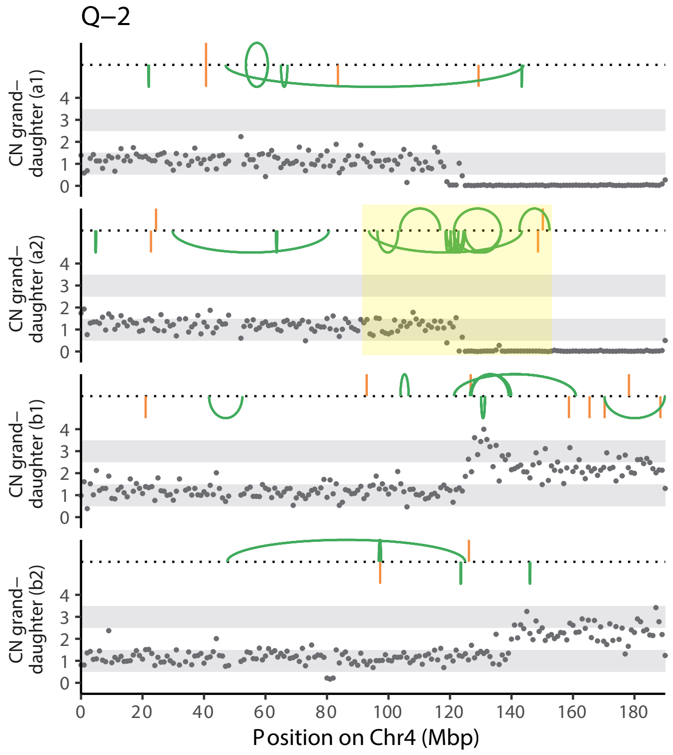


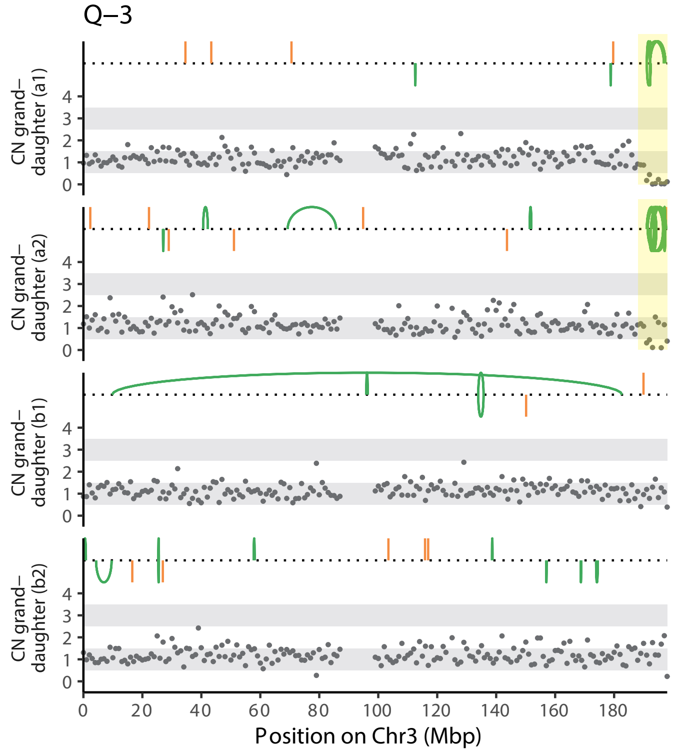

### Supplemental Figure 12

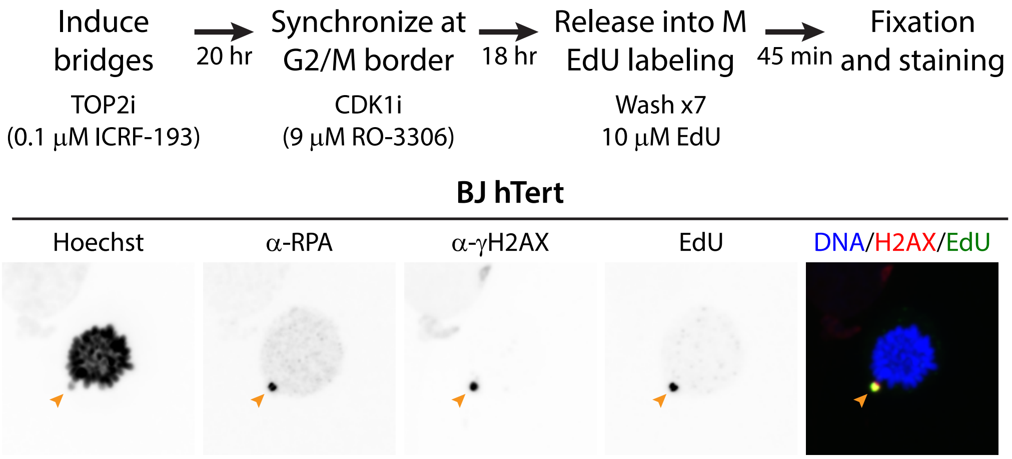

### Supplemental Figure 13

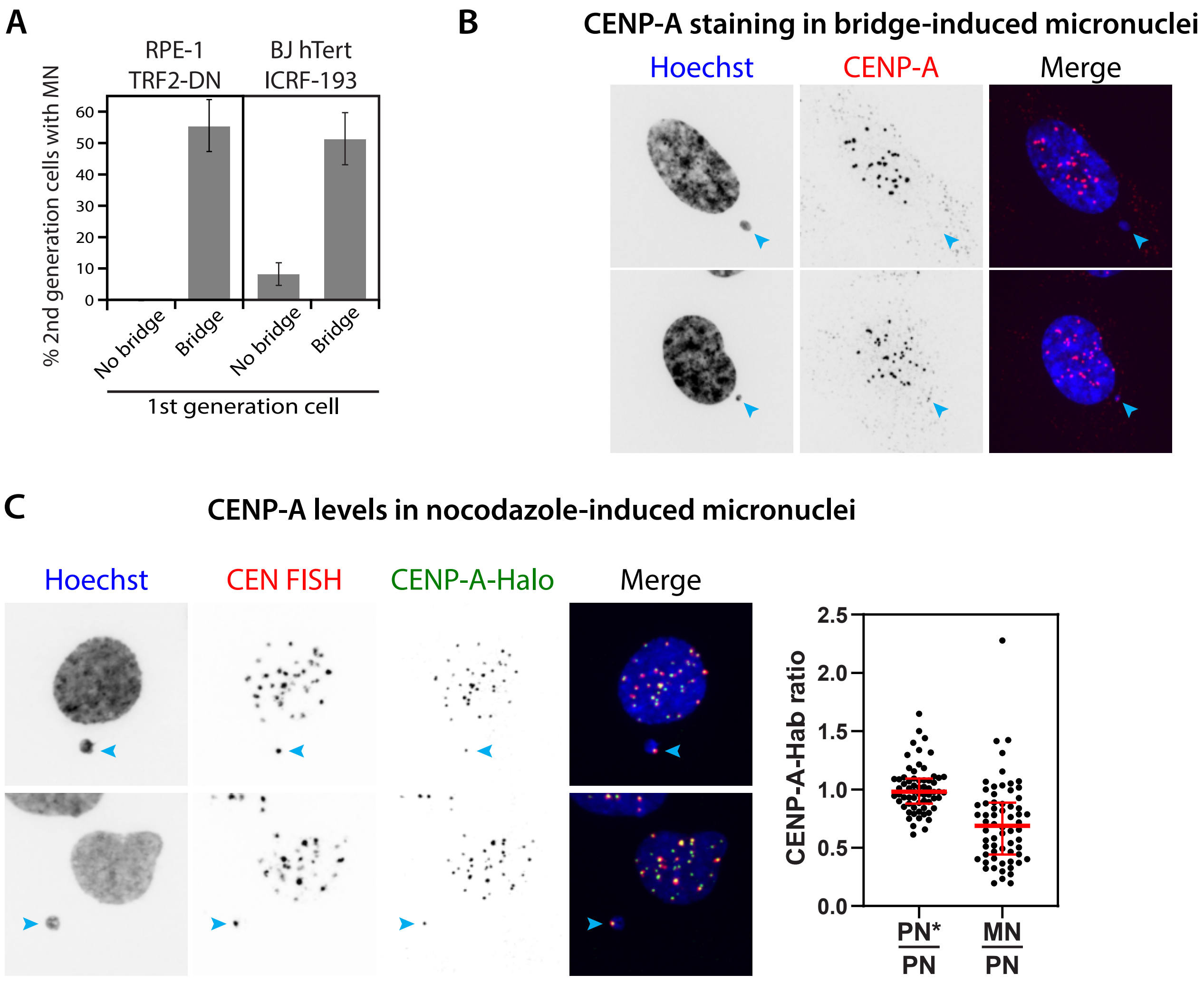

### Supplemental Figure 14

A

Mitotic replication on micronuclear chromosomes

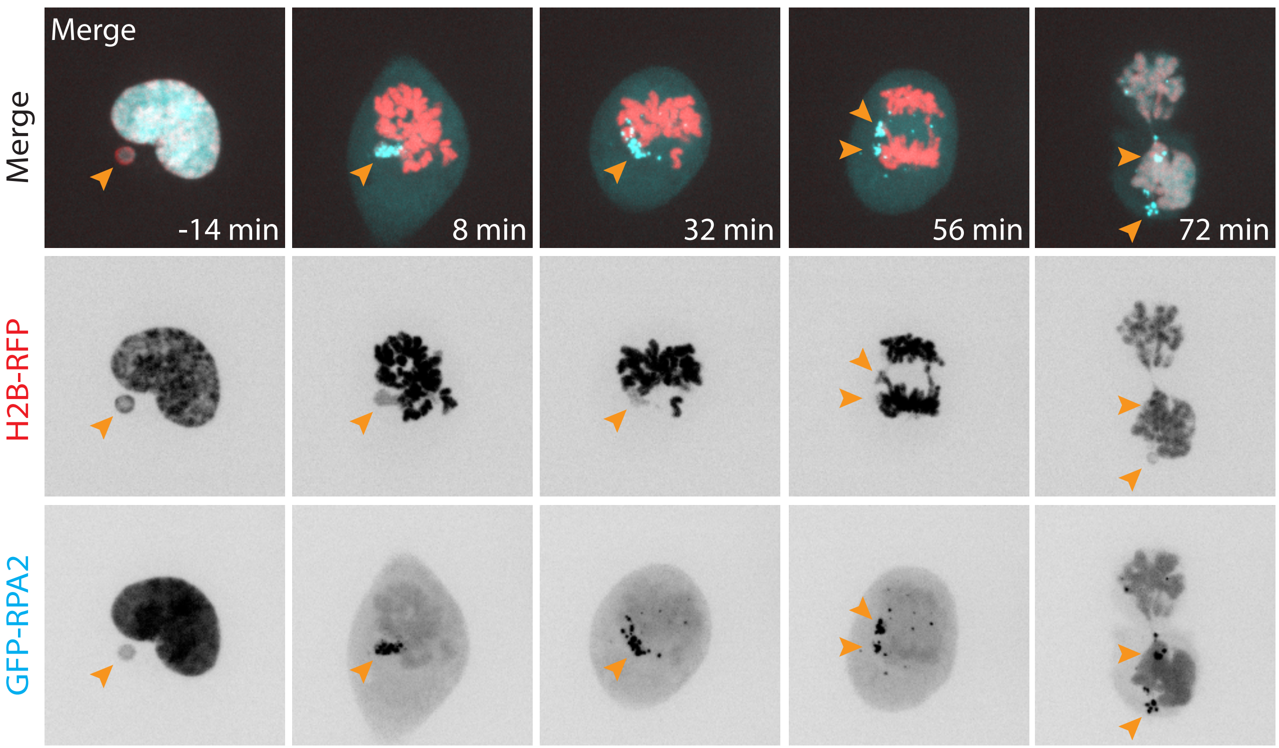


B

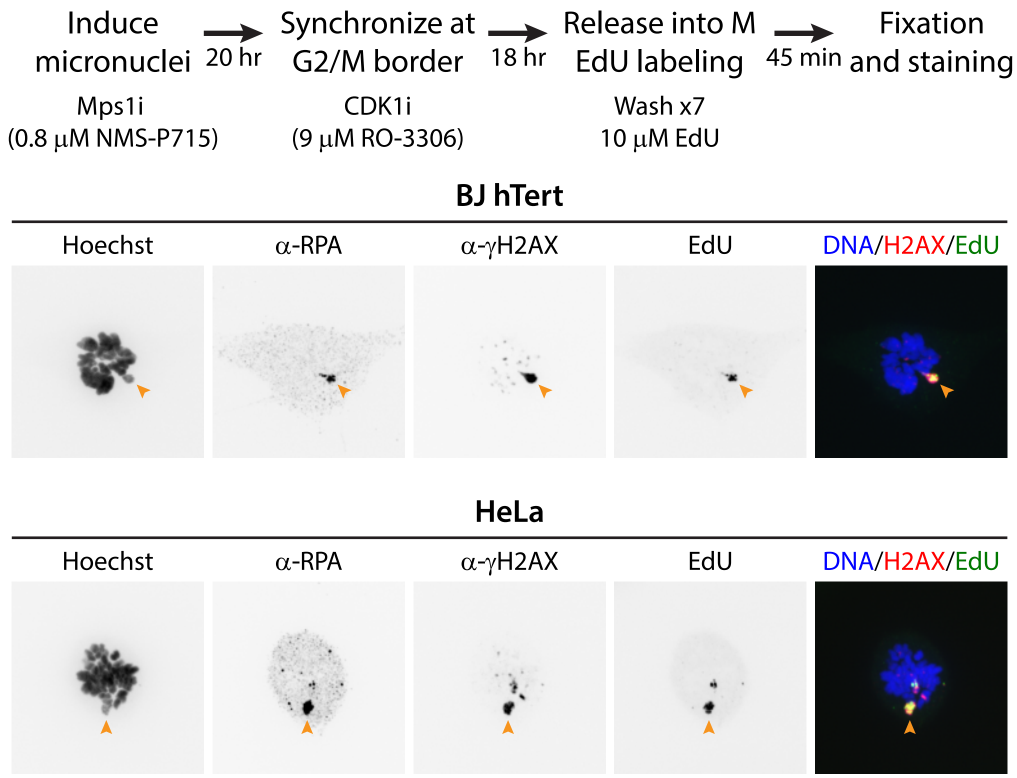

#### Supplemental Figure 15

Chromosomes from intact micronuclei undergo chromothripsis after passing through the next mitosis (all examples from this study).


MN1 **Chromothripsis/fragmentation on chromosomes 10A (1:0) and 21B (2:1)**. . . . . . . . . . . . . . . . . . . . 1

MN2 Asymmetric copy number of chromosome 8B (1:0). . . . . . . . . . . . . . . . . . . . . . . . . . . . . . . . . . . . . . 4

MN3 **Chromothripsis/fragmentation on both chromosome 2 homologs: 2A (2:1) and 2B (1:0)**. . . . . . . . . . . 7

MN4 Asymmetric copy number of chromosome 2B (1:0) with partial replication and fragmentation. . . . . . . . 10

MN5 **Chromothripsis/fragmentation on chromosome 2B (1:0) with partial replication**. . . . . . . . . . . . . . 13

MN6 **Chromothripsis/fragmentation on chromosome 1B (1:0)**. . . . . . . . . . . . . . . . . . . . . . . . . . . . . . 16

MN7 **Chromothripsis/fragmentation on chromosome 11B (1:0)**; daughter b likely in G2 with un-replicated Chr. 11 in persistent MN. . . . . . . . . . . . . . . . . . . . . . . . . . . . . . . . . . . . . . . . . . . . . . . . . . . . 19

MN8 **Complex rearrangements involving chromosomes 1B p-arm (2:1), 10A (2:1), and 18B(2:1)**. . . . . . . . . 22

MN9 **Chromothripsis on chromosome 1A (1:0)**; daughter b likely in G2 with Chr. 1A in persistent MN. . . . . . 25

MN10 **Complex rearrangement on chromosome 1B q-arm (2:0)**. . . . . . . . . . . . . . . . . . . . . . . . . . . . . . 28

**Micronucleus Pair MN1 | Chromothripsis/fragmentation on chromosomes 10A (1:0) and 21B (2:1).**

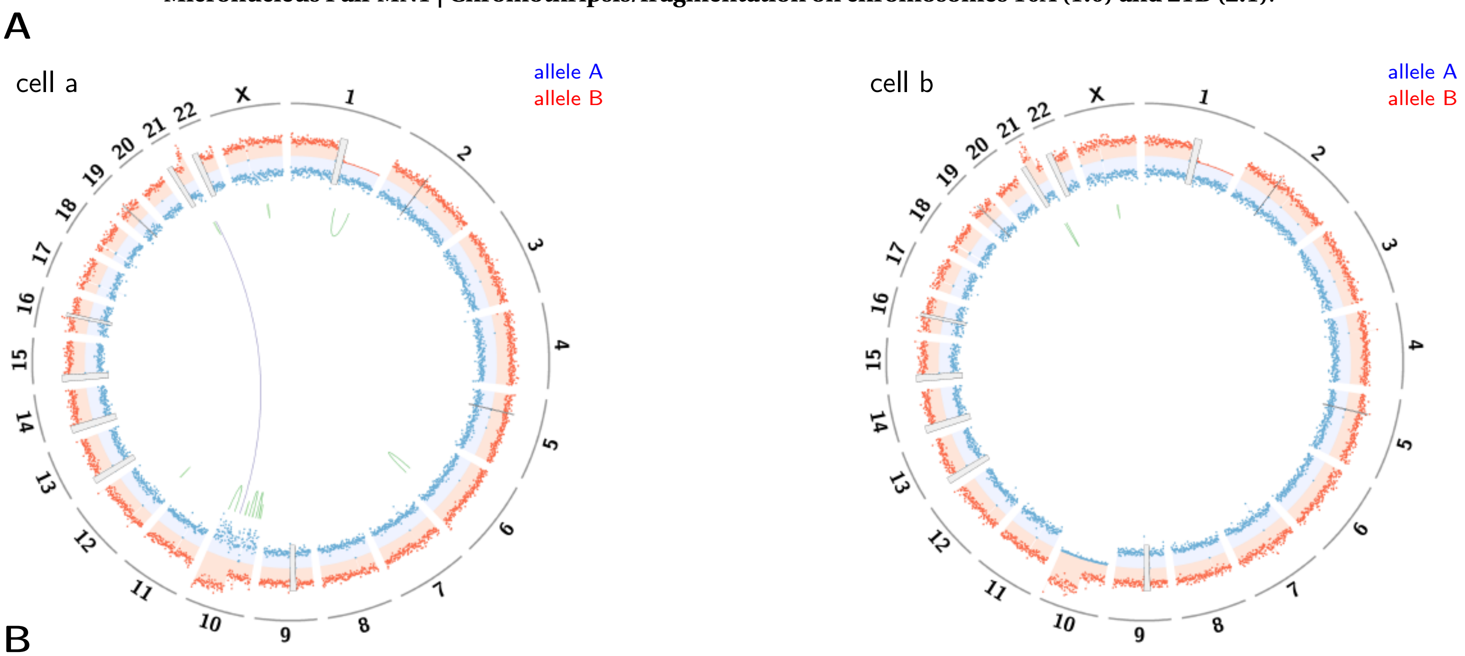


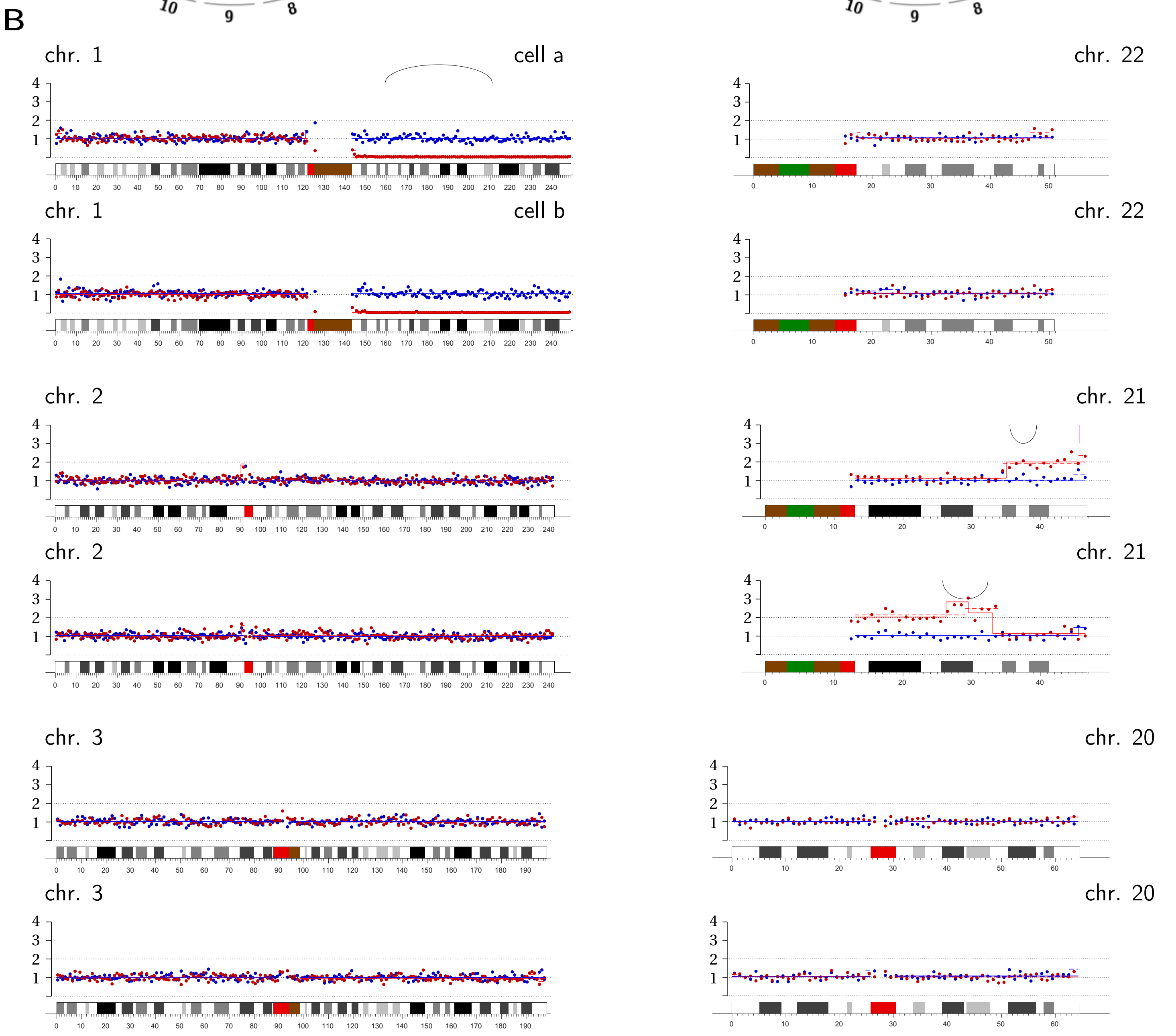

chr. 4

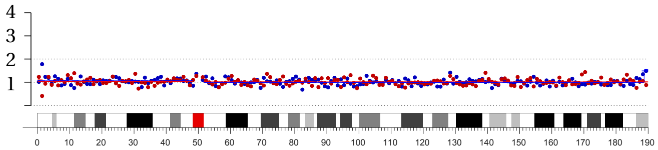


chr. 4

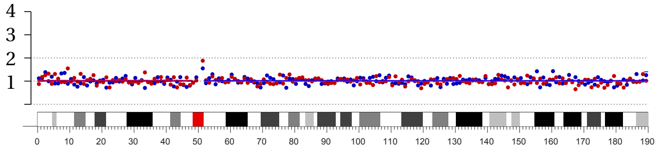


chr. 19

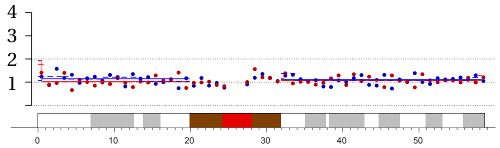


chr. 19

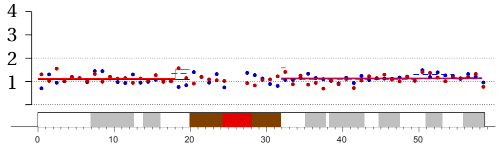


chr. 5

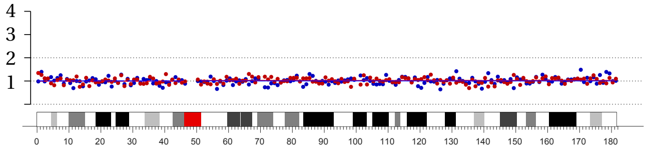


chr. 5

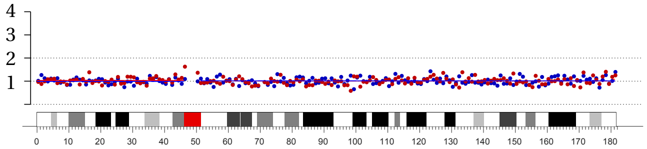


chr. 18

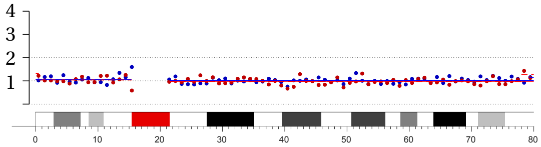


chr. 18

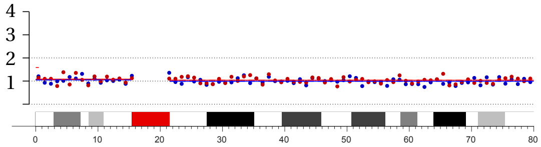


chr. 6

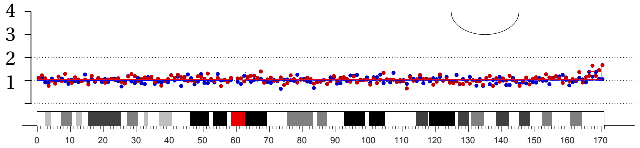


chr. 6

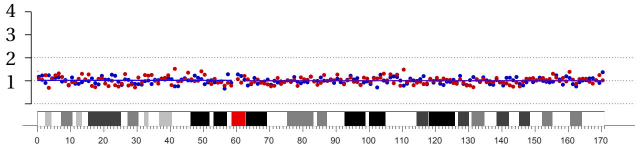


chr. 17

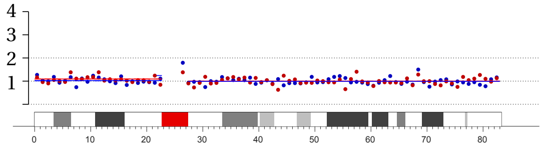


chr. 17

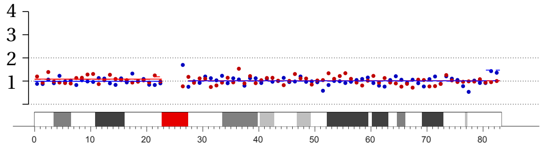


chr. 7

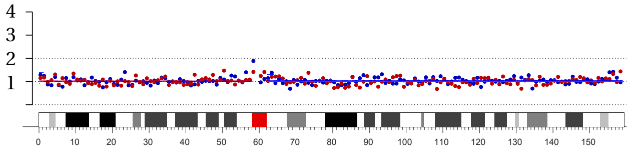


chr. 7

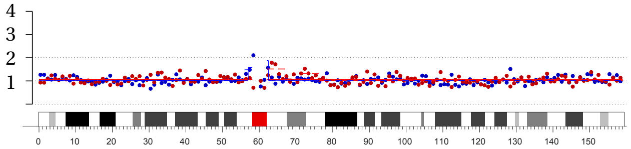


chr. 16

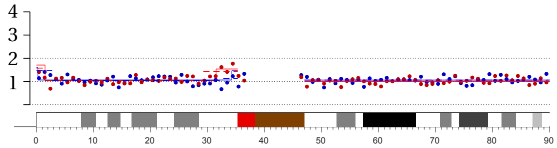


chr. 16

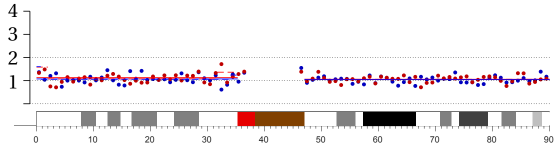

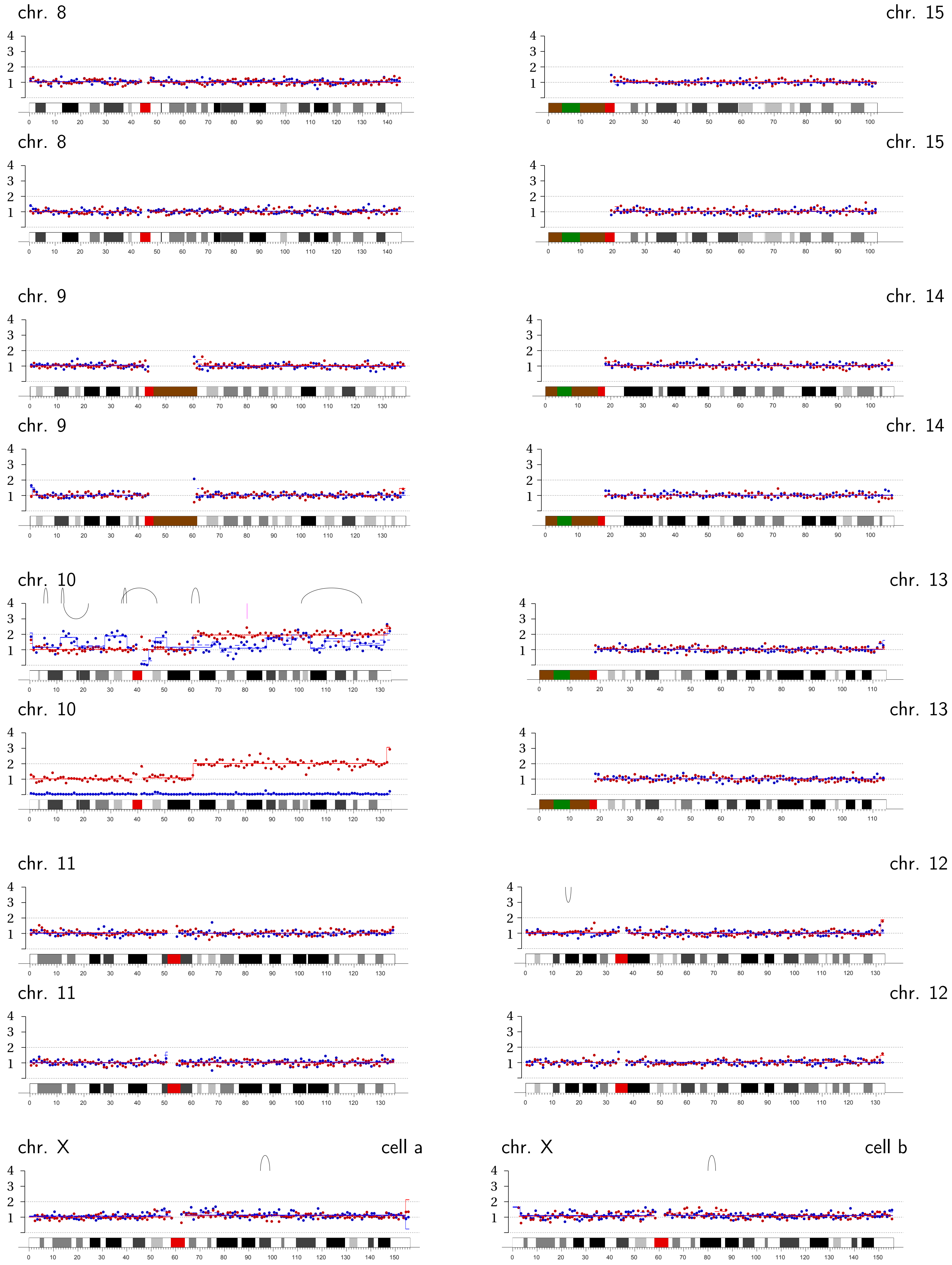
chr. 8
chr. 15
chr. 8
chr. 15
chr. 9
chr. 14
chr. 9
chr. 14
chr. 10
chr. 13
chr. 10
chr. 13
chr. 11
chr. 12
chr. 11
chr. 12
chr. X
cell a
chr. X
cell b

**Micronucleus Pair MN2 |** Asymmetric copy number of chromosome 8B (1:0).

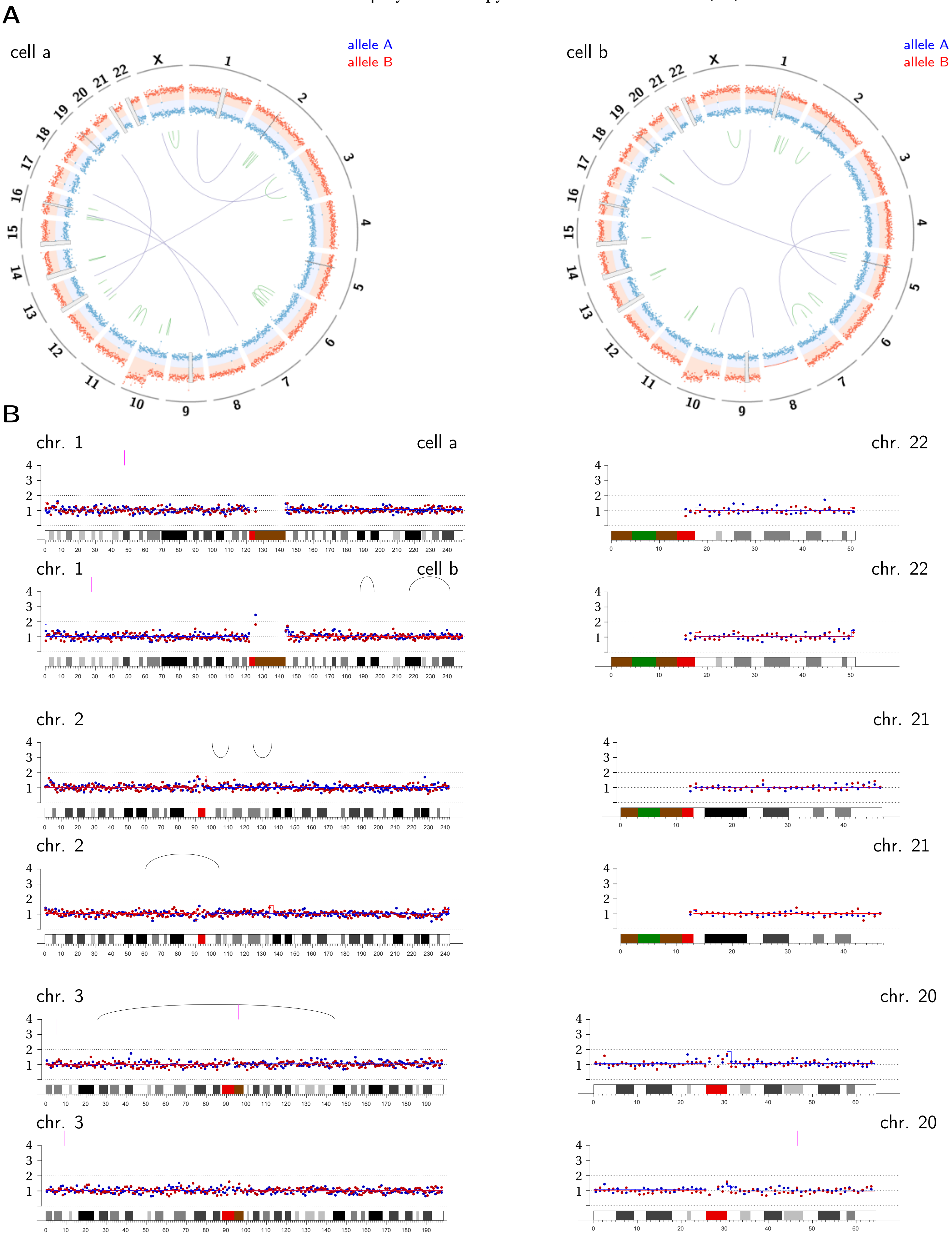

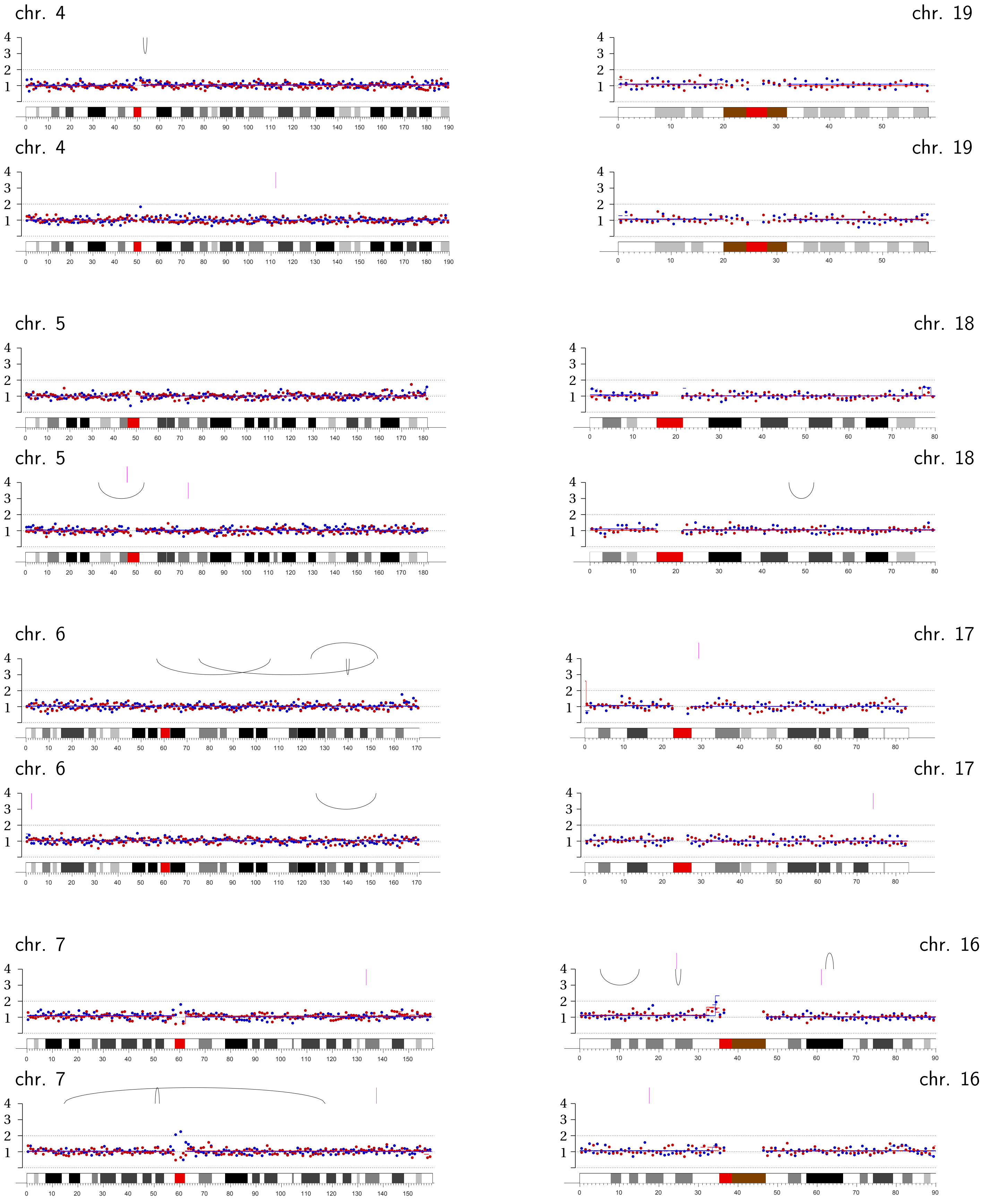
chr. 4
chr. 4
chr. 19
chr. 19
chr. 5
chr. 5
chr. 18
chr. 18
chr. 6
chr. 6
chr. 17
chr. 17
chr. 7
chr. 7
chr. 16
chr. 16

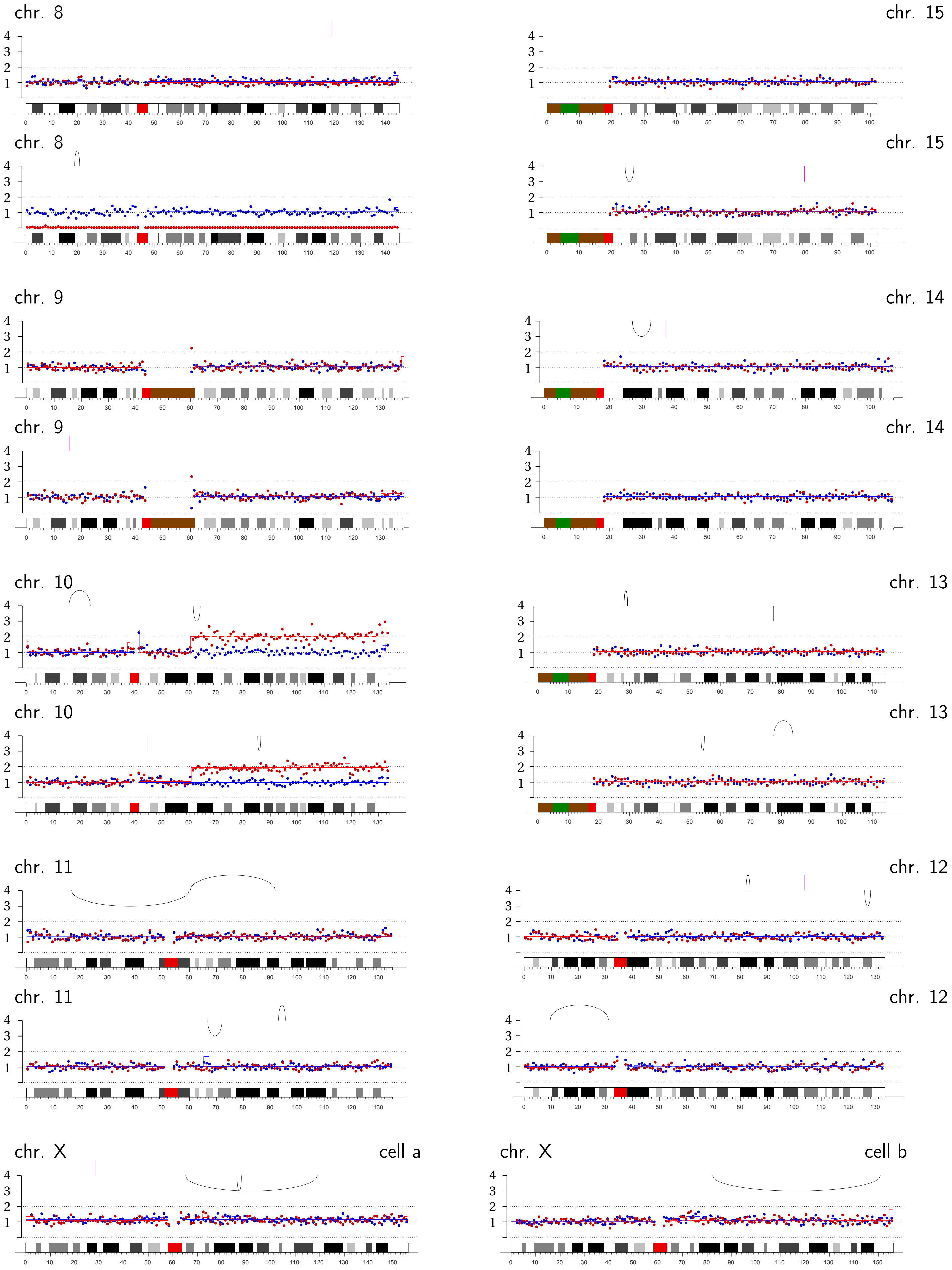
chr. 8
chr. 15
chr. 8
chr. 15
chr. 9
chr. 14
chr. 9
chr. 14
chr. 10
chr. 13
chr. 10
chr. 13
chr. 11
chr. 12
chr. 11
chr. 12
chr. X
cell a
chr. X
cell b

**Micronucleus Pair MN3 | Chromothripsis/fragmentation on both chromosome 2 homologs: 2A (2:1) and 2B (1:0).**

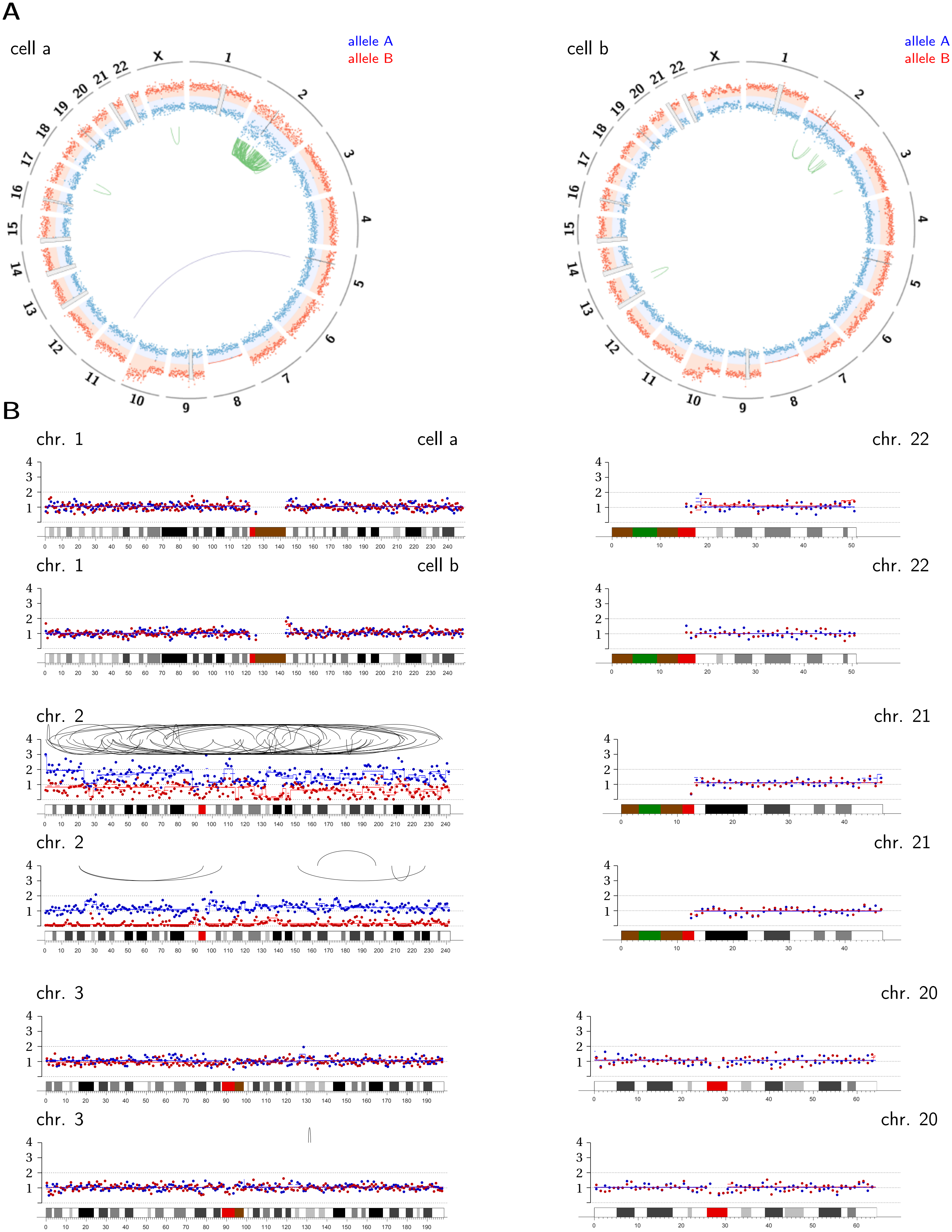

chr. 4

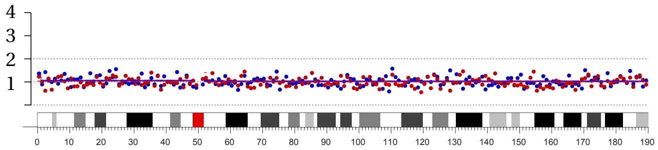

chr. 4

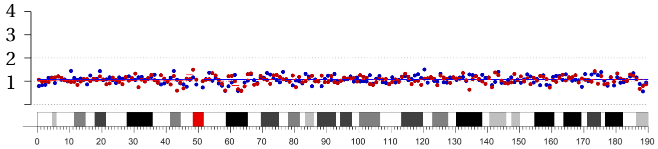

chr. 5

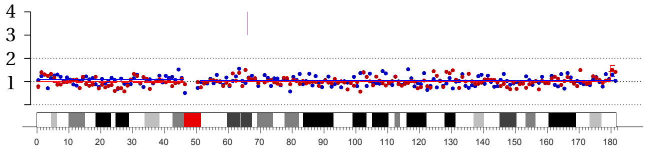

chr. 5

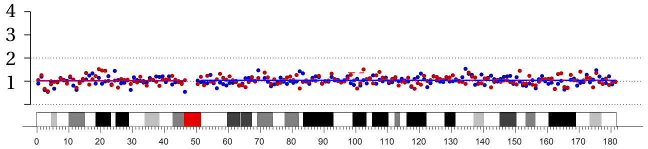

chr. 6

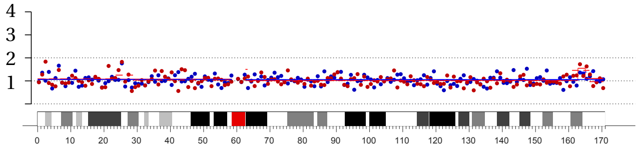

chr. 6

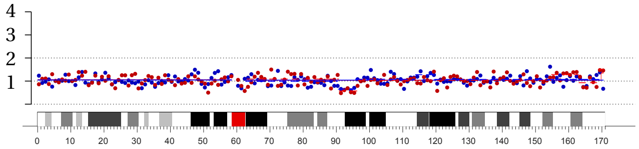

chr. 7

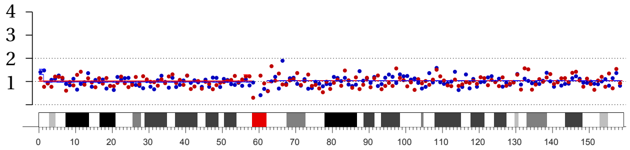

chr. 7

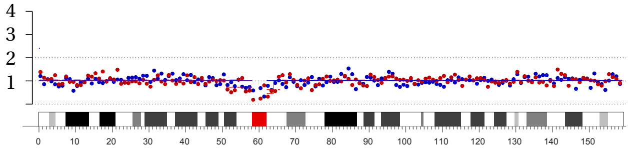

chr. 19

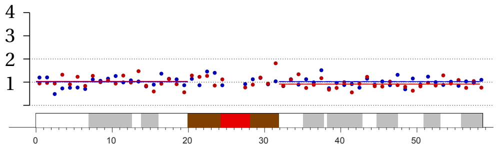

chr. 19

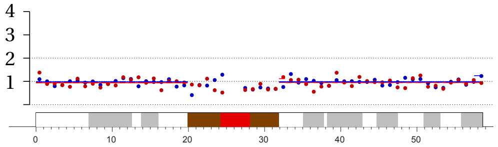

chr. 18

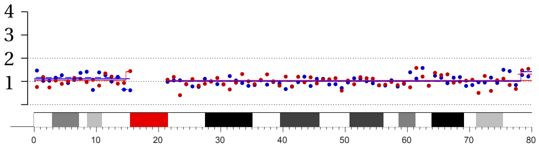

chr. 18

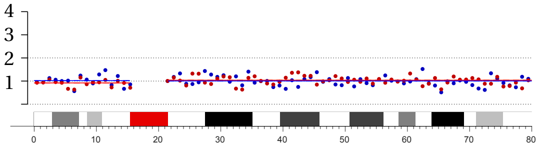

chr. 17

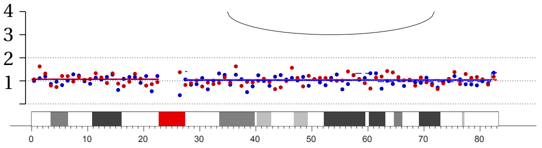

chr. 17

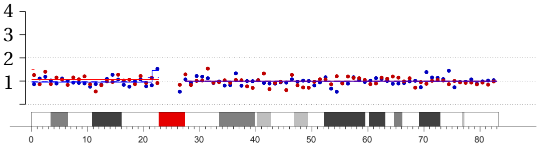

chr. 16

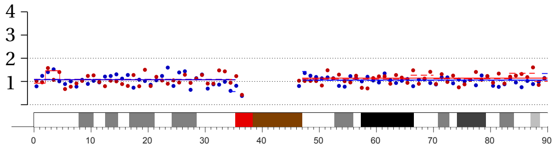

chr. 16

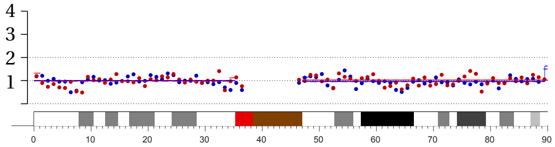

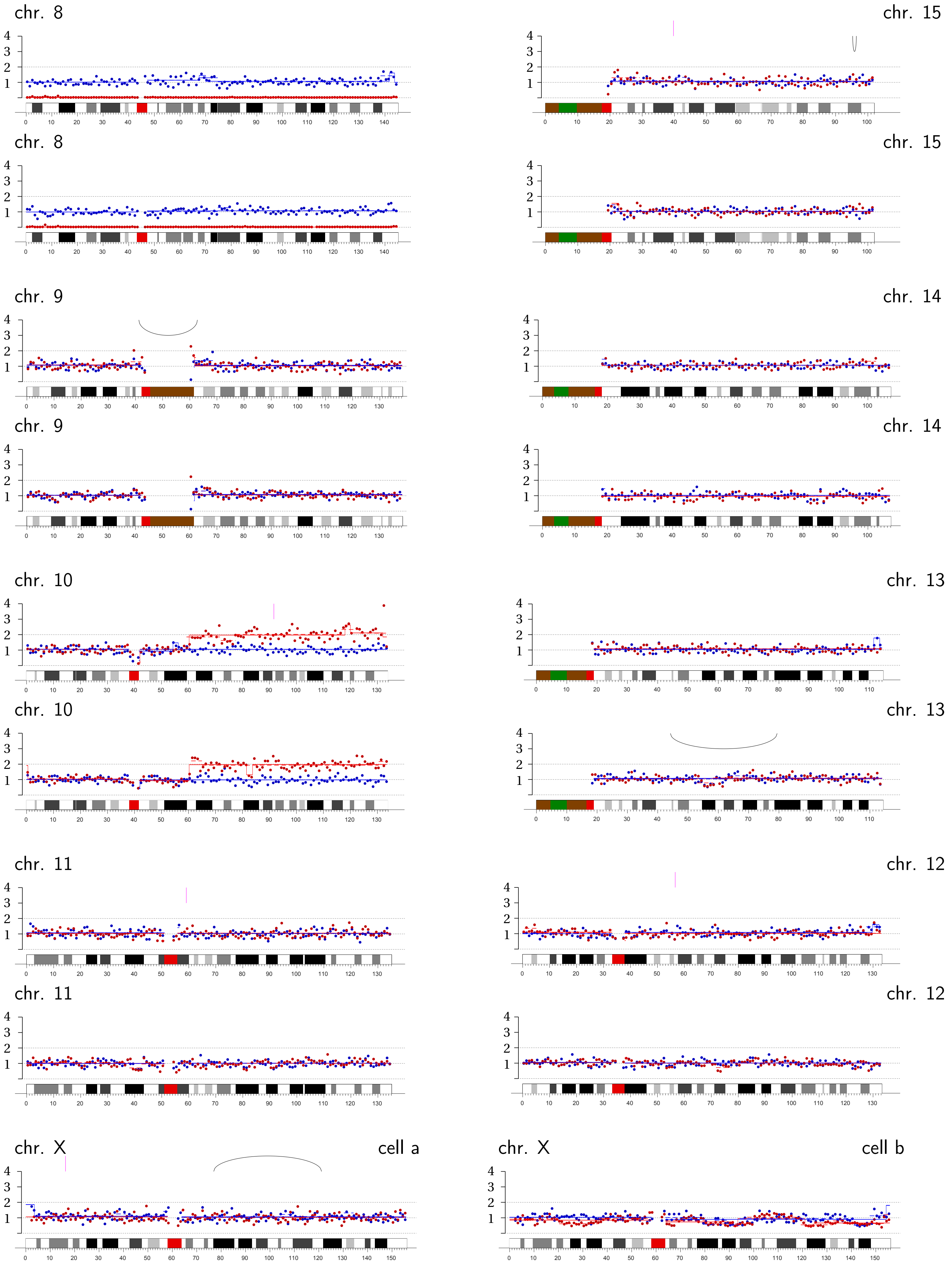
chr. 8
chr. 15
chr. 8
chr. 15
chr. 9
chr. 14
chr. 9
chr. 14
chr. 10
chr. 13
chr. 10
chr. 13
chr. 11
chr. 12
chr. 11
chr. 12
chr. X
cell a
chr. X
cell b

**Micronucleus Pair MN4** | Asymmetric copy number of chromosome 2B (1:0) with partial replication and fragmentation.

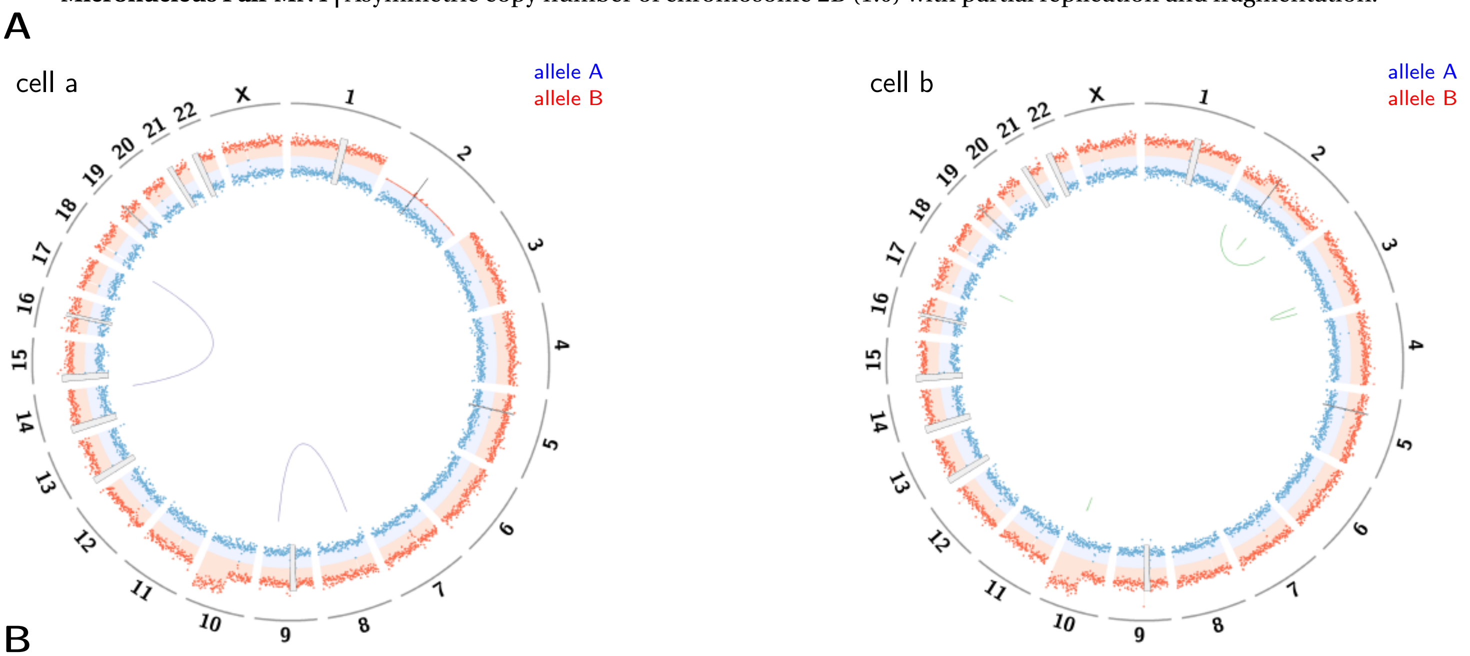


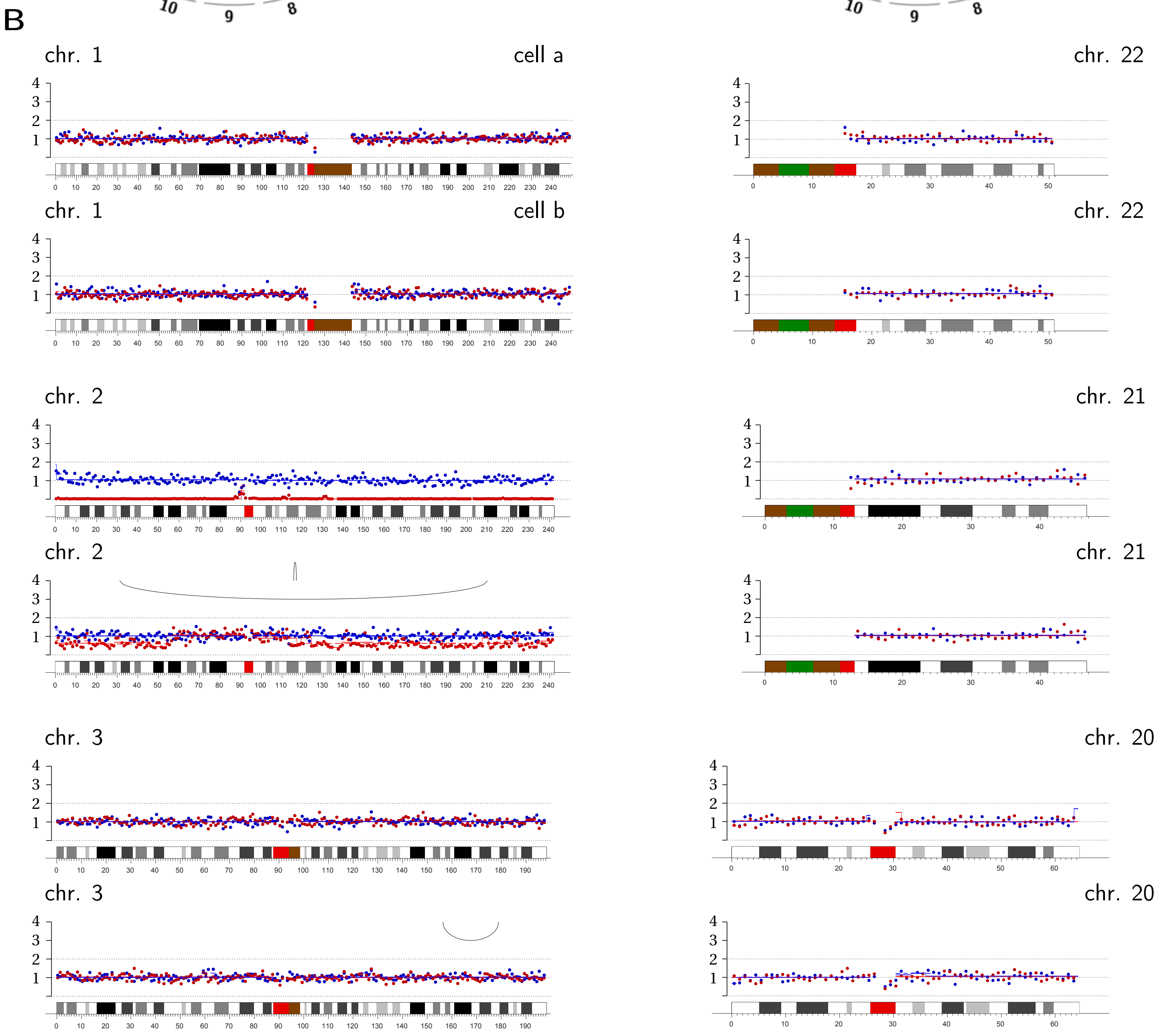

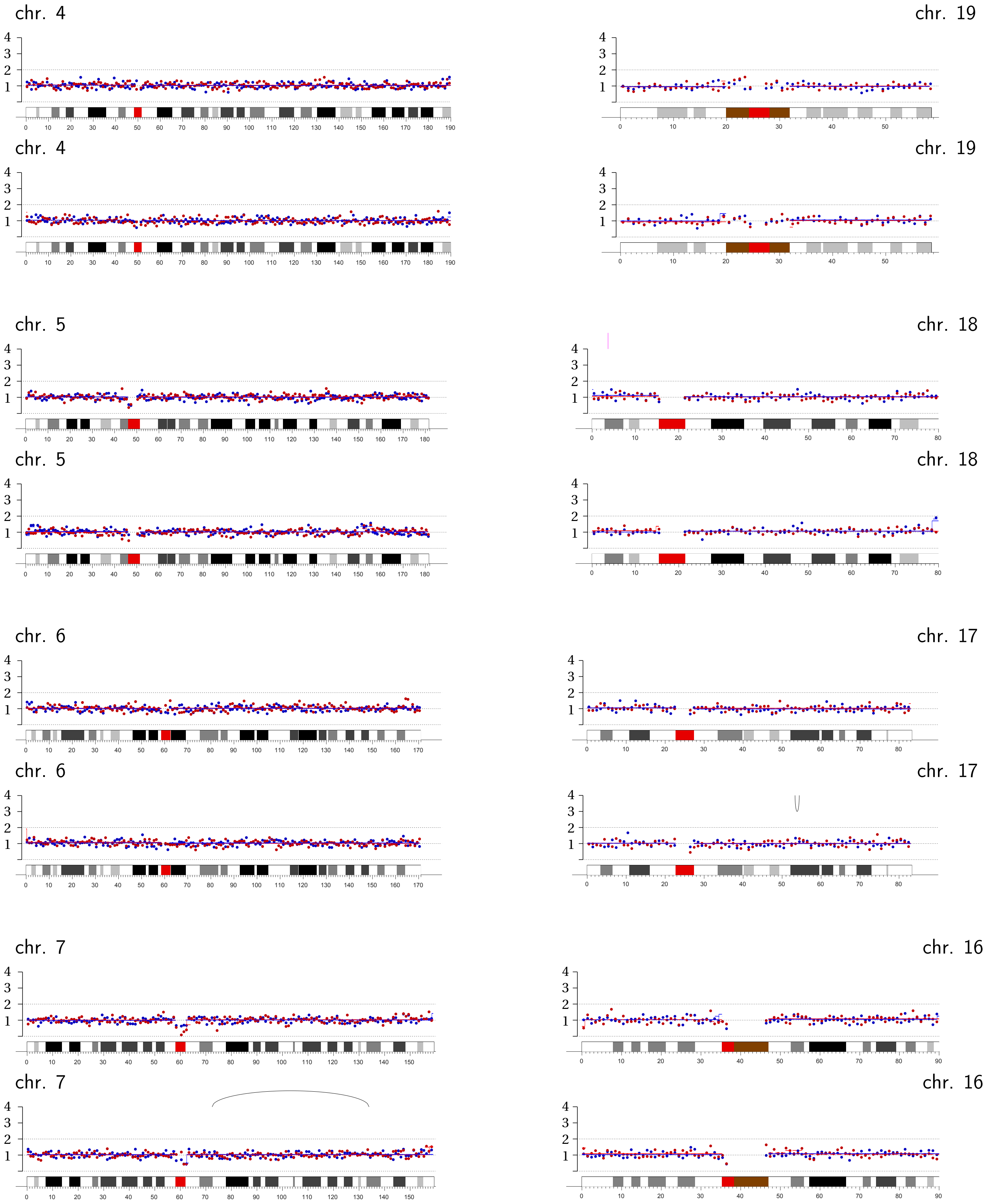

chr. 4
chr. 19
chr. 4
chr. 19
chr. 5
chr. 18
chr. 5
chr. 18
chr. 6
chr. 17
chr. 6
chr. 17
chr. 7
chr. 16
chr. 7
chr. 16

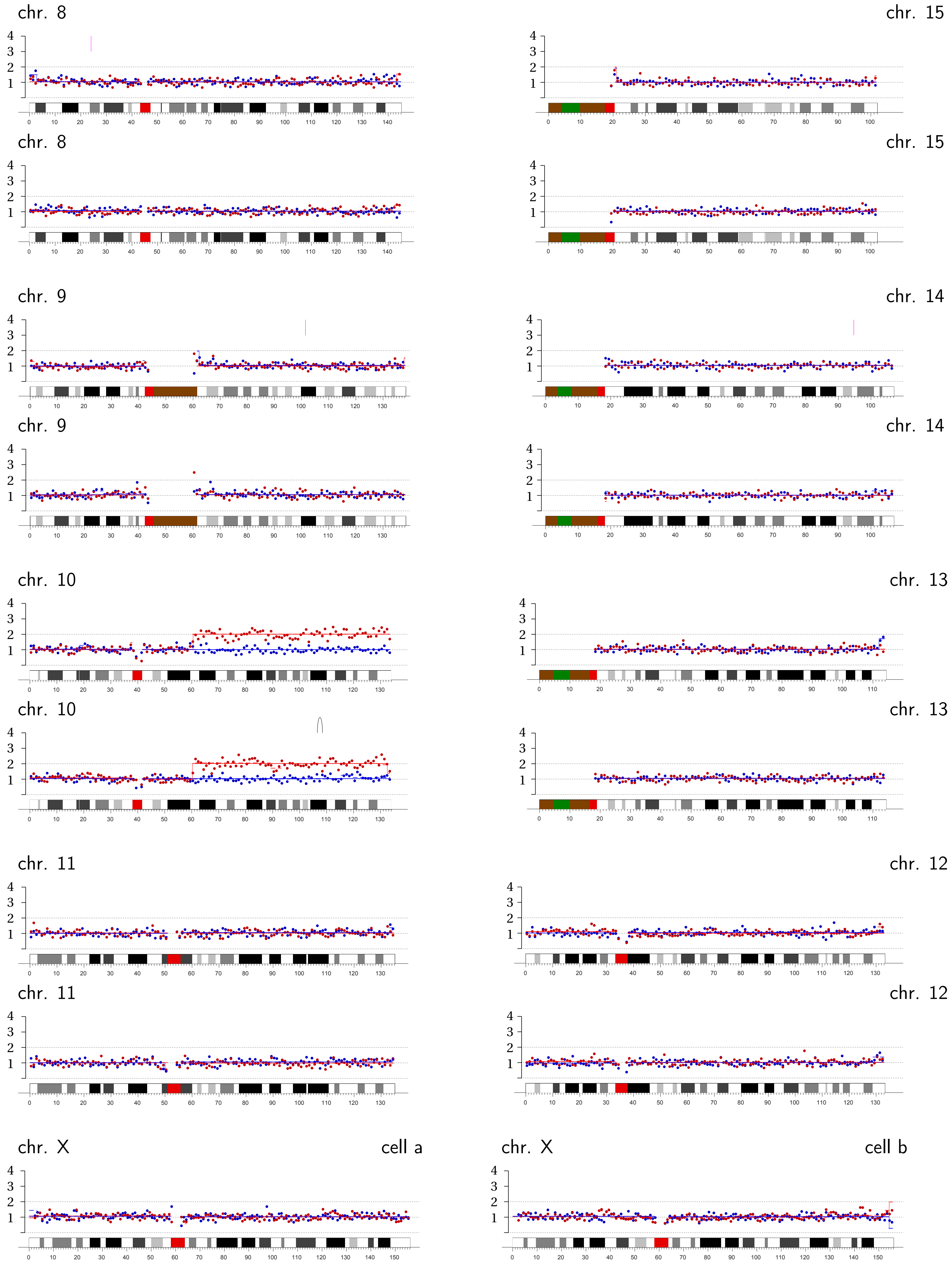
chr. 8
chr. 15
chr. 8
chr. 15
chr. 9
chr. 14
chr. 9
chr. 14
chr. 10
chr. 13
chr. 10
chr. 13
chr. 11
chr. 12
chr. 11
chr. 12
chr. X
cell a
chr. X
cell b

**Micronucleus Pair MN5 | Chromothripsis/fragmentation on chromosome 2B (1:0) with partial replication.**

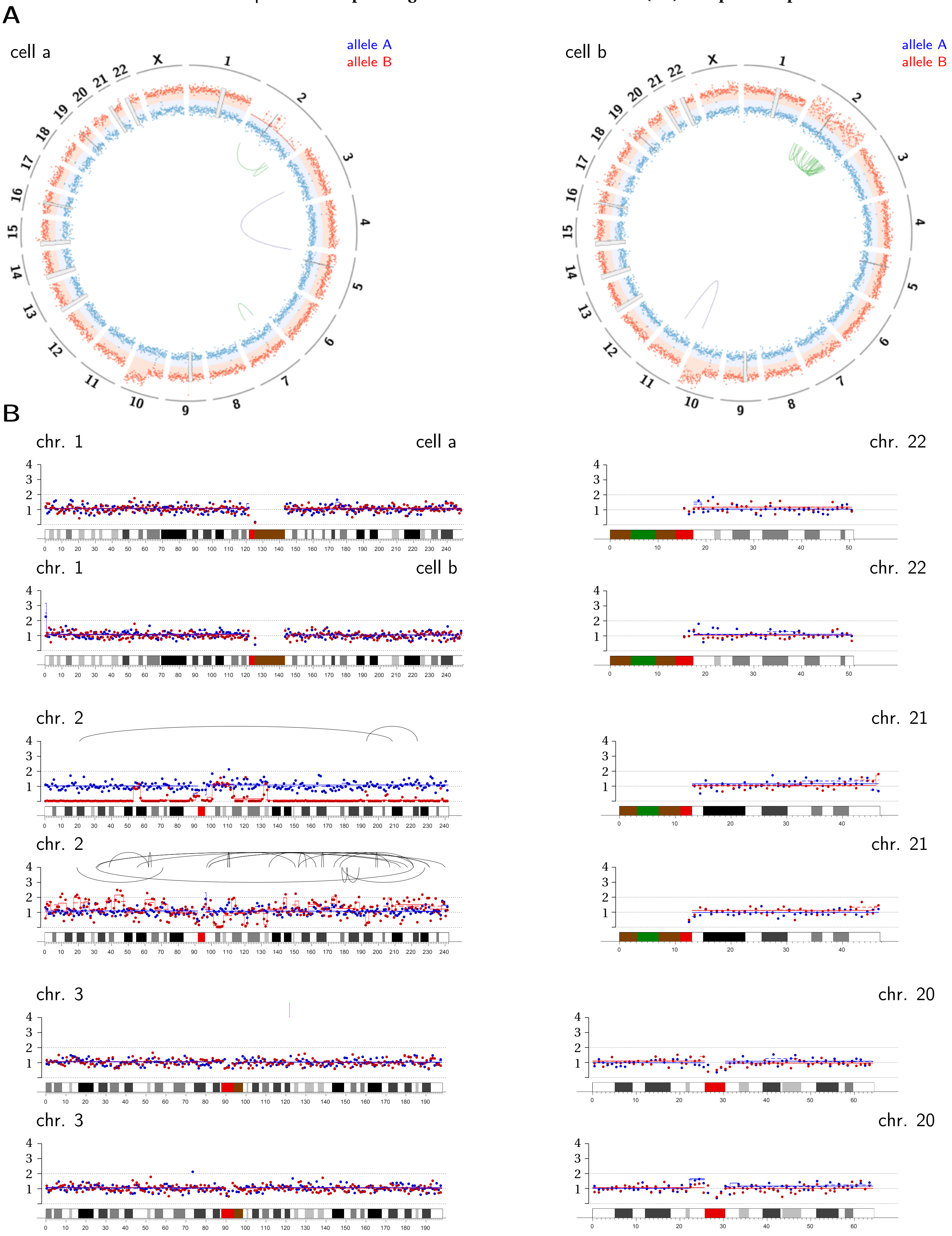

chr. 4

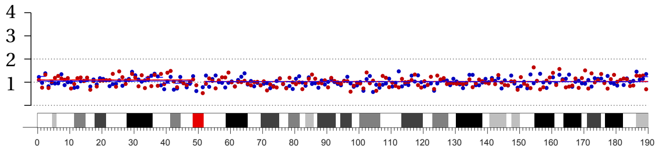

chr. 4

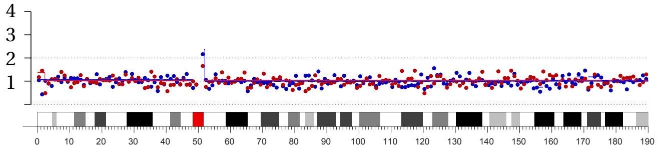

chr. 5

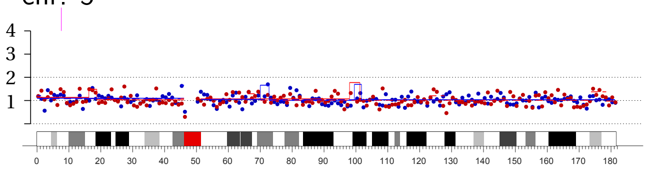

chr. 5

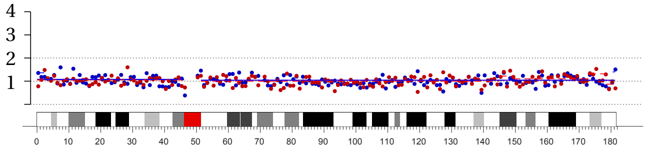

chr. 6

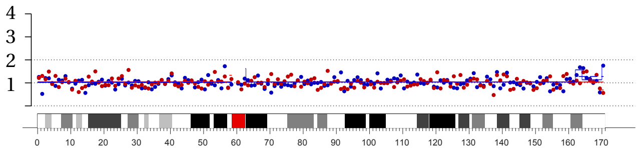

chr. 6

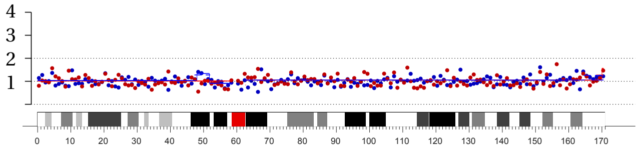

chr. 7

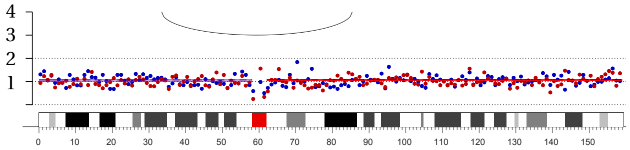

chr. 7

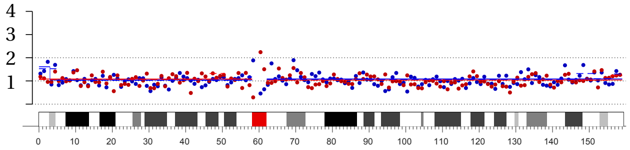

chr. 19

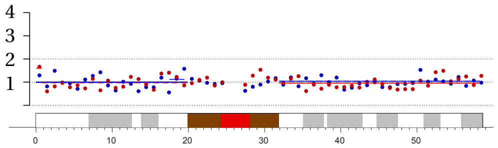

chr. 19

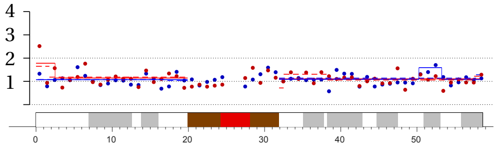

chr. 18

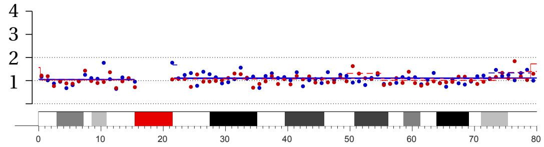

chr. 18

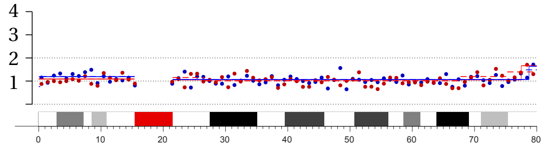

chr. 17

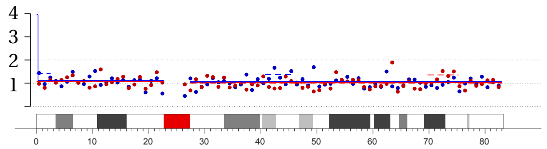

chr. 17

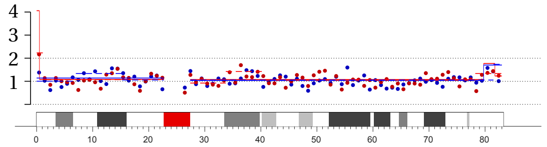

chr. 16

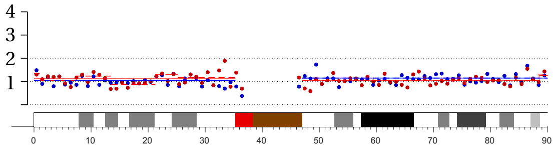

chr. 16

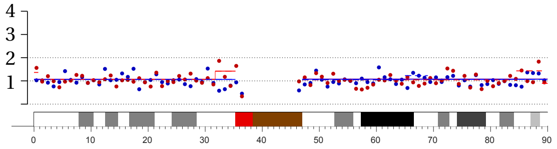

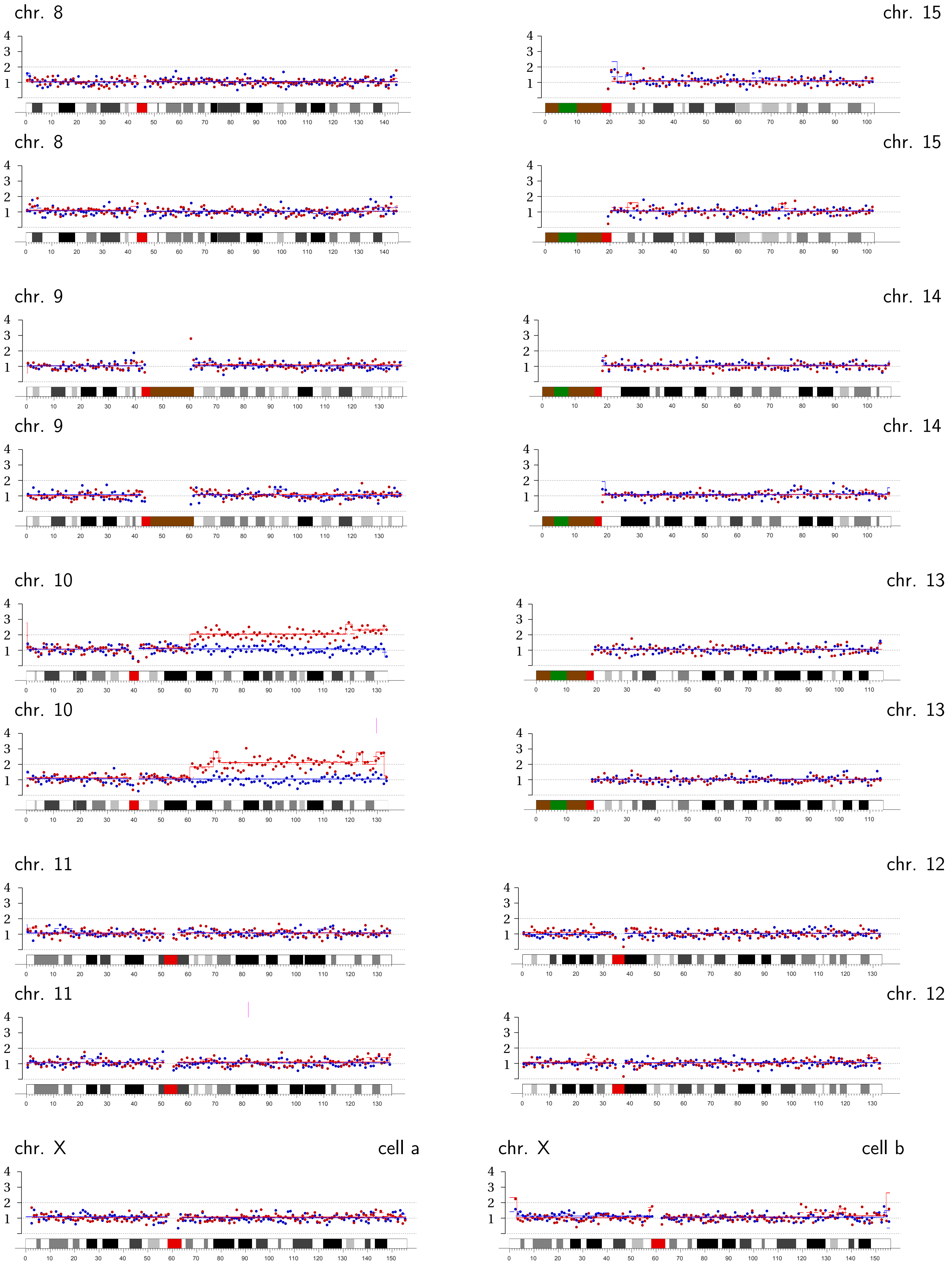
chr. 8
chr. 15
chr. 8
chr. 15
chr. 9
chr. 14
chr. 9
chr. 14
chr. 10
chr. 13
chr. 10
chr. 13
chr. 11
chr. 12
chr. 11
chr. 12
chr. X
cell a
chr. X
cell b

**Micronucleus Pair MN6 | Chromothripsis/fragmentation on chromosome 1B (1:0).**

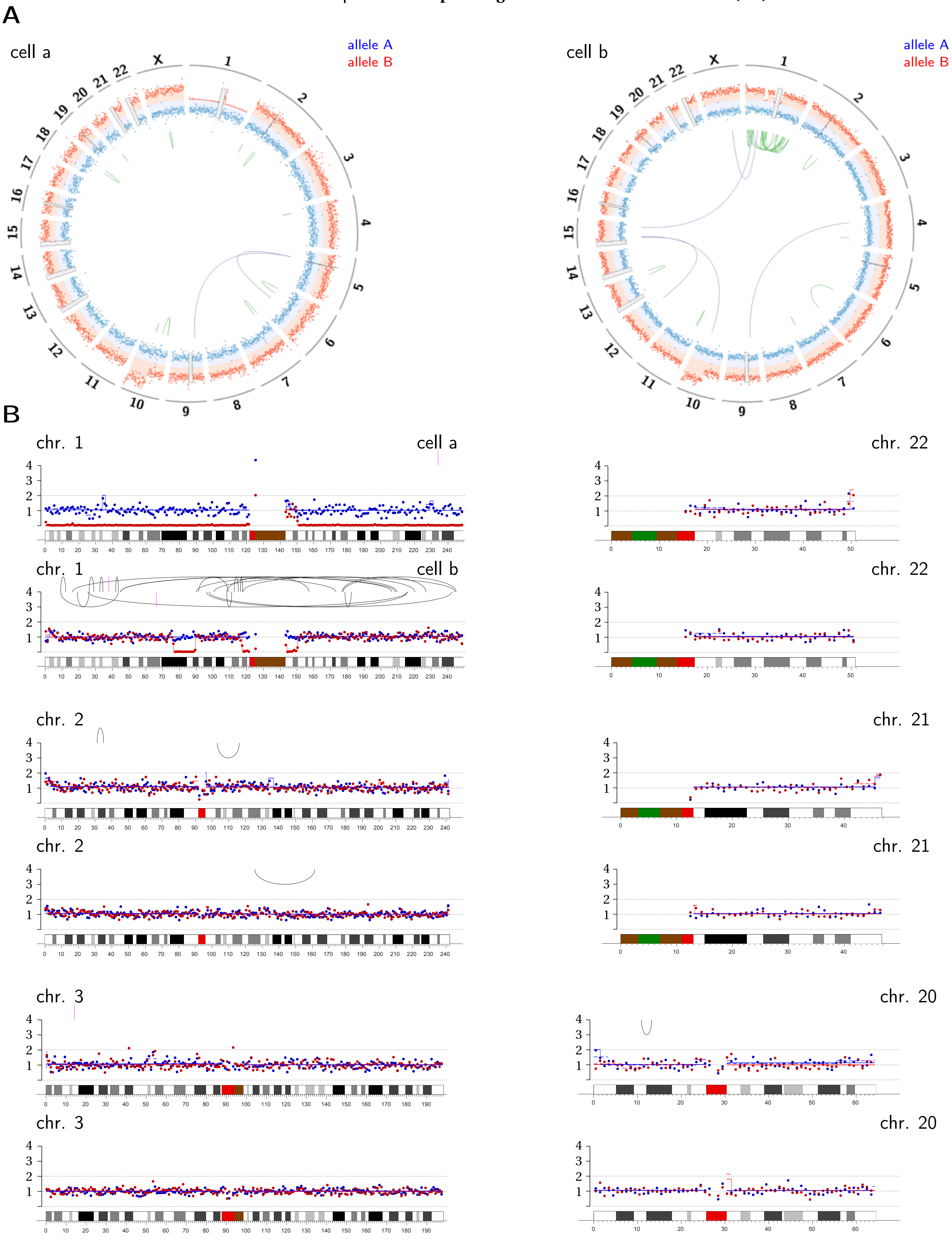

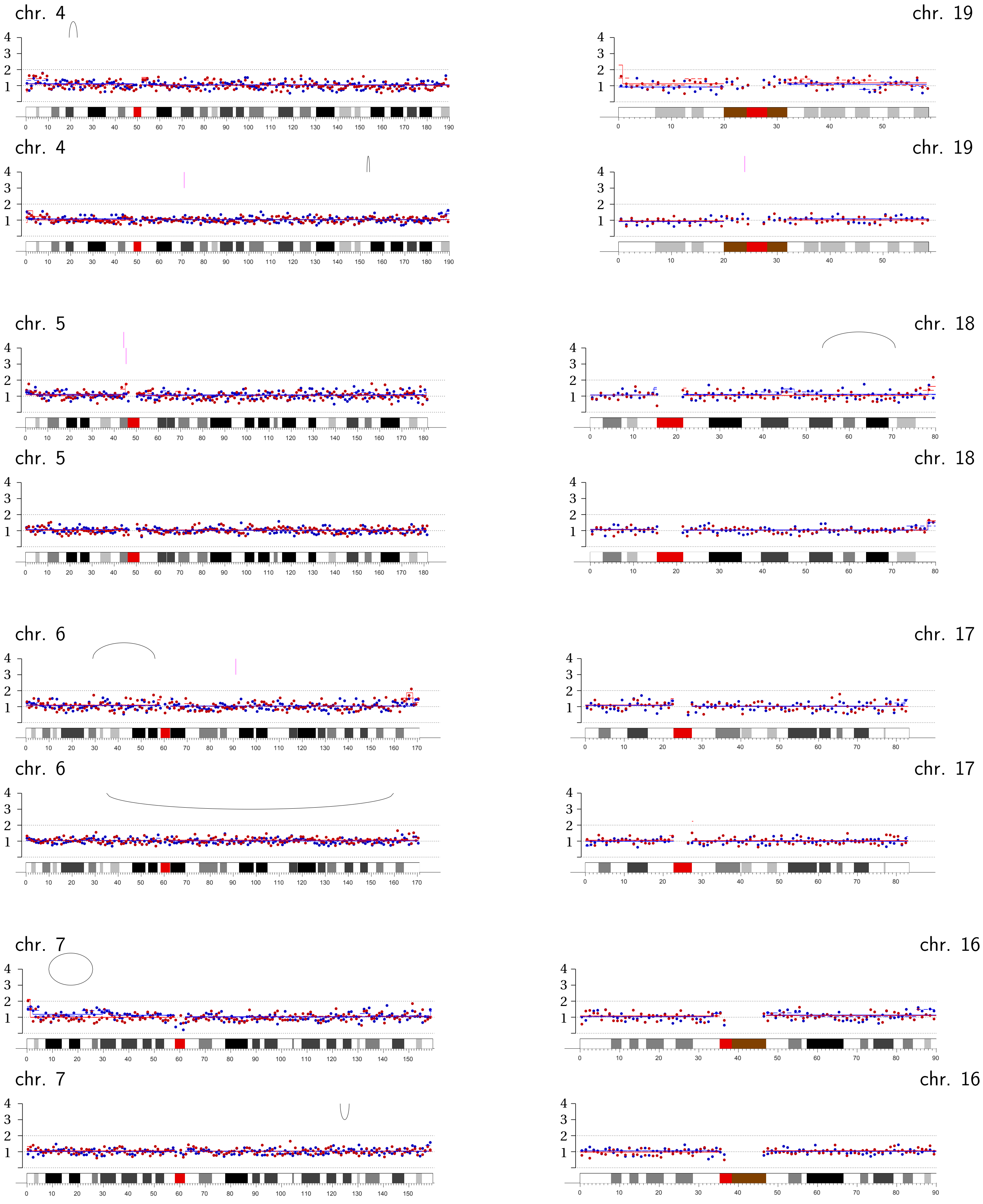
chr. 4
chr. 19
chr. 4
chr. 19
chr. 5
chr. 18
chr. 5
chr. 18
chr. 6
chr. 17
chr. 6
chr. 17
chr. 7
chr. 16
chr. 7
chr. 16

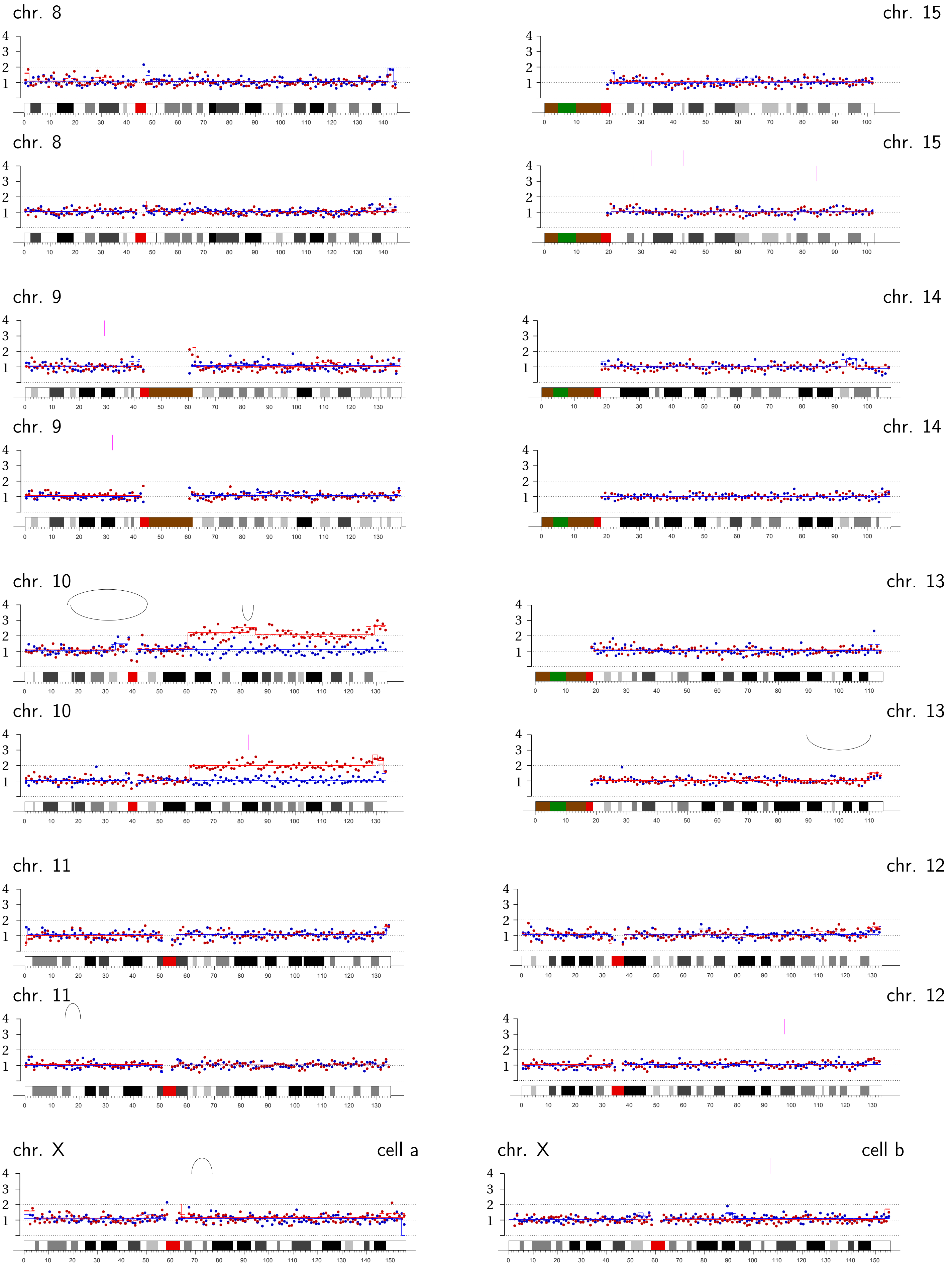
chr. 8
chr. 15
chr. 8
chr. 15
chr. 9
chr. 14
chr. 9
chr. 14
chr. 10
chr. 13
chr. 10
chr. 13
chr. 11
chr. 12
chr. 11
chr. 12
chr. X
cell a
chr. X
cell b

**Micronucleus Pair MN7 | Chromothripsis/fragmentation on chromosome 11B (1:0)**; daughter b likely in G2 with under-replicated Chr. 11 in persistent MN.

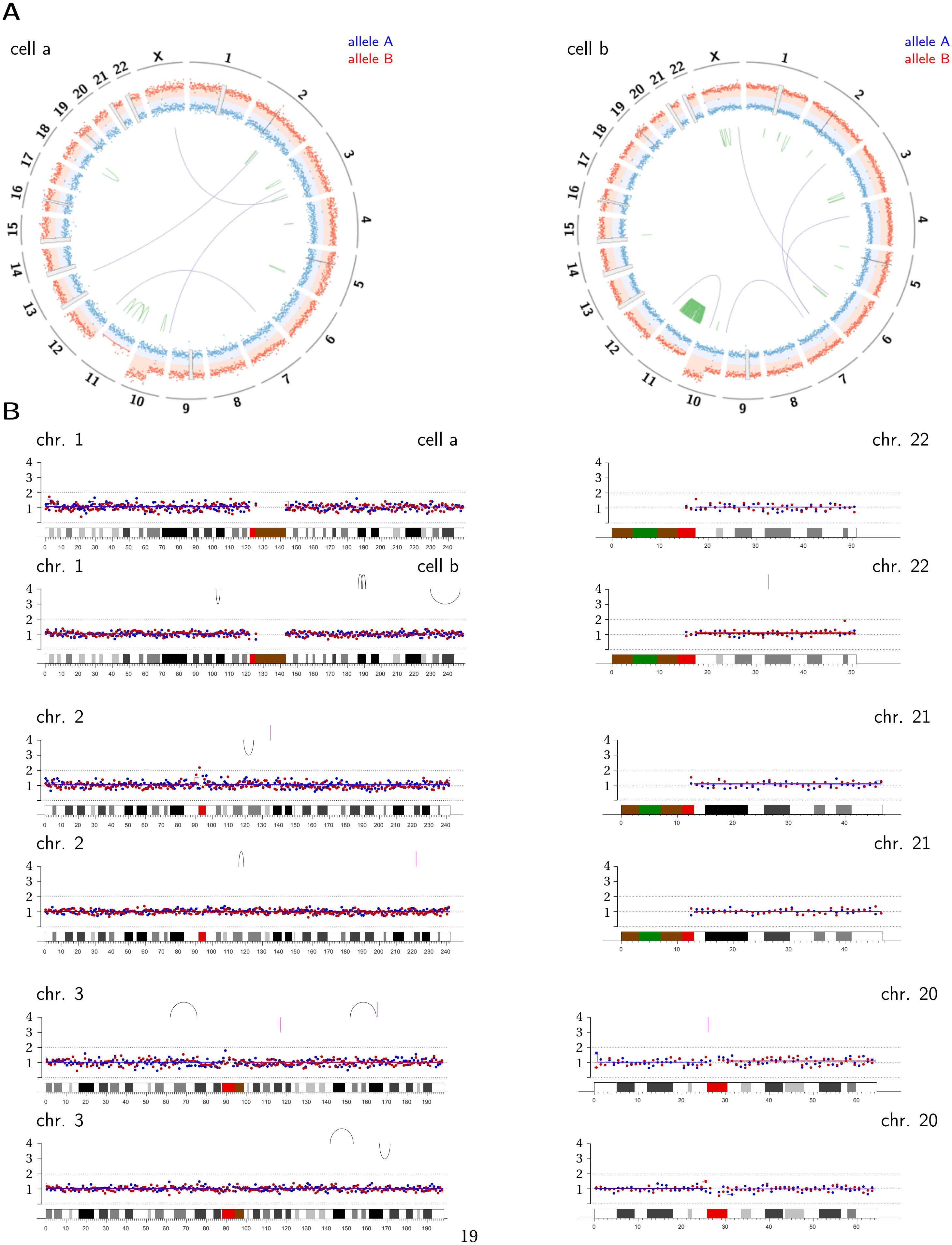

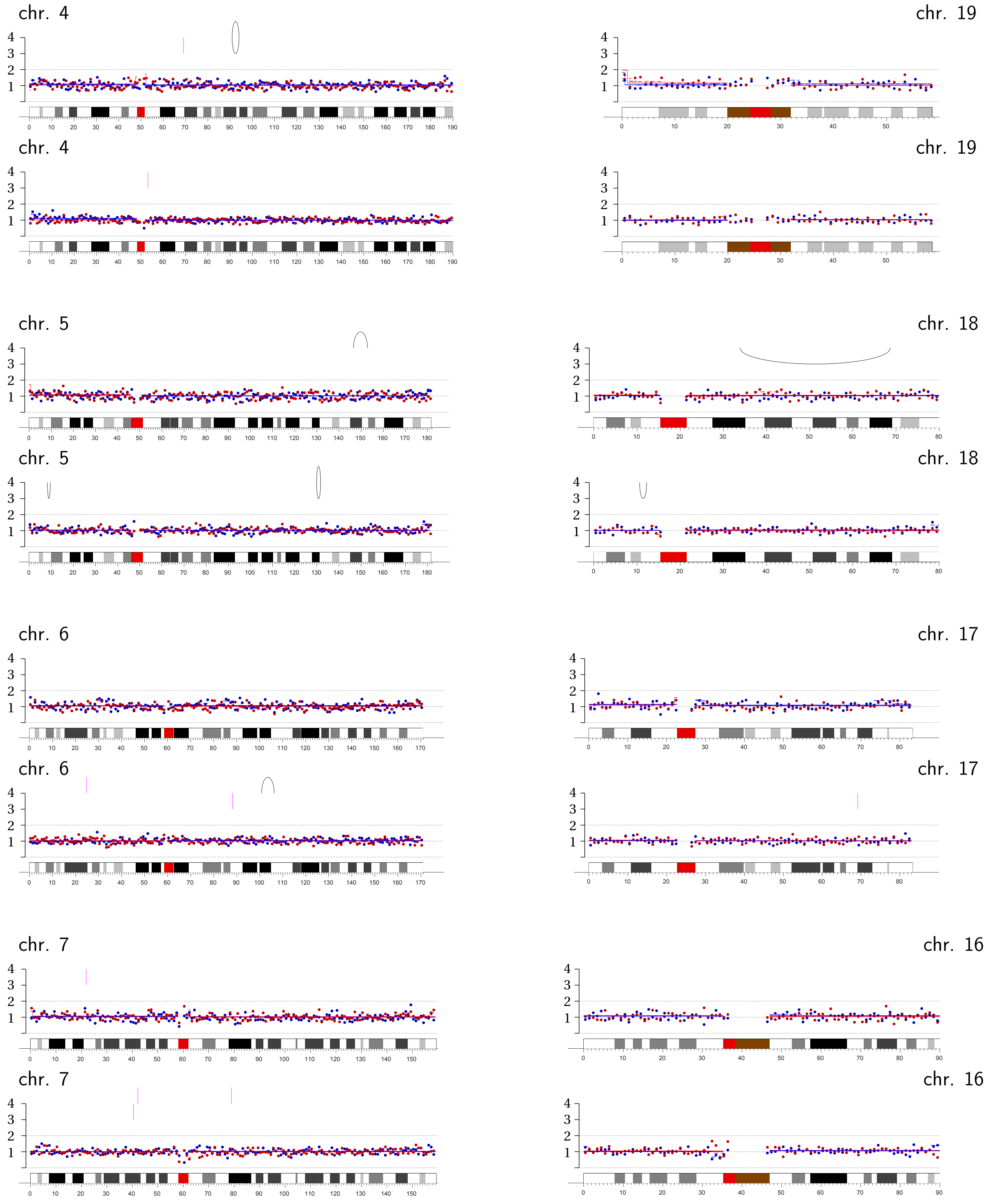
chr. 4
chr. 4
chr. 5
chr. 5
chr. 6
chr. 6
chr. 7
chr. 7
chr. 19
chr. 19
chr. 18
chr. 18
chr. 17
chr. 17
chr. 16
chr. 16

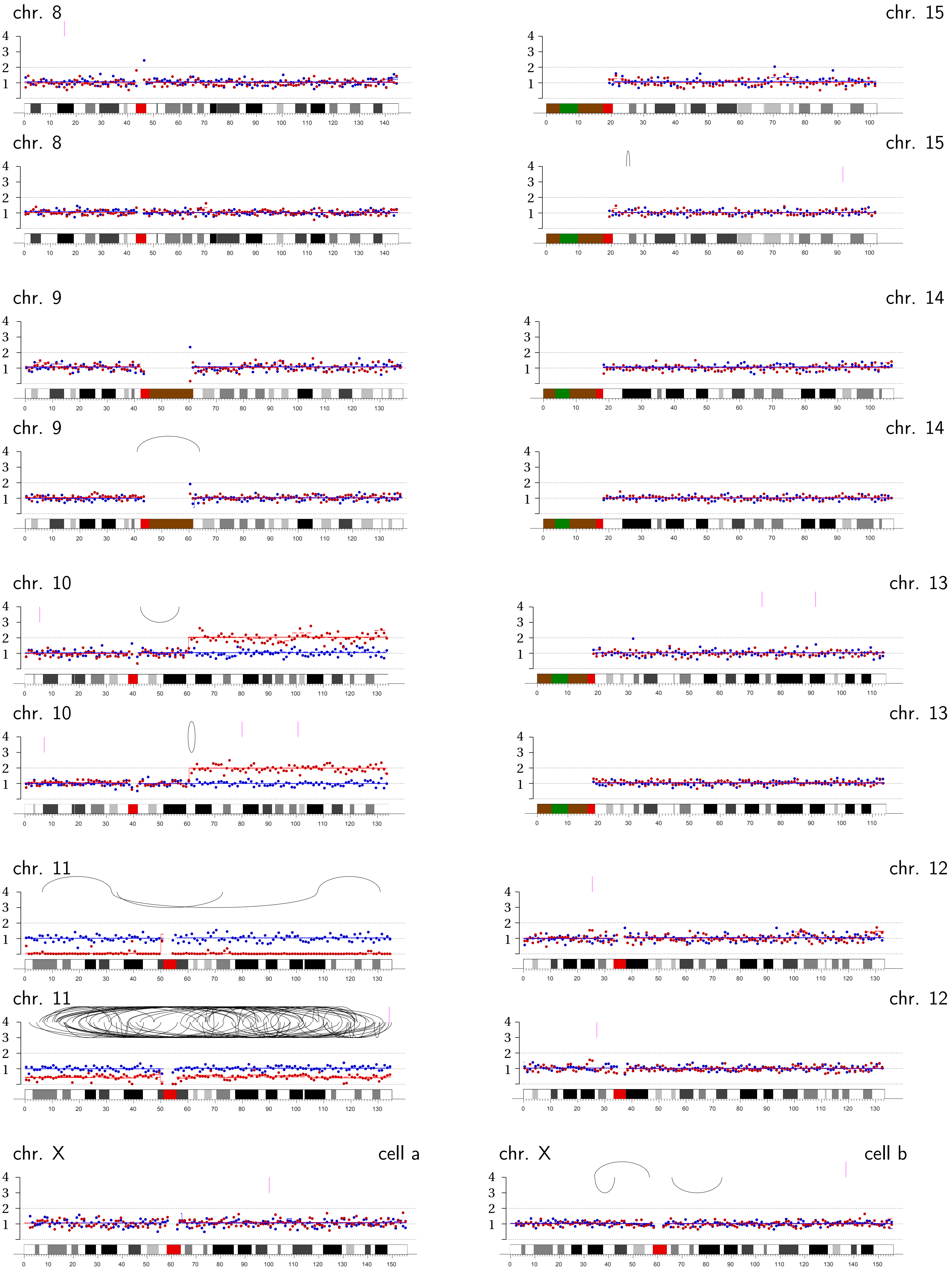
chr. 8
chr. 15
chr. 8
chr. 15
chr. 9
chr. 14
chr. 9
chr. 14
chr. 10
chr. 13
chr. 10
chr. 13
chr. 11
chr. 12
chr. 11
chr. 12
chr. X
cell a
chr. X
cell b

**Micronucleus Pair MN8 | Complex rearrangements involving chromosomes 1B p-arm (2:1), 10A (2:1), and 18B(2:1).**

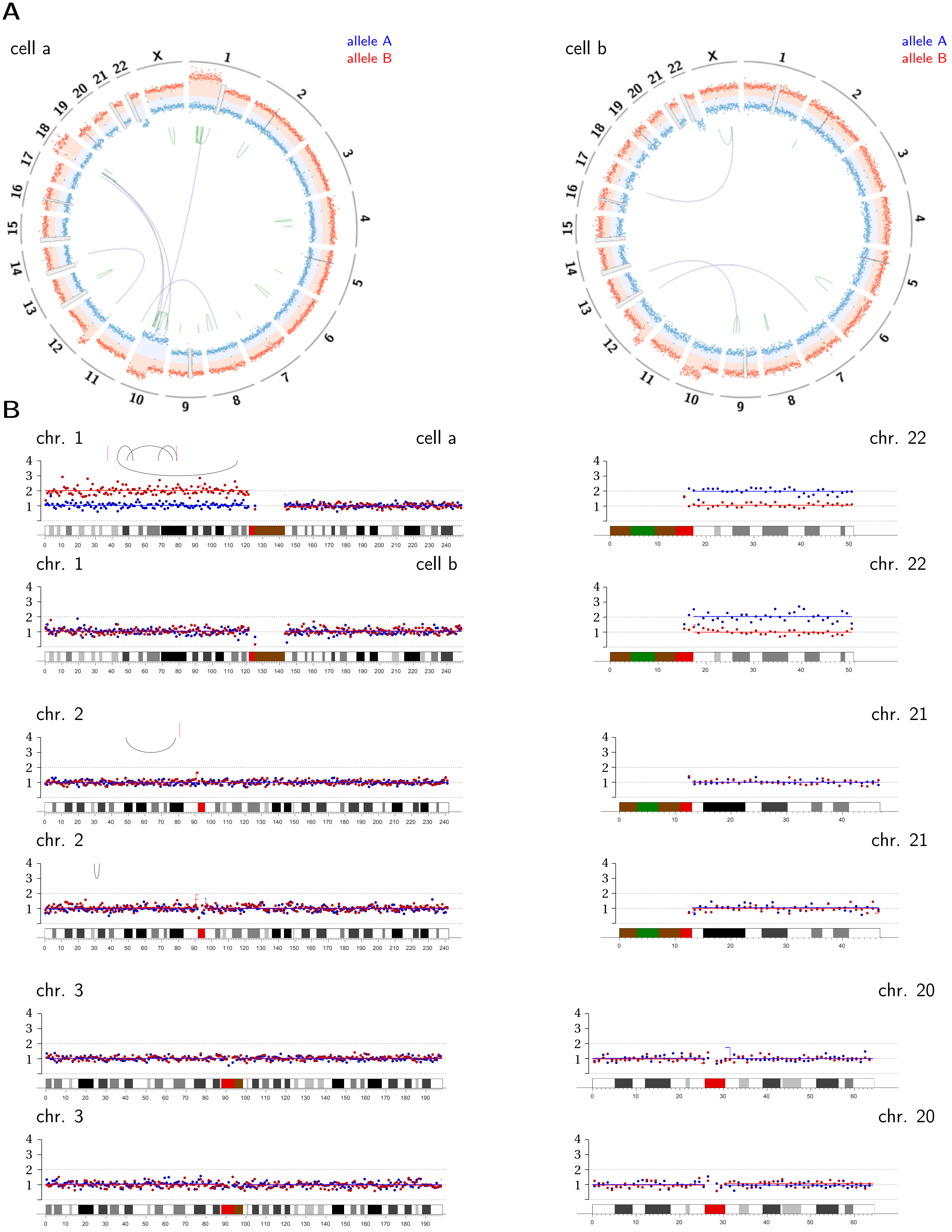

chr. 4

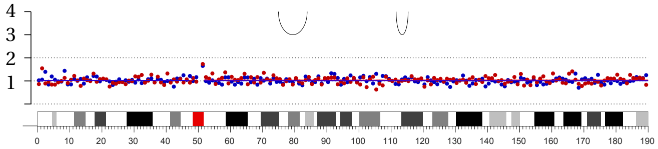

chr. 4

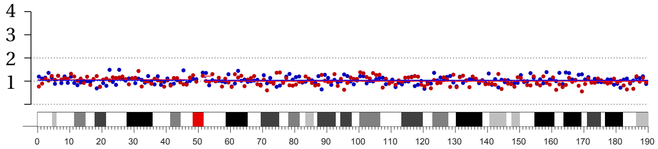

chr. 5

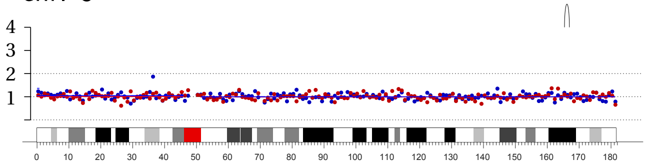

chr. 5

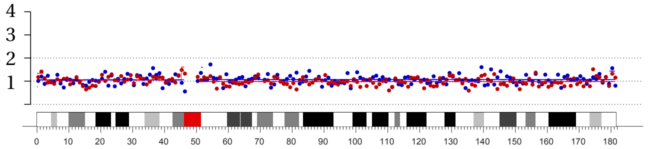

chr. 6

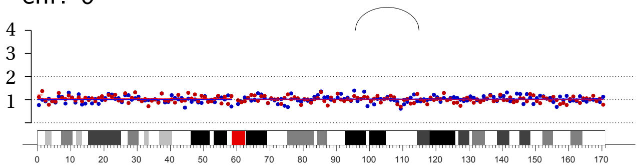

chr. 6

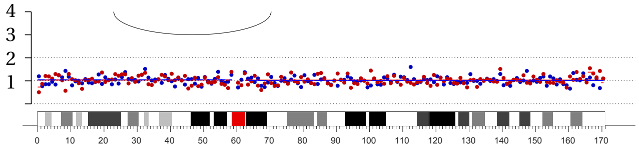

chr. 7

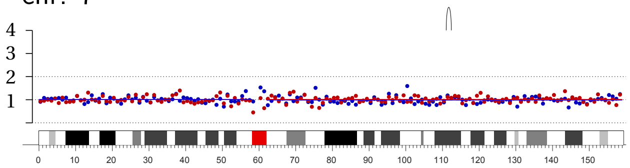

chr. 7

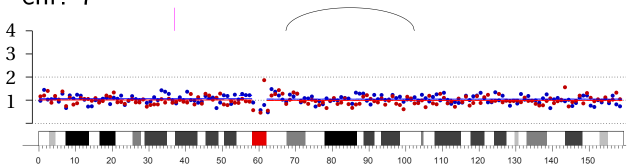

chr. 19

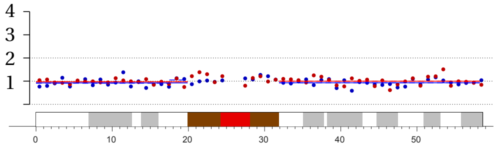

chr. 19

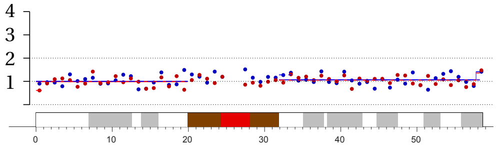

chr. 18

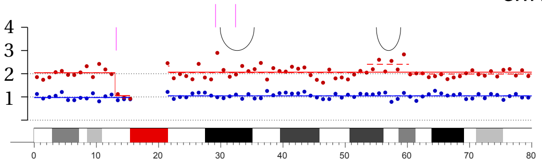

chr. 18

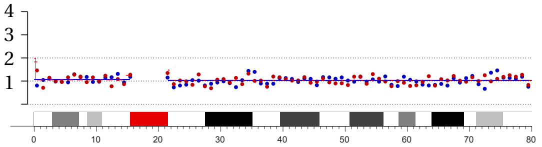

chr. 17

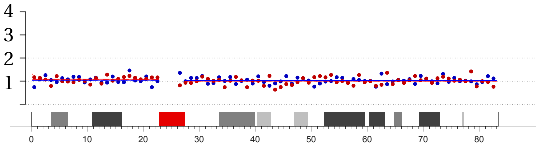

chr. 17

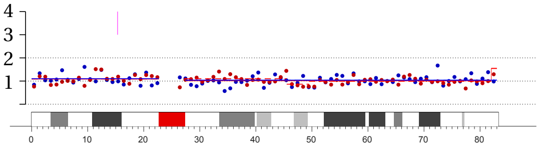

chr. 16

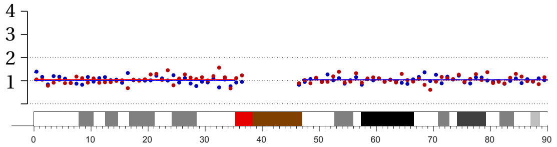

chr. 16

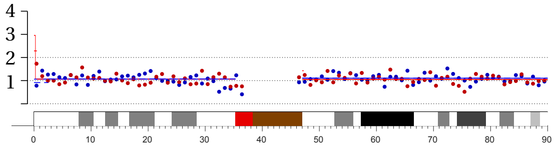

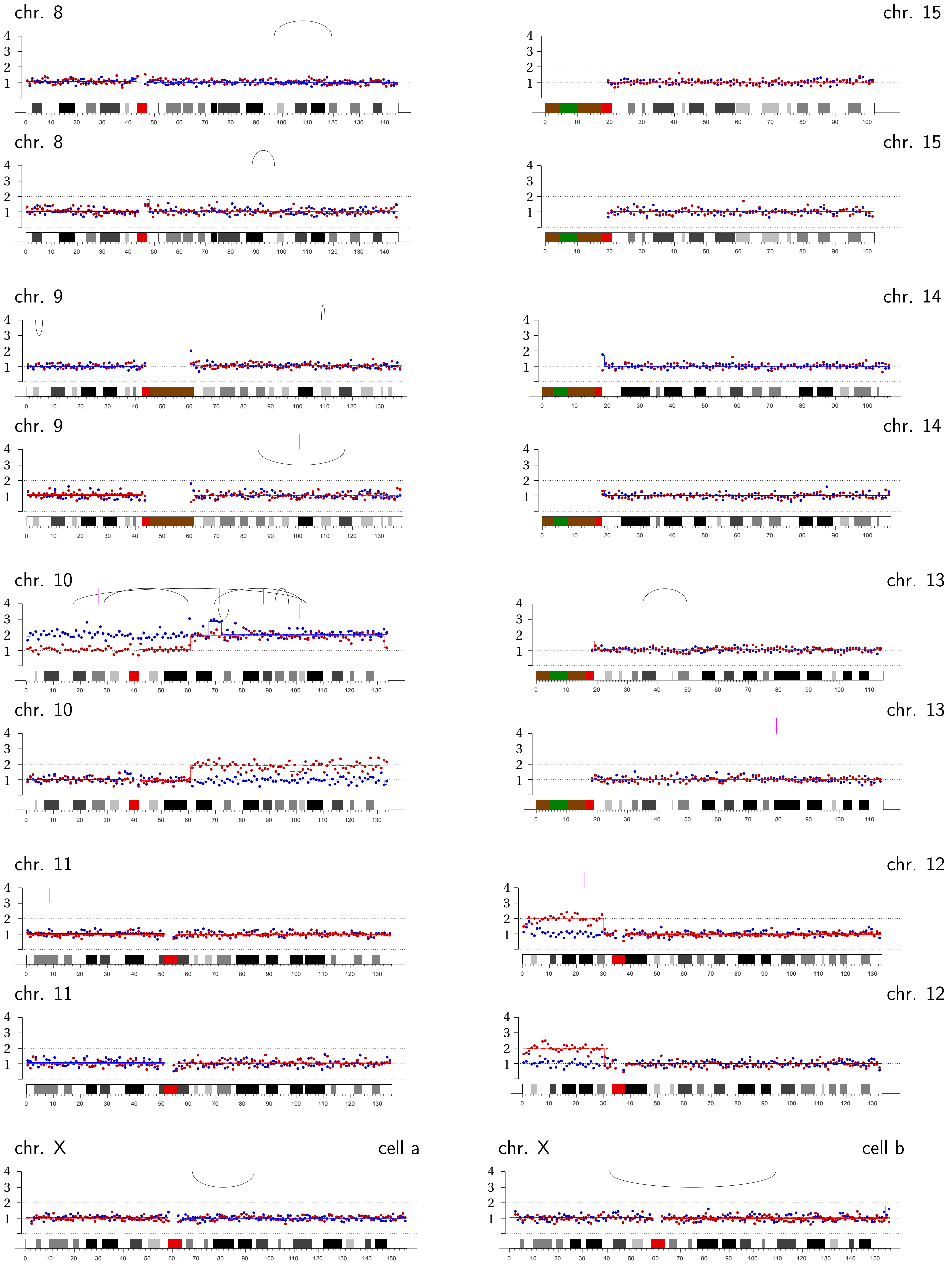
chr. 8
chr. 15
chr. 8
chr. 15
chr. 9
chr. 14
chr. 9
chr. 14
chr. 10
chr. 13
chr. 10
chr. 13
chr. 11
chr. 12
chr. 11
chr. 12
chr. X
cell a
chr. X
cell b

**Micronucleus Pair MN9 | Chromothripsis on chromosome 1A (1:0)**; daughter b likely in G2 with under-replicated Chr. 1A in persistent MN.

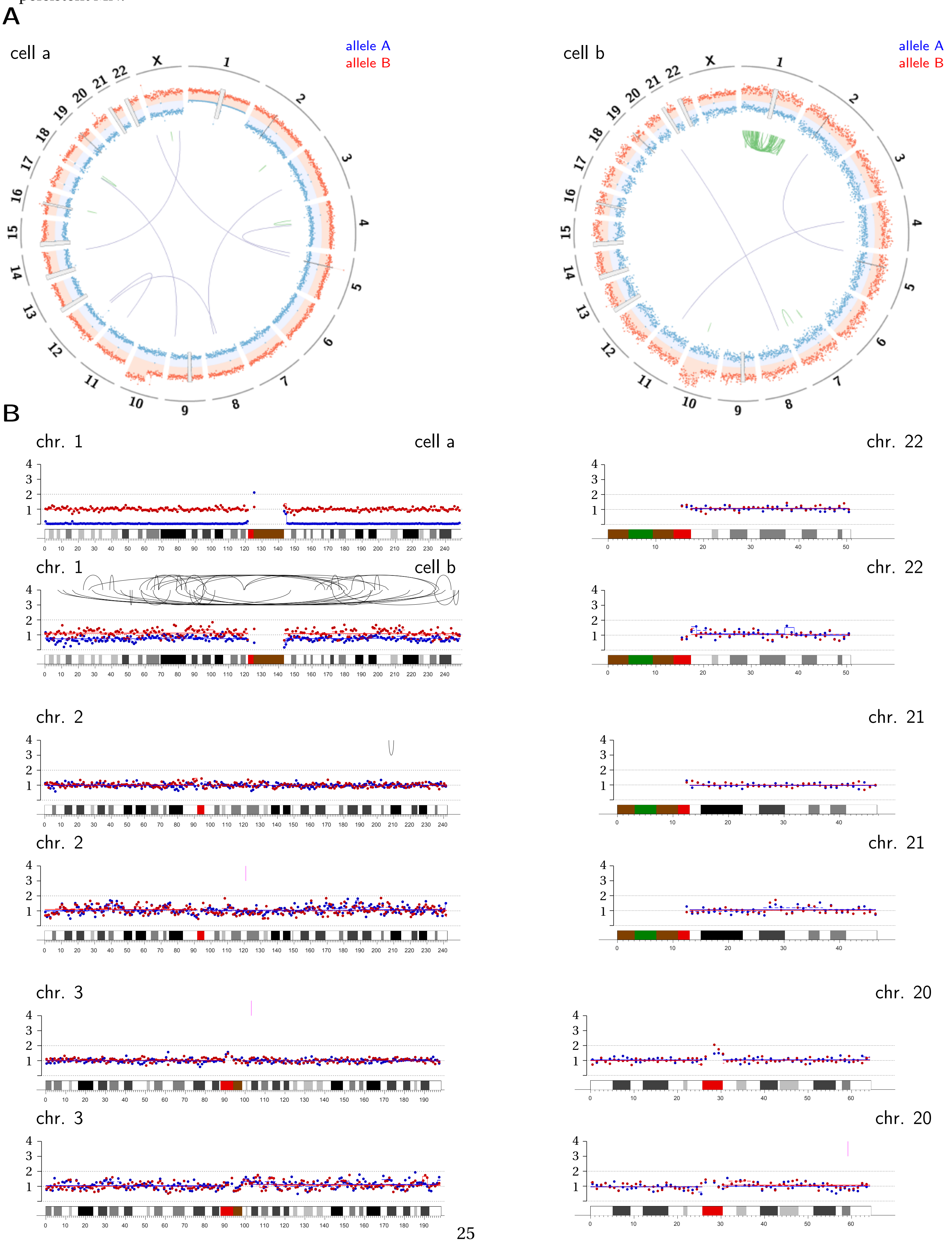

chr. 4

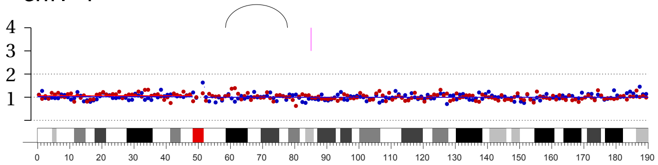

chr. 4

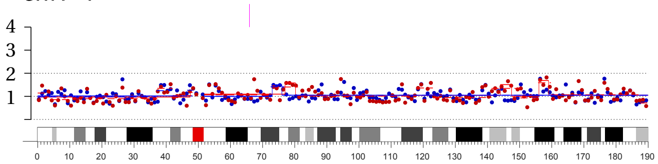

chr. 5

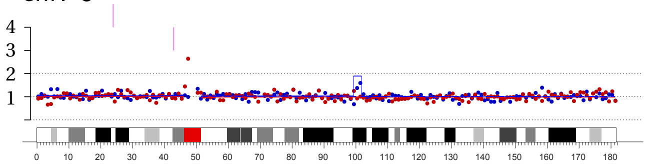

chr. 5

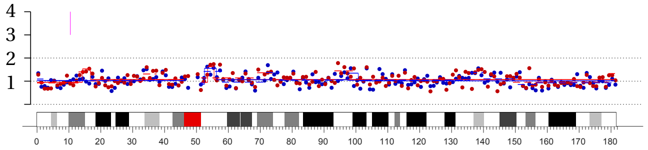

chr. 6

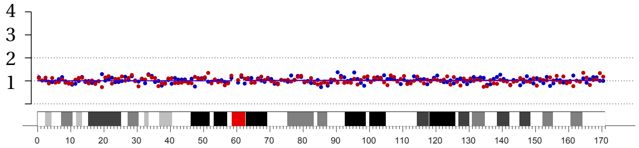

chr. 6

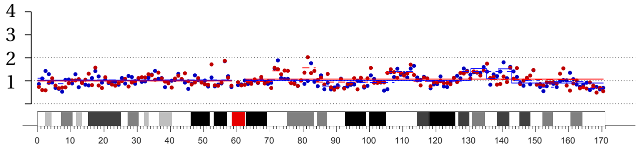

chr. 7

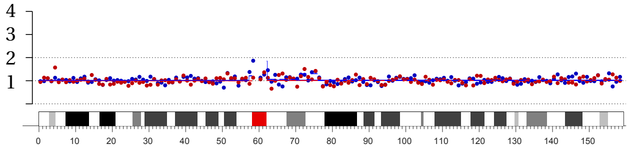

chr. 7

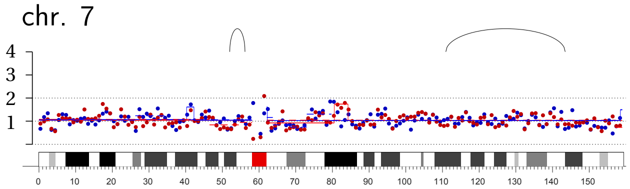

chr. 19

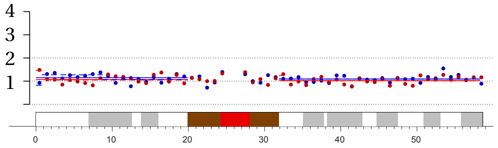

chr. 19

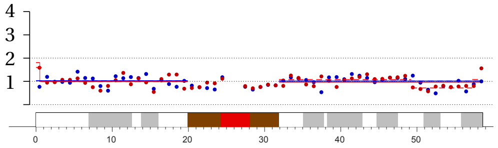

chr. 18

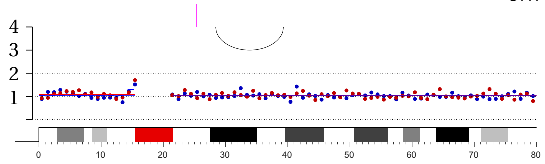

chr. 18

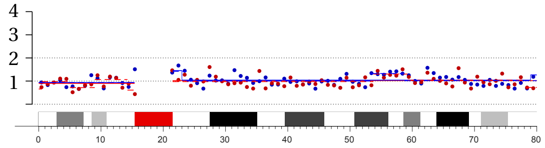

chr. 17

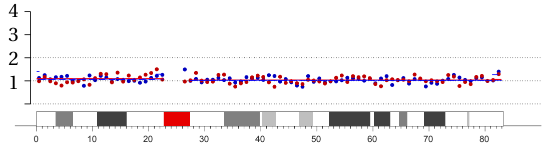

chr. 17

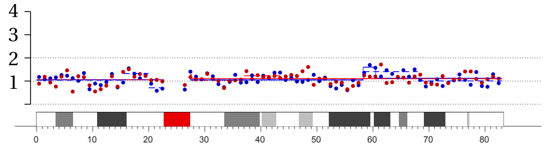

chr. 16

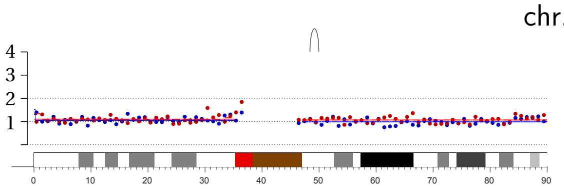

chr. 16

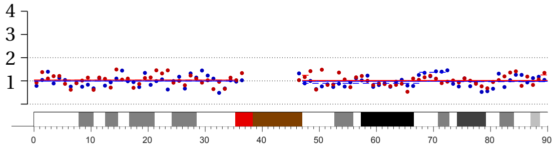

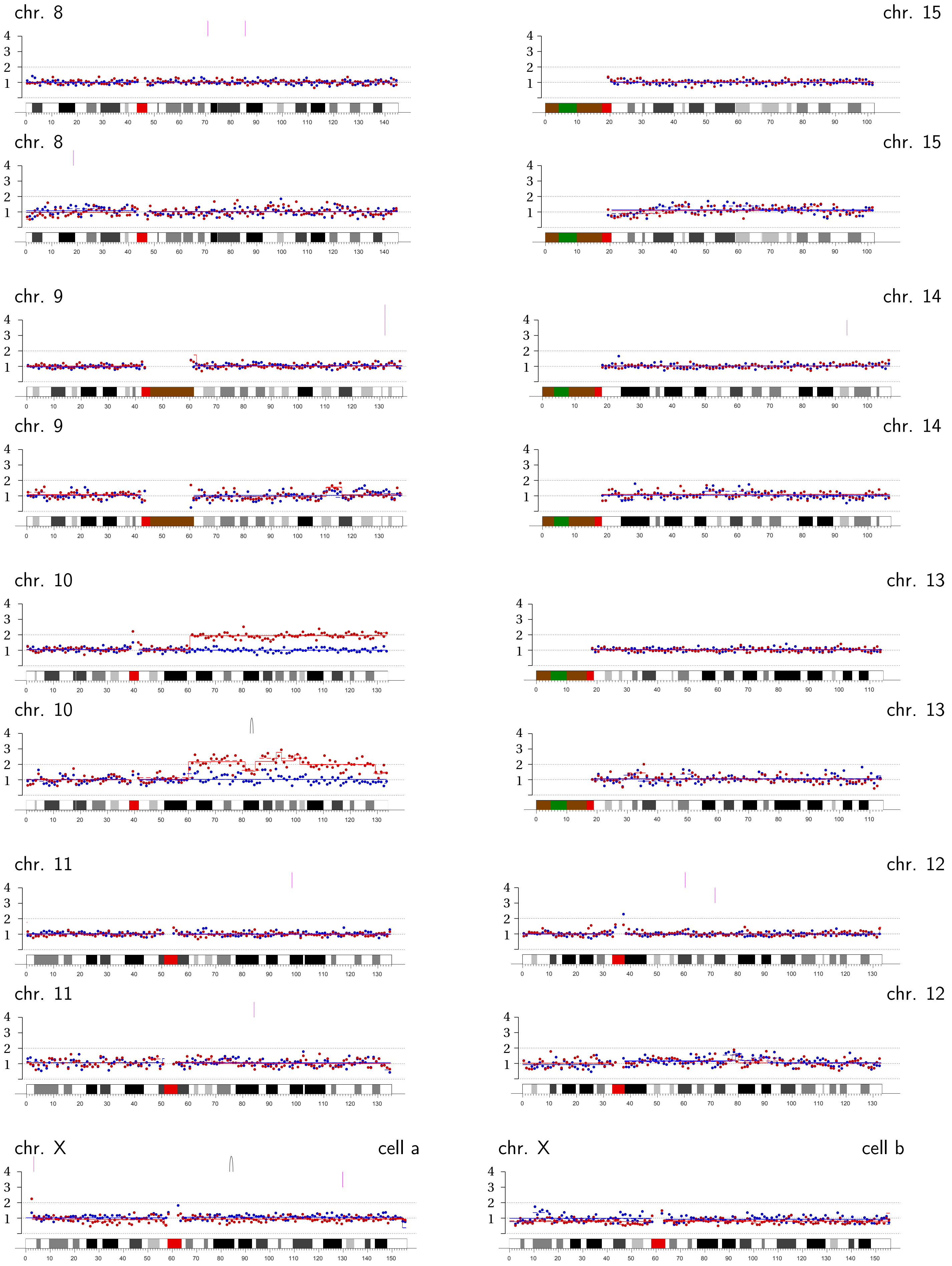
chr. 8
chr. 15
chr. 8
chr. 15
chr. 9
chr. 14
chr. 9
chr. 14
chr. 10
chr. 13
chr. 10
chr. 13
chr. 11
chr. 12
chr. 11
chr. 12
chr. X
cell a
chr. X
cell b

**Micronucleus Pair MN10 | Complex rearrangement on chromosome 1B q-arm (2:0).**

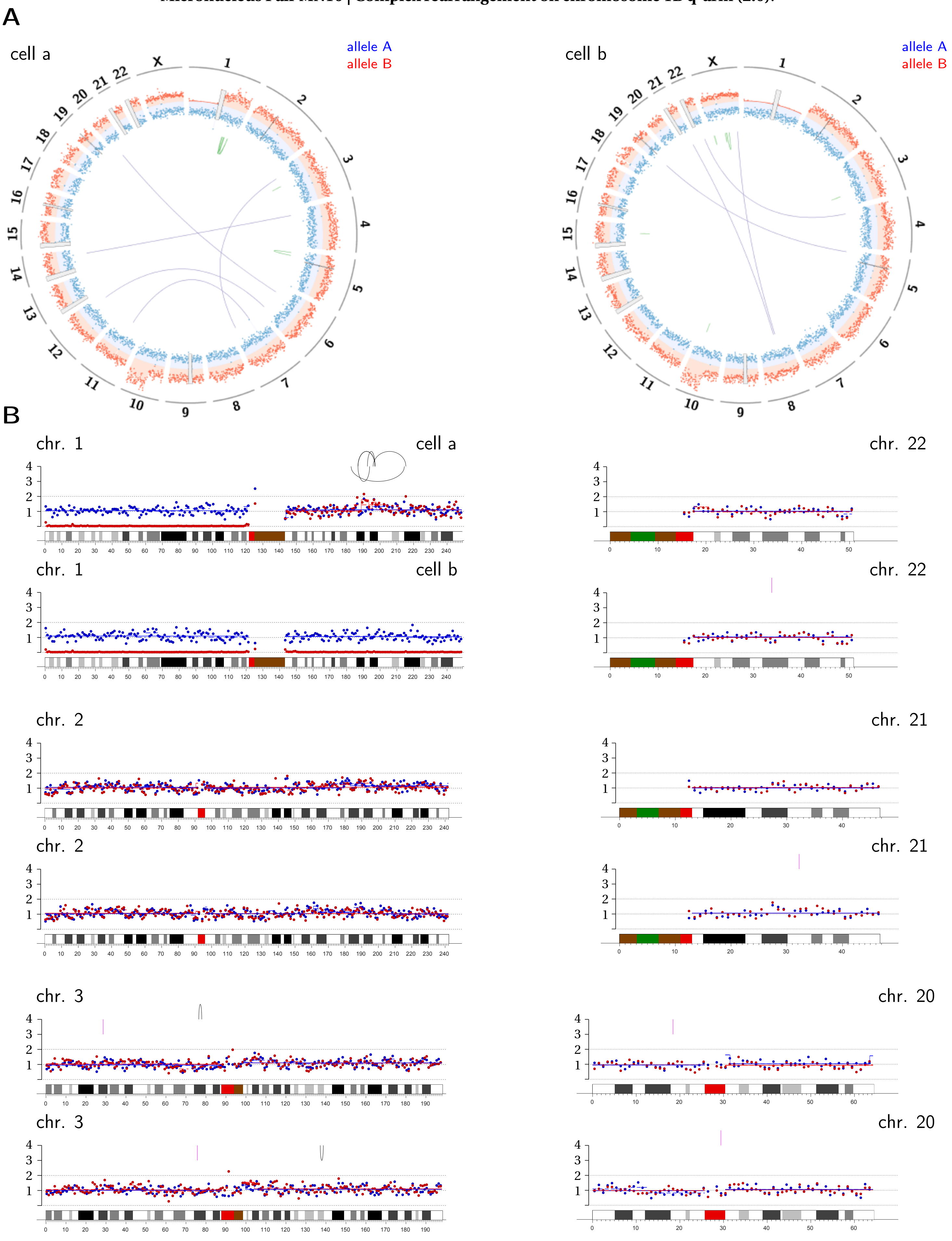

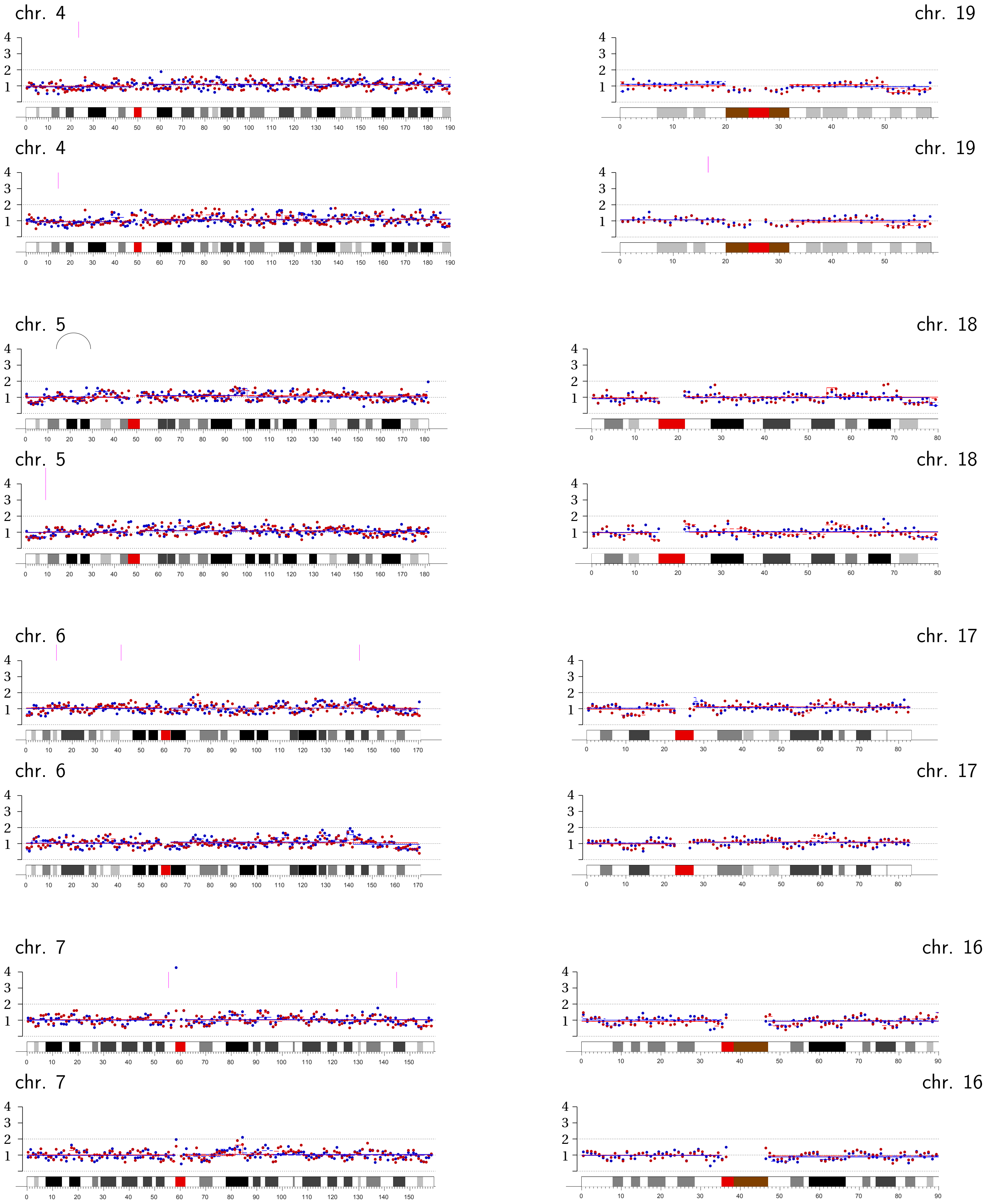
chr. 4
chr. 19
chr. 4
chr. 19
chr. 5
chr. 18
chr. 5
chr. 18
chr. 6
chr. 17
chr. 6
chr. 17
chr. 7
chr. 16
chr. 7
chr. 16

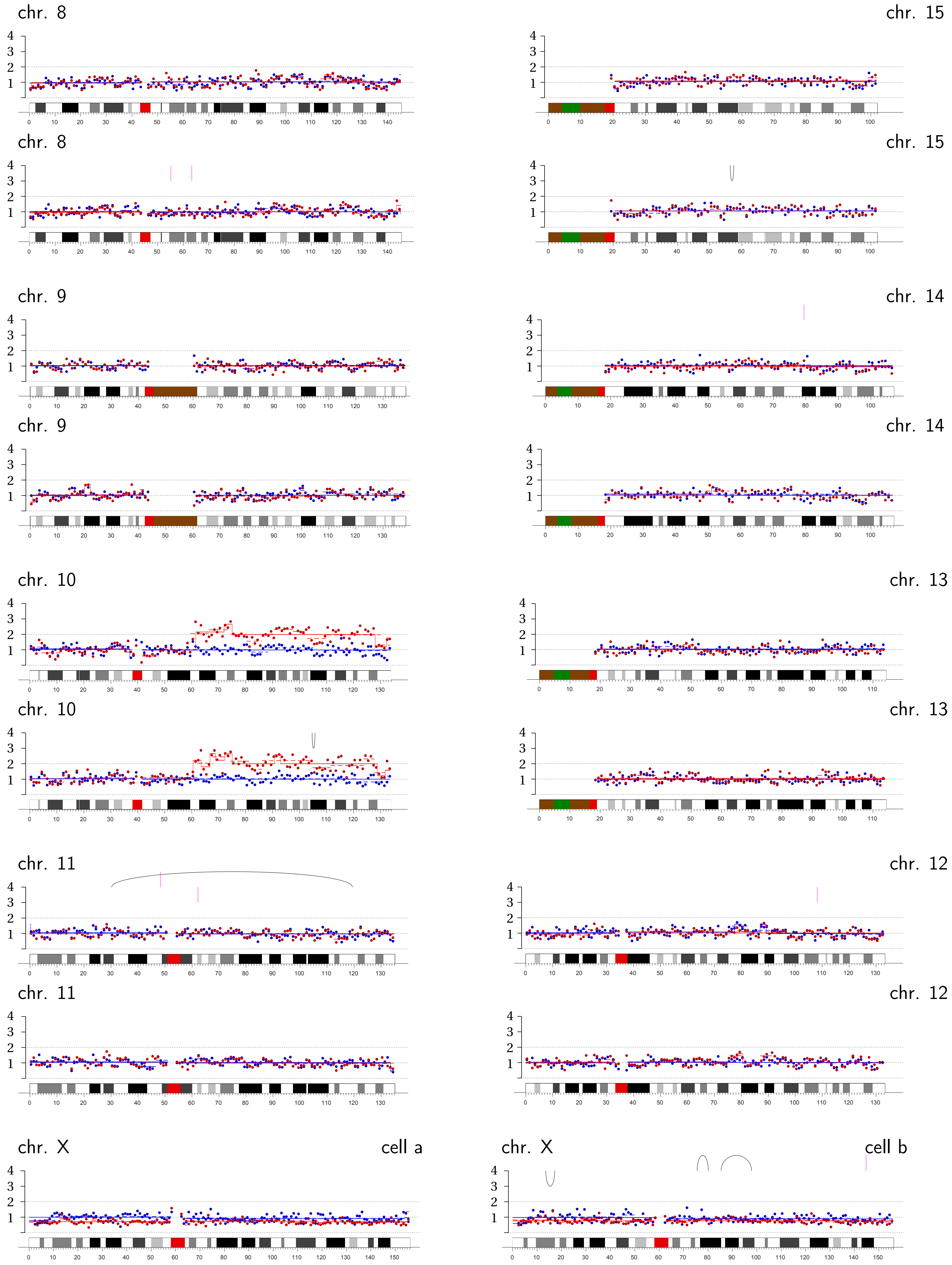
chr. 8
chr. 15
chr. 8
chr. 15
chr. 9
chr. 14
chr. 9
chr. 14
chr. 10
chr. 13
chr. 10
chr. 13
chr. 11
chr. 12
chr. 11
chr. 12
chr. X
cell a
chr. X
cell b

Supplemental Figure 16

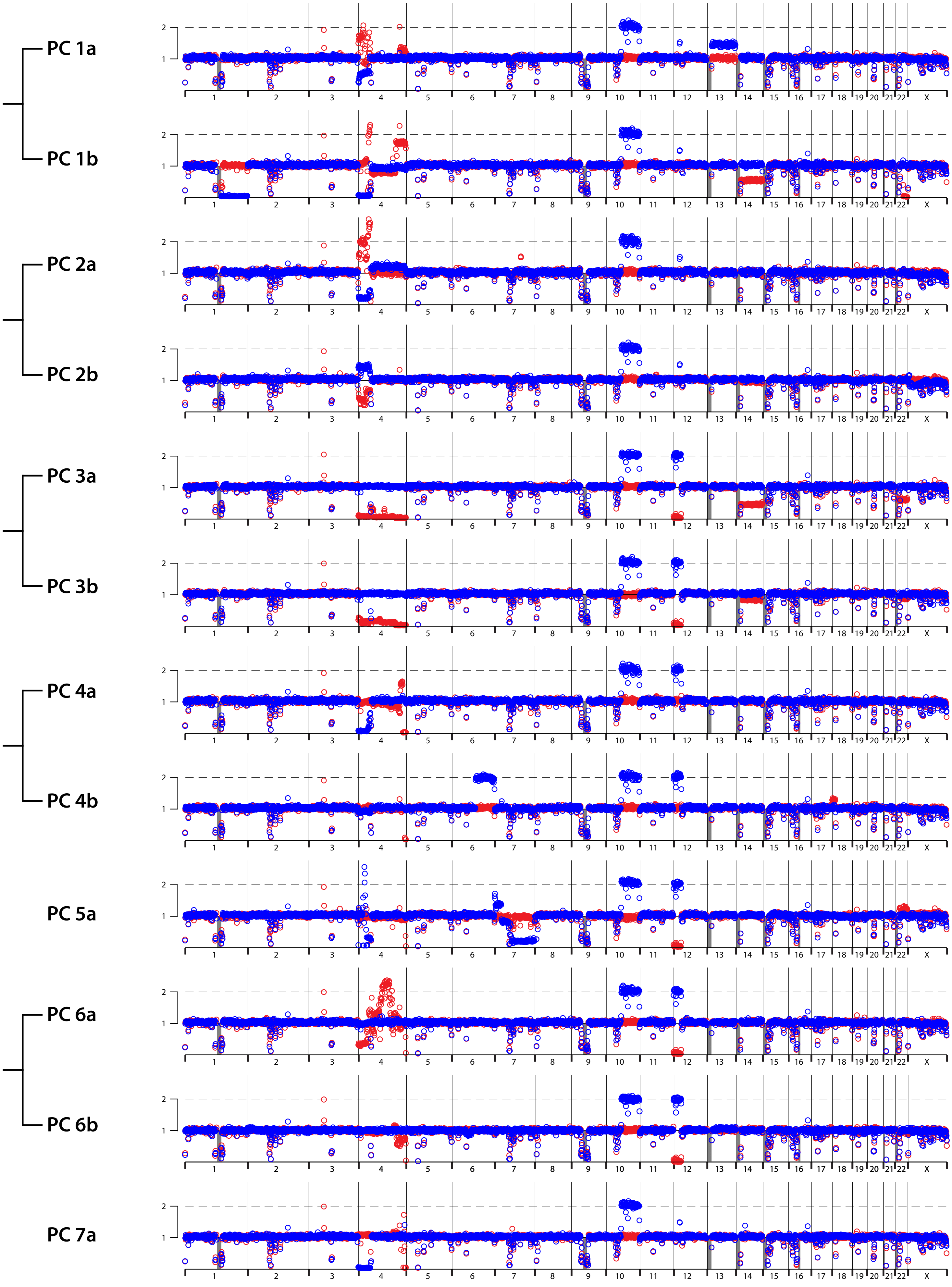

Supplemental Figure 17

**A** **Primary Clone 1a**

45~46,X,add(X)(q28),add(4)(p16),?psu dic(4)(4;14)(p11;p11.2),i(13)(q10)[cp5]

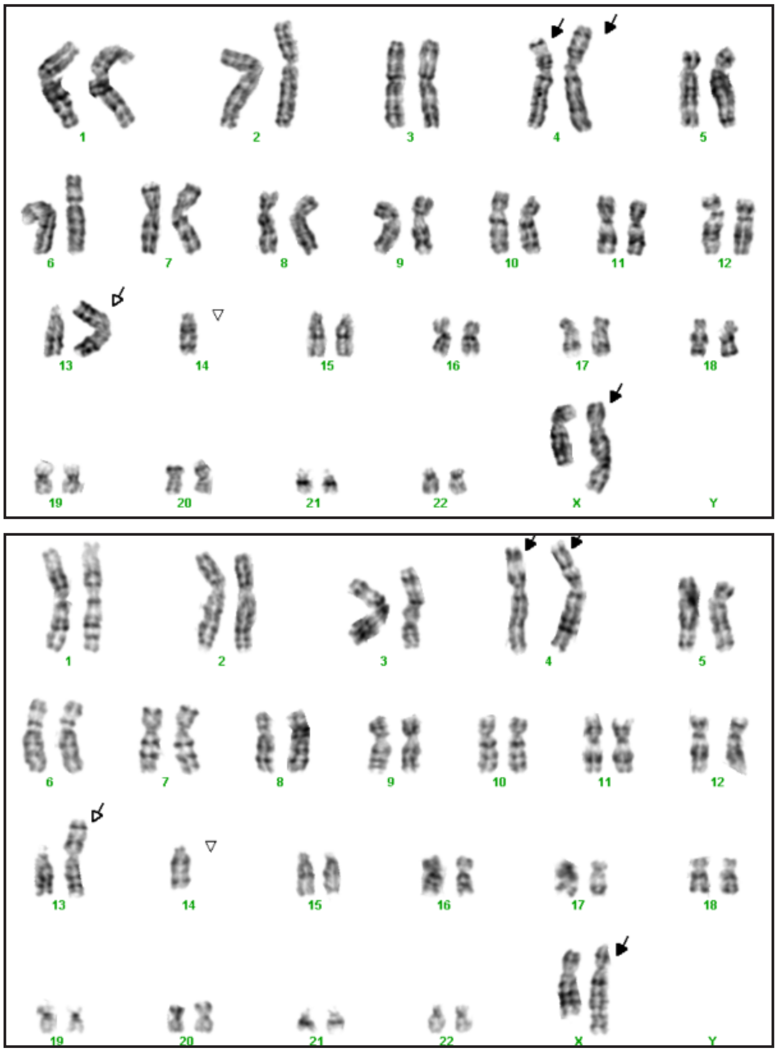

**B** **Primary Clone 6b**

44~46,X,add(X)(q28),add(4)(q35),-22,add(22)(p11.2)[cp7]

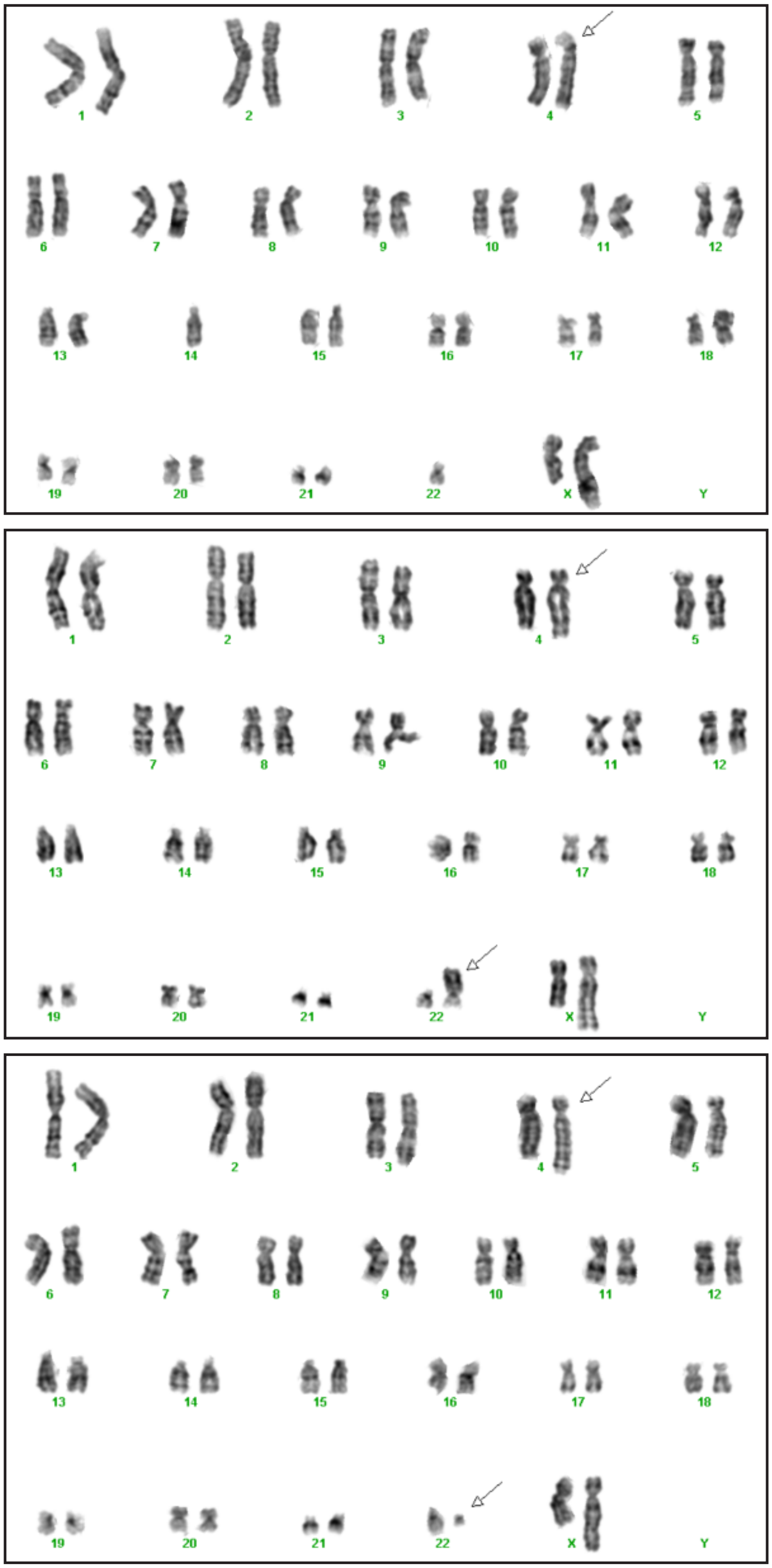

### Supplemental Figure 18

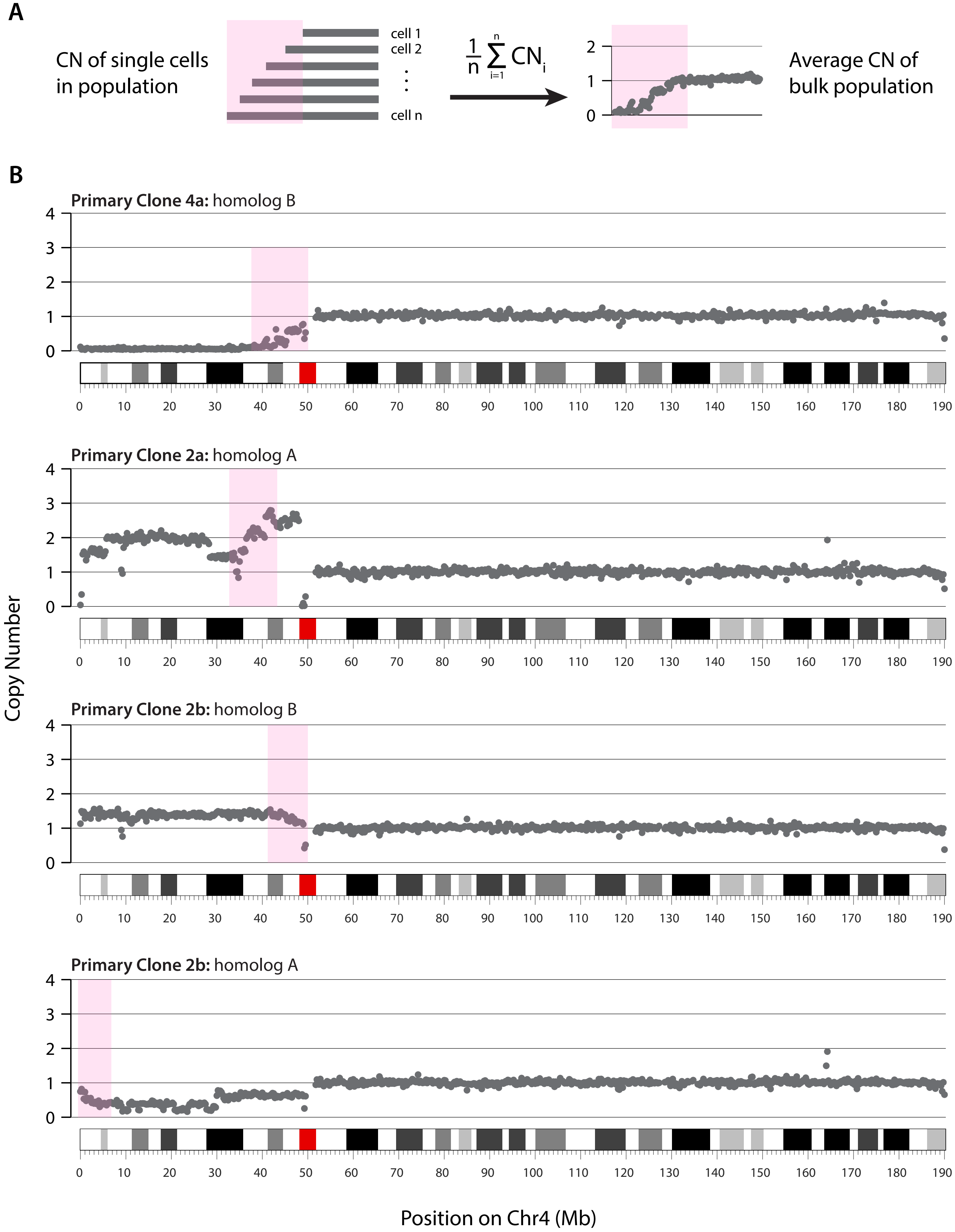

### Supplemental Figure 19

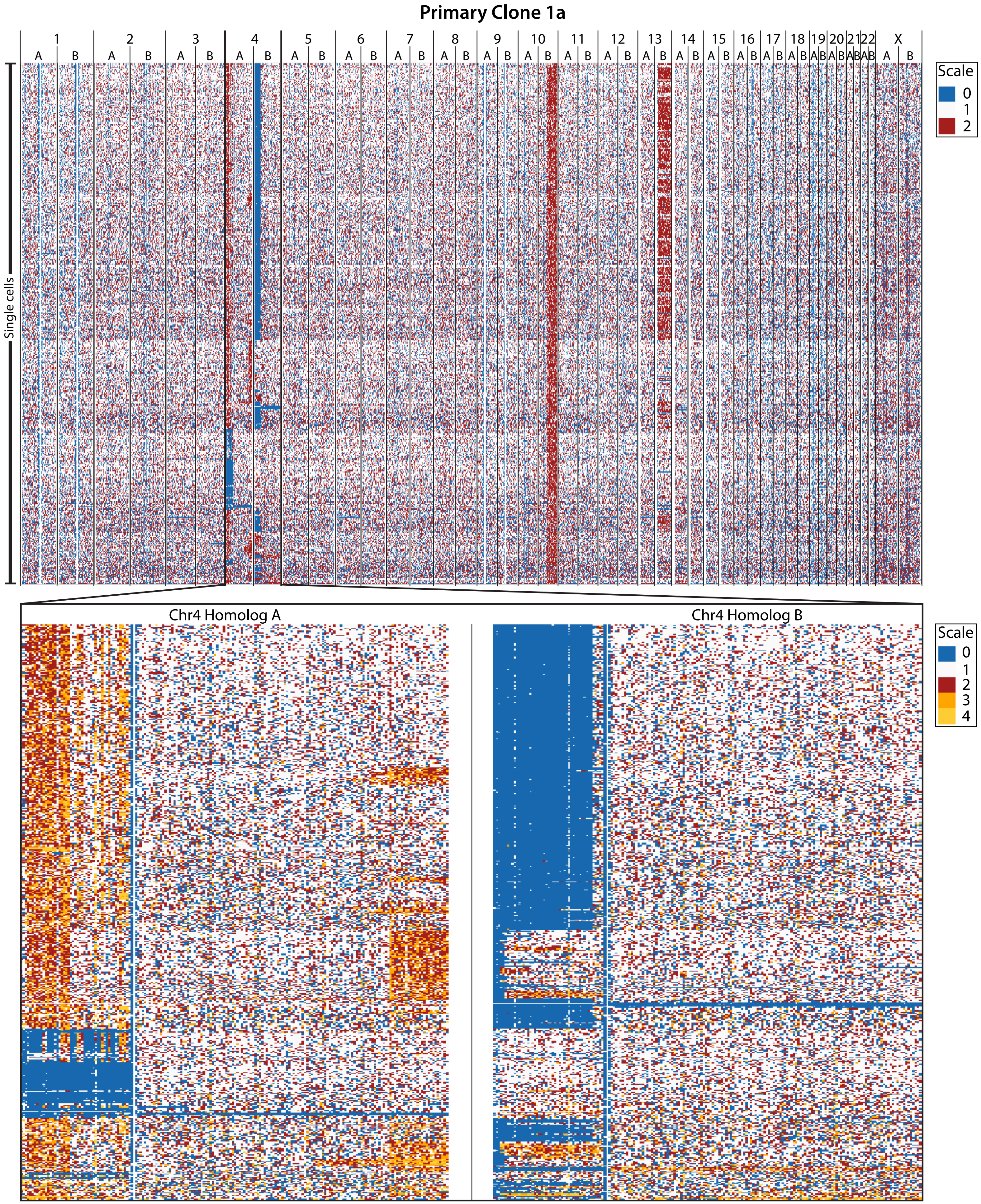

Supplemental Figure 19

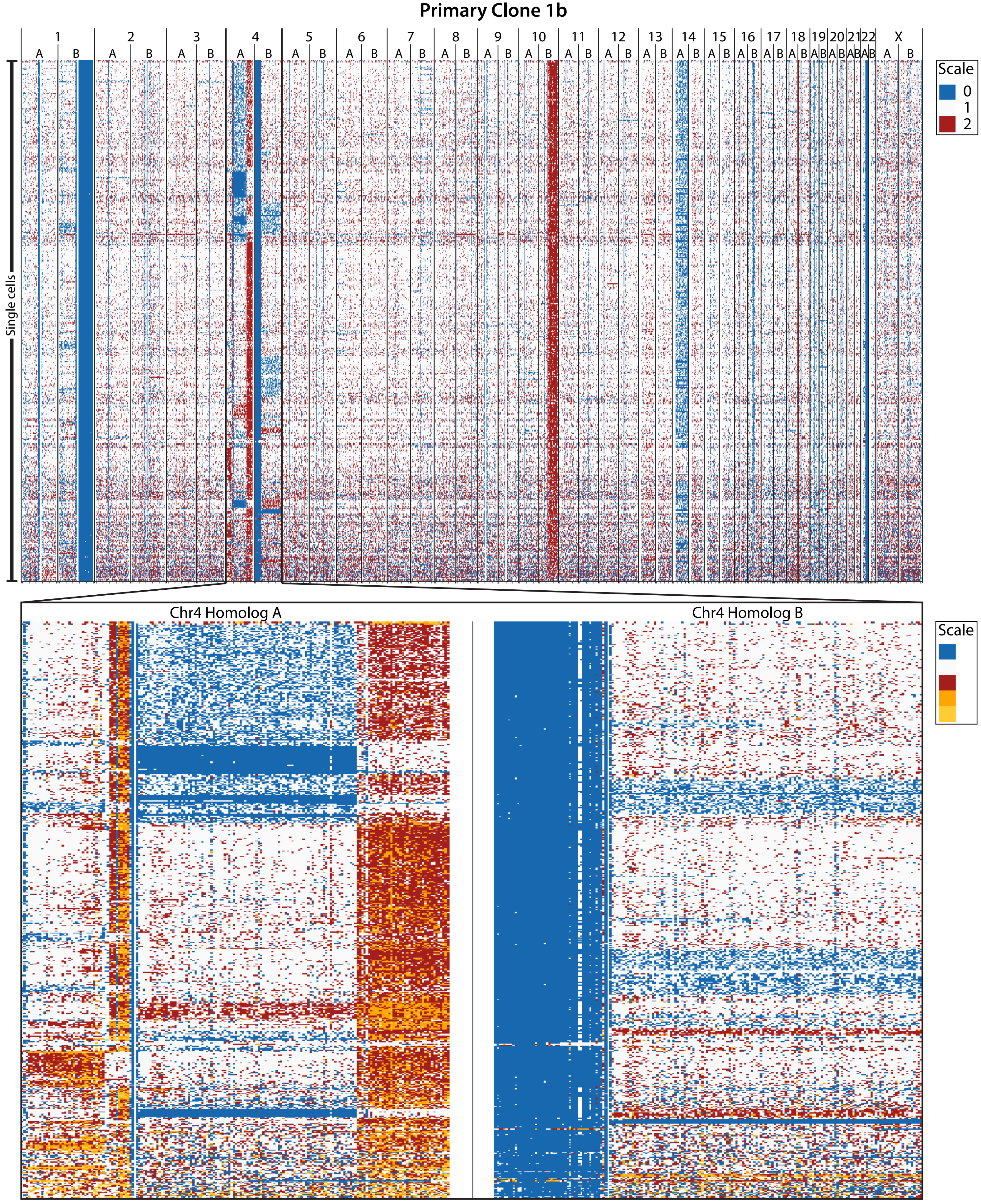

Supplemental Figure 19

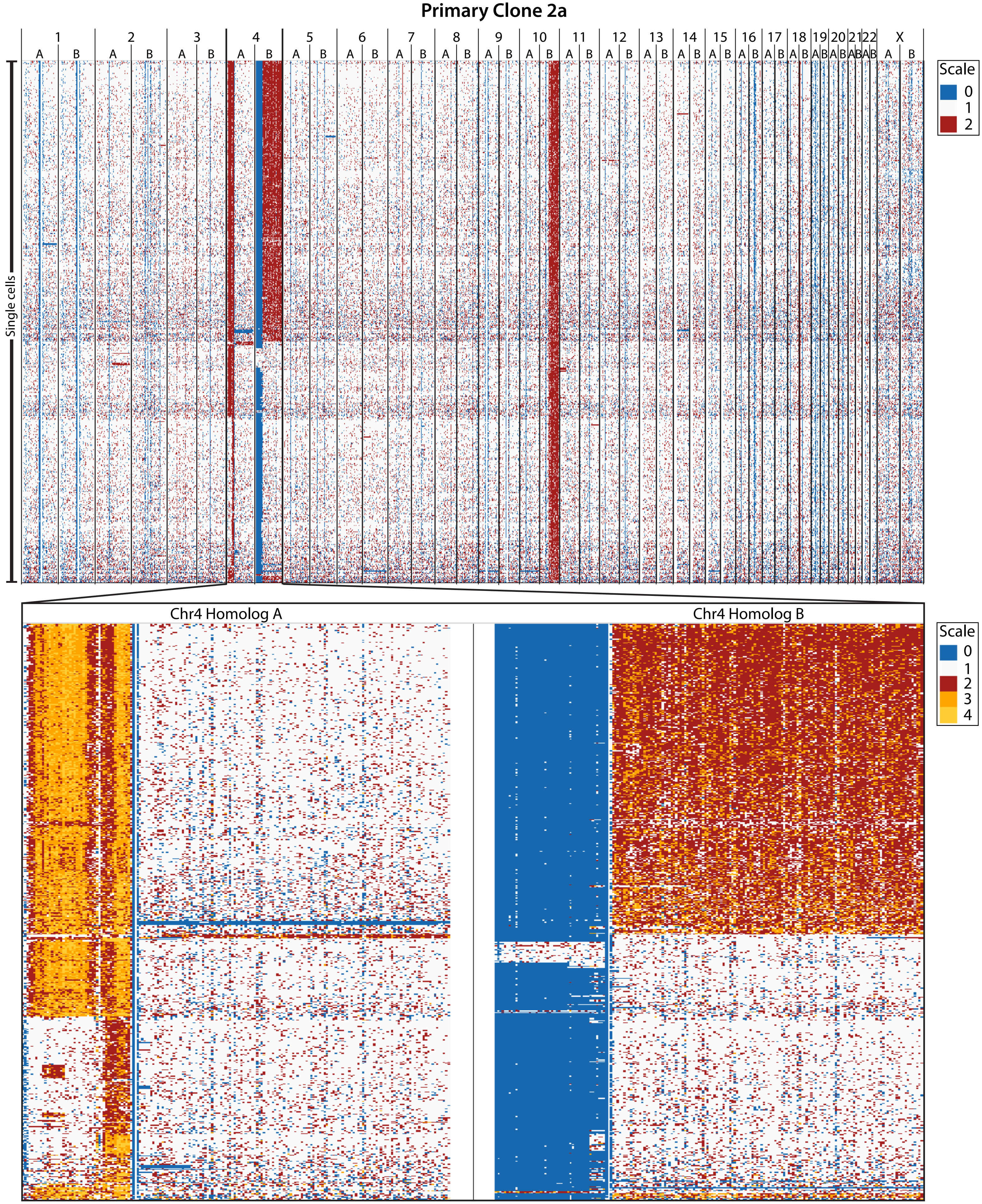

### Supplemental Figure 19

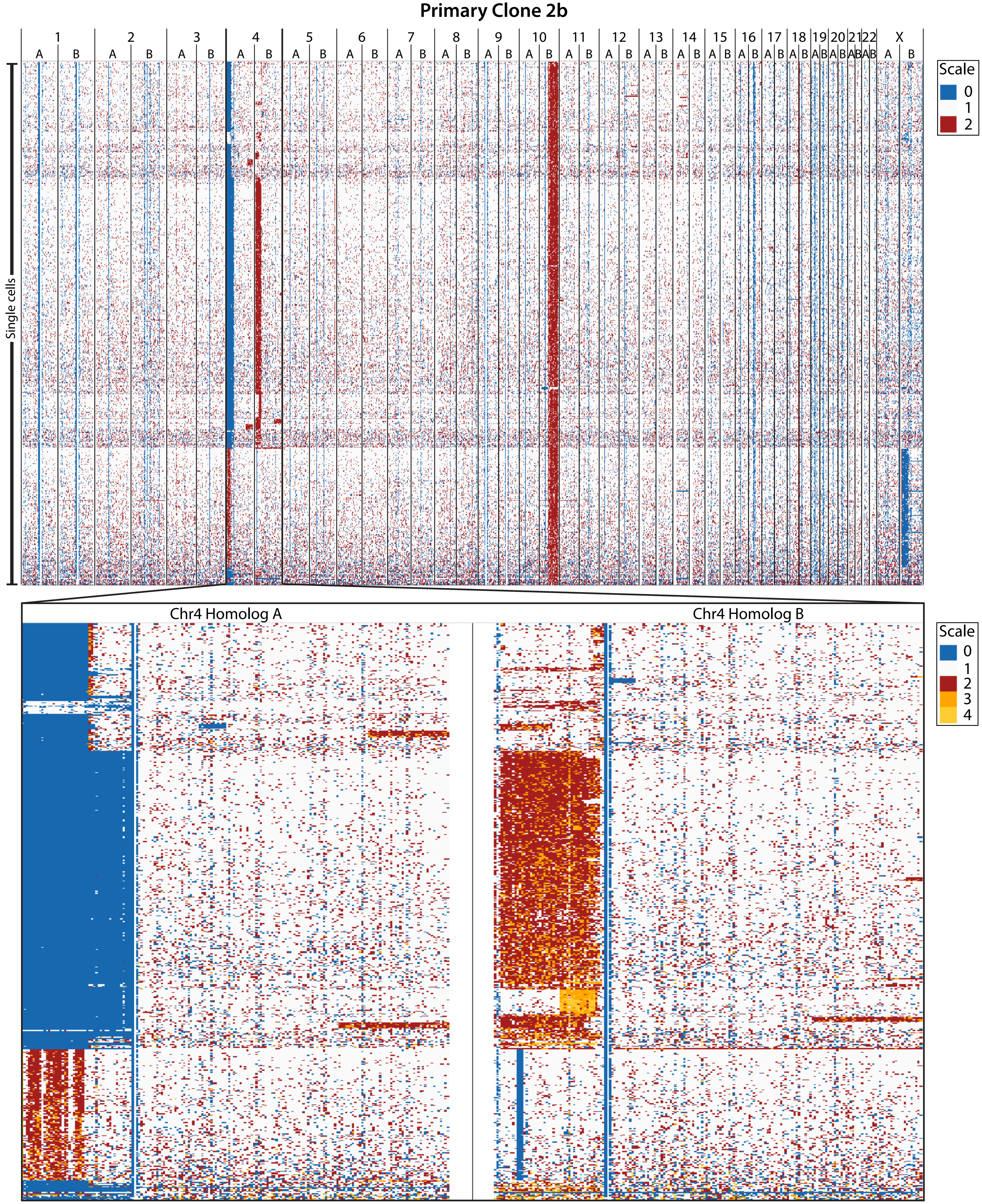

Supplemental Figure 19

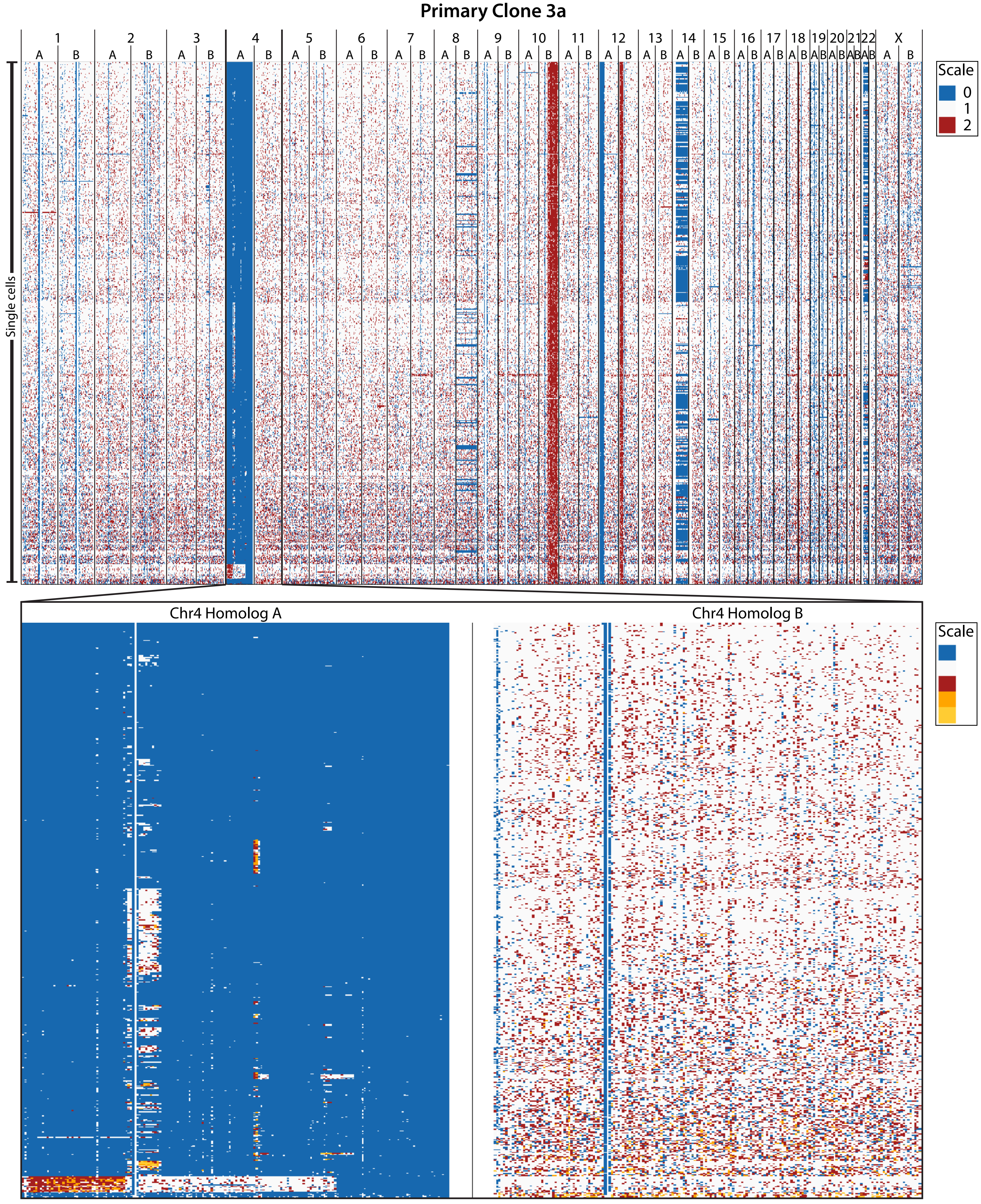

Supplemental Figure 19

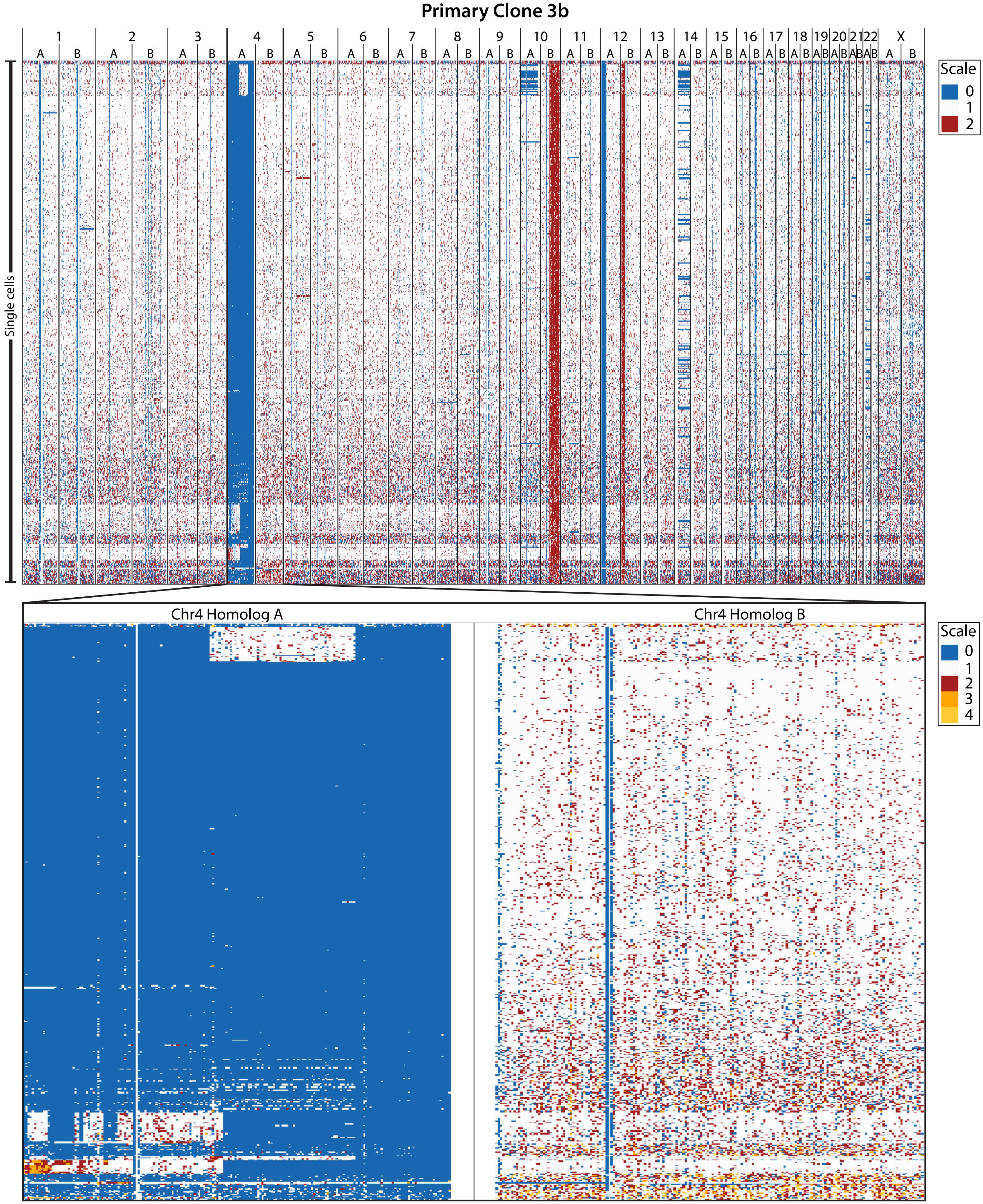

### Supplemental Figure 19

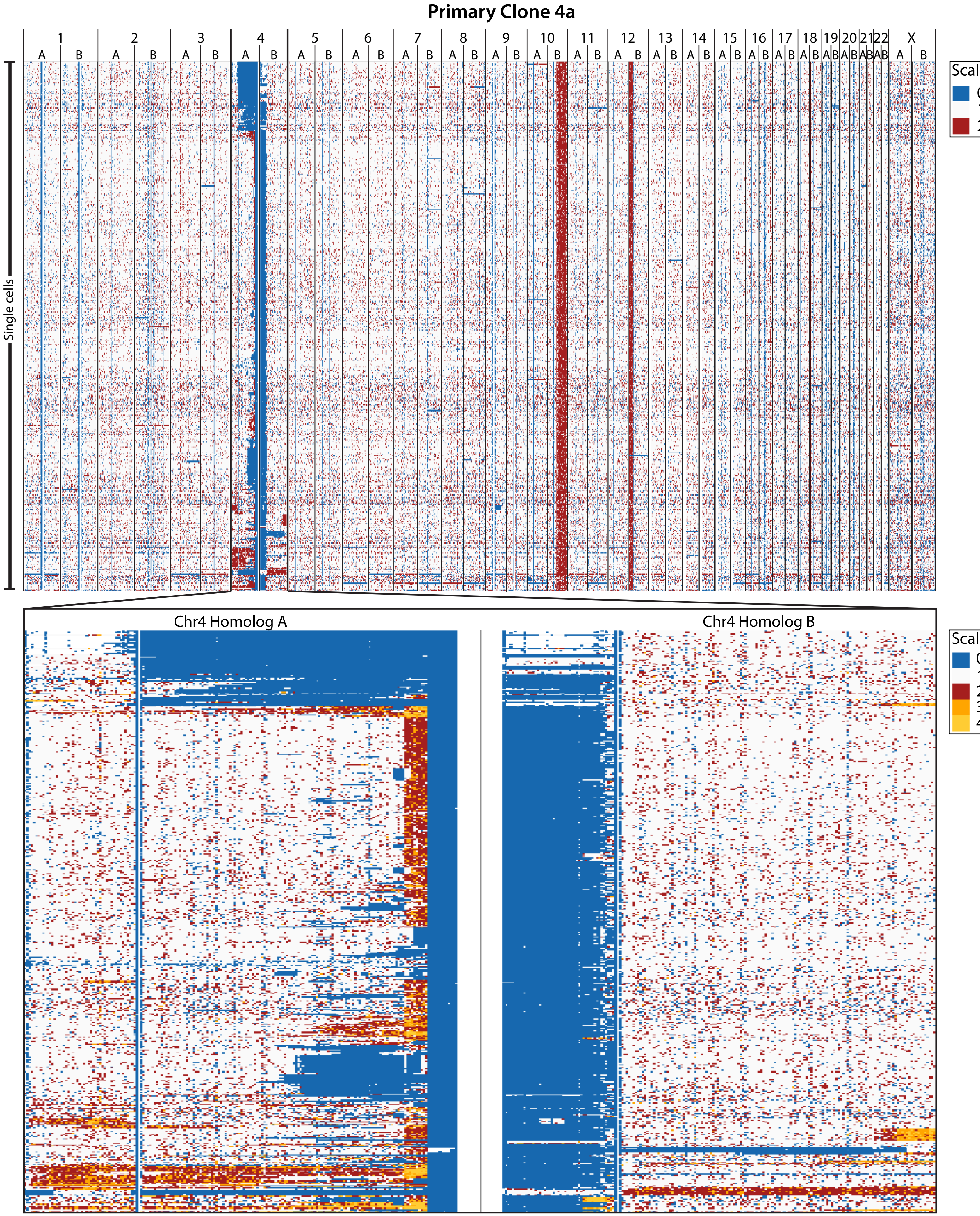

Supplemental Figure 19

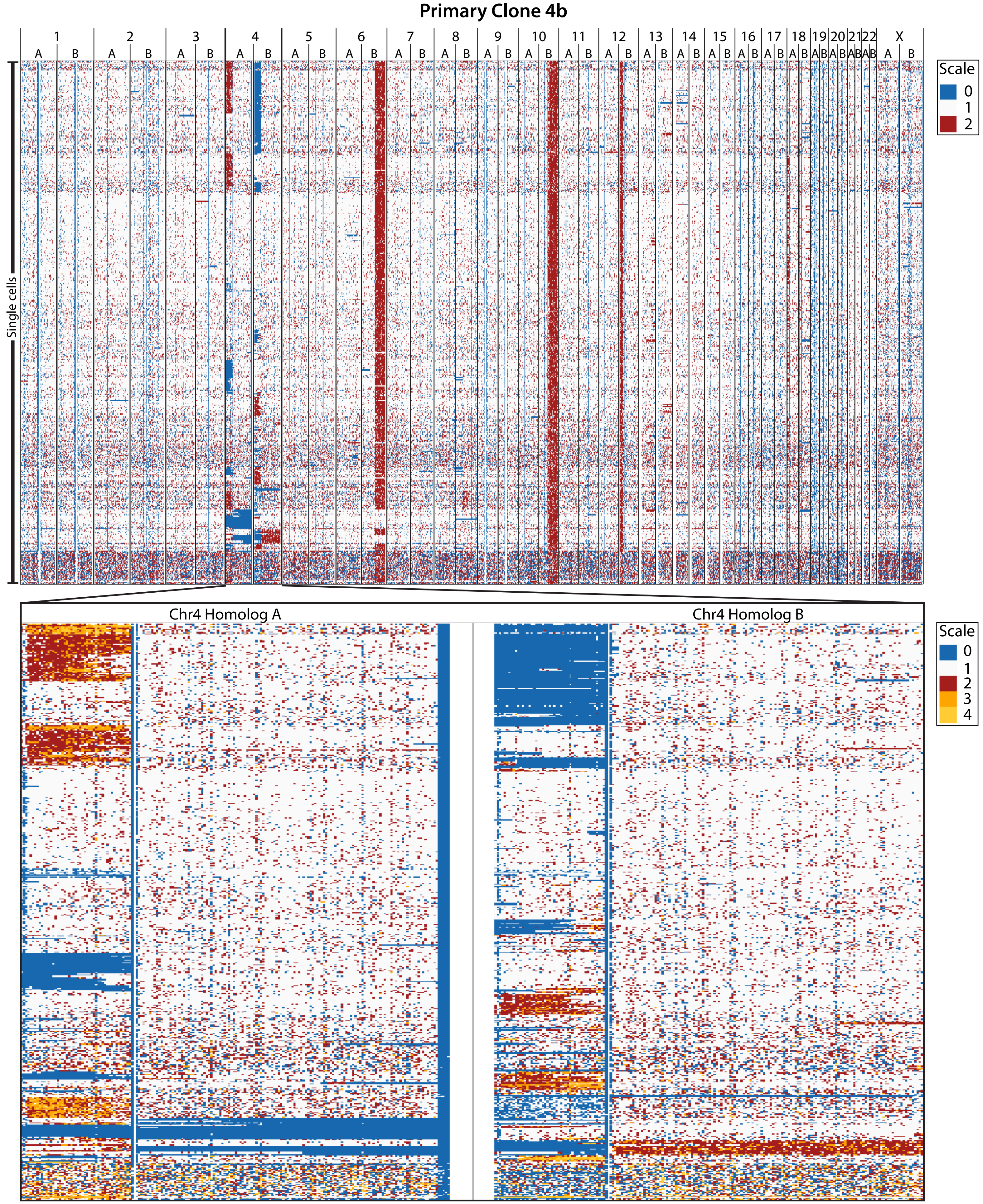

### Supplemental Figure 19

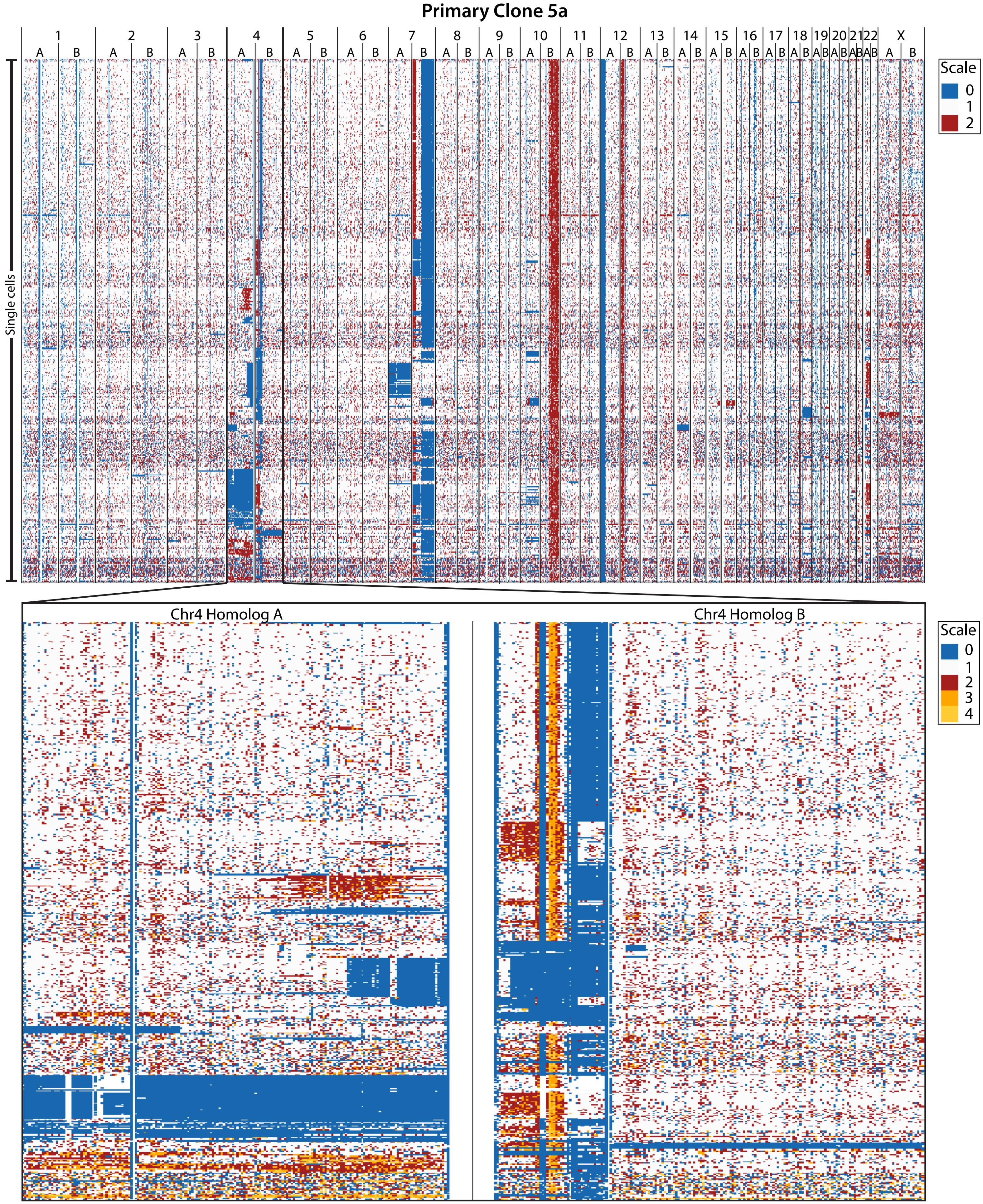

### Supplemental Figure 20

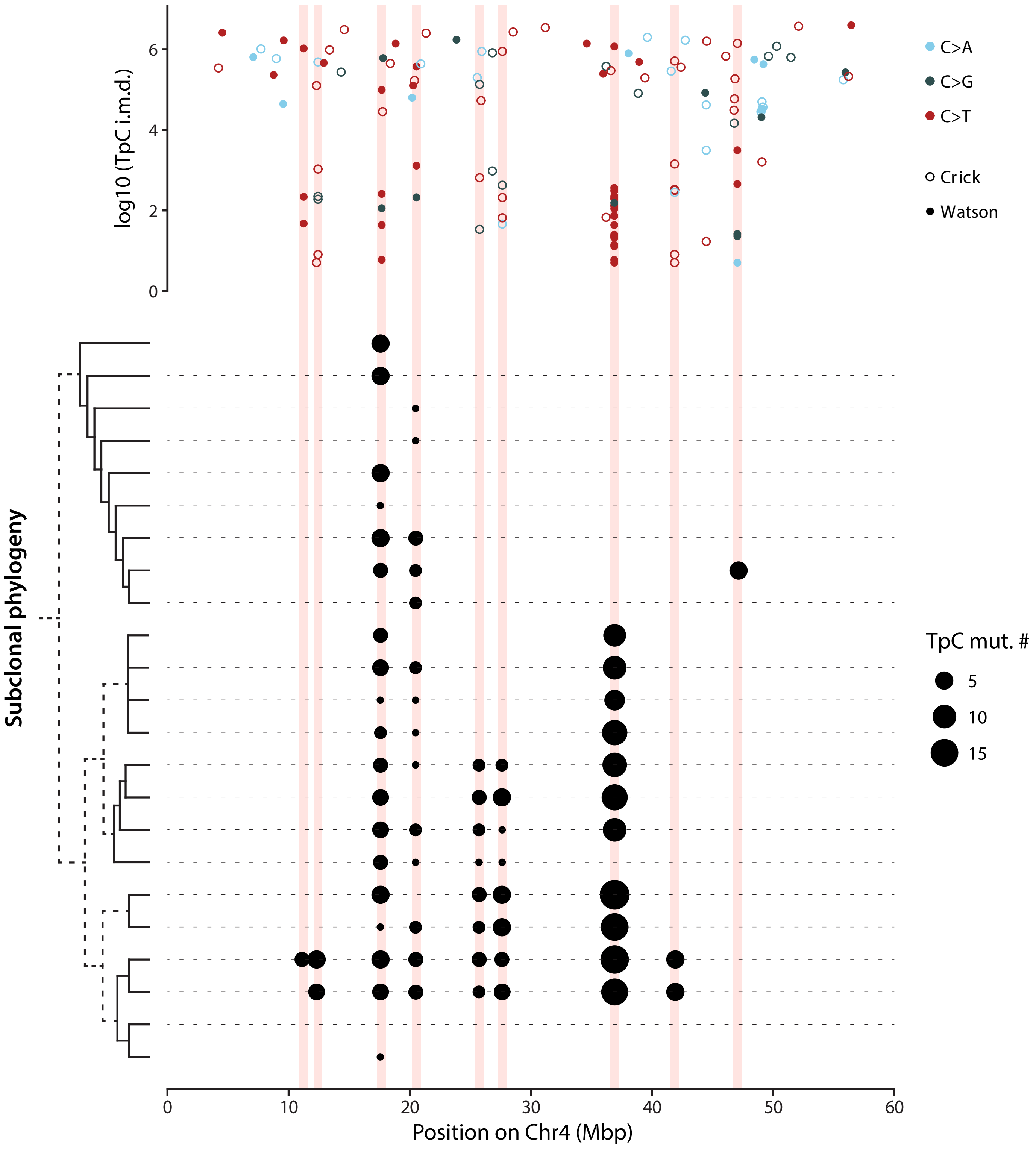

Supplemental Figure 21

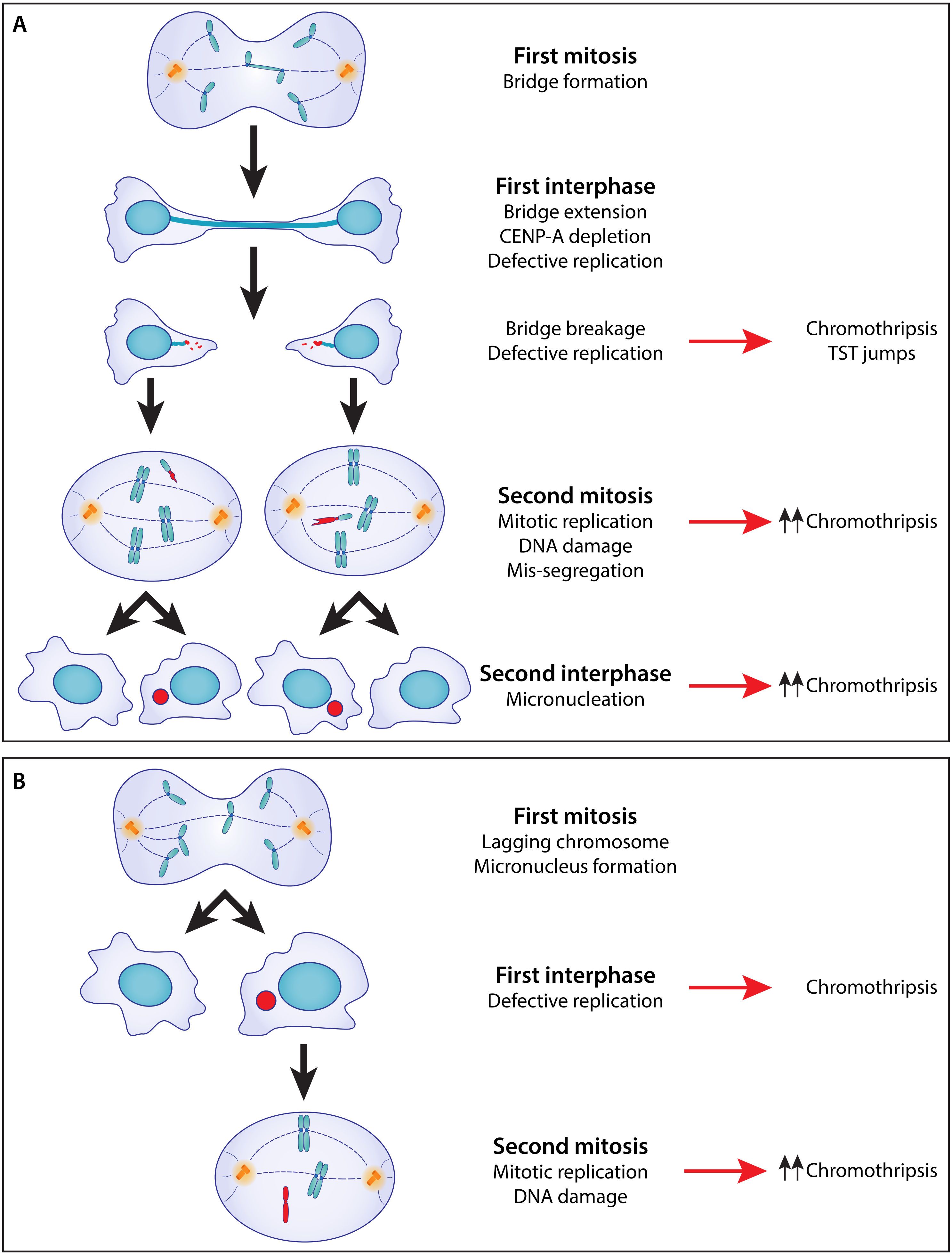
